# Tryptoline Stereoprobe Elaboration Identifies Inhibitors of the GRPEL1-HSPA9 Chaperone Complex

**DOI:** 10.1101/2025.10.20.683548

**Authors:** Rachel E. Hayward, Raymond F. Berkeley, Zijian Gao, Maximilian Garhammer, Marc A. Morizono, Evert Njomen, Haoxin Li, Kristen E. DeMeester, Victor Cociorva, Mark A. Herzik, F. Ulrich Hartl, Bruno Melillo, Benjamin F. Cravatt

**Affiliations:** Department of Chemistry, Scripps Research, La Jolla, CA, USA; Department of Chemistry, University of Oxford, Oxford, UK; Department of Cellular Biochemistry, Max Planck Institute of Biochemistry, Martinsried, Germany; Department of Chemistry and Biochemistry, University of California, San Diego, La Jolla, CA, USA

## Abstract

Activity-based protein profiling has identified hundreds of proteins from diverse classes that react at specific cysteine residues with stereochemically defined electrophilic compounds (stereoprobes) in human cells. The structure-activity relationships underlying these stereoprobe-protein interactions, however, remain poorly understood. Here we show that the protein interaction landscape of tryptoline acrylamide stereoprobes can be profoundly altered by structural modifications distal to the acrylamide reactive group. The majority of stereoprobe liganding events occurred at non-orthosteric sites and mostly evaded assignment by the machine learning-based co-folding model Boltz-2, which instead tended to misplace the stereoprobes in orthosteric pockets (an outcome we term “orthostery burnout”). We found that stereoprobes reacting with C124 in the nucleotide exchange factor GRPEL1 disrupt interactions with the mitochondrial HSP70 chaperone HSPA9/mortalin, leading to impairments in mitochondrial protein import and induction of mitophagy. Our results highlight tryptoline acrylamides as a versatile source of covalent ligands targeting non-orthosteric sites on proteins, including tool compounds that perturb the mitochondrial HSP70 chaperone system.

## Introduction

Most proteins in the human proteome lack chemical probes^1–4^. The identification of small-molecule ligands for proteins that are challenging to assay or do not display well-defined binding pockets represents a particular challenge for chemical biology and drug discovery^5^. Many innovative and broadly applicable approaches have emerged to discover ligands for proteins, including fragment-based drug discovery^6–9^, DNA-encoded libraries^10–12^, affinity-selection^13^ and tethering-based^14^ mass spectrometry, and structure-guided computational methods^15,16^. Each of these methods typically studies purified proteins isolated from the cellular environment. Chemical proteomic strategies such as activity-based protein profiling (ABPP) have the advantage of evaluating small molecules for interactions with endogenous proteins in native biological systems^3,17–19^, which can facilitate the discovery of ligands that bind to dynamic and regulated protein domains and complexes that are challenging to reconstitute in purified, recombinant systems^20–23^.

ABPP has been used in combination with focused libraries of electrophilic small molecules to generate covalent ligandability maps of diverse types of human cells, including cancer cell lines^24–26^ and primary immune cells^27^. By balancing features of recognition and reactivity, electrophilic compounds can engage more shallow and dynamic binding pockets on proteins, thereby expanding the scope of non-orthosteric sites that can be targeted by small molecules^18,19,28–30^. These irreversible small molecule-protein interactions may also improve selectivity by targeting isotype-restricted nucleophilic residues within sets of related proteins^23,31^ and produce extended pharmacological effects that are maintained until new protein target synthesis occurs in cells^3,18,19,28–30,32–34^. ABPP studies performed to date have identified first-in-class covalent ligands that perturb historically challenging protein types, including adaptors^21,35^, RNA-binding proteins^35,36^, and transcription factors^22,37,38^.

While ABPP is a high-content method that can evaluate thousands of proteins in parallel for interactions with small molecules, it has limited throughput owing to the constraints of mass spectrometry (MS)-based analysis. The design of electrophilic compound libraries that maximize the structural diversity presented to the proteome is accordingly a crucial feature of ABPP efforts aimed at broadly interrogating the ligandability of biological systems. ABPP experiments leveraging electrophilic fragments that target cysteine residues have provided initial deep covalent ligandability maps of human cells^24–26,39^. However, these fragment-cysteine interactions generally displayed low potency and selectivity, which complicates studies of their functional effects in cells. More recently, we have designed more elaborated libraries of electrophilic compounds for ABPP that are constructed around diverse conformationally constrained and sp^3^-rich cores capable of displaying variable recognition and reactive groups, as well as latent (e.g., alkyne) affinity handles^21,27,35,40,40–42^. These libraries are also synthesized in stereopure form such that biological systems are exposed to physicochemically matched pairs of enantiomeric compounds (stereoprobes), and ABPP results can then be interpreted for stereoselective small molecule-protein interactions, a hallmark of specific molecular recognition events^43^. ABPP experiments performed with stereoprobes further facilitate downstream functional studies by providing pairs of active and inactive enantiomeric compounds for each liganding event of interest.

Among the different classes of stereoprobes investigated by ABPP, the tryptoline acrylamides have shown a remarkably diverse array of interactions with the human proteome. We recently found, for instance, that a focused library of ∼ 28 tryptoline acrylamides tested at low µM concentrations in human cancer cells engaged >300 proteins with high occupancy and stereoselectivity^21^. The tryptoline acrylamides may owe their versatility to a Goldilocks-like balance of reactivity-and recognition-dependent interactions with proteins that is enabled by a scaffold displaying at least 4-5 points for straightforward synthetic diversification. The respective contributions of covalent and non-covalent interactions to defining the scope of proteins engaged by tryptoline acrylamides, however, remains largely unexplored. Indeed, the most common sites of structural diversification studied to date are proximal to the acrylamide reactive group and may therefore simultaneously impact both its intrinsic electrophilicity and steric accessibility^21,22,42,44^. Our previous studies nonetheless provided initial evidence of the importance of recognition groups on the tryptoline core, as we identified several liganding events dependent on an alkyne group located at the C6 position distal to the acrylamide reactive group^21^.

Motivated to more thoroughly understand the contribution of molecular recognition to the covalent ligandability profiles of electrophilic stereoprobes, we report here the design and ABPP analysis of eight sets of tryptoline acrylamides featuring elaborated distal (C6 and C7) ring modifications. Across > 24,000 cysteines quantified in human cancer cells, we identified ∼100 sites on structurally and functionally diverse proteins that were liganded by tryptoline acrylamides, of which 54 showed preferential reactivity with the elaborated analogs. Only a small subset of these liganding events occurred at orthosteric pockets, and these events were predicted with high confidence by the machine learning (ML) co-folding program Boltz-2. In contrast, Boltz-2 mostly failed to predict the much larger category of non-orthosteric liganding events, which were instead often incorrectly localized by the program to orthosteric pockets on the same protein. We validated the stereoselectivity, chemoselectivity, and site-specificity of several non-orthosteric liganding events mediated by elaborated tryptoline acrylamides with recombinant proteins. We further found that one such interaction occurring at C124 of the nucleotide exchange factor GRPEL1 disrupted binding of this protein to the mitochondrial HSP70 chaperone HSPA9 (or mortalin or GRP75), leading to blockade of protein import into the mitochondria, impaired cellular respiration, and activation of the integrated stress response and mitophagy. Our findings illuminate a profound impact of distal structural modifications on the protein interaction profiles of tryptoline acrylamide stereoprobes, enabling the discovery of ligands targeting non-orthosteric sites on a wide array of proteins.

## Results

### Proteome-wide ligandability maps of elaborated tryptoline acrylamides

Previous ABPP studies of tryptoline acrylamides revealed that several cysteine liganding events in human cancer cells strongly depended on the presence of an alkyne handle at the C6 position of the tryptoline core^21^ (**Fig. 1a, b** and **Extended Data Fig. 1a**). These results were somewhat unexpected, given the relatively small size of the propargyl ether appendage and its location at a site remote from the acrylamide reactive group. To further explore the impact of distal core modifications on the proteome reactivity of tryptoline acrylamides, we generated a series of elaborated analogs bearing substituents of varying size, flexibility, and lipophilicity (phenylacetylene, benzodioxolyl, fluorophenylether, and *N*-methyl triazole) positioned at the C6 or C7 positions (**Fig. 1c**). Given their unhindered positions, we expected such appendages to have a minimal impact on the geometry of the tryptoline core itself, which we corroborated *in silico* (**Extended Data Fig. 1b**). We therefore anticipated that the eight sets of stereoprobes would collectively provide insights into not only the stereoselectivity (each set contains all four possible stereoisomers), but also chemoselectivity and regioselectivity of elaborated tryptoline acrylamide-cysteine interactions in human cells.

**Fig. 1.**
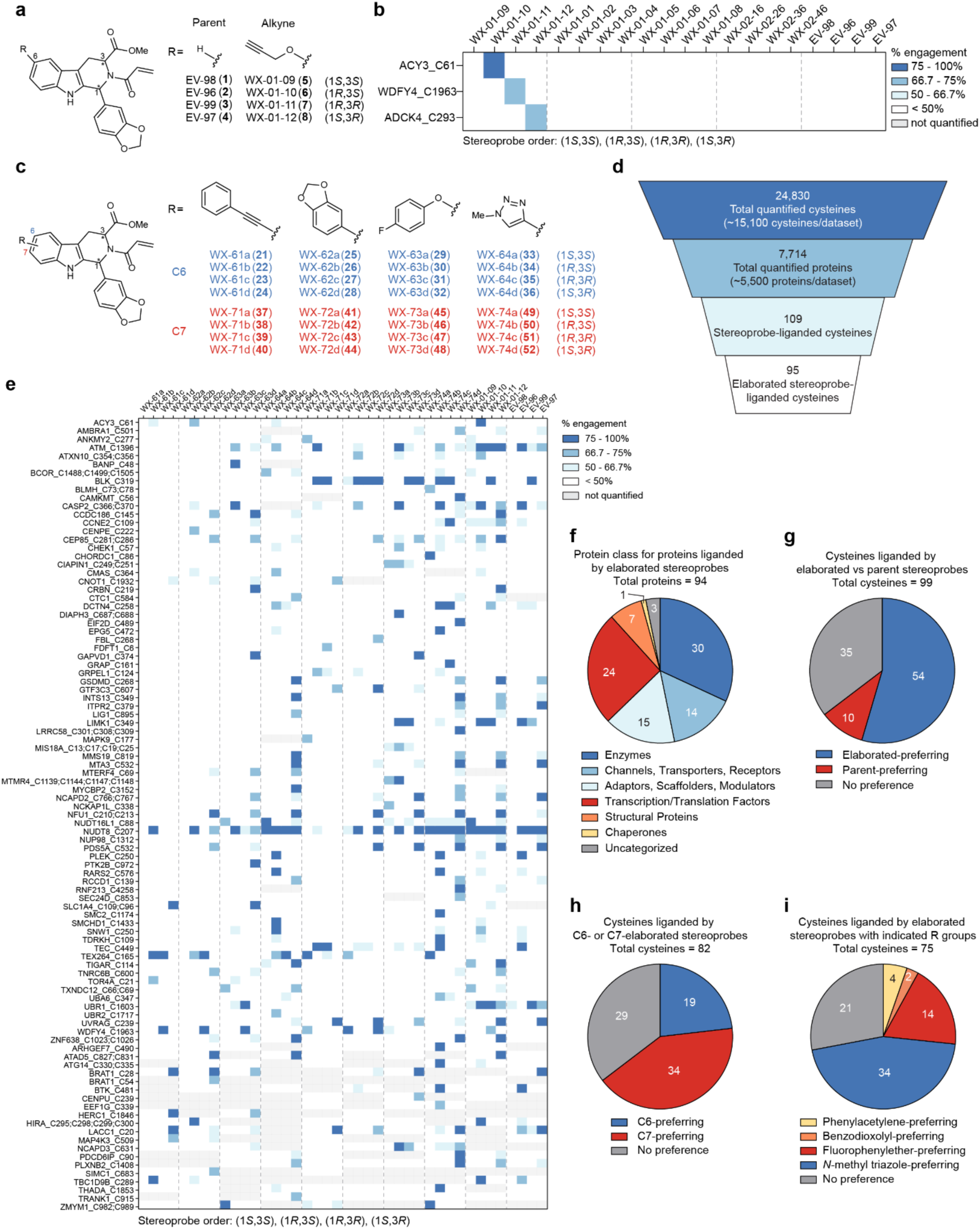
Design and chemical proteomic analysis of elaborated tryptoline acrylamide stereoprobes. **a**, Structures of original parent and alkynylated tryptoline acrylamide stereoprobes^21,27^. **b,** Heatmap showing representative cysteines displaying preferential stereoselective reactivity with C6-alkyne modified tryptoline acrylamide stereoprobes WX-01-10, -11, and -12. Cysteine-directed ABPP data were from Ramos cells, as reported previously^21^, and represent average values from 4 independent experiments per stereoprobe. **c,** Structures of tryptoline acrylamides with elaborations at the C6 (blue) and C7 (red) positions. **d,** Total numbers of proteins/cysteines, and their stereoselectively liganded subsets, quantified by cysteine-directed ABPP for parent and/or elaborated stereoprobes in Ramos cells. **e,** Heatmap showing the 95 cysteines liganded by the elaborated stereoprobes compared to alkyne and parent stereoprobes. Cysteines separated by a semi-colon (;) are located on the same tryptic peptide. Cysteines are sorted alphabetically then by number of experiments quantified (9-10 experiments and 1-8 experiments; also see **Extended Data Fig. 1d**). Cysteine-directed ABPP data represent average values from 4-6 independent experiments per stereoprobe. **f,** Functional class distribution of elaborated stereoprobe-liganded proteins assigned as described previously using GO (Panther) and Uniprot annotations^21,27^. **g,** Pie chart showing number of stereoselectively liganded cysteines preferentially engaged (> 1.5-fold IA-DTB blockade) by elaborated versus parent stereoprobes of the same stereoconfiguration. Cysteines not quantified in either elaborated or parent stereoprobe datasets were excluded from the analysis (10 total cysteines). **h,** Pie chart showing number of stereoselectively liganded cysteines preferentially engaged (> 1.5-fold IA-DTB blockade) by C6-elaborated versus C7-elaborated stereoprobes with the same R group and stereoconfiguration. Cysteines not quantified in either C6-and C7-elaborated stereoprobe datasets of the same R group and stereoconfiguration were excluded from the analysis (13 total cysteines). **i,** Pie chart showing number of stereoselectively liganded cysteines preferentially engaged by a specific elaborated chemotype (R group) as determined by comparing data for stereoprobes having different R groups at the same position (C6 or C7) and of the same stereoconfiguration (> 1.5-fold IA-DTB blockade by the reference stereoprobe compared to the median IA-DTB blockade by the three R-group analogs). Cysteines not quantified in all datasets required for comparison were excluded from this analysis (20 total cysteines).

We performed cysteine-directed ABPP experiments in the Ramos human B lymphoid cancer line, which we previously evaluated with C6-alkyne-modified tryptoline acrylamide stereoprobes^21^ (WX-01-09/10/11/12; **Fig. 1a**). Ramos cells were treated with elaborated stereoprobes (C6-modified: WX-61a – WX-64d (**21-36**); C7-modified: WX-71a – 74d (**37**-**52**); **Fig 1c**; 20 µM each), as well as unelaborated parent stereoprobes^27^ (EV-96-99; **Fig. 1a**; 20 µM each), for 3 h and analyzed by cysteine-directed ABPP as described previously^21^ (**Extended Data Fig. 1c**). In this approach, electrophilic compounds are broadly assessed for the blockade of cysteine reactivity with an iodoacetamide-desthiobiotin (IA-DTB) probe. Across >24,000 cysteines quantified in total (an average of ∼15,100 cysteines per ABPP dataset) from >7,000 proteins, we assigned cysteines as being stereoselectively liganded if i) they showed a > 66.7% decrease in IA-DTB reactivity in cells treated with a stereoprobe, and ii) this decrease was at least 2.5-fold greater than that observed for the enantiomer of the stereoprobe^21^. Stereoprobe liganding events associated with tryptic peptides containing more than one cysteine were assigned as potentially occurring on any of these cysteines, but were considered a single liganding event for further analyses. Based on these criteria, 95 cysteines were identified as liganded by the elaborated stereoprobes (**Fig. 1d, e** and **Supplementary Dataset 1**). These liganded cysteines were found in structurally and functionally diverse proteins (**Fig. 1f**), and the vast majority (∼80%) were quantified in at least one replicate ABPP experiment performed with each of the 10 stereoprobe sets (**Extended Data Fig. 1d**), which facilitated the analysis of structure-activity relationships (SAR).

Of the 95 cysteines liganded by the elaborated stereoprobes, 54 were preferentially engaged (> 1.5-fold) in comparison to the parent stereoprobes (**Fig. 1g**). Far fewer cysteines (10 in total) showed preferential engagement by the parent stereoprobes (**Fig. 1g**), with the remaining cysteines being similarly engaged by both elaborated and parent stereoprobes (**Fig. 1g**). Exemplary ligandability profiles for cysteines in each category are shown in **Extended Data Fig. 1e**. We observed a similar breakdown of liganding preferences in comparisons of elaborated vs C6-alkyne stereoprobes (WX-01-09/10/11/12), although with a somewhat higher number of shared cysteine liganding events for these two classes of tryptoline acrylamides (**Extended Data Fig. 1f**). Within the elaborated stereoprobe set, we also observed clear SAR, as most of the cysteines liganded by these compounds showed a strong preference for modifications at either the C6 or C7 position (**Fig. 1h**), and, even within each position, most of the liganded events preferred a single R-group structural elaboration (**Fig. 1i** and **Extended Data Fig. 1g**). As was found previously and consistent with differences in intrinsic electrophilicity^21^, the trans stereoisomers ((1*S*,3*R*) and (1*R*,3*S*)) generally engaged more cysteines than the cis stereoisomers ((1*R*,3*R*) and (1*S*,3*S*)), although all four stereochemical configurations nonetheless liganded distinct sets of cysteines (**Extended Data Fig. 1g**).

Taken together, our cysteine-directed ABPP data indicated that C6 and C7 core elaborations exerted a strong influence on the ligandability maps of tryptoline acrylamide stereoprobes, leading to the loss of some cysteine interactions while enhancing many others. We considered the stringency of the SAR of these stereoprobe-cysteine interactions, which reflected, in most instances, stereoselectivity, regioselectivity, and chemoselectivity, as an opportunity to evaluate their predictability by modern machine learning (ML)-based co-folding methods.

### Boltz-2 analysis of tryptoline acrylamide-liganded proteins

Recent advances in ML have led to the introduction of tools for predicting the three-dimensional structure of biomacromolecules and their complexes^45–50^. The accuracy, scalability, and ease of use of ML-enabled tools have facilitated *en masse* protein structure predictions for entire proteomes^51–53^. By comparison, similar ML-based co-folding tools for predicting protein-ligand complexes are in earlier stages of development^54–57^. We considered our stereoprobe ligandability maps, which included both well-characterized small molecule-binding sites in proteins (e.g., kinase active sites) and many previously unannotated sites in proteins lacking ligands, to offer a unique opportunity for evaluating the ability of co-folding tools to predict diverse types of protein-ligand interactions. Indeed, because our ligandability maps report on covalent binding to specific cysteine residues, they provide a defined location for each small molecule interaction in protein structures. And, even though ML programs have been trained mostly on reversible small molecule-protein interactions, we interpreted the stereoselectivity and, in most cases, additional regioselectivity and chemoselectivity of the covalent stereoprobe binding events to reflect a degree of molecular recognition suitable for minimally testing whether ML programs can place ligands in proximity to their target cysteines.

We generated noncovalent protein-ligand complex predictions for the stereoselective liganding events identified for stereoprobes **1-8** and **21-52** by cysteine-directed ABPP using Boltz-2^54^, a recently introduced open-source biomolecular foundation model for macromolecular complex prediction inspired by the framework of AlphaFold3. In total, this effort generated predictions for 102 unique proteins with one or more of 32 unique tryptoline acrylamide stereoprobes. An additional 31 stereoprobe-liganded proteins ranging from 169 to 796 kDa failed to generate predictions due to memory constraints and were excluded from further analysis. For each protein-ligand pair, Boltz-2 inference was performed with 10 replicates, leading to a total of 1,710 models for 171 protein-ligand pairs representing 218 unique protein-ligand-cysteine combinations. Boltz-2 consistently produced models of protein-ligand pairs with valid bond lengths and angles, as well as intermolecular distances free of clashes^58^ (**Supplementary Dataset 2**). We quantified the accuracy of the ligand site prediction by measuring distances from the liganded cysteine (as determined by ABPP) and the modeled ligand in each replicate of Boltz-2 inference (**Fig. 2a**). For this analysis, we considered the distance from the nearest atoms in the liganded cysteine and the modeled ligand without regard for the pose of the ligand or orientation of the reactive acrylamide with respect to the cysteine.

**Fig. 2.**
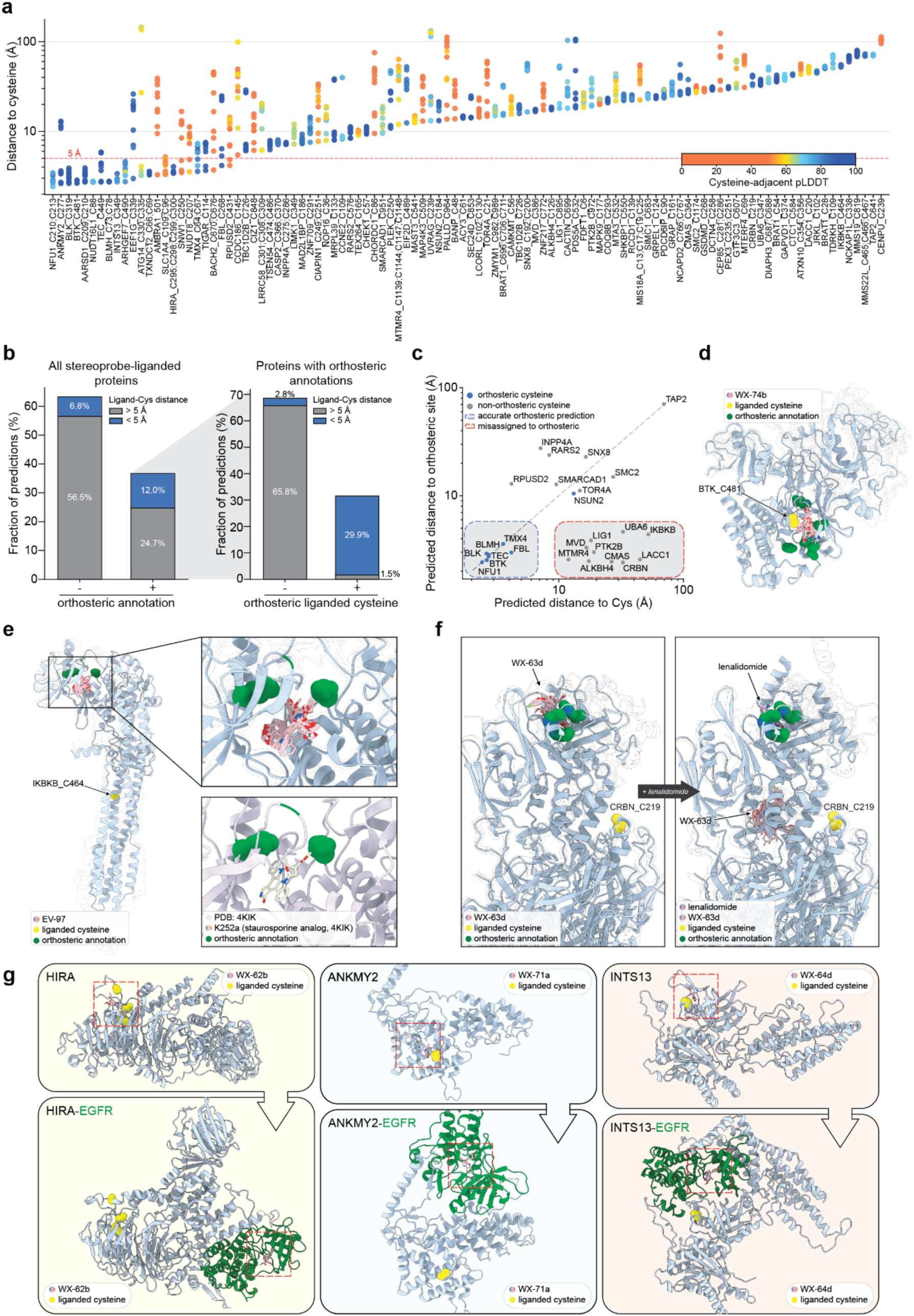
Boltz-2 analysis of tryptoline acrylamide-liganded proteins. **a**, Waterfall plot showing cysteine-ligand distances for each protein identified as having a tryptoline acrylamide stereoprobe-liganded cysteine (**Supplementary Dataset 1**). The ten diffusion samples for the protein-stereoprobe pair with the closest stereoprobe-cysteine distance are shown. Proteins are sorted by closest stereoprobe distance and points are colored according to the mean pLDDT of all residues within 5 Å of the liganded cysteine. **b**, *Left*: Stacked bar graphs showing the fraction of proteins with and without orthosteric site annotations in UniProt colored by predicted stereoprobe proximity to the liganded cysteine (blue: < 5 Å (correct predictions); gray: > 5 Å (incorrect predictions)). *Right*: Stacked bar graphs, where each bar shows the fraction of liganded cysteines occupying (*right*) or not occupying (*left*) orthosteric sites for the subset of proteins with orthosteric site annotations in UniProt, and where each bar is then further color-coded for correct (blue) and incorrect (gray) predictions as defined in left panel. **c**, Scatter plot representing the relationship between the shortest distance between the stereoprobe and the liganded cysteine (x-axis) versus the stereoprobe and the orthosteric site (y-axis) for each protein with an unambiguous orthosteric assignment. **d**, Structures for ten diffusion samples of predictions for WX-74b binding to BTK. The rank 0 BTK structure is blue, ranks 1-9 are transparent. Orthosteric annotations on the rank 0 structure are denoted in green. The liganded cysteine BTK_C481 is yellow. All WX-74b ligands are pink. **e**, Structures for ten diffusion samples of predictions for EV-97 binding to IKBKB. The rank 0 IKBKB structure is blue, ranks 1-9 are transparent. Orthosteric annotations on the rank 0 structure are denoted in green. The liganded cysteine IKBKB_C464 is yellow. All EV-97 ligands are pink. Insets show the orthosteric site for IKBKB with incorrect predictions of EV-97 binding to this site (*top inset*) or with an orthosteric ligand K252a binding to this site in an experimental structure (PDB: 4KIK). **f**, Structures for ten diffusion samples of predictions of WX-63d binding to a CRBN:DDB1^ΔBPB^ complex in the presence (*left*) or absence (*right*) of the orthosteric ligand lenalidomide. The rank 0 CRBN:DDB1^ΔBPB^ structure is blue, ranks 1-9 are transparent. Orthosteric annotations on the rank 0 structure are denoted in green. The liganded cysteine CRBN_C219 is yellow. All WX-63d ligands are pink. **g**, Structures representing accurate Boltz-2 protein-ligand co-folding predictions for representative non-orthosteric stereoprobe liganding events with HIRA, ANKMY2, and INTS13 proteins (*top*), and the corruption of these predictions by fusing the proteins to a protein with a well-defined orthosteric pocket (EGFR kinase domain (green); *bottom*). The liganded cysteines in each image are yellow. Red dashed boxes highlight the location of stereoprobes in each co-folding prediction.

These measurements revealed mixed performance with the ligands in our panel. Of the 102 proteins, only 23 produced at least one model with the ligand within 5 Å of the known liganded cysteine, which we considered a successful prediction (**Fig. 2a, b**). Most of the successful predictions exhibited high confidence pLDDT scores near the cysteine, but a handful (six) showed lower pLDDT scores that also appeared to correlate with wider distributions of ligand-cysteine distances across the replicate predictions (**Fig. 2a**).

By further cross-referencing existing data on functional sites of the modeled proteins (i.e., active site and binding site annotations from UniProt, the majority of which are derived from experimental structures^59,60^), we found that the subset of proteins with accurate ligand predictions was enriched in proteins containing known orthosteric small molecule-binding pockets (**Fig. 2b-d**), including kinases (e.g., BTK (**Fig. 2d**), BLK, TEC) and metabolic enzymes (e.g., BLMH, TIGAR). Interestingly, among proteins with orthosteric site annotations, the success of predictions was very high for liganded cysteines located within 5 Å of the orthosteric site (239 out of 252 predictions) and dramatically lower for liganded cysteines located elsewhere in the protein (22 out of 548 predictions) (**Fig. 2b**, right bar graph). Further assessment of the latter category revealed a large cluster of proteins with liganded cysteines at non-orthosteric sites for which Boltz-2 incorrectly predicted stereoprobe binding at the orthosteric sites (e.g., IKBKB, CRBN, LIG1, MVD) (**Fig. 2c**). As one example, we show predictions for the kinase IKBKB, where the liganded cysteine C464 is located on a non-kinase domain helix distal (> 20 Å) from the ATP-binding pocket, but Boltz-2 nonetheless consistently places the stereoprobe ligand in this orthosteric pocket, which is the site where previously reported ligands (e.g., staurosporine) bind in crystal structures of IKBKB in the PDB^61^ (**Fig. 2e**).

We wondered whether the Boltz-2 predictions could be improved by filling the orthosteric site on a protein with a known ligand. To this end, we co-folded the E3 ligase CRBN with tryptoline acrylamide WX-63d (a ligand for C219 of CRBN) in the presence and absence of the orthosteric ligand lenalidomide^62–64^. The predictions were also performed in the presence of a DDB1^ΔBPB^ construct used previously for structural studies of CRBN:DDB1 complexes^65^. In the absence of lenalidomide, Boltz-2 incorrectly placed WX-63d in the orthosteric site of CRBN that binds lenalidomide and the natural C-terminal cyclic imide substrates^66,67^ of this E3 ligase (**Fig. 2f**, left). In the presence of lenalidomide, WX-63d was displaced from the orthosteric pocket, but was not redirected to bind in proximity to C219 and instead positioned at a distinct site on CRBN (**Fig. 2f**, right). A similar outcome was observed for spirocycle acrylamide stereoprobes previously found to engage C287 of CRBN^40^ (**Extended Data Fig. 2a**), as well as for tryptoline acrylamide EV-97 and IKBKB when cofolded with the orthosteric ligand K252a (**Extended Data Fig. 2b**), indicating that, even when the orthosteric pocket is blocked on proteins, Boltz-2 still struggles to locate non-orthosteric ligandable sites.

Despite the challenges that Boltz-2 faced in assigning non-orthosteric stereoprobe liganding events to proteins *with* orthosteric pockets, we were intrigued to see multiple examples of successful prediction of non-orthosteric liganding events on proteins *lacking* orthosteric sites (e.g., HIRA, ANKMY2, and INTS13) (**Fig. 2a, g**). While we do not yet understand why Boltz-2 succeeded with these proteins, we note that, in each case, the liganded cysteine and its surrounding region had a high pLDDT score (**Fig. 2a**). The success with HIRA, in particular, is striking given that this protein is > 100 kDa in size, and Boltz-2 consistently placed the stereoprobe ligand in proximity to candidate liganded cysteine (C295) (**Fig. 2a**), which is predicted to reside on the side of the donut-like WD40 repeat domain of the protein (**Fig. 2g**). This cysteine is distant from the more common central ligand binding pocket presented by WD40-like folds as well as the side pocket D-box binding site reported in the structure of apcin bound to Cdc20 (**Fig. 2g** and **Extended Data Fig. 3a**)^68,69^. We finally evaluated the robustness of these successful Boltz-2 predictions, by artificially fusing each stereoprobe-liganded protein to another protein with a known orthosteric pocket (the kinase domain of EGFR)^70^. For each fusion protein, Boltz-2 redirected the stereoprobe ligands to the ATP-binding pocket of EGFR (**Fig. 2g**), suggesting that correct non-orthosteric liganding predictions made by Boltz-2 can be readily corrupted when contesting with an orthosteric pocket on the same protein.

The results of our Boltz-2 analysis of covalent stereoprobe-protein interactions indicate that this program predicts the stereoprobe binding events that occur at established orthosteric sites with high success rates but has difficulty locating non-orthosteric liganding events. This challenge was compounded for proteins with orthosteric pockets, which often served as sites for the incorrect placement of ligands. We should finally note that, because the version of Boltz-2 available at the time of analysis lacked effective stereochemical restraints for small molecules, our predictions sampled all stereoisomers of each stereoprobe. When we reviewed the stereoconfiguration of ligands found in correct predictions, we did not observe a substantial enrichment for experimentally determined stereoisomers from ABPP experiments (32% correct stereoconfiguration vs 25% expected for random assignment; **Extended Data Fig. 3b** and **Supplementary Dataset 2**), indicating that, in these instances, Boltz-2 was capable of locating the liganded pockets, but not the type of chemistry that preferentially bound/reacted with these pockets. Later versions of Boltz-2 that include effective stereochemical restraints were confirmed to accurately recapitulate the stereochemistry of the ligands specified in the input without yielding an improvement in the accuracy of the predicted ligand placement with respect to the liganded cysteine (**Extended Data Fig. 3b, c**).

### Characterization of representative stereoprobe-cysteine interactions

Several of the correct Boltz-2 predictions were confidently assigned as orthosteric liganding events based on the corresponding cysteines also having been targeted by other covalent ligands (e.g., the BTK, TEC, and BLK kinases^71^) or representing conserved catalytic residues (e.g., BLMH, TMX4). There were, however, also less well-characterized cases, such as the methyltransferase FBL, which was liganded by the elaborated stereoprobe WX-72d at C268 (**Fig. 3a**), a cysteine that resides at the edge of the FBL active site (**Fig. 3b**). Boltz-2 predicted that this liganding event occurs in the *S*-adenosyl methionine (SAM) cofactor-binding pocket (**Figs. 2c** and **3b**). We confirmed using established gel-ABPP methods^42,72^ that recombinant WT-FBL, but not a C268S-FBL mutant, stereoselectively reacted with an alkyne analog of WX-72d (WX-72d-yne) in HEK293T cells (**Fig. 3c, d**) or cell lysates (**Extended Data Fig. 4a**), and that this interaction was stereoselectively and chemoselectively blocked by pre-treatment with WX-72d (**Fig. 3e** and **Extended Data Fig. 4b**). Interestingly, however, co-treatment with SAM did not block, but rather increased the stereoselective reactivity of WX-72d-yne with WT-FBL in a concentration-dependent manner (**Fig. 3f, g** and **Extended Data Fig. 4c**). In contrast, SAH did not affect WX-72d-yne-WT-FBL interactions (**Fig. 3f, g** and **Extended Data Fig. 4c**). SAM also enhanced WX-72d-yne reactivity with endogenous FBL in Ramos cell lysates as determined by protein-directed ABPP (**Extended Data Fig. 4d**). We interpret these results to indicate that WX-72d-yne binds to a pocket proximal to, but not overlapping with, the SAM-binding pocket in FBL, thus underscoring the importance of experimentally testing ML predictions even in cases where the small molecule is placed in an orthosteric pocket proximal to a site of liganding mapped by chemical proteomics. Notably, attempts to predict the presence of a proximal pocket with Boltz-2 by filling the orthosteric pocket with a molecule of SAM were unsuccessful, as this experiment resulted in the misplacement of WX-72d-yne at a site distal to C268 (**Extended Data Fig. 4e**).

**Fig. 3.**
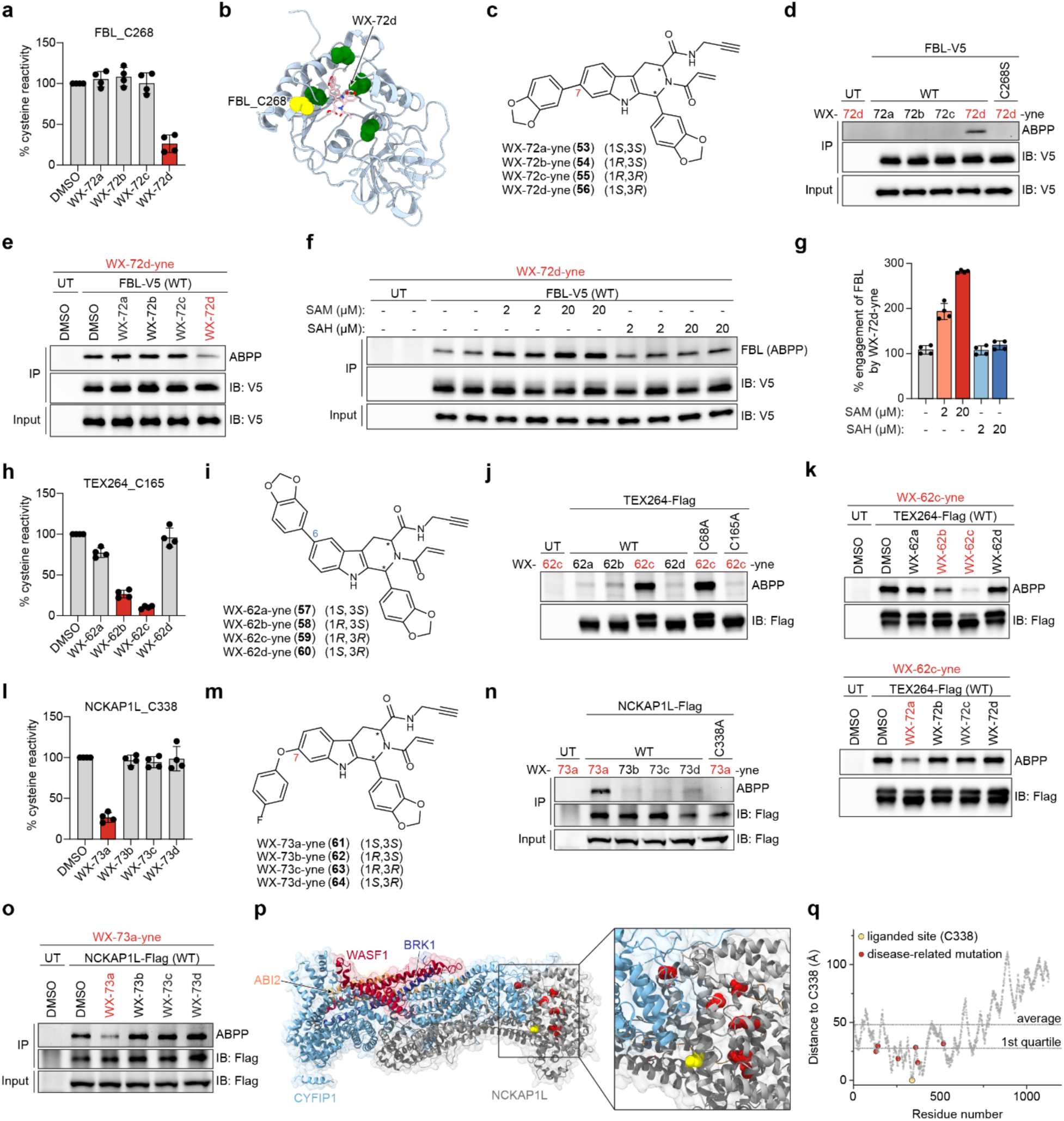
Characterization of elaborated stereoprobe-cysteine interactions. **a**, Cysteine-directed ABPP results showing stereoselective engagement of FBL_C268 by WX-72d. Data represent average values ± SD for four independent experiments. **b,** Boltz-2 prediction for WX-72d binding to FBL showing ligand placement proximal to C268 (yellow) in the SAM/SAH binding pocket. **c,** Structures of alkynylated elaborated stereoprobes used to confirm liganding of FBL. **d,** Gel-ABPP data showing stereoselective engagement of recombinant FLAG epitope-tagged WT-FBL, but not a C268S-FBL mutant by WX-72d-yne (5 µM, 1 h) in HEK293T cells. UT, untransfected cells. **e**, Gel-ABPP data showing stereoselective blockade of WX-72d-yne (5 µM, 1 h) engagement of recombinant WT-FBL by WX-72d (20 µM, 1 h pretreatment) in HEK293T cells. **f,** Gel-ABPP data showing increased engagement of WT-FBL by WX-72d-yne in the presence of SAM but not SAH. Experiments were performed in lysates of HEK293T cells stably expressing WT-FBL. **g,** Quantification of gel-ABPP signals from **f** normalized to DMSO signals. Data represent average values ± SD for four independent experiments. **h,** Cysteine-directed ABPP results showing enantioselective engagement of TEX264_C165 by WX-62b and WX-62c. Data represent average values ± SD for four independent experiments. **i,** Structures of alkynylated elaborated stereoprobes used to confirm liganding of TEX264. **j,** Gel-ABPP data showing stereoselective engagement of recombinant FLAG epitope-tagged WT-TEX264 and a C68A-TEX264 mutant, but not a C165A-TEX264 mutant, by WX-62c-yne (5 µM, 1 h) in HEK293T cells. **k,** Gel-ABPP data showing enantioselective blockade of WX-62c-yne (5 µM, 1 h) engagement of recombinant WT-TEX264 by C6 elaborated probes WX-62b and WX-62c and C7 elaborated probe WX-72a (20 µM, 2 h) in HEK293T cells. **l,** Cysteine-directed ABPP results showing stereoselective engagement of NCKAP1L_C338 by WX-73a. Data represent average values ± SD for four independent experiments. **m,** Structures of alkynylated elaborated stereoprobes used to confirm liganding of NCKAP1L. **n,** Gel-ABPP data showing stereoselective engagement of recombinant FLAG epitope-tagged WT-NCKAP1L, but not a C338A-NCKAP1L mutant by WX-73a-yne (5 µM, 1 h) in THP1 cells. **o,** Gel-ABPP data showing stereoselective blockade of WX-73a-yne (5 µM, 1 h) engagement of WT-NCKAP1L by WX-73a (20 µM, 1 h pretreatment) in THP1 cells. **p,** Overlay of NCKAP1L AlphaFold-predicted structure (AF-P55160-F1-model_v4) with the WAVE regulatory complex crystal structure containing the NCKAP1L paralog NCKAP1 (PDB: 3P8C) showing location of C338 (yellow) relative to NCKAP1L hotspot mutations (red: M371V, R258L, P359L, V519L, R129W, V141F) that lead to human immunological disorder. **q,** Graph showing distance of NCKAP1L residues mutated in immunological disorders (red) to NCKAP1L_C338 (yellow) in AF-P55160-F1-model_v4. For **d-f, j, k, n, o**, gel-ABPP data represent stereoprobe treatments of cells transiently (TEX264) or stably (FBL and NCKAP1L) expressing epitope-tagged WT or C-to-A/S mutant. ABPP signals were measured by CuAAC conjugation to rhodamine-azide tag followed by SDS-PAGE and in-gel fluorescence scanning; data are from single experiments representative of at least two independent experiments.

We next turned our attention to the most populated category of liganding events emerging from the Boltz-2 analysis – incorrect predictions occurring on proteins that lack orthosteric annotations (**Fig. 2b**). We selected three proteins from this category for confirmatory studies – TEX264, NCKAP1L, and GRPEL1 – that are from different structural and functional classes and, to our knowledge, have heretofore lacked small-molecule ligands.

TEX264 is an integral membrane endoplasmic reticulum autophagy receptor^73–76^ and was liganded exclusively by elaborated tryptoline acrylamides, with the largest change in reactivity observed for WX-62c at C165 along with lower magnitude decreases in C68 reactivity (**Fig. 3h** and **Extended Data Fig. 4f**). Gel-ABPP experiments verified stereoselective reactivity of recombinant WT-TEX264 by WX-62c-yne in HEK293T cells, and mutagenesis supported C165 as the site of covalent engagement (**Fig. 3i, j**). We also verified stereoselective blockade of WX-62c-yne reactivity with WT-TEX264 by both C6 (WX-62c) and C7 (WX-72a) benzodioxolyl-modified tryptoline acrylamides (**Fig. 3k** and **Extended Data Fig. 4g**), while the corresponding unsubstituted analog EV-99 was inactive (**Extended Data Fig. 4g**). These gel-ABPP data for recombinant WT-TEX264 matched the SAR observed for endogenous TEX264 by cysteine-directed ABPP (**Extended Data Fig. 4f**).

NCKAP1L (or HEM1) is an immune-enriched scaffolding component of the WAVE regulatory complex (WRC) involved in actin remodeling^77–79^. NCKAP1L was liganded at C338 exclusively by the C7 fluorophenylether stereoprobe WX-73a (**Fig. 3l** and **Extended Data Fig. 4h**). We confirmed the striking stereo-and chemo-selectivity of this interaction with alkyne analog WX-73a-yne and recombinant WT-NCKAP1L stably expressed in THP-1 cells and further verified the loss of stereoprobe reactivity with a C338A-NCKAP1L mutant (**Fig. 3m, n** and **Extended Data Fig. 4i**). Of note, C338 is not found in the paralogous protein NCKAP1 and is also proximal to several missense mutations in NCKAP1L that disrupt WRC signaling and cause immunodysregulatory disorders in humans^77,80^ (**Fig. 3p, q**). These findings suggest that ligands targeting NCKAP1L_C338 may provide a way to modulate the function of this scaffolding protein in a paralog-restricted manner.

GRPEL1 is a nucleotide exchange factor for the mitochondrial chaperone HSPA9 (or mortalin or mtHSP70)^81,82^, which regulates the import, folding, and degradation of proteins to support mitochondrial biogenesis and homeostasis^83,84^. The C7 phenylacetylene (1*R*,3*S*) and (1*R*,3*R*) stereoprobes WX-71b and WX-71c, respectively, each enantioselectively liganded C124 in GRPEL1 in Ramos cells (**Fig. 4a**), and recombinant WT-GRPEL1 stably expressed in HCT-116 cells exhibited a similar enantioselective reactivity profile with alkyne tryptoline acrylamides WX-01-06 and WX-01-07 (**Fig. 4b**). These alkyne stereoprobes did not react with a C124A-GRPEL1 mutant (**Fig. 4b**). WX-71b and WX-71c, but not their respective enantiomers WX-71d and WX-71a, blocked the WX-01-07-WT-GRPEL1 interaction, with WX-71b showing the higher potency (IC_50_ value = 2.0 µM; 95% CI = 1.5 – 3 µM; **Fig. 4c** and **Extended Data Fig. 5a**).

**Fig. 4.**
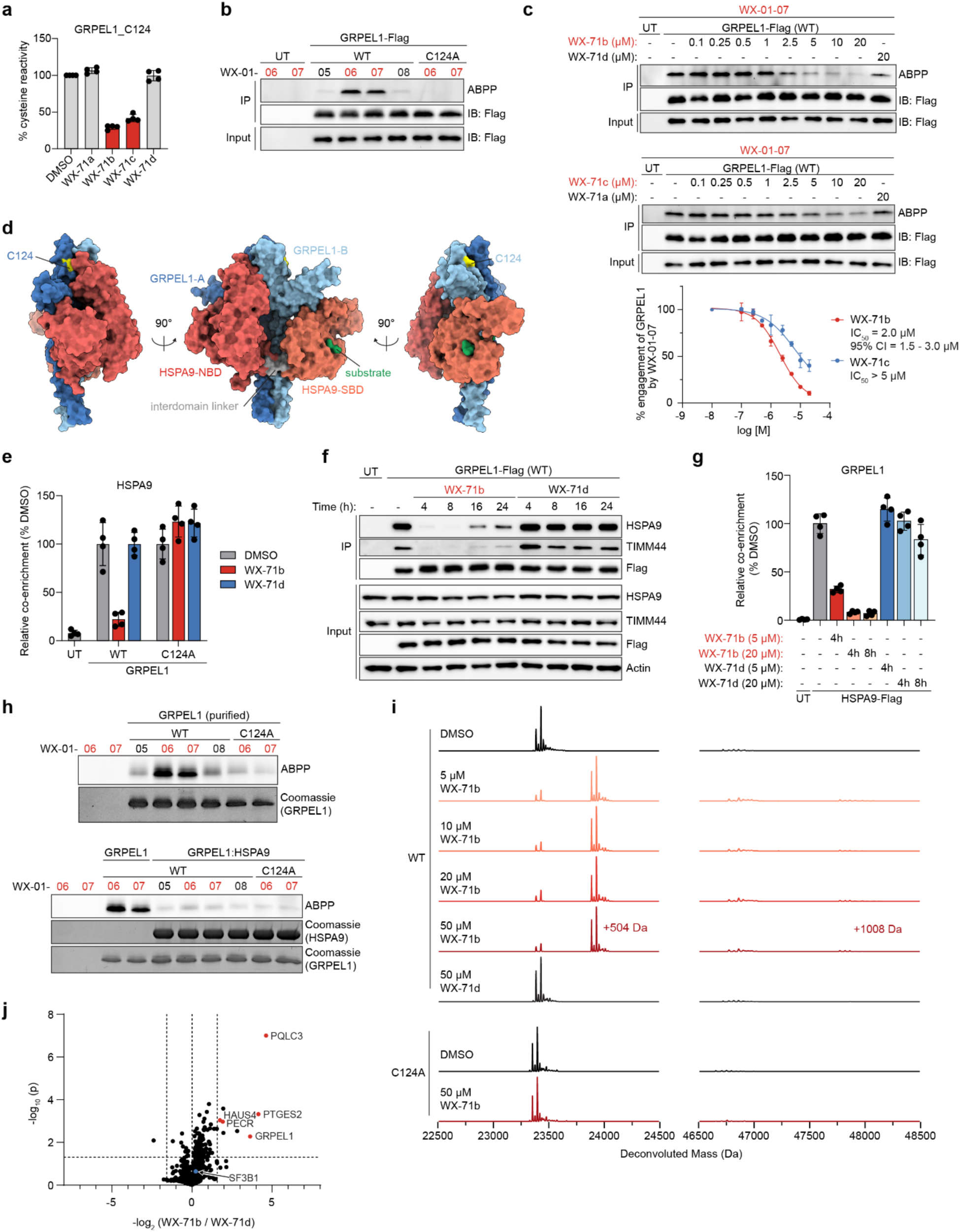
Tryptoline acrylamide stereoprobes disrupt GRPEL1:HSPA9 interactions. **a**, Cysteine-directed ABPP data showing enantioselective engagement of GRPEL1_C124 by WX-71b and WX-71c. Data represent average values ± SD of four independent experiments. **b,** Gel-ABPP data showing enantioselective engagement of recombinant FLAG epitope-tagged WT-GRPEL1, but not a C124A-GRPEL1 mutant by WX-01-06 and -07 (5 µM, 1 h) in HCT-116 cells. UT, untransfected cells. **c,** Gel-ABPP data showing concentration-dependent and enantioselective blockade of WX-01-07 (5 µM, 1 h) engagement of WT-GRPEL1 by WX-71b (top gel) and WX-71c (bottom gel) (2 h pretreatment). Top, representative gel-ABPP data; bottom, quantification of gel-ABPP data. Data represent average values ± SD for 2 independent experiments. **d,** Cryo-EM structure^85^ of GRPEL1 dimer (blue) in complex with HSPA9 (red/orange) (PDB: 9BLS) showing location of stereoprobe-liganded cysteine GRPEL1_C124 (yellow) at the GRPEL1:HSPA9 interface. SBD: substrate binding domain; NBD: nucleotide binding domain; GRPEL1-A, protomer 1; GRPEL1-B, protomer 2. **e,** IP-MS data showing stereoselective disruption of recombinant GRPEL1 interactions with endogenous HSPA9 following treatment with WX-71b or enantiomeric control WX-71d (20 µM, 4 h) in HCT-116 cells. HSPA9 signals for treatment groups were normalized to DMSO signals within each WT- and C124A-GRPEL1 group. Data represent average values ± SD for four independent experiments. **f,** IP-western blotting data showing time-dependent and enantioselective disruption of recombinant GRPEL1 interactions with endogenous HSPA9 and TIMM44 following treatment with WX-71b or enantiomeric control WX-71d (20 µM, 4-24 h) in HCT-116 cells. **g,** IP-MS data showing stereoselective disruption of recombinant FLAG epitope-tagged HSPA9 interactions with endogenous GRPEL1 following treatment with WX-71b or WX-71d (5 or 20 µM, 4 or 8 h) in HCT-116 cells. GRPEL1 signals for each treatment group were normalized to DMSO signals. Data represent average values ± SD for four independent experiments. **h,** Gel-ABPP data showing enantioselective and site-specific engagement of purified GRPEL1 (top) but not purified GRPEL1 in complex with HSPA9 (bottom) by WX-01-06 and -07 (10 µM, 2 h). **i,** Intact protein MS data for purified WT-GRPEL1 and C124A-GRPEL1 (3.5 µM) incubated with the indicated concentrations of WX-71b or WX-71d (4 h). Proteins were analyzed by time-of-flight (TOF)-LC/MS. Data show deconvoluted mass spectra from a single experiment representative of two independent experiments. **j,** Volcano plot for protein-directed ABPP data in parental HCT-116 cells showing blockade of proteins enriched by WX-01-06 by pre-treatment with WX-71b or WX-71d. Dashed vertical lines mark proteins showing greater than 3-fold stereoselective blockade by WX-71b or WX-71c. Proteins stereoselectively engaged by WX-71b that also show less than 2-fold engagement by WX-71c are marked in red. SF3B1 is in blue. Data represent average values ± SD for four independent experiments. Statistical significance was assessed using Welch two sample t-tests. For **b**, **f**, and **h** data are from a single experiment representative of at least two independent experiments.

Our follow-up studies with recombinant proteins thus generally verified the stereoselectivity and chemoselectivity of the non-orthosteric covalent liganding events mapped for elaborated tryptoline acrylamides by cysteine-directed ABPP. We next set out to determine the functional impact of such liganding events using GRPEL1 as a case study.

### GRPEL1 ligands disrupt the GRPEL1-HSPA9 protein complex

Structures of the human GRPEL1:HSPA9 and *E. coli* GrpE:DnaK complexes indicate that C124 (V108 in GrpE) is located near the interface of GRPEL1 and HSPA9^85,86^ (**Fig. 4d**). We therefore hypothesized that the GRPEL1 ligands might perturb the GRPEL1:HSPA9 complex in cells.

Consistent with this premise, we found by immunoprecipitation (IP)-MS and IP-Western blotting experiments that WX-71b, but not its inactive enantiomer WX-71d, disrupted GRPEL1 binding to HSPA9 in cells expressing a FLAG epitope-tagged WT-GRPEL1 protein, but not a C124A-GRPEL1 mutant (**Fig. 4e, Extended Data Fig. 5b**, and **Supplementary Dataset 3**). WX-71b also stereoselectively and site-specifically blocked GRPEL1 interactions with other mitochondrial and mitochondrial-associated proteins (**Fig. 4f** and **Extended Data Fig. 5c**). The disruption of the GRPEL1:HSPA9 complex was sustained for up to 24 h (**Fig. 4f**), matching the duration of GRPEL1_C124 engagement by WX-71b as determined by gel-ABPP (**Extended Data Fig. 5d**). IP-MS experiments performed in the converse direction in HCT-116 cells stably expressing FLAG epitope-tagged HSPA9 protein further verified the stereoselective disruption of the GRPEL1:HSPA9 complex by WX-71b (5 or 20 µM, 4 or 8 h treatment; **Fig. 4g, Extended Data Fig. 5e**, and **Supplementary Dataset 3**). Most of the other HSPA9-associated proteins were unaffected by the GRPEL1 ligand (**Extended Data Fig. 5e**), suggesting that other HSPA9 interactors may stay bound to this protein in the absence of GRPEL1.

We found that the tryptoline acrylamide stereoprobes also stereoselectively and site-specifically reacted with recombinant purified WT-GRPEL1 protein^85^, as determined by gel-ABPP (with alkynes WX-01-06 and WX-01-07; **Fig. 4h**) or intact MS (with WX-71b; **Fig. 4i** and **Extended Data Fig. 5f**). Co-incubation with purified HSPA9 blocked stereoselective engagement of WT-GRPEL1 by WX-01-06 and WX-01-07 (**Fig. 4h**), supporting that the covalent liganding of GRPEL1_C124 is mutually exclusive with HSPA9 binding.

Having established that tryptoline acrylamides stereoselectively and site-specifically disrupt GRPEL1 interactions with HSPA9 in cells, we next investigated how the GRPEL1 ligands impacted mitochondrial biology.

### GRPEL1 ligands disrupt mitochondrial protein import and promote mitophagy

Previous ABPP studies have identified the splicing factor SF3B1 as a frequent target of (1*R*,3*S*) tryptoline acrylamides^21,22,35^. These compounds react with C1111 in a functional pocket of SF3B1 that also binds natural products such as pladienolide B, leading to global perturbations in RNA splicing and general impairments in cell growth^35^. The liganding of SF3B1 accordingly has the potential to confound the interpretation of biological effects of (1*R*,3*S*) tryptoline acrylamide interactions with other proteins in human cells, a technical challenge that we have previously addressed by generating drug-resistant C1111S-SF3B1-expressing cells^22^. Here, however, we found by protein-directed ABPP that the GRPEL1 ligand WX-71b did not substantially engage SF3B1 and only showed a handful of other stereoselective off-targets in HCT-116 cells (HAUS4, PECR, PTGES2; **Fig. 4j** and **Extended Data Fig. 5g**). These experiments also showed that WX-71b does not engage GRPEL2 (**Extended Data Fig. 5g**), a paralog of GRPEL1 that shares ∼45% sequence identity, including C124 (C126 in GRPEL2)^87^.

Encouraged that the C7-phenylacetylene group in WX-71b removed SF3B1 as an off-target of concern for cellular studies, we proceeded to investigate the effects of this compound on mitochondrial biology. The mtHSP70 chaperone complex plays a critical role in the import of proteins into the mitochondria^88^. Using a doxycycline-inducible mitochondrial-localized EGFP reporter system^89–91^, we found that mitochondrial protein import in HCT-116 cells was disrupted by WX-71b, but not by the inactive enantiomer WX-71d (20 µM each compound, 8 h; **Fig. 5a, b** and **Extended Data Fig. 6a**). Additional experiments were performed in HCT-116 cell models stably expressing WT-GRPEL1 or a C124A-GRPEL1 mutant against a background of CRISPR/Cas9-mediated disruption of endogenous GRPEL1 (sgGRPEL1 cells; **Extended Data Fig 6b**), which revealed that WX-71b impaired EGFP mitochondrial import in cells expressing WT-, but not C124A-GRPEL1 (**Fig. 5c, d**).

**Fig. 5.**
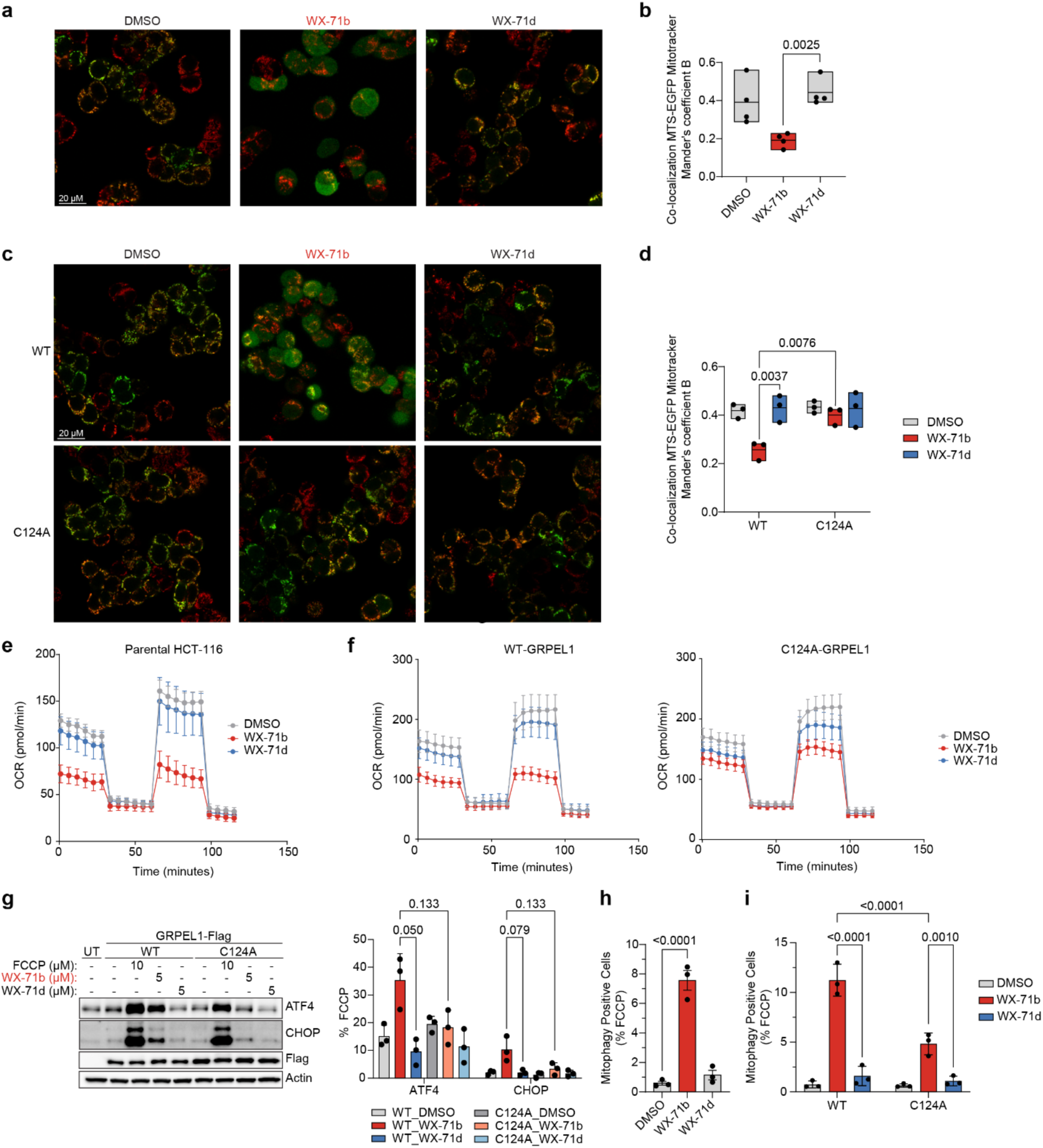
GRPEL1 stereoprobes modulate mitochondrial protein import and function. **a**, Microscope images showing localization of EGFP and Mitotracker Deep Red FM in MTS-EGFP-inducible parental HCT-116 cells treated with doxycycline (0.5 µg/mL) and WX-71b or WX-71d (20 µM, 8 h). **b,** Quantification of **a** where mitochondrial localization of EGFP was determined by colocalization with Mitotracker Deep Red FM using the Imaris 10.2 colocalization module. For each treatment group, 10 images per independent experiment were analyzed (**Extended Data Fig. 5a** shows one independent experiment with 10 images). Data represent average values ± SD of four independent experiments. Statistical significance was assessed using Welch two sample t-test. **c,** Microscope images showing localization of EGFP and Mitotracker Deep Red FM in MTS-EGFP-inducible sgGRPEL1 HCT-116 cells expressing recombinant FLAG epitope-tagged WT-GRPEL1 (top) or C124A-GRPEL1 (bottom) treated with doxycycline (0.5 µg/mL) and DMSO or WX-71b or WX-71d (20 µM, 8 h). **d,** Quantification of **c**, as described in panel **b**. Data represent average values ± SD of three independent experiments. Statistical significance was assessed using two-way ANOVA with Šídák’s multiple comparisons test (reported p values are adjusted p values). **e, f,** Mitochondrial respiration measured by a Seahorse mitochondrial stress test in (**e**) parental HCT-116 cells or (**f**) sgGRPEL1 HCT-116 cells expressing recombinant FLAG epitope-tagged WT-GRPEL1 (left) or C124A-GRPEL1 (right) treated with DMSO or WX-71b or WX-71d (5 µM, 8 h). Data represent average values ± SD of ten technical replicates from a single experiment representative of three independent experiments. **g,** Left, immunoblotting of integrated stress response (ISR) markers ATF4 and CHOP in sgGRPEL1 HCT-116 expressing recombinant Flag epitope-tagged WT- or C124A-GRPEL1 treated with DMSO or WX-71b or WX-71d (5 µM, 8 h). Right, quantification of blotting signals. Protein signals were normalized to FCCP-treated samples. Data represent average values ± SD of three independent experiments. Statistical significance was assessed using unpaired-multiple t-tests (Holm-Šídák approach to multiple comparisons; reported p values are adjusted p values). **h,** Parental HCT-116 cells stably expressing mt-mKeima and Parkin were treated with DMSO, FCCP (10 µM), or WX-71b or WX-71d (20 µM, 8 h), and mitophagy flux was measured by flow cytometry. Cells showing increased 561 nm/405 nm mt-mKeima ratios were considered mitophagy-positive. Data represent average values ± SD of three independent experiments. Statistical significance was assessed using ordinary one-way ANOVA with Dunnett’s multiple comparison (reported p values are adjusted p values). **i,** sgGREPL1 HCT-116 cells expressing mt-mKeima, Parkin, and Flag epitope-tagged WT- or C124A-GRPEL1 were treated with DMSO, FCCP (10 µM), or WX-71b or WX-71d (20 µM, 8 h), and mitophagy flux was measured by flow cytometry. Cells showing increased 561 nm/405 nm mt-mKeima ratios were considered mitophagy-positive. Data represent average values ± SD of three independent experiments. Statistical significance was assessed using two-way ANOVA with Šídák’s multiple comparisons test (reported p values are adjusted p values).

We next examined mitochondrial protein import more globally by pulse-chase SILAC (stable isotope labeling by amino acids in cell culture) quantitative proteomics^90,92^. WT- and C124A-GRPEL1 cells grown in light amino acid media were pretreated with DMSO or WX-71b or WX-71d (20 µM, 4 h), after which the cells were shifted to heavy amino acid media for 8 h in the continued presence of stereoprobes (5 µM each) (**Extended Data Fig. 6c**). Cells were then lysed and mitochondria isolated and analyzed by quantitative proteomics, which revealed that WX-71b-treated WT-GRPEL1 cells showed a general decrease in heavy amino acid-labeled mitochondrial proteins in comparison to control groups (WX-71d-treated WT-GRPEL1 cells or WX-71b-treated C124A-GRPEL1 cells; **Extended Data Fig. 6d** and **Supplementary Dataset 3**). The stereoselective impairment in import caused by WX-71b in WT-GRPEL1 cells was observed for proteins from different sub-mitochondrial compartments, with inner mitochondrial membrane (IMM) and matrix proteins showing the strongest effect (**Extended Data Fig. 6e**).

In addition to suppressing mitochondrial protein import, WX-71b (5 µM, 8 h) also stereoselectively decreased basal and maximal respiration in both parental and WT-GRPEL1-expressing cells (**Fig. 5e, f**), and this effect was markedly attenuated in C124A-GRPEL1-expressing cells (**Fig. 5f**). We interpret the low residual effect of WX-71b in C124A-GRPEL1-expressing cells to possibly reflect perturbation of the remaining endogenous WT-GRPEL1 in these cells, as the population-based CRISPR-Cas9 disruption of the *GRPEL1* gene was incomplete (**Extended Data Fig. 6b**).

Mitochondrial stress caused by impairments in protein import and respiration can lead to activation of the integrated stress response (ISR)^93,94^. We found that WX-71b (5 µM, 8 h) promoted a stereoselective increase in ISR biomarkers ATF4 and CHOP in WT-GRPEL1-, but not C124A-GRPEL1-expressing cells (**Fig. 5g**). This effect was, however, much less dramatic than that caused by the oxidative phosphorylation uncoupling agent carbonyl cyanide *p*-(trifluoromethoxy) phenylhydrazone (FCCP) (**Fig. 5g**). HSPA9 knockdown has also been reported to induce PINK1-dependent mitophagy^89^. To evaluate the impact of GRPEL1 ligands on mitophagy, we established an HCT-116 cell model expressing Parkin (PRKN) and a mitochondrial matrix-targeted (mt)-mKeima reporter protein and then treated these cells with DMSO or stereoprobe (WX-71b or WX-71d; 20 µM, 8 h) and measured mitophagy flux by flow cytometry^89,95,96^. This experiment revealed that WX-71b, but not WX-71d induced mitophagy (**Fig. 5h** and **Extended Data Fig. 6f**). We observed a similar stereoselective increase in mitophagy for WX-71b in WT-GRPEL1-expressing cells that was much greater in magnitude than in C124A-GRPEL1-expressing cells (**Fig. 5i** and **Extended Data Fig. 6f**).

Our functional studies thus indicate that acute chemical disruption of the GRPEL1:HSPA9 complex leads to rapid impairments in mitochondrial import and respiration and corresponding activation of the ISR and mitophagy in human cells.

## Discussion

Chemical proteomics holds great promise for enriching our understanding of small molecule-protein interactions in cells and identifying first-in-class ligands for proteins. The limited throughput of untargeted MS analysis, however, makes chemical proteomics more suited for evaluating focused libraries of small molecules. Original cysteine-directed ABPP studies performed with fragment electrophiles provided initial covalent ligandability maps of human cells that included non-orthosteric sites on structurally and functionally diverse classes of proteins^24–26,39^. But, many of these fragment-cysteine interactions displayed low potency and selectivity.

The more recent introduction of structurally elaborated electrophilic compounds bearing defined stereocenters (stereoprobes) into ABPP workflows has furnished ligands that engage proteins with high stoichiometry, stereoselectivity, and site-specificity at low-µM concentrations in cells^21,27,35,40–42^. These features have in turn facilitated direct utilization of stereoprobes in cell biology experiments.

So far, efforts to diversify stereoprobes have mostly involved designing focused compound sets with alternative core structures^27,40,41,97^. Among these stereoprobe classes, the tryptoline acrylamides have been found to target non-orthosteric pockets on a wide range of proteins, including adaptors, RNA-binding proteins, and transcription factors^21,22,35,98^. Our studies described herein demonstrate that appendage modifications to the tryptoline core can serve as an additional source of molecular recognition capable of shaping and expanding the landscape of stereoprobe-protein interactions in cells. The distal location of these appendages relative to the acrylamide reactive group may suggest their contributions reflect direct improvements in reversible binding affinity, but we cannot exclude alternative modes of enhancing stereoprobe reactivity with proteins (e.g., orienting the acrylamide for optimal nucleophilic attack by cysteines)^99^. Distinguishing among these possibilities is further complicated by the generally negligible activity displayed by electrophilic stereoprobes with cysteine mutants of target proteins (**Fig. 4e** and refs. ^21,22,40,42,97^), which indicates only a modest amount of reversible binding affinity is required to promote substantial gains in covalent engagement.

The >100 stereoselective and chemoselective tryptoline acrylamide-cysteine interactions identified herein included both established sites of covalent drug action (e.g., orthosteric pockets on kinases like BTK, BLK, and TEC) and many other non-orthosteric liganding events. We surmised that this diverse dataset could be instructive for evaluating the performance of state-of-the-art ML co-folding tools aimed at predicting small molecule-protein interactions. In our analysis, we did not assess fine-grained structural features such as ligand poses, and we further acknowledge that noncovalent protein-ligand structure predictions may not fully capture the molecular basis for covalent liganding events. With these caveats in mind, we sought mainly to evaluate the ability of the ML model Boltz-2 to identify sites in proteins that had been experimentally defined by ABPP to bind small molecules with a well-defined SAR, as reflected by the proximity of the stereoprobe location in co-folded structure predictions to the liganded cysteine determined by ABPP.

The high success rate of Boltz-2 at assigning stereoprobe liganding events that occur at orthosteric pockets supports the potential for this model to accurately predict a specific subset of covalent liganding events in the proteome. However, Boltz-2 rarely predicted the location of the much larger subset of ligand interactions occurring at non-orthosteric sites. While we do not yet understand the full set of factors limiting the success of Boltz-2 with non-orthosteric liganding events, the strong tendency of this program to redirect stereoprobes to established orthosteric pockets points to a challenge with overfitting likely due to the predominance of orthosteric small molecule-protein structures in the Protein Data Bank (PDB) on which Boltz-2 is trained. A similar outcome was recently reported for a set of 17 proteins with structurally defined orthosteric and allosteric ligands, where Boltz-1/1x proved capable of predicting the binding poses of orthosteric ligands, but generally mislocated the allosteric ligands, often placing them at orthosteric sites^57^. These cases of “orthostery burnout” may point to the need to re-train ML co-folding tools on distinct datasets enriched in more diverse types of small molecule-protein interactions in order to avoid biased predictions derived from memorization^55^ of common orthosteric ligand poses.

When considering other ways to improve the computational prediction of non-orthosteric small molecule-protein interactions, we note that several of the liganded cysteines mapped by ABPP occurred at sites on proteins for which the basis for stereoselective engagement is difficult to glean from single-protein models. Such liganding events may occur at composite pockets formed by two or more proteins^44,97,100^ or in a context-dependent manner^22^ not captured by the structural data used in co-folding model training. Methods like ABPP that map protein-small-molecule interactions in native biological environments may therefore provide valuable case studies, if not more global datasets (see below), for both testing and training ML programs aimed at predicting the diverse types of non-orthosteric liganding events that occur on proteins in cells.

We verified several of the non-orthosteric liganding events mapped by cysteine-directed ABPP with recombinant proteins in heterologous cell systems and, for GRPEL1, with purified protein. While our initial attempts to determine a structure of GRPEL1 covalently bound to WX-71b have not been successful, the disruptive impact of this small molecule on the GPREL1:HSPA9 complexes is consistent with the interface-proximal location of the liganded C124 as revealed in a recent cryo-EM structure of the complex^85^. The acute impact of stereoprobe disruption of the GRPEL1:HSPA9 complex reverberated across multiple aspects of mitochondrial biology, including impairments of protein import and respiration, as well as induction of mitophagy. These data generally align with the molecular and cellular phenotypes associated with deleterious mutations in HSPA9 that cause EVEN-PLUS syndrome and congenital sideroblastic anemias^101–105^. The GRPEL1 ligands should further offer complementary chemical tools compared to direct HSPA9 inhibitors, which show variable degrees of selectivity across HSP70 paralogs and other proteins^106^.

The GRPEL1 ligands additionally increased ISR, although this effect was much more attenuated than the ISR induction observed with FCCP, a direct uncoupler of oxidative phosphorylation. Whether the more prolonged pharmacological perturbation of the mtHSP70 chaperone complex by GPREL1 ligands will cause greater mitochondrial and cell stress is an important question for future investigation. Even if such an outcome is observed, the temporary induction of mitophagy through partial and/or transient disruption of the GRPEL1:HSPA9 complex may have translational value for diseases caused by the aberrant accumulation of damaged mitochondria (e.g., Parkinson’s disease)^107–109^.

Projecting forward, we are encouraged by the number of cysteines showing preferred engagement by elaborated tryptoline acrylamide, which suggests that continued efforts to expand the ligandability of the human proteome may benefit from synthetically accessible modifications to established stereoprobe cores. The tryptoline scaffold, in this regard, provides straightforward routes for multiple sites of synthetic modification (**Fig 1a, c** and **Extended Data Fig. 1a**)^21,42^. Nonetheless, we should note that the elaborated tryptoline acrylamides examined herein are large in size (average MW 507 g/mol), suggesting that further optimization of their potency and selectivity for individual proteins of interest may require derivations rather than additions to the compound structures. We also believe that the burgeoning ligandability maps generated by ABPP have the potential to improve the performance of ML programs at predicting non-orthosteric small molecule-binding pockets in proteins. Such pockets are often shallow and dynamic, and these cryptic features present challenges for current ML programs trained mostly on more well-defined orthosteric sites in protein structures. Here, covalent ligandability maps, have the advantage of not only trapping small molecule binding events at dynamic pockets, but also precisely locating such events in proteins through the MS assignment of liganded cysteines. When such covalent small molecule-protein interactions further display the types of stereoselective and chemoselective SAR described herein, they would appear poised to offer robust opportunities for both testing and training ML programs. We also anticipate an important future contribution from additional chemical proteomic approaches that map the sites of reversible small molecule-protein interactions in cells^110^.

Finally, we continue to see the assignment of functional effects to stereoprobe-protein interactions as an important challenge. For interactions that occur at sites proximal to a biomolecular interface, as observed for the GRPEL1 ligands, experimentally testable hypotheses can be put forward (i.e., does the ligand disrupt or stabilize the biomolecular interaction?). But, many liganding events mapped by ABPP occur at non-orthosteric sites of unclear structural or functional impact. The future integration of covalent ligandability maps with growing databases of genotype (genetic variant)-phenotype relationships^111^ within the predicted^45^ or solved structures of proteins may provide a useful way to prioritize and guide the functional investigation of the nascent small molecule-protein interactions discovered by chemical proteomics.

## Methods

### Research ethics and regulations

All experiments were performed in compliance with protocols approved by The Scripps Research Institute Institutional Review Board.

### Cell lines and cell culture

All cell lines were obtained from American Type Culture Collection (ATCC). Ramos (ATCC, CRL-1596), THP-1 (ATCC, TIB-202), HEK293T (ATCC, CRL-3216), and HCT-116 (ATCC, CCL-247) cells were grown in Roswell Park Memorial Institute medium (RPMI; Ramos and THP-1), or Dulbecco’s modified Eagle medium (DMEM; HEK293T and HCT-116), supplemented with 10% fetal bovine serum (FBS), 2 mM L-glutamine, penicillin (100 U/mL), and streptomycin (100 μg/mL), in a humidified, 37 °C/5% CO_2_ tissue culture incubator.

### Reagents

Additional reagents, source and catalog numbers are available in Supplementary Tables 1-3.

### Cysteine-directed ABPP

#### *In situ* treatment and sample processing

Cysteine-directed ABPP was carried out as previously reported^21^ with slight modifications. In brief, Ramos cells (suspension cells: 10 mL at 3 x 10^6^ cells/mL; seeded at least 1 h before treatment) were treated with DMSO or 20 μM stereoprobes for 3 h. The cells were washed two times with chilled DPBS and immediately processed or stored at −80 °C. The cell pellets were resuspended in 500 μL of cold DPBS and lysed by sonication (2 × 15 pulses, 10% power output). The total protein content of whole-cell lysates was measured using a Pierce BCA protein assay kit. The samples were normalized to 2 mg/mL and 500 μL (1 mg of proteome) was treated with 5 μL of 10 mM IA-DTB (in DMSO) for 1 h at r.t. with occasional vortexing. Proteins were precipitated out of solution by the addition of chilled HPLC-grade methanol (600 μL), chloroform (200 μL) and water (100 μL), followed by vigorous vortexing and centrifugation at 16,000 × g for 10 min, to create a disk. Without disrupting the protein disk, both top and bottom layers were aspirated, and the protein disk washed with 1 mL of cold methanol and centrifuged at 16,000 × g for 10 min. The pellets were allowed to air dry (just enough get rid of methanol droplets) and then resuspended in 90 μL of denaturing/reducing buffer (9 M urea, 10 mM DTT, 50 mM triethylammonium bicarbonate (TEAB) pH 8.5). The samples were reduced by heating at 65 °C for 20 min, followed by the addition of 10 μL (500 mM) iodoacetamide for 30 min at 37 °C to cap free cysteines. The samples were then centrifuged at maximum speed (16,000 × g for 2 min) to pellet any insoluble precipitate and probe-sonicated once more to ensure complete resuspension, and then diluted with 300 μL of 50 mM TEAB pH 8.5 to reach a final urea concentration of 2 M. Trypsin (4 μL of 0.25 μg/μL in trypsin resuspension buffer with 25 mM CaCl_2_) was added to each sample and digested at 37 °C overnight. Digested samples were then diluted with 300 μL of wash buffer (50 mM TEAB pH 8.5, 150 mM NaCl, 0.2% NP-40) containing streptavidin-agarose beads (50 μL of 50% slurry/ sample) and were rotated at r.t. for 2 h. The samples were centrifuged (2,000 × g, 2 min) and the entire content transferred to BioSpin columns and washed (3 × 1 mL wash buffer this time containing 0.1% NP-40, 3 × 1 mL DPBS, 3 × 1 mL water). Enriched peptides were eluted twice from the beads with 200 μL of 50% acetonitrile with 0.1% formic acid and dried with a SpeedVac. IA-DTB-labeled and enriched peptides were resuspended in 100 μL EPPS buffer (200 mM, pH 8.0) with 30% acetonitrile, vortexed and water bath-sonicated. The samples were TMT labeled by the addition of 3 μL of 20 mg/mL (in dry acetonitrile) of corresponding TMT^10^plex tag, vortexed and incubated at r.t. for 1.5 h. TMT labeling was quenched with the addition of hydroxylamine (3 μL of a 5% solution in H_2_O) and incubated for 15 min at r.t. The samples were then acidified with 5 μL of formic acid, combined and dried using a SpeedVac. The samples were desalted with Sep-Pak and then high-pH-fractionated by HPLC (described in the following) into a 96-well plate and recombined into 12 fractions (total).

#### HPLC fractionation

The cysteine-directed ABPP samples were resuspended in 500 μL of 5% acetonitrile, 0.1% formic acid and fractionated with an Agilent HPLC system into a 96-deep-well plate containing 20 μL of 20% formic acid to acidify the eluting peptides, as previously reported.^21^ The peptides were eluted onto a capillary column (ZORBAX 300 Extend-C18, 3.5 μm) and separated at a flow rate of 0.5 mL min^−1^ using the following gradient: 100% buffer A from 0 min to 2 min, 0–13% buffer B from 2 min to 3 min, 13–42% buffer B from 3 min to 60 min, 42–100% buffer B from 60 min to 61 min, 100% buffer B from 61 min to 65 min, 100–0% buffer B from 65 min to 66 min, 100% buffer A from 66 min to 75 min, 0–13% buffer B from 75 min to 78 min, 13–80% buffer B from 78 min to 80 min, 80% buffer B from 80 min to 85 min, 100% buffer A from 86 min to 91 min, 0–13% buffer B from 91 min to 94 min, 13–80% buffer B from 94 min to 96 min, 80% buffer B from 96 min to 101 min, and 80–0% buffer B from 101 min to 102 min (buffer A, 10 mM aqueous NH_4_HCO_3_; buffer B acetonitrile). The plates were evaporated to dryness using a SpeedVac and peptides resuspended in 80% acetonitrile, with 0.1% formic acid, and combined to a total of 12 fractions (for example, fraction 1 = well 1A + 1B … 1H, fraction 2 = well 2A + 2B … 2H) (3 × 400 μL / column). Samples were dried with a SpeedVac, and the resulting 12 fractions were resuspended in buffer 5% acetonitrile, 0.1% formic acid in water and analyzed by MS.

#### Data processing

Cysteine engagement ratios (DMSO versus compound) were calculated for each peptide–spectra match by dividing each TMT reporter ion intensity by the average intensity for the DMSO channels. Peptide–spectra matches were then grouped based on protein ID and residue number (for example, NFU1 C210), excluding peptides with summed reporter ion intensities for the DMSO channels of <10,000 unless otherwise noted and a coefficient of variation for DMSO channels of >0.5. Replicate channels were grouped across each experiment, and average values were computed for each cysteine site. A variability metric was also computed across replicate channels, which equaled the ratio of the median absolute deviation to the average and was expressed as a percentage. A cysteine site was considered enantioselectively liganded if the variability corresponding to the probe leading to the highest blockade of IA-DTB did not exceed 20%, and at least one of the following additional criteria were met: (1) the average IA-DTB blockade by the probe was >66.7% and >2.5-fold that of its enantiomer, and either (a) the same probe led to <25% IA-DTB blockade of at least one other cysteine in the same protein (b) the same cysteine site was deemed liganded in another cysteine-directed ABPP experiment in this study or (c) the protein had 3-fold enantioselective enrichment as reported previously^21^.

### Protein-directed ABPP

#### *In situ* treatment

Protein-directed ABPP was carried out as previously reported^21^ with some modifications. Briefly, Ramos cells (suspension: 10 mL at 3 x 10^6^ cells/mL; seeded 1 h before treatment) or HCT-116 cells (adherent: 10 mL at 0.75 x 10^6^ cells/mL in a 10-cm dish; seeded 24 h before treatment) were treated with DMSO or non-alkyne competitor for 2 h. The cells were further treated with 5 μM alkyne probes for 1 h, then the cells were washed two times with chilled DPBS, and immediately processed or stored at −80 °C. The cell pellets were resuspended in 500 μL of cold DPBS and lysed by sonication (2 × 15 pulses, 10% power output). The total protein content of the whole-cell lysates was measured using a Pierce BCA protein assay kit. The samples were normalized to 2 mg/mL and 500 μL (1 mg of proteome) treated with 55 μL of click mix (30 μL of 1.7 mM TBTA in 4:1 t-BuOH:DMSO, 10 μL of 50 mM CuSO_4_ in H_2_O, 5 μL of 10 mM biotin-PEG4-azide (BroadPharm, catalog number BP-22119) in DMSO, 10 μL of freshly prepared 50 mM tris(2-carboxyethyl)phosphine in DPBS) for 1 h at r.t. with vigorous vortexing every 15 min. Proteins were precipitated out of solution by the addition of chilled HPLC-grade methanol (600 μL), chloroform (200 μL) and water (100 μL), followed by vigorous vortexing and centrifugation at 16,000 × g for 10 min, to create a disk. Without disrupting the protein disk, both top and bottom layers were aspirated, and the protein disk re-sonicated in 500 μL of methanol and centrifuged at 16,000 × g for 10 min. After complete aspiration of the methanol, protein pellets were frozen at −80 °C or immediately resuspended in 500 μL of freshly made 8 M urea in DPBS, followed by the addition of 10 μL of 10% SDS, and pellet was sonicated to clarity. The samples were reduced with 25 μL of 200 mM dithiothreitol (DTT) at 65 °C for 15 min, followed by alkylation with 25 μL of 400 mM iodoacetamide at 37 °C for 30 min. The samples were quenched with 130 μL of 10% SDS, transferred to a 15-mL tube, and the total volume was brought to 6 mL with DPBS (0.2% final SDS). Washed streptavidin beads (Thermo, catalog number 20353; 100 μL of 50% slurry/sample) was then added and the probed-labeled protein enriched for 1.5 h at r.t. with rotation. After incubation, the beads were pelleted (2 min × 2,000 × g) and washed with 0.2% SDS in DPBS (2 × 10 mL), DPBS (1 × 5 mL, then transferred to a protein low-bind Eppendorf safe-lock tube), HPLC water (2 × 1 mL) and 200 mM 4-(2-hydroxyethyl)-1-piperazinepropanesulfonic acid (EPPS; 1 × 1 mL) at r.t. Enriched proteins were digested on-bead overnight with 200 μL of trypsin mix (2 M urea, 1 mM CaCl_2_, 10 μg/mL trypsin, 200 mM EPPS, pH 8.0). The beads were spun down, supernatant was collected and 100 μL of acetonitrile (30% final) was added, followed by 6 μL of 20 mg/mL (in dry acetonitrile) of the corresponding TMT^16^plex tag for 1.5 h at r.t. with vortexing every 30 min. TMT labeling was quenched by the addition of hydroxylamine (6 μL 5% solution in H_2_O) and incubated for 15 min at r.t. Samples were then acidified with 5 μL 100% formic acid, combined and dried with a SpeedVac. Samples were desalted with a Sep-Pak column and then high-pH spin column fractionated into ten fractions using peptide desalting spin columns (as described below).

#### High-pH spin column fractionation

High-pH fractionation was carried out as previously reported^21^ using peptide desalting spin columns (Thermo 89852). Samples were resuspended in 300 μL of buffer A (5% acetonitrile, 0.1% formic acid) by water bath sonication and bound to the spin columns. Bound peptides were then washed once with water, once with 5% acetonitrile in 10 mM NH_4_HCO_3_, and eluted into 30 fractions with an increasing gradient of acetonitrile. Every tenth fraction was combined (for example, 1, 10, and 30) and dried with a SpeedVac to dryness. Each of the resulting ten fractions was resuspended in buffer A (5% acetonitrile, 0.1% formic acid) and analyzed by MS.

*In vitro* treatment with SAM and sample processing. HEK293T cell pellets were lysed in 500 μL of DPBS. Samples were normalized to 2 mg/mL in 500 μL (1 mg of proteome) and 5 μL of 100X SAM was added before being treated with 1 μL of 500X probe for 1 h at r.t. Cell lysates were then treated with click mix and processed for MS analysis as described for *in situ* treatment above.

#### Data processing

Enrichment ratios (probe versus probe) were calculated for each peptide– spectra match by dividing each TMT reporter ion intensity by the sum intensity for all the channels. Peptide–spectra matches were then grouped based on protein ID and (excluding peptides with summed reporter ion intensities <10,000) coefficient of variation of >0.5, and <2 distinct peptides.

### IP-MS

#### *In situ* treatment and sample processing

Cells stably expressing FLAG epitope-tagged protein of interest (HCT-116: 7.5 × 10^6^ or 5 × 10^6^ cells in a 10-cm dish overnight) were treated with DMSO or 20 μM stereoprobes for 4 h. The cells were collected and washed twice with cold DPBS then frozen at −80 °C. Ice-thawed pellets were resuspended in 1 mL of IP lysis buffer (50 mM EPPS pH 8.0, 150 mM NaCl, 1% NP-40, 10% glycerol) supplemented with EDTA-free complete protease inhibitor. Lysing was achieved by rotating the samples at 4 °C for 1 h. The lysate was clarified by spinning at 16,000 × g for 5 min and the supernatant was assayed for total protein using BCA reagent. Lysate (1-3 mg) was mixed with 40 μL of prewashed anti-FLAG magnetic beads for 2-4 h at 4 °C with rotation. The samples were washed three times with IP wash buffer (25 mM EPPS pH 8.0, 150 mM NaCl, 0.2% NP-40) and once with 50 mM EPPS pH 8.0. Enriched proteins were eluted off the beads by boiling with 40 μL of 8 M urea in 50 mM EPPS pH 8.0 at 65 °C for 10 min, and the supernatant was collected with a magnetic stand into new tubes. Sample was then reduced with 2 μL of 200 mM DTT for 15 min at 65 °C and alkylated with 2 μL of 400 mM iodoacetamide for 30 min at 37 °C. Samples were further diluted to 2 M urea with 115 μL of 50 mM EPPS pH 8.0 and trypsin (4 μL of 0.25 μg/µL in 50mM EPPS with 25 mM CaCl_2_) was added to each sample and digested at 37 °C overnight. Acetonitrile (75 μL) was added to each sample and TMT labeled and desalted as described in ‘Protein-directed ABPP’. Combined samples were high-pH spin column fractionated and combined into three fractions.

#### Data processing

Processing and analysis by MS followed the protein-directed ABPP protocol described above with the following changes. Protein signals were normalized to the FLAG epitope-tagged protein pulled down within each treatment group and then to DMSO signals across treatment groups, unless otherwise noted.

### Pulse-chase SILAC

#### *In situ* treatment and sample processing

Cells were cultured in light or heavy DMEM minus L-Lysine and L-Arginine (Life Technologies 88364) supplemented with 10% dialyzed FBS (Omega Scientific FB-03), 2 mM L-glutamine, penicillin (100 U/mL), streptomycin (100 μg/mL), and 100 μg/mL [^13^C_6_, ^15^N_4_] L-Arginine-HCl and [^13^C_6_, ^15^N_2_] L-Lysine-HCl (Sigma 608033 and 608041) or L-Arginine-HCl and L-Lysine-2HCl (Sigma A6969 and L5751) for 2 weeks before treatment. HCT-116 cells (adherent: 10 mL at 0.75 × 10^6^ cells/mL in a 10-cm dish; seeded in light or heavy amino acid media 24 h before treatment) were pre-treated with DMSO or 20 µM stereoprobe for 4 h, switched to heavy amino acid media containing 5 µM stereoprobe for an additional 8 h, and then the cells were washed twice with chilled DPBS, and stored at −80 °C. Light-only and heavy-only controls were handled identically but maintained in their respective media throughout (no media exchange).

Mitochondria were isolated from cell pellets using Thermo Scientific Mitochondria Isolation Kit for Cultured Cells (89874). Mitochondria were then resuspended in 80 µL of DPBS with 150 mM NaCl and 2% CHAPS and lysed by vortexing. Protein content was measured using a standard Micro BCA^TM^ assay (Thermo Scientific) and a volume corresponding to 50 µg was transferred to a new low-bind Eppendorf tube (containing 48 mg urea) and brought to 100 µL total volume.

Samples were reduced with DTT (5 µL 200 mM stock in H_2_O, 10 mM final concentration) and incubated at 65 °C for 15 minutes, then alkylated with iodoacetamide (5 µL 400 mM stock in H_2_O, 20 mM final concentration) incubated at 37 °C for 30 min in the dark. Samples were precipitated with the addition of ice-cold MeOH (250 µL), CHCl_3_ (50 µL), and H_2_O (200 µL), and vortexed and centrifuged (16,000 × g, 10 minutes, 4 °C). The supernatant above and below the protein disc was aspirated and an additional 1 mL of ice-cold MeOH was added. The samples were again vortexed and centrifuged (16,000 × g, 10 min, 4 °C), the supernatant aspirated and the protein pellets were allowed to air dry and be stored at -80 °C or proceeded to resuspension and digestion. Samples were resuspended in 100 µL EPPS buffer (200 mM pH 8.0) using a probe sonicator (1 × 10 pulses, 10% output). Proteomes were first digested with Lys-C (2 µg/µL in HPLC grade water, 1 µL per sample) for 2 h at 37 °C then 2 µg of trypsin (1 mM CaCl_2_ final concentration) was added and samples were incubated at 37 °C overnight. Peptide concentrations were estimated using Micro BCA^TM^ assay (Thermo Scientific), and a volume corresponding to 25 µg was transferred to a new low-bind Eppendorf tube. Samples were diluted with acetonitrile (final concentration 30%) and then TMT labeled with 6 µL of corresponding TMT tag (20 mg/mL), vortexed, and incubated at room temperature for 1.5 h. TMT labeling was quenched with the addition of hydroxylamine (6 µL 5% solution in H_2_O) and incubated for 15 minutes at r.t. Samples were then acidified with 5 µL formic acid and an volume corresponding to 10 µg (per channel) was combined and dried using a SpeedVac. Samples were desalted via Sep-Pak and then high-pH spin column fractionated as described above.

#### Data Processing

Processing and analysis by MS followed the protein-directed ABPP protocol described above with the following changes. Each data set was independently searched with light and heavy parameter files; for the light search, all other amino acids were left at default masses; for the heavy search, static modifications on lysine (+8.0142 Da) and arginine (+10.0083 Da) were specified. Channels were averaged across replicates, light only values were subtracted from heavy peptide values and normalized to WT_DMSO or C124A_DMSO channels. Enantioselectivity and site-specificity ratios were subsequently calculated to identify proteins with reduced protein abundance in the mitochondria correlating with the stereoselective and site-specific engagement of GRPEL1_C124 by WX-71b.Finally, Human MitoCarta3.0 was used for annotation of mitochondrial proteins^112^. Submitochondrial localizations were also annotated based on MitoCarta3.0.

### TMT LC–MS analysis

Samples were analyzed by liquid chromatography tandem mass-spectrometry using an Orbitrap Eclipse Tribrid Mass Spectrometer (Thermo Scientific) or an Orbitrap Fusion Tribrid Mass Spectrometer (Thermo Scientific) coupled to an UltiMate 3000 Series Rapid Separation LC system and autosampler (Thermo Scientific Dionex). The peptides were eluted onto an EASY-Spray HPLC column (Thermo ES902, ES903) using an Acclaim PepMap 100 (Thermo 164535) loading column, and separated at a flow rate of 0.25 μL/min. Data were acquired using a data-dependent acquisition with a 2.5 second cycle time.

For Orbitrap Eclipse Tribrid Mass Spectrometer, the scan sequence began with an MS1 master scan (Orbitrap analysis, resolution 120,000, 400-1700 m/z, RF lens 30%, standard automatic gain control, maximum injection time 50 ms, centroid mode) with dynamic exclusion enabled (repeat count 1, duration 30 s). The top precursors were then selected for MS2 analysis within a 2.5 duty cycle. MS2 analysis consisted of quadrupole isolation (isolation window 0.7) of precursor ion followed by High energy collision-induced dissociation (HCD 36%) detected via rapid scan in ion trap. (AGC 180%, max injection time 120 ms). The top 10 SPS precursors were then selected for MS3 and subjected to HCD fragmentation (55% collision energy) and detection in orbitrap (30k resolution, turboTMT enabled, 500% normalized AGC target, 120 ms max injection time).

For Orbitrap Fusion Tribrid Mass Spectrometer, the scan sequence began with an MS1 master scan (Orbitrap analysis, resolution 120,000, 400-1700 m/z, RF lens 60%, custom automatic gain control normalized AGC target 50%, maximum injection time 50 ms, centroid mode) with dynamic exclusion enabled (repeat count 1, duration 15s). The top precursors were then selected for MS2 analysis within a 2.5 duty cycle. MS2 analysis consisted of quadrupole isolation (isolation window 0.7) of precursor ion followed by collision-induced dissociation (collision energy 35%) detected via rapid scan in ion trap. (normalized AGC target 180%, max injection time 120 ms, centroid). The top 10 SPS precursors were then selected for MS3 and subjected to HCD fragmentation (55% collision energy) and detection in orbitrap (50k resolution, 300% normalized AGC target, 120 ms max injection time, centroid).

Raw files were uploaded to the Integrated Proteomics Pipeline (IP2, version 6.0.2) available at http://ip2.scripps.edu/ ip2/mainMenu.html, and MS2 and MS3 files were extracted from the raw files using RAW Converter (version 1.1.0.22, available at http://fields.scripps.edu/rawconv/) and searched using the ProLuCID algorithm using a reverse concatenated, non-redundant variant of the Human UniProt database (release 2016-07). Cysteine residues were searched with a static modification for carboxyamidomethylation (+57.02146 Da). For cysteine-directed ABPP, a dynamic modification for IA-DTB labeling (+398.25292 Da) was included with a maximum number of two differential modifications per peptide. N termini and lysine residues were also searched with a static modification corresponding to the TMT tag (+229.1629 Da for 10plex and +304.2071 Da for 16plex). Peptides were required to be at least six amino acids long. ProLuCID data were filtered through DTASelect (version 2.0) to achieve a peptide false-positive rate below 1%. The MS3-based peptide quantification was performed with reporter ion mass tolerance set to 20 ppm with the Integrated Proteomics Pipeline (IP2).

### Boltz-2 Analysis

To acquire a dataset of Boltz-2-derived protein-ligand structures, relevant UniProt accession numbers and ligand SMILES strings were parsed from processed chemical proteomics data reported in **Supplementary Dataset 1**, Tabs 4 and 2, respectively, to generate Boltz-2 submission scripts. MSAs were prepared using MMseqs2 via the ‘--use_msa_server’ argument provided by Boltz-2^113^. Predictions for every unique protein-ligand pair were run with 10 diffusion samples (‘--diffusion_samples 10’) and with inference time potentials (via ‘--use_potentials’) using Boltz-2.2.0. All predictions were performed without restraints. Predictions used in the generation of Extended Data Figure 2c and 2d were performed using the same parameters and input data with Boltz-2.2.1. All predictions were conducted using RTX A6000 GPUs available on-site at Scripps Research.

Ligand positions were evaluated using the shortest through-space distances between any atom in the ligand and any atom in the cysteine liganded in cysteine-directed ABPP experiments. In cases where multiple cysteines are present on the tryptic peptide associated with the liganding event, distances between all cysteines were calculated. Proximal pLDDT calculations were performed using data associated with all residues harboring any atom within 5 Å of the ligand or the liganded cysteine. When relevant, an arbitrary 5 Å distance threshold was used to discriminate between correct and incorrect predictions. The relative orientation of the ligand, the ligand acrylamide, or the cysteine sulfhydryl was not considered. To complete our structural analysis, we evaluated the ligand geometry and the ligand-protein interface using PoseBusters 0.4.6 and evaluated the protein structure using Phenix 1.21.2^58,114^. Ligand stereochemistry was determined using RDKit 2024.09.5^115^.

To identify orthosteric sites on proteins, a complete list of orthosteric annotations for each of the proteins in our analysis was prepared by collating “Active site” and “Binding site” features and their associated ‘ligand’, ‘namè, ‘location’, ‘notè, and ‘description’ fields for each protein’s entry in UniProt. For annotations with residue ranges, the center residue was considered. Mutagenesis annotations were not included in the analysis. An extensive list of metal cofactors was also excluded. Metal cofactors annotated as “catalytic”, such as the zinc cofactor in LACC1, were reintroduced into the analysis. Distance calculations were performed using the same approach described for ligand-cysteine distances. To extend this analysis, we generated orthostery classifications for each of the liganded cysteines in the proteins in our evaluation. We sought to faithfully represent the more complex relationship between the orthosteric site and the liganded cysteine, for which many edge cases exist that were not captured by 5 Å cutoffs. Using 𝑑_*LC*_ as the minimum distance between the modeled ligand and the cysteine (Å), 𝑑_*LO*_ as the minimum distance between the modeled ligand and the nearest orthosteric annotation (Å), and 𝑑_*CO*_ as the minimum distance between the known liganded cysteine and the nearest orthosteric site (Å), we designed the following per-prediction classification scheme for each liganded cysteine. Cysteine categories are named for their relationship to the nearest orthosteric annotation:

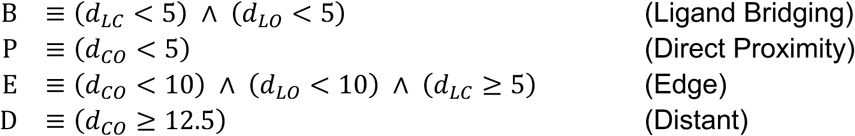

Using these categories as predicates, we classified each cysteine as orthosteric, non-orthosteric, or ambiguous:

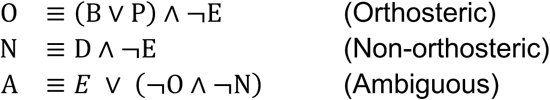

All calculations and subsequent analysis are available online at https://github.com/cravattlab/Hayward_2025_Boltz_analysis. In addition to previously mentioned tools, data preparation, analysis, and visualization were performed using tools from the SciPy ecosystem and Biopython^116–119^.

### Cloning and mutagenesis

All full-length plasmids were obtained from either OriGene in pCMV6 vector with a C-terminal Myc-DDK (FLAG) epitope tag or from GenScript in pcDNA3.1-C-(k) DYK (FLAG), as shown in Supplementary Table 2. Mutagenesis was carried out using a Q5 site-directed mutagenesis kit (New England BioLabs, E0554S), using the primers shown in Supplementary Table 3.

### Generation of NCKAP1L, FBL, and GRPEL1 stable cell lines

Full-length NCKAP1L, FBL, or GRPEL1 (WT and C-to-A/S mutant) carrying C-terminal epitope tags (FLAG or V5) were generated using Gateway cloning into lentiviral vectors using primers shown in Supplementary Table 3. To generate lentivirus, 3 × 10^5^ HEK293T cells were seeded in 6-well plates overnight, in 2 mL of DMEM. Protein-encoding viral vectors, lentiviral packaging vector (pCMV-dR8.91) and envelope vector (VSV-G) were mixed in a 6:6:1 ratio in OPTI-MEM medium, and 3 μg of PEI (1 mg/mL, Polysciences) was added per μg total plasmids. The DNA:PEI complex was incubated at r.t. for 20 min and the complex added dropwise to the HEK293T cells. Media were replaced with fresh DMEM (Corning) with 30% FBS plus pen-strep and 2 mM glutamine, 6-16 h post transfection. The virus was collected at 48 h post transfection and filtered through a 0.45-μm syringe filter (Millipore) to eliminate floating cells. For transduction, 0.1-0.5 million cells were mixed with 100-500 µL of viral supernatant in 1-2 mL supplemented with 6 μg/mL polybrene. Cells were spin-infected at 930 × g at 30 °C for 1-2 h and incubated for 24 h at 37 °C. Selection was initiated with 2 μg/mL puromycin (NCKAP1L), 7 μg/mL blasticidin (FBL), or 1 mg/mL geneticin (GRPEL1) for 1-2 weeks. Selected pools were characterized by immunoblots (FLAG or V5).

### Generation of GRPEL1 CRISPR/Cas9 knockout cells

Stable knockout cells were generated by the transduction of cells with LentiCRISPR v2-Blast carrying sgGRPEL1. Briefly, sgRNA (sgGRPEL1-05_sense5′-CACCGAGCTGAAGGAGACTGTGGTG) were annealed and cloned into LentiCRISPR-v2 Blast^120^. Viral supernatants were generated as described above, and GRPEL1 stable cell lines were transduced and selected with 5 μg/mL of blasticidin for one week. Selected stable pools were evaluated by immunoblot with anti-GRPEL1 antibody.

### Protein purification

#### Plasmid/construct design

Plasmids encoding full-length genes for human HSPA9 and human GRPEL1 were prepared according to previously published protocols^85^, except for using WT-HSPA9 in place of R126W-HSPA9. The C124A mutant of GRPEL1 (C124A-GRPEL1) was generated using Gibson assembly. All oligonucleotides were ordered from Integrated DNA Technologies and all plasmid sequences were confirmed by GENEWIZ. Verified plasmid sequences were transformed into the *E. coli* LOBSTR^121^ strain for recombinant protein expression.

#### Recombinant protein expression, purification, and complex formation

A pETDuet-1 vector encoding for HSPA9 was transformed into the *E. coli* LOBSTR^121^ strain and grown in LB media containing 50 µg/mL ampicillin and 50 µM ZnCl_2_ at 37 °C until the OD_600_ reached 1.0. Protein expression was induced by the addition of 300 µM isopropyl β-D-1-thiogalactopyranoside (IPTG) and allowed to express for 4 h at 37 °C. Cells were centrifuged, collected, and resuspended in buffer A (20 mM HEPES (pH 8), 100 mM NaCl, 50 µM ZnCl2, 5 mM β-mercaptoethanol (BME), 10 mM imidazole, 1 mM MgCl_2_, 1 mM CaCl_2_, 1 µg/mL polyethyleneimine (PEI), 1 mM phenylmethylsulfonyl fluoride (PMSF), 5 mM benzamidine, 10 µM leupeptin, 1 µM pepstatin A, and 2 µg/mL aprotinin) with a small amount of DNase I (Millipore Sigma). Cells were lysed using a Branson SFX550 sonifier on ice. Lysed cells were centrifuged at 20,000 × g for 30 min at 4 °C, and the clarified supernatant was loaded onto a 5 mL HisTrap HP column (Cytiva) pre-equilibrated with buffer A at 1 mL/min using a peristaltic pump. The column was washed with buffer B (20 mM HEPES (pH 8), 100 mM NaCl, 50 µM ZnCl_2_, 5 mM BME, 40 mM imidazole, 1mM ATP, 1mM MgCl_2_ and 0.1% Triton X-100 (Fisher Scientific)) until protein was no longer observed in the elution, as visualized by mixing with Bradford reagent. The target protein was eluted using 5 column volumes (CVs) of buffer C (20mM HEPES (pH 8), 100mM NaCl, 50 µM ZnCl_2_, 5mM BME, and 300mM imidazole). The eluates were pooled, concentrated, and applied to a Superdex 200 Increase 10/300 GL (Cytiva) that had been pre-equilibrated with buffer D (20 mM HEPES (pH 8), 100 mM NaCl, and 0.5 mM TCEP). Peak fractions were collected and analyzed using SDS-PAGE. The purest fractions were combined, concentrated to 1.7 mg/mL using a 10 kDa MWCO centrifugal filter (Millipore Sigma), and flash frozen in liquid nitrogen (LN_2_) for storage.

WT-GRPEL1 was expressed and purified according to previously published protocols^85^. Briefly, WT-GRPEL1, with C-terminal tandem Strep-Tag II and hexahistidine tags, was expressed in the *E. coli* LOBSTR^121^ strain. Following cell lysis, WT-GRPEL1 was purified using Nickel affinity chromatography (Ni Sepharose High-Performance resin (Cytiva)), Strep affinity chromatography (Strep-Tactin Superflow resin (IBA Lifesciences)), and size exclusion chromatography (Superdex 200 Increase 10/300 GL (Cytiva)). The cleanest fractions were combined, concentrated to 0.8 mg/mL using a 3 kDa MWCO centrifugal filter (Millipore Sigma), and flash frozen in LN_2_ for storage.

A pRSFDuet-1 vector encoding for the C124A-GRPEL1 mutant was transformed into the *E. coli* LOBSTR^121^ strain. Recombinant protein expression and purification of C124A-GRPEL1 were performed as previously described^85^ for WT-GRPEL1. Following cell lysis, C124A-GRPEL1 was purified using Nickel affinity chromatography (Ni Sepharose High-Performance resin (Cytiva)), Strep affinity chromatography (Strep-Tactin Superflow resin (IBA Lifesciences)), and size exclusion chromatography (Superdex 200 Increase 10/300 GL (Cytiva)). The cleanest fractions were combined, concentrated to 1.32 mg/mL using a using a 3 kDa MWCO centrifugal filter (Millipore Sigma), and flash frozen in LN_2_ for storage.

Preparation of the HSPA9:GRPEL1 complex was performed as previously described^85^, except for using WT-HSPA9 in place of R126W-HSPA9. Fractions consisting of HSPA9:GRPEL1 complex, as visualized by SDS-PAGE, were concentrated to ∼0.5 mg/mL and were flash frozen in LN_2_ for storage.

### Gel-ABPP with recombinant proteins

#### Transient transfection

HEK293T cells (3 × 10^5^) were seeded in a six-well plate overnight and transfected with 1-2 μg of FLAG-epitope tag plasmids (depending on difficulty of expression) using polyethylenimine (PEI) at a ratio of 1:3 (DNA:PEI) for 48 h. The cells were treated with alkyne probe only, for 1 h or with competitor probe for 1-2 h, followed by alkyne probe for a further 1 h. The cells were then collected and washed three times with chilled Dulbecco’s phosphate-buffered saline (DPBS). The cell pellets were resuspended in 200 μL of cold DPBS and lysed by sonication (2 × 15 pulses, 10% power output). The total protein content of the whole-cell lysates was measured using a Pierce BCA protein assay kit. Samples were normalized to 1 mg/mL and a 50 μL sample was treated with 6 μL of click mix (3 μL of 1.7 mM tris((1-benzyl-4-triazolyl)methyl) amine (TBTA) in 4:1 t-BuOH:DMSO, 1 μL of 50 mM CuSO_4_ in H_2_O, 1 μL of 1.25 mM rhodamine–polyethylene glycol–azide in DMSO, 1 μL of freshly prepared 50 mM tris(2-carboxyethyl)phosphine in DPBS) for 1 h at r.t. with vigorous vortexing every 15 min. The click reaction was quenched by the addition of 18 μL of 4X SDS gel loading buffer, and the samples were resolved on SDS–PAGE gels and imaged by in-gel fluorescent scanning using a BioRad imager with Image Lab software version 6.1.

#### Stable cell lines

Cells stably expressing epitope-tagged protein of interest (HCT-116 and HEK293T: 1 x 10^6^ cells in a 6-well plate overnight; THP-1: 3 mL at 3 x 10^6^ cells/mL; seeded 1 h before treatment) were treated with alkyne probe only, for 1 h or with competitor probe for 1-2 h, followed by alkyne probe for a further 1 h. The cells were then collected and washed three times with chilled DPBS. The cell pellets were resuspended in 100-200 μL of IP lysis buffer (50 mM EPPS pH 8.0, 150 mM NaCl, 1% NP-40, 10% glycerol, 1 mM MgCl_2_/1:5,000 benzonase (only for FBL)) and cell pellets were lysed by sonication (2 × 15 pulses, 10% power output). The lysate was clarified by spinning at 16,000 × g for 5 min. and the supernatant was assayed for total protein using BCA reagent. Protein concentrations for all the samples were adjusted to 1 mg/mL, 15 µL of proteome was taken for the input, 5 µL of 4× SDS gel loading buffer was added, and the samples were heated at 95 °C for 10 min. The remaining lysate (150-500 µg) was mixed with 15 μL of prewashed anti-FLAG or anti-V5 magnetic beads at 4 °C with rotation for 1-2 h or overnight, respectively. The samples were washed three times with IP wash buffer (25 mM EPPS pH 8.0, 150 mM NaCl, 0.2% NP-40) and once with DPBS. Beads were resuspended in 25 µL of DPBS for on-bead click reaction with 3 μL of click mix (1.5 μl of 1.7 mM tris((1-benzyl-4-triazolyl)methyl) amine (TBTA) in 4:1 t-BuOH:DMSO, 0.5 μL of 50 mM CuSO_4_ in H_2_O, 0.5 μL of 1.25 mM rhodamine–polyethylene glycol–azide in DMSO, 0.5 μL of freshly prepared 50 mM tris(2-carboxyethyl)phosphine in DPBS) for 1 h at r.t. with vigorous vortexing every 15 min. The click reaction was quenched by the addition of 9 μL of 4X SDS gel loading buffer, and proteins were eluted off the beads by boiling for 10 min. The supernatant was collected with a magnetic stand into new tubes and input and IP samples were resolved on SDS–PAGE gels and IP samples were imaged as described above.

#### Gel-ABPP with purified GRPEL1

Purified GRPEL1 was diluted to 0.5 µM in cold DPBS (total 50 µL) and samples were treated with 10 µM alkyne probe for 2 h at r.t. with pipet mixing every 30 min. Click reaction and SDS-PAGE / in-gel fluorescent scanning were performed as described above.

#### *In vitro* Gel-ABPP for FBL

Cell pellets were lysed by sonication (2 × 15 pulses, 10% power output) in DPBS. Samples were normalized to 1 mg/mL and 500 μl of lysate was treated with 1 μl of 500× alkyne probe for 1 h at r.t. After treatment, 15 µL of proteome was taken for the input, 5 µL of 4× SDS gel loading buffer was added, and the samples were heated at 95 °C for 10 min. The remaining lysate was mixed with 2X IP lysis buffer (containing MgCl_2_ and benzonase) and 15 μl of prewashed anti-V5 magnetic beads overnight at 4 °C with rotation. The samples then were washed and processed as described above for stable cell lines.

For SAM/SAH gel-ABPP, samples were normalized to 1 mg/mL and 5 µL of 100x SAM or SAH was added to 500 μl of lysate 10 min before alkyne treatment.

### NCKAP1L Distance to C338

NCKAP1L residue distances from C338 were calculated using the package ’bio3d’ (v2.4-5)^122–124^ for R (v4.3.2).

### Intact MS

Purified recombinant GRPEL1 (3.5 µM) in DPBS was treated with DMSO or stereoprobes. Proteins were concentrated to 10 µM and desalted into water (Zeba Spin Desalting column, 40K, MWCO). For ESI-TOF analysis, samples were injected into Agilent 6230 TOF LC/MS with a dual AJS ESI ion source (Agilent Technologies) equipped with PLRP-S 1000 Å, 5 µm column (PL1912-1502). Samples were separated and eluted by 30-minute gradient of 5 % - 95 % acetonitrile in water containing 0.1 % formic acid (0.3 mL/min). The instrument operated on positive ion mode, with the following settings: VCap = 4000, Nozzle voltage = 2000, fragmentor = 250, skimmer1 = 65, octopoleRFpeak = 750, gas temperature = 350 °C, gas flow = 8, nebulizer = 40, sheath gas temp = 400 °C, and sheath gas flow = 12. LC/MS data was processed using MassHunter BioConfirm software (Agilent) to calculate deconvoluted mass and peak intensities of each protein species.

### MTS-EGFP mitochondrial fluorescence import assay

#### Generation of MTS-EGFP cells

MTS-EGFP (a gift from the Wiseman Lab) was cloned into pCW57.1 (Addgene #41393) by Gateway cloning. Viral supernatants were generated as described above, and cells were transduced and selected with 1 μg/mL of puromycin for one week. Selected stable pools were evaluated by fluorescence microscopy.

#### MTS-EGFP import assay

Cells (7 × 10^5^) stably expressing doxycycline-inducible MTS-EGFP were seeded in 35 mm poly-D-lysine coated dishes (Matek P35GC-1.5-14-C) overnight and cotreated with 0.5 μg/mL doxycycline and 20 µM stereoprobes 8 h prior to microscopy. The cells were then stained with 200 nM MitoTracker Deep Red FM (Cell signaling 8778) for 30 min in pre-warmed DMEM medium. Cells were washed with DPBS and incubated in DMEM during measurements. The Zeiss LSM 880 Airyscan Confocal laser scanning microscope with a Plan-Apochromat 63x NA 1.4 objective was used to take live-cell images with 488 nm excitation 493-598 nm emission for EGFP and 633 nm excitation 638-755 nm emission for MitoTracker Deep Red FM. Pixel resolution was 0.085 microns/pixel. For each treatment condition, 10 images per biological replicate were analyzed using Imaris 10.2 colocalization module (Oxford Instruments). The co-localization between MTS-EGFP and MitoTracker Deep Red FM was determined using Mander’s coefficients to get an estimate to the amount of protein import into the mitochondria.

### Seahorse Assay

Mitochondrial respiration parameters were measured using an XF96 Extracellular Flux Analyzer (Seahorse Bioscience) according to the manufacturer’s protocol. In brief, 10,000 cells/well were plated on an XFe96/XF plates (pre-coated with poly-D-lysine) in their standard growing media. After 48 h, cells were treated with DMSO or stereoprobe and incubated at 37 °C, 5% CO_2_ for 8h. Seahorse media (serum-free DMEM (Sigma-Aldrich D5030) containing 50 mM glucose, 10 mM sodium pyruvate, 2 mM glutamine and 10 mM HEPES pH 7.4) was used to wash and remove the standard growing media from the cell plate before a mitochondrial stress test was performed consisting of 3 min cycles of mixing and 2 min cycles of measurements. Basal respiration measurement was followed by injections of oligomycin (2 µM), FCCP (1 µM) and rotenone/antimycin (2 µM each).

### Integrated Stress Response Immunoblots

HCT-116 cells (8 × 10^5^) were seeded in 6-well plates overnight before being treated with DMSO or 5 μM stereoprobes for 8 h. The cells were collected and washed with cold DPBS and immediately processed or stored at −80 °C. Pellets were resuspended in IP lysis buffer (50mM EPPS pH 8.0, 150mM NaCl, 1% NP-40, 10% glycerol) supplemented with EDTA-free complete protease inhibitor and PhosSTOP and lysed by sonication (2 × 15 pulses, 10% power output). Lysates were cleared by centrifugation (5 min, 16,000 × g, 4 °C), and the total protein content of the lysates was measured using a Pierce BCA protein assay kit. Protein concentrations for all the samples were adjusted to ∼2 mg/mL, 4× SDS gel loading buffer was added (25 µL to 75 µL of proteome), and the samples were heated at 95 °C for 10 min. The proteins were resolved by SDS–PAGE and immunoblotted with the respective antibody in 3% BSA in TBST.

### Mitophagy flux mt-mKeima assay

#### Generation of mt-mKeima cell lines

To generate recombinant retro- and lentiviruses, 2.5 x 10^6^ HEK293T LentiX cells (Takara Bio) were plated in a 10 cm cell culture dish. For mt-mKeima encoding lentivirus, LentiX cells were co-transfected with 0.82 pmol pHAGE-mt-mKeima (Addgene #131626), 0.65 pmol psPAX2 (Addgene #12260), and 0.36 pmol pMD2.G (Addgene #12259). For TagBFP-Parkin encoding retrovirus, LentiX cells were transfected with 0.82 pmol pBMN BFP-Parkin (Addgene #186221), 0.65 pmol pBS-CMV-gagpol (Addgene #35614), and 0.36 pmol pMD2.G (Addgene #12259). The plasmids were diluted in OptiMEM I Reduced Serum Media (ThermoFischer Scientific), before a volume of XtremeGene HD Transfection Reagent (Roche) corresponding to 4 times the combined mass of all three plasmids was added. The resulting transfection mix was gently vortexed and incubated at room temperature for 15 minutes. The media in the cell culture plate was replaced with fresh growth media, and the transfection mix was added dropwise to the cell culture plate. After 16 hours, the media was exchanged for fresh growth medium. The virus was collected at 72 h post transfection and filtered through a 0.45-μm PES membrane filter (Sartorius) to eliminate floating cells.

For transduction, 80,000 HCT-116 cells were plated into 12-well plates overnight. On the day of, the media was exchanged for full growth medium containing 10 µg/mL Polybrene (Merck) and different volumes of lentivirus-containing supernatant were added. After 48 hours the media was aspirated and cells were expanded into 6-well plates. Cells positive for TagBFP fluorescence and mt-mKeima fluorescence after excitation at 488 nm were enriched by fluorescence activated cell sorting (FACS) using a CytoFLEX SRT Benchtop Cell Sorter (Beckmann Coulter).

#### Mitophagy assay

A matrix-targeted version of the fluorescent protein mKeima was used to determine the fraction of live cells undergoing mitophagy. mKeima is highly acid sensitive and the excitation spectrum shifts upon acidification.

HCT-116 mt-mKeima BFP-Parkin cells (1 x 10^5^) were plated in 24-well plates in duplicates per condition. On the day of, cells were treated with DMSO, 10 µM FCCP (Cayman Chemicals), or 20 µM stereoprobe for 8 hrs. Cells were harvested using TrypLE (ThermoFischer Scientific) and pelleted by centrifugation for 5 min at 350 x g at 4°C. Pelleted cells were resuspended in PBS and analyzed on a Attune NxT Flow Cytometer (ThermoFisher Scientific) running Attune Cytometric Software 5.1.2111.1. Mt-mKeima in a neutral pH environment was excited at 405 nm and emission was recorded with a 605/40 nm filter (channel VL-3). Mt-mKeima in an acidic environment was excited at 561 nm and emission was recorded with a 620/15 nm filter (channel YL-2). Cells were first gated based on forward scatter (FSC-H) and side scatter (SSC-A) to eliminate cellular debris. Per condition, at least 100,000 events were analyzed. For quantification of mitophagy-positive cells, YL-2A vs VL-3A fluorescence was plotted in FlowJo software v10.7.1 for each sample. A polygonal gate was then drawn above the x-y scatter of a sample only undergoing basal mitophagy (i.e. a DMSO treated sample). The percentage of mitophagy-positive cells was then analyzed for each sample.

## Data availability

The mass spectrometry proteomics data have been deposited to the ProteomeXchange Consortium via the PRIDE^125^ partner repository with the dataset identifier PXD069511. Previously published datasets^21^ relevant to this study are under data identifier PXD042541. Raw proteomic files were searched using the ProLuCID algorithm using a reverse concatenated, non-redundant variant of the Human UniProt database (release 201607). Processed proteomic data are provided in Supplementary Datasets 1 and 3.

## Code availability

Custom codes used for Boltz-2 data analysis are available at https://github.com/cravattlab/Hayward_Boltz_analysis.

## Supporting information

Supplementary Chemistry Information

## Acknowledgements

This work was supported by the National Cancer Institute (R35 CA231991), an HHMI Hanna H. Gray Fellowship (GT15176; E.N.), a Jane Coffin Childs Memorial Fellowship (K.E.D.), the Damon Runyon Cancer Research Foundation (DRG: 2406-20; H.L.), the NCI (K99/R00 CA290143; H.L), the NIH (R35-GM138206; M.A.H., T32-GM008326; M.A.M.), the Searle Scholars Program (M.A.H.), and the Cottrell Scholars Program (M.A.H.). We thank Xuedong Liu and Bing Chen (WuXi AppTec) for small-molecule synthesis, JC Ducom (High Performance Computing Core Scripps Research) for assistance and helpful discussions on Boltz-2, Kathryn Spencer and Scott Henderson (Core Microscopy Facility Scripps Research) for assistance with microscopy, Corey Hoang and Elizabeth Billings (The Center for Metabolomics and Mass Spectrometry Scripps Research) for obtaining intact MS data, Michael Won for assistance with intact MS experiments, Jolene Diedrich and Antonio Pinto for assistance with proteomics (Proteomics Core Scripps Research), Parth Jariwala and Enrique Saez for assistance with Seahorse assays, Chris Reinhardt and Wieland Goetzke for assistance with high-resolution mass spectrometry and review of synthetic chemistry data. Molecular graphics and analyses were performed with UCSF ChimeraX, developed by the Resource for Biocomputing, Visualization, and Informatics at the University of California, San Francisco (UCSF), with support from National Institutes of Health (NIH) grant R01-GM129325 and the Office of Cyber Infrastructure and Computational Biology, National Institute of Allergy and Infectious Diseases. The funders had no role in study design, data collection and analysis, decision to publish or preparation of the manuscript.

## Author contributions

R.E.H., B.M. and B.F.C. conceived the study. R.E.H., E.N. and K.E.D. performed proteomic experiments. R.E.H., B.M. and B.F.C., performed analysis of proteomic data. R.F.B., Z.G., B.M., and B.F.C. generated and analyzed Boltz-2 data. R.E.H., R.F.B., B.M., and B.F.C. wrote the paper. R.E.H. and V.C. confirmed protein–stereoprobe interactions by gel-ABPP. R.E.H. performed all functional assays, except for the mitophagy flux mt-mKeima assay, which was carried out by M.G. and F.U.H. Compound synthesis and characterization were supervised by B.M. and B.F.C. Additional resources for the study were contributed by M.A.M., H.L., and M.A.H. All authors edited and approved the paper.

---

**Extended Data Fig. 1.**
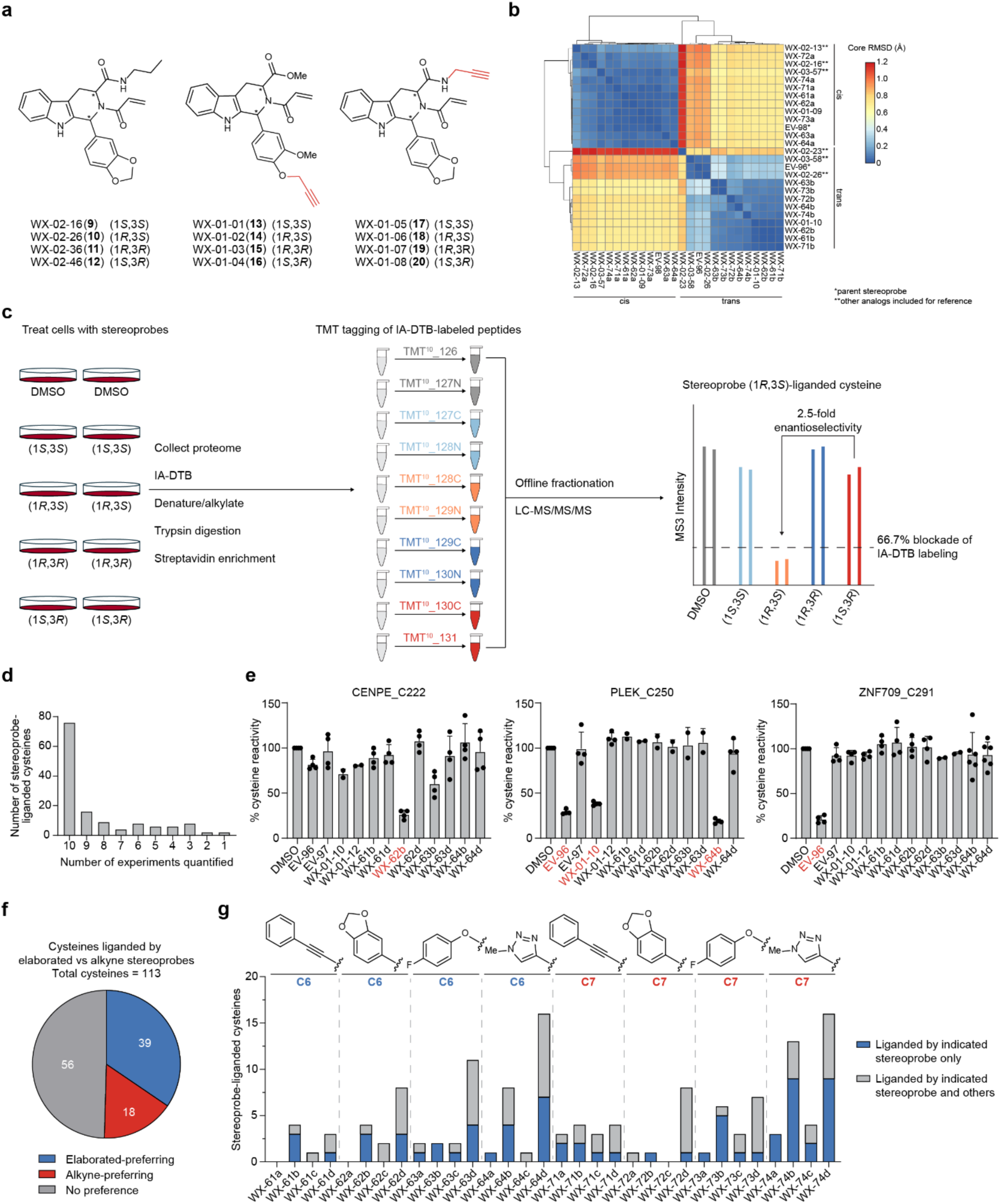
Chemical proteomic analysis of elaborated tryptoline acrylamide stereoprobes. **a**, Structures of parent and alkynylated tryptoline acrylamide stereoprobes used previously^21^. **b,** Heatmap showing DFT calculations for elaborated tryptoline acrylamide stereoprobes compared to unelaborated counterparts (* parent stereoprobes, ** other analogs included for analysis), suggesting minimal impact of C6-and C7-substitution on tryptoline acrylamide core geometry. **c,** Workflow for cysteine-directed ABPP experiments where stereoprobe reactivity with cysteines is determined by blockade of iodoacetamide-desthiobiotin (IA-DTB) labeling, streptavidin enrichment, and identification and quantification by multiplexed (tandem mass tagging, TMT^10p^^lex^) MS-based proteomics, as described previously^21^. **d,** Bar graph showing the number of experiments in which stereoprobe-liganded cysteines were quantified (shown for all cysteines liganded by elaborated, parent, and/or alkyne stereoprobes). **e,** Bar graphs showing representative liganding profiles for cysteines preferentially engaged by elaborated (left) or parent stereoprobes (right) or showing no preference (middle). Data represent average values ± SD of four independent experiments. **f,** Pie chart showing number of cysteines preferentially engaged (> 1.5-fold) by elaborated stereoprobes or alkyne stereoprobes. Cysteines not quantified in either elaborated or alkyne stereoprobe datasets were excluded from the analysis (12 total cysteines). **g,** Bar graph comparing the number of liganded cysteines for each elaborated tryptoline acrylamide stereoprobe where blue and grey designate cysteines that were engaged solely by the indicated stereoprobe vs multiple stereoprobes, respectively.

**Extended Data Fig. 2.**
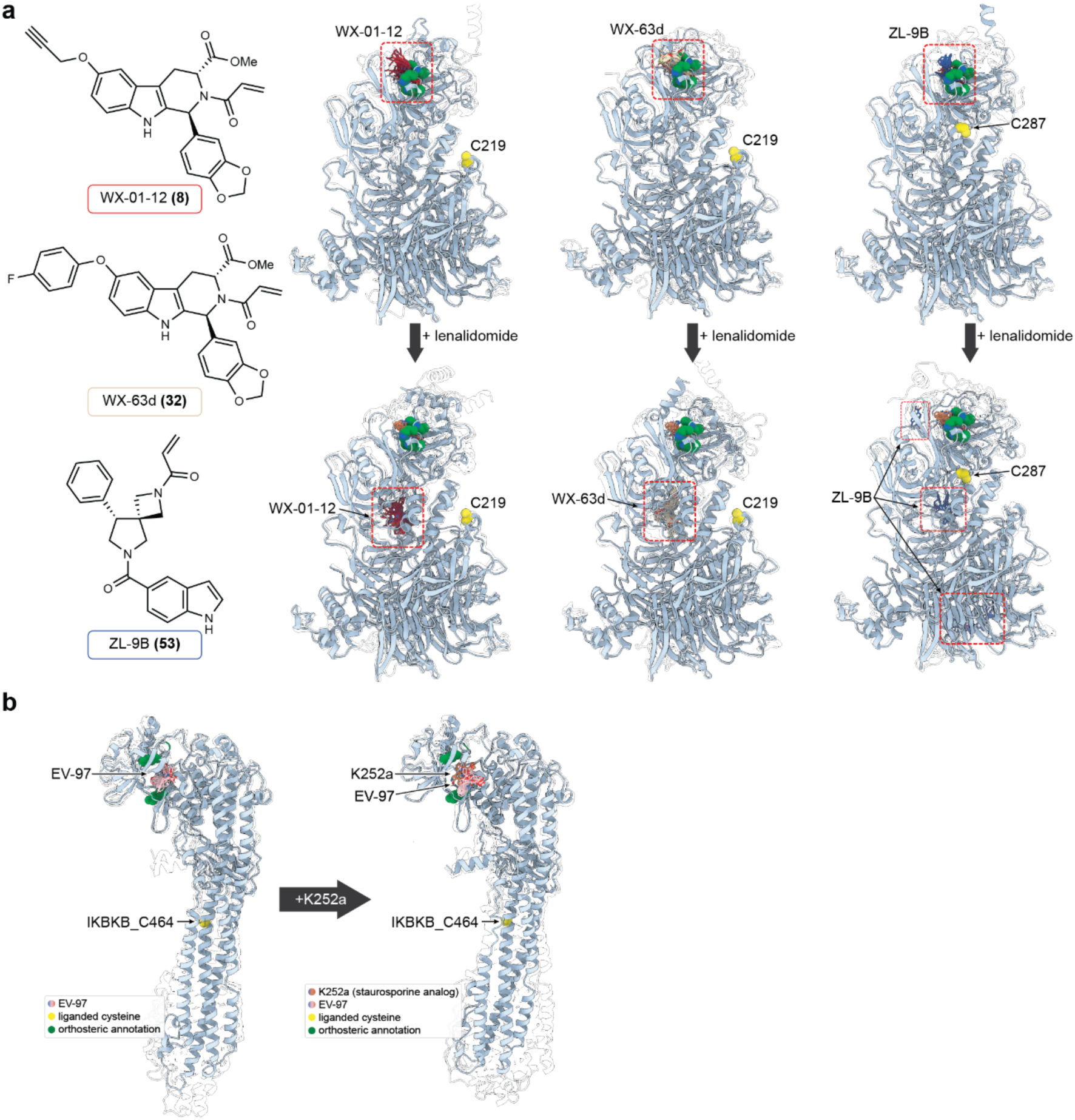
Boltz-2 analysis of tryptoline acrylamide-liganded proteins with virtual competition by orthosteric ligands. **a**, A virtual competition assay comparing Boltz-2 predictions for various allosteric stereoprobe ligands for CRBN with and without the orthosteric ligand lenalidomide. Allosteric ligands targeting CRBN_C219 (WX-01-12; WX-63d) or CRBN_C287 (ZL-9B) are shown (far left) alongside CRBN:DDB1^ΔBPB^ predicted structures with WX-01-12 (left), WX-63d (center), and ZL-9B (right); generated with allosteric ligands alone (top), or with allosteric ligands and lenalidomide (bottom). Boltz-2 rank 0 models are light blue, rank 1-9 models are transparent. Liganded cysteines are colored yellow. Ligands colored per their assignment at far left, lenalidomide poses are coral. **b**, A virtual competition assay comparing Boltz-2 predictions for IKBKB and EV-97 with (*right*) and without (*left*) K252a, a staurosporine analog known to bind the ATP-binding pocket as shown in PDB 4KIK. Boltz-2 rank 0 models are light blue, rank 1-9 models are transparent. EV-97 is pink, K252a is plum.

**Extended Data Fig. 3.**
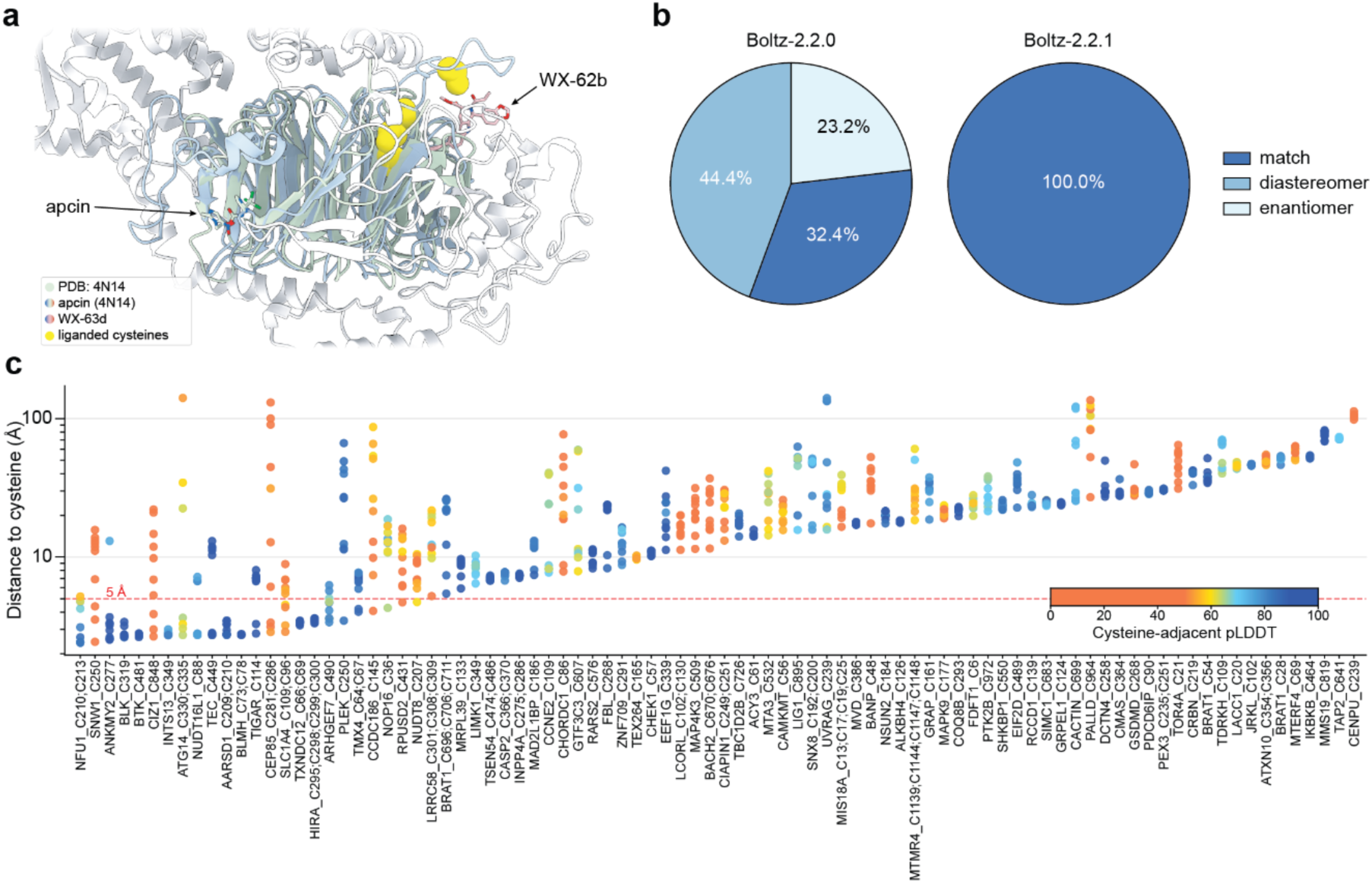
Additional Boltz-2 analysis of tryptoline acrylamide-liganded proteins. **a**, Rank 0 structure of HIRA co-folded with WX-62b aligned with the crystal structure of the WD40 domain protein Cdc20 in complex with the small molecule apcin (PDB 4N14). The WD40 domain of HIRA is light blue, while the remainder of HIRA is white. Candidate liganded cysteines are yellow spheres (note that the tryptic peptide for HIRA found to be engaged by WX-62b in cysteine-directed ABPP experiments has four cysteines (C295, C298-C300) representing candidates for the liganding event). WX-62b is pink. The 4N14 structure, including the ligand apcin, is green. **b**, Pie charts representing the distribution of stereochemistries for predicted stereoprobe ligands with respect to the stereochemistry of the experimentally determined stereoprobe ligands (determined by cysteine-directed ABPP) and specified in the input for all structural predictions for Boltz-2.2.0 (no template-aware stereochemical guidance) and Boltz-2.2.1 (adds template-aware stereochemical guidance). **c**, Waterfall plot showing cysteine-ligand distances as predicted by Boltz-2.2.1 for each protein identified as having a tryptoline acrylamide stereoprobe-liganded cysteine (**Supplementary Dataset 1**). The ten diffusion samples for the protein-stereoprobe pair with the closest stereoprobe-cysteine distance are shown. Proteins are sorted by closest stereoprobe distance and points are colored according to the mean pLDDT of all residues within 5 Å of the liganded cysteine.

**Extended Data Fig. 4.**
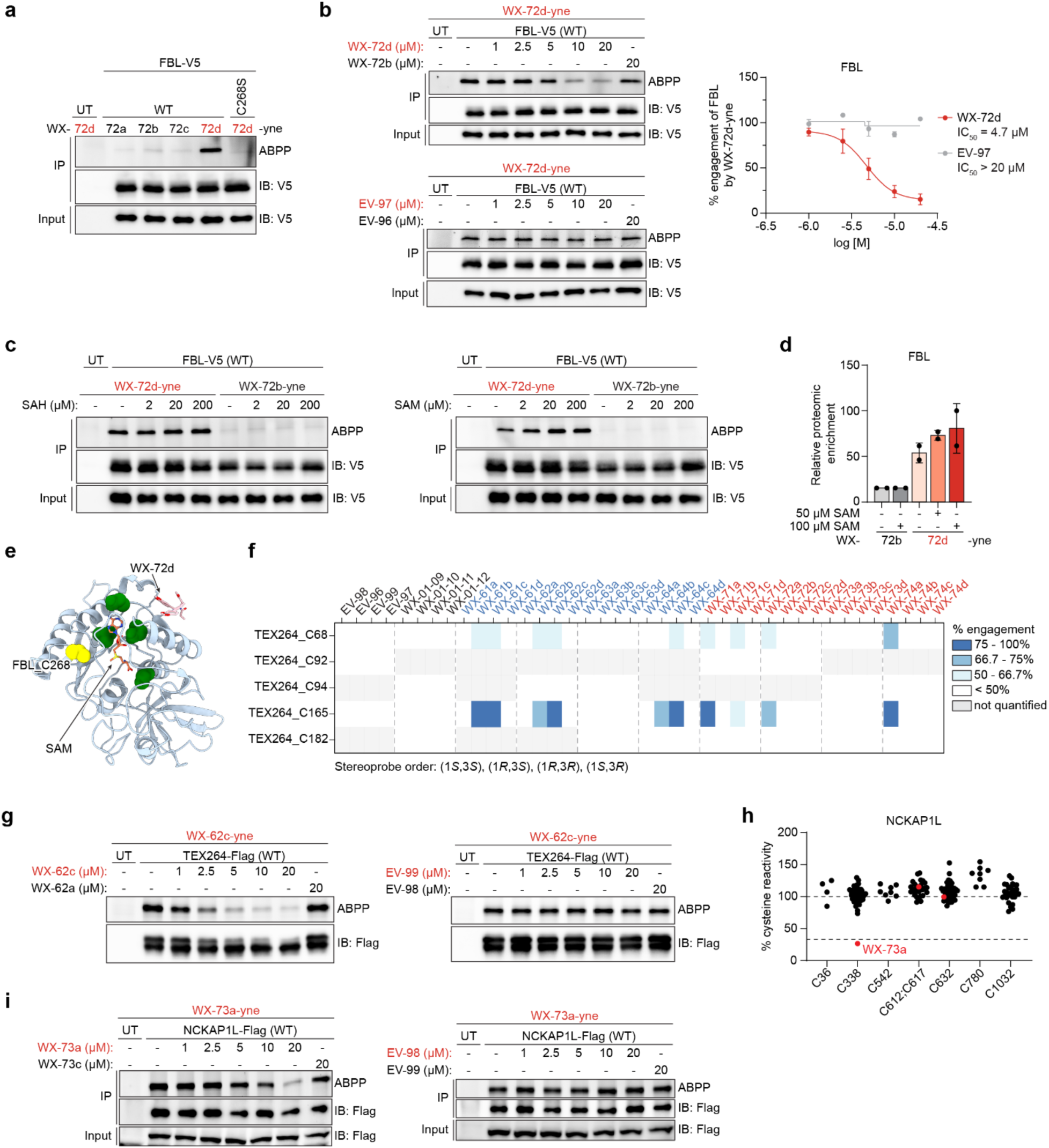
Characterization of elaborated stereoprobe-cysteine interactions. **a**, Gel-ABPP data showing stereoselective engagement of recombinant WT-FBL, but not a C268S-FBL mutant by WX-72d-yne (5 µM, 1 h) in lysates of HEK293T cells. UT, untransfected cells. **b,** Gel-ABPP data showing concentration-dependent and enantioselective blockade of WX-72d-yne (5 µM, 1 h) engagement of WT-FBL by WX-72d (top gel) but not parent probe EV-97 (bottom gel) (2 h pretreatment). Left, representative gel-ABPP data; right, quantification of gel-ABPP data. Data represent average values ± SD for 2 independent experiments. **c,** Gel-ABPP data showing increased enantioselective liganding of recombinant WT-FBL by WX-72d-yne in the presence of SAM (right) but not SAH (left). Experiments were performed in lysates of HEK293T cells stably expressing WT-FBL. **d,** Protein-directed ABPP data showing relative enrichment of endogenous FBL in Ramos cell lysates treated with WX-72d-yne or inactive enantiomer WX-72b-yne (5 µM, 1 h) in the presence and absence of SAM (50 or 100 µM). Data represent average values ± SD for two independent experiments. **e,** Boltz-2 prediction for WX-72d binding to FBL showing WX-72d placement in the presence of SAM, which occupies the SAM/SAH binding pocket, causing displacement of WX-72d to a site distal to liganded C268. **f,** Heatmap showing cysteine-directed ABPP data for TEX264, identifying decreases in the IA-DTB reactivity for C165 and C68 in cysteine-directed ABPP experiments of Ramos cells treated with the indicated stereoprobes. Blue and red text represent C6- and C7-elaborated stereoprobes. **g,** Gel-ABPP data showing concentration-dependent and enantioselective blockade of WX-62c-yne (5 µM, 1 h) engagement of recombinant TEX264 by WX-62c (left), but not parent stereoprobe EV-99 (right) (2 h pretreatment) in HEK293T cells. **h,** Cysteine-directed ABPP data for all quantified cysteines in NCKAP1L and all stereoprobes (elaborated, parent, and alkyne stereoprobes), identifying WX-73a (red) as the sole stereoprobe that liganded C338.The dashed line marks 66.7% IA-DTB blockade used to define a liganded cysteine event. Cysteines separated by a semi-colon (;) are located on the same tryptic peptide. **i,** Gel-ABPP data showing concentration-dependent and enantioselective blockade of WX-73a-yne (5 µM, 1 h) engagement of recombinant NCKAP1L by WX-73a (left) but not parent probe EV-98 (right) (2 h pretreatment) in HEK293T cells. For **f** and **h**, cysteine-directed ABPP data represent average values, and only cysteines quantified from at least two independent experiments are shown (from a total of 4-6 experiments performed per stereoprobe). For **a**, **g**, and **i**, data are from a single experiment representative of at least two independent experiments

**Extended Data Fig. 5.**
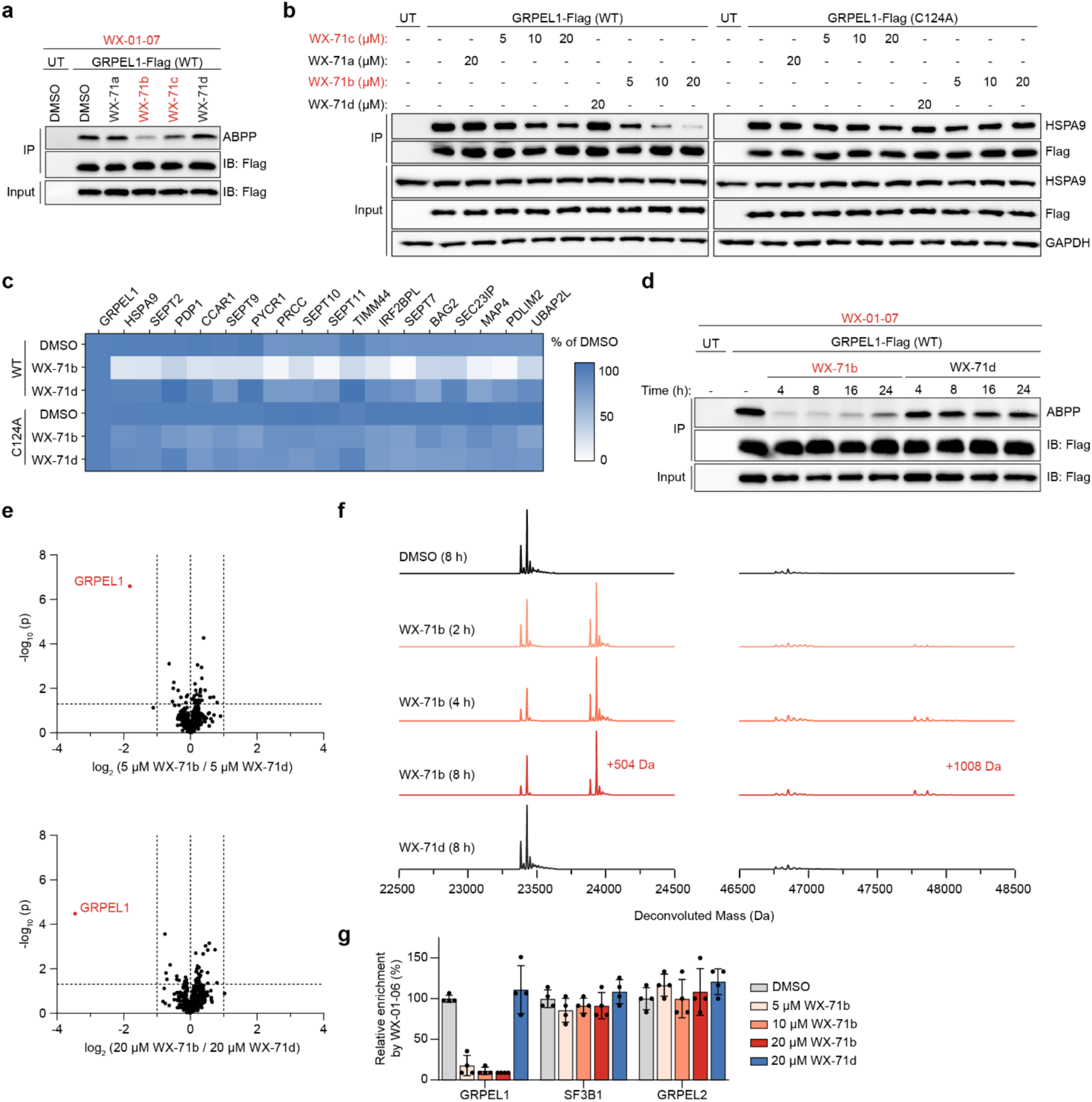
Tryptoline acrylamide stereoprobes disrupt GRPEL1:HSPA9 interactions. **a**, Gel-ABPP data showing enantioselective blockade of WX-01-07 (5 µM, 1 h) engagement of WT-GRPEL1 by WX-71b and WX-71c (20 µM, 1 h) in HCT-116 cells. UT, untransfected cells. **b,** IP-western blotting data showing concentration-dependent and enantioselective disruption of recombinant WT-GRPEL1 interactions with endogenous HSPA9 following treatments with WX-71b and WX-71c or enantiomeric controls WX-71d and WX-71a (indicated concentrations, 4 h) in HCT-116 cells. **c,** Heatmap of IP-MS data showing stereoselective disruption of protein interactions for recombinant WT-GRPEL1, but not C124A-GRPEL1, following treatment with WX-71b vs WX-71d (20 µM, 4 h) in HCT-116 cells. Displayed proteins exhibited at least 3-fold enrichment in WT-GRPEL1-expressing cells compared to untransfected HCT-116 cells. Protein signals were normalized to GRPEL1 signals within each treatment group and then to DMSO signals across treatment groups and are listed from left-to-right following GRPEL1 by average spectral count values. Data represent average values ± SD for four independent experiments. **d,** Gel-ABPP data showing time-dependent and enantioselective blockade of WX-01-07 (5 µM, 1 h) engagement of WT-GRPEL1 by WX-71b or WX-71d (20 µM, indicated times). **e,** Volcano plots of IP-MS data showing stereoselective disruption of protein interactions for recombinant HSPA9 following treatment with WX-71b vs WX-71d (5 or 20 µM, 4 h) in HCT-116 cells. Protein signals were normalized to HSPA9 within each treatment group and then to DMSO signals across treatment groups. Data represent average values ± SD of four independent experiments. Statistical significance was assessed using Welch two sample t-tests. **f,** Intact protein MS data for purified WT-GRPEL1 (3.5 µM) incubated with the indicated times with WX-71b or WX-71d (10 µM). Proteins were analyzed by time-of-flight (TOF)-LC/MS. **g,** Protein-directed ABPP data showing concentration-dependent and enantioselective blockade of WX-01-06 (5 µM, 1 h) engagement of GRPEL1, but not SF3B1 or GRPEL2, by WX-71b. Data represent average values ± SD of four independent experiments. For **a, b,** and **d**, data are from a single experiment representative of at least two independent experiments

**Extended Data Fig. 6.**
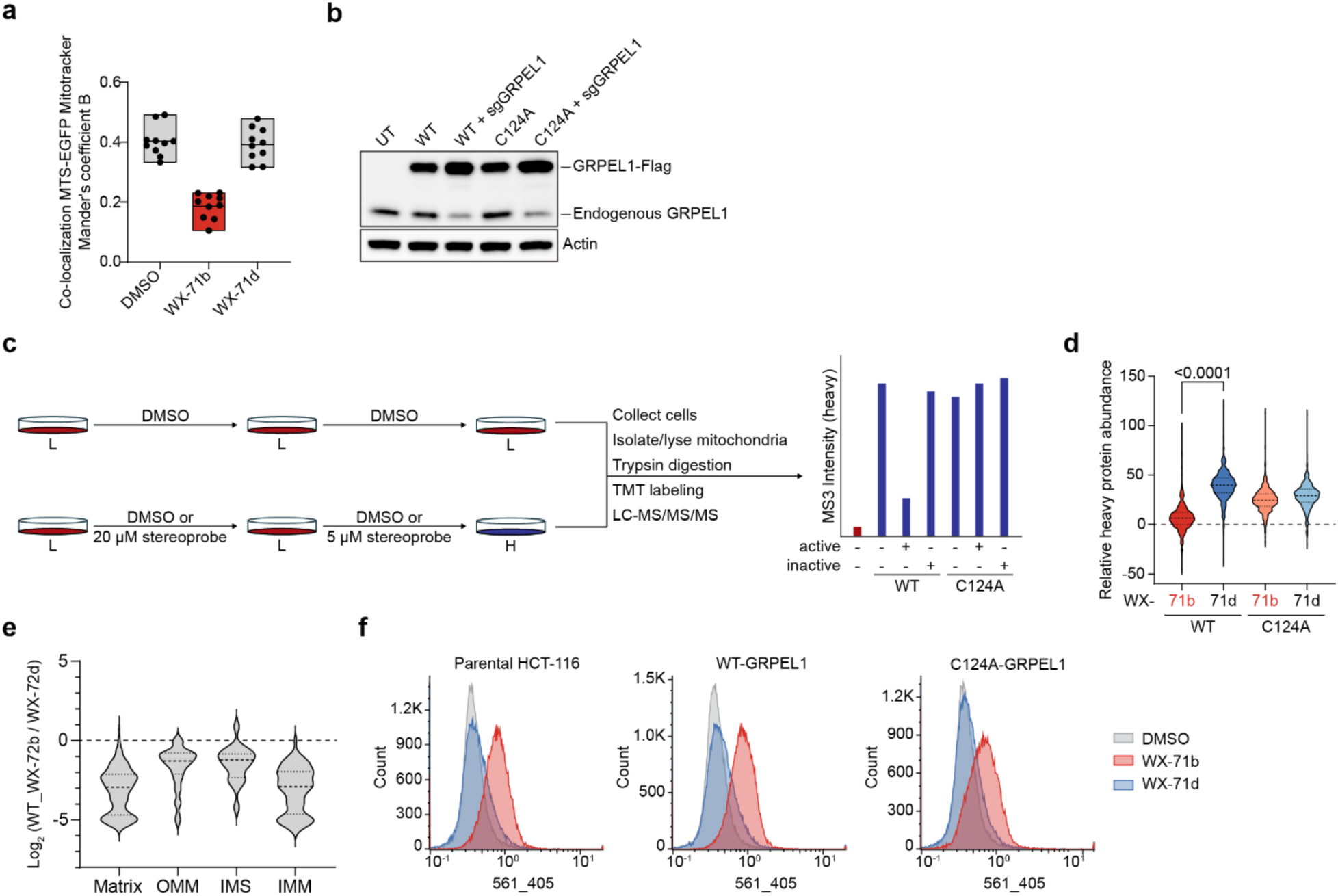
GRPEL1 stereoprobes modulate mitochondrial protein import and function. **a**, Colocalization of MTS-EGFP and Mitotracker Deep Red FM for 10 images from a single independent experiment performed in MTS-EGFP-inducible parental HCT-116 cells treated with doxycycline (0.5 µg/mL) and DMSO or WX-71b or WX-71d (20 µM, 8 h) as described in Fig. 5a. Data represent average values ± SD of ten technical replicates. **b,** Generation of sgGRPEL1 cells. HCT-116 cells stably expressed Flag epitope-tagged WT or C124A-GRPEL1 were subject to CRISPR/Cas9 disruption of endogenous GRPEL1 and analyzed at the population level. **c,** Workflow for pulsed-SILAC labeling with tandem-mass tag (TMT^16p^^lex^)-based multiplexing. sgGRPEL1 HCT-116 cells expressing Flag-epitope tagged WT-or C124A-GRPEL1 were pretreated in light media with DMSO or WX-71b or WX-71d (20 µM, 4 h) followed by shifting the cells to heavy amino acid media in the continued presence of stereoprobes (5 µM, 8 h). Mitochondria were biochemically enriched and analyzed by quantitative proteomics. **d,** Violin plot showing heavy-labeled protein abundance for the indicated treatment groups in pulse-SILAC experiments. Protein signals were corrected to light-labeled protein signals and normalized to heavy-labeled DMSO-treated WT-GRPEL1 or C124A-GRPEL1 signals. Statistical significance evaluated with parametric, two-tailed, paired t-test. **e,** Violin plot showing relative heavy-labeled protein abundance for WT-GRPEL1 cells across the indicated submitochondrial localizations as annotated from MitoCarta3.0 in combination with information retrieved from Uniprot and Human Protein Atlas. OMM, outer mitochondrial membrane; IMS, mitochondrial intermembrane space; IMM, inner mitochondrial membrane. **f,** Histograms (cell count (y axis)) versus mt-mKeima excitation (x axis)) showing mitophagy induction in (left) parental HCT-116 cells or sgGRPEL1 cells expressing WT-GRPEL1 (middle) or C124A-GRPEL1 (right) and also expressing mt-mKeima and Parkin treated with DMSO or WX-71b or WX-71d (20 µM, 8 h). Data show a single experiment representative of three independent experiments (Fig. 5h, i present quantification).

