## Supplementary Chemistry Information for "Tryptoline Stereoprobe Elaboration Identifies Inhibitors of the GRPEL1-HSPA9 Chaperone Complex"

### Supplementary Information: Synthesis and Characterization of Novel Compounds

#### General considerations

All NMR spectra were recorded at 298 K unless otherwise noted. $^1$H NMR spectra were recorded on Bruker Avance III 400, Avance III HD 400, Avance Neo 400 spectrometers or a ZKNJ QOne Quantum-I Plus 400 spectrometer ($^1$H, 400 MHz). $^1$H NMR data are reported as follows: chemical shift (δ), multiplicity (s = singlet, d = doublet, t = triplet, m = multiplet; br. = broad), coupling constants, and integration. Chemical shifts are reported in parts per million (ppm) using the appropriate solvent as reference.[1] Analytical supercritical fluid chromatography (SFC) was performed on a Shimadzu LC system (flow rate: 3 mL/min, back pressure: 100 Bar, column temperature: 35 ºC) equipped with a polydiode array detector. Tandem liquid chromatography/mass spectrometry (LC-MS) was performed on an Agilent 1200 series LC-MSD system equipped with an Agilent G6110A mass detector, alternatively a Shimadzu LC-20AD or AB series LC-MS system equipped with Shimadzu SPD-M20A or SPDM40 mass detectors. Mass measurements for high-resolution mass spectrometry (HRMS) were performed on a Waters Xevo G2-XS TOF calibrated against sodium formate clusters and using a LeuEnk lockmass. Expected monoisotopic masses were calculated using MassLynx 4.1 and the m/z values for calibrant and lockmass were MassLynx-default values.

#### Experimental procedures and analytical data

The compounds used in this study were synthesized by adapting previously reported protocols.[2-5] Representative new experimental procedures (synthesis of WX-61c, WX-62c, and WX-63c) as well as analytical data for all new stereoprobes are provided.

*Synthesis of WX-61c (WX-02-420)*

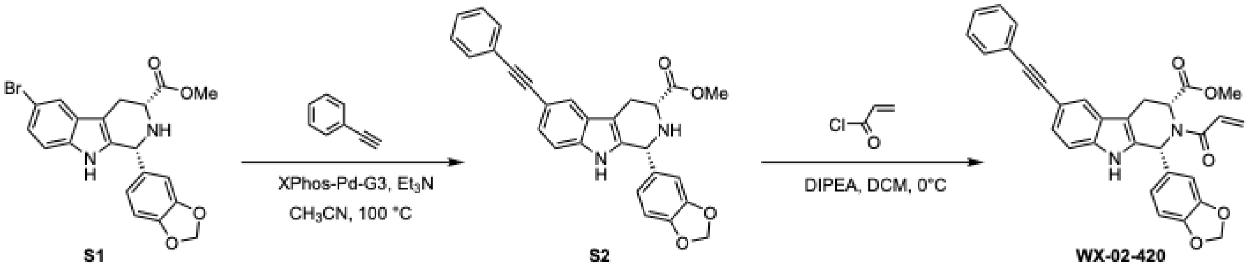


methyl (1*R*,3*R*)-1-(benzo[*d*][1,3]dioxol-5-yl)-6-(phenylethynyl)-2,3,4,9-tetrahydro-1*H*-pyrido[3,4-*b*]indole-3-carboxylate (S2)

A mixture of S1 (140 mg, 326 µmol, 1.0 equiv), phenylacetylene (167 mg, 1.63 mmol, 5.0 equiv), triethylamine (99.0 mg, 978 µmol, 3.0 equiv), XPhos-Pd-G3 (27.6 mg, 32.6 µmol, 10 mol%) in acetonitrile (2 mL) was degassed and purged with $N_2$ (× 3), and then the mixture was warmed to 100 °C and stirred for 16 h under $N_2$ atmosphere. Upon reaction completion, the mixture was partitioned between ethyl acetate 60 mL and brine (40 mL) and the aqueous layer was extracted with ethyl acetate (40 mL × 3). The organic layers were combined, dried over sodium sulfate, filtered, and concentrated under reduced pressure. The resulting residue was purified by prep-

TLC ($SiO_2$, petroleum ether/EtOAc = 2:1) and HPLC to give S2 (80 mg, 51% yield) as a yellow solid.
$^1$H NMR (400 MHz, $CD_3OD$) δ 8.27 (br. s, 1H), 7.68 (t, *J* = 1.2 Hz, 1H), 7.54 – 7.46 (m, 2H), 7.40 – 7.29 (m, 3H), 7.25 (d, *J* = 1.2 Hz, 2H), 6.94 (dd, *J* = 8.0, 1.7 Hz, 1H), 6.88 (d, *J* = 7.9 Hz, 1H), 6.84 (d, *J* = 1.7 Hz, 1H), 5.98 (t, *J* = 1.0 Hz, 2H), 5.36 (d, *J* = 2.7 Hz, 1H), 4.15 (dd, *J* = 11.4, 4.6 Hz, 1H), 3.85 (s, 3H), 3.30 – 3.24 (m, 1H), 3.04 (ddd, *J* = 15.4, 11.4, 2.5 Hz, 1H); 1 exchangeable proton not observed.
LC-MS m/z calc. for $C_{28}H_{23}N_2O_4$ $[M+H]^+$ 451.2 found 451.2.

methyl (1*R*,3*R*)-2-acryloyl-1-(benzo[*d*][1,3]dioxol-5-yl)-6-(phenylethynyl)-2,3,4,9-tetrahydro-1*H*-pyrido[3,4-*b*]indole-3-carboxylate (WX-61c (WX-02-420))
To a precooled (0 ºC) solution of S2 (60.0 mg, 133 µmol, 1.0 equiv) in DCM (2 mL) were added DIPEA (51.6 mg, 400 µmol, 3.0 equiv) and acryloyl chloride (36.2 mg, 400 µmol, 3.0 equiv). The mixture was stirred at 0 °C for 1 h. Upon reaction completion, the mixture was concentrated under reduced pressure. The resulting residue was purified by prep-TLC ($SiO_2$, petroleum ether/EtOAc = 2:1) and prep-HPLC (mobile phase: A: water (10 mM $NH_4HCO_3$), B: acetonitrile; gradient: 50%-83% B over 8 min) to give WX-61c (WX-02-420) (42.0 mg, 63% yield) as a yellow solid.
$^1$H NMR (400 MHz, DMSO-$d_6$) δ 11.17 (s, 1H), 7.81 (s, 1H), 7.58 – 7.52 (m, 2H), 7.47 – 7.24 (m, 5H), 7.06 – 6.66 (m, 4H), 6.52 – 6.43 (m, 1H), 6.27 (dd, *J* = 16.6, 2.2 Hz, 1H), 5.98 (d, *J* = 6.8 Hz, 2H), 5.82 (d, *J* = 10.5 Hz, 1H), 5.48 (d, *J* = 6.6 Hz, 1H), 3.50 (d, *J* = 16.2 Hz, 1H), 3.30 (s, 1H), 3.03 (s, 3H).
HRMS m/z calc. for $C_{31}H_{25}N_2O_5$ $[M+H]^+$ 505.1763 found 505.1758.

*Synthesis of WX-62c (WX-02-424)*

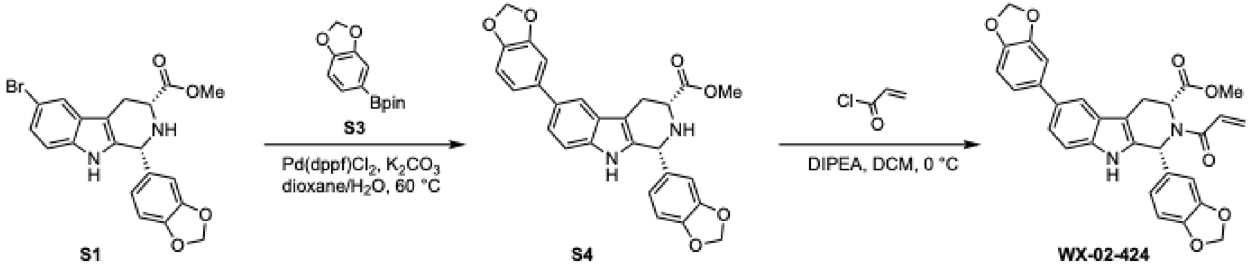


methyl (1*R*,3*R*)-1,6-bis(benzo[*d*][1,3]dioxol-5-yl)-2,3,4,9-tetrahydro-1*H*-pyrido[3,4-*b*]indole-3-carboxylate (S4)
A mixture of S1 (180 mg, 419 µmol, 1.0 equiv), pinacol ester S2 (156 mg, 629 µmol, 1.5 equiv), Pd(dppf)$Cl_2$ (30.7 mg, 41.9 µmol, 10 mol%), $K_2CO_3$ (116 mg, 839 µmol, 2.0 equiv) and water (2 mL) in dioxane (6 mL) was degassed and purged with $N_2$ (× 3). The mixture was then warmed to 60 ºC and stirred for 1 h under $N_2$ atmosphere. Upon reaction completion, the mixture was partitioned between ethyl acetate (60 mL) and brine (40 mL) and extracted with ethyl acetate (40 mL × 3). The organic layers were combined, dried over sodium sulfate, filtered, and concentrated under reduced pressure. The resulting residue was purified by column chromatography ($SiO_2$, petroleum ether/EtOAc = 10:1 to 1:1) to give S4 (130 mg, 90% w/w [pinacol], 50% yield) as a yellow solid.
$^1$H NMR (400 MHz, $CDCl_3$) δ 7.65 (d, *J* = 1.7 Hz, 1H), 7.45 (br. s, 1H), 7.32 (dd, *J* = 8.4, 1.8 Hz, 1H), 7.26 – 7.22 (m, 1H), 7.14 – 7.06 (m, 2H), 6.92 – 6.87 (m, 2H), 6.85 – 6.76 (m, 2H), 6.00 (s,

2H), 5.96 (s, 2H), 5.30 (s, 1H), 5.20 (br. s, 1H), 4.02 – 3.94 (m, 1H), 3.82 (s, 3H), 3.25 (dd, *J* = 15.7, 4.1 Hz, 1H), 3.02 (ddd, *J* = 14.5, 11.1, 2.6 Hz, 1H).
LC-MS m/z calc. for $C_{27}H_{23}N_2O_6$ $[M+H]^+$ 471.2 found 471.1.

methyl (1*R*,3*R*)-2-acryloyl-1,6-bis(benzo[*d*][1,3]dioxol-5-yl)-2,3,4,9-tetrahydro-1*H*-pyrido[3,4-*b*]indole-3-carboxylate (WX-62c (WX-02-424))
To a precooled (0 ºC) solution of S4 (40.0 mg, 90% w/w [pinacol], 76.5 µmol, 1.0 equiv) in DCM (1 mL) were added DIPEA (33.0 mg, 255 µmol, 3.3 equiv) and acryloyl chloride (11.5 mg, 127 µmol, 1.7 equiv). The mixture was stirred at 0 °C for 1 h. Upon reaction completion, the mixture was concentrated under reduced pressure. The resulting residue was purified by prep-HPLC (mobile phase: A: water (10 mM $NH_4HCO_3$), B: acetonitrile; gradient: 45%-75% B over 8min) to give WX-62c (WX-02-424) (26.0 mg, 62% yield) as an off-white solid.
$^1$H NMR (400 MHz, $CD_3OD$) δ 7.67 (t, *J* = 1.3 Hz, 1H), 7.37 – 7.28 (m, 2H), 7.17 – 7.09 (m, 2H), 7.03 – 6.98 (m, 1H), 6.97 – 6.85 (m, 2H), 6.82 (br. s, 1H), 6.74 – 6.66 (m, 1H), 6.65 – 6.56 (m, 1H), 6.35 – 6.22 (m, 1H), 5.97 (s, 2H), 5.91 (s, 2H), 5.87 – 5.79 (m, 1H), 5.41 – 5.27 (m, 1H), 3.73 – 3.57 (m, 1H), 3.24 – 2.97 (m, 4H); 1 exchangeable proton not observed.
HRMS m/z calc. for $C_{30}H_{25}N_2O_7$ $[M+H]^+$ 525.1662 found 525.1679.

*Synthesis of WX-63c (WX-02-427)*

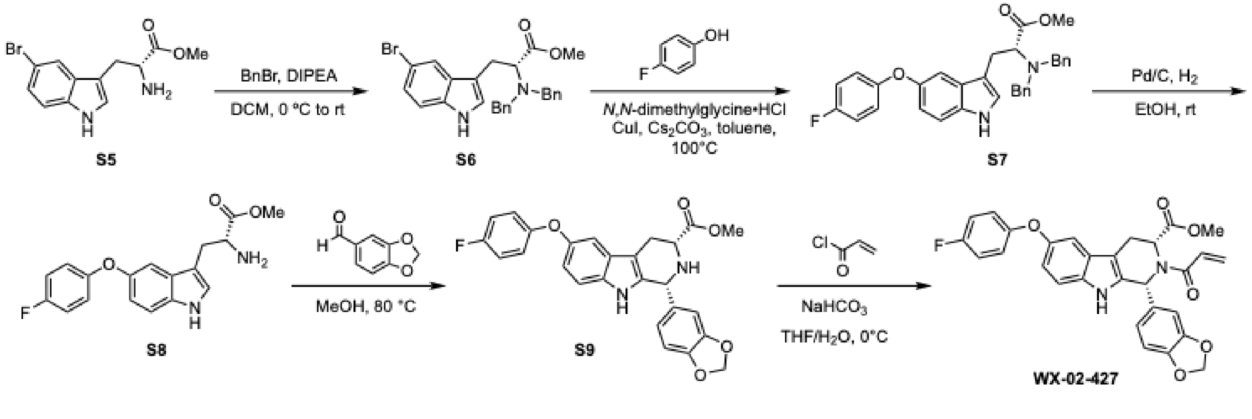


methyl (*R*)-3-(5-bromo-1*H*-indol-3-yl)-2-(dibenzylamino)propanoate (S6)
To a precooled (0 ºC) solution of S5 (3.5 g, 11.8 mmol, 1.0 equiv) in DCM (50 mL) were added DIPEA (5.33 g, 41.2 mmol, 3.5 equiv) and BnBr (12.1 g, 70.7 mmol, 6.0 equiv). The mixture was allowed to warm to rt and stirred for 16 h. Upon reaction completion, the mixture was concentrated under reduced pressure. The resulting residue was purified by column chromatography ($SiO_2$, petroleum ether/EtOAc = 100:1 to 5:1) to give S6 (4.7 g, 80% yield) as a yellow solid.
$^1$H NMR (400 MHz, $CD_3OD$) δ 7.33 – 7.16 (m, 12H), 7.12 (dd, *J* = 8.5, 1.9 Hz, 1H), 6.92 (s, 1H), 3.95 (d, *J* = 13.7 Hz, 2H), 3.73 – 3.64 (m, 4H), 3.49 (d, *J* = 13.7 Hz, 2H), 3.25 (dd, *J* = 14.3, 8.7 Hz, 1H), 3.03 (dd, *J* = 14.3, 6.4 Hz, 1H); 1 exchangeable proton not observed.
LC-MS m/z calc. for $C_{26}H_{26}BrN_2O_2$ $[M+H]^+$ 477.1 found 477.2.

methyl (*R*)-2-(dibenzylamino)-3-(5-(4-fluorophenoxy)-1*H*-indol-3-yl)propanoate (S7)
To a solution of S6 (4.6 g, 9.64 mmol, 1.0 equiv) and 4-fluorophenol (4.93 g, 44.0 mmol, 4.6 equiv) in toluene (50 mL) were added *N,N*-dimethylglycine•HCl (403 mg, 2.89 mmol, 30 mol%), $Cs_2CO_3$ (6.28 g, 19.3 mmol, 2.0 equiv) and CuI (184 mg, 964 µmol, 10 mol%). The mixture was warmed

to 100 °C and stirred for 12 hours under $N_2$ atmosphere. Upon reaction completion, the mixture was concentrated under reduced pressure. The resulting residue was purified by column chromatography ($SiO_2$, petroleum ether/EtOAc = 100:1 to 5:1) to give S7 (4.0 g, 93% w/w [DMF], 77% yield) as a yellow solid.
$^1$H NMR (400 MHz, $CD_3OD$) δ 7.57 – 7.33 (m, 11H), 7.19 (s, 1H), 7.08 – 6.94 (m, 3H), 6.94 – 6.80 (m, 3H), 4.84 – 4.51 (m, 4H), 4.27 (dd, *J* = 9.1, 5.9 Hz, 1H), 3.69 – 3.50 (m, 5H); 1 exchangeable proton not observed.
LC-MS m/z calc. for $C_{32}H_{30}FN_2O_3$ $[M+H]^+$ 509.2 found 509.3.

methyl (*R*)-2-amino-3-(5-(4-fluorophenoxy)-1*H*-indol-3-yl)propanoate (S8)
To Pd/C (419 mg, 393 µmol, 10% w/w, 21 mol%) under $N_2$ atmosphere were added sequentially EtOH (10 mL) and S8 (930 mg, 1.84 mmol, 1.0 equiv). The reaction mixture was degassed and purged with $N_2$ (× 3), then degassed and purged with $H_2$ (× 3). The reaction mixture was stirred at rt for 12 h under $H_2$ atmosphere (15 psi). Upon reaction completion, the mixture was filtered and concentrated under reduced pressure. The resulting residue was purified by prep-HPLC (A: $H_2O$ (10 mM HCl (36%)), B: acetonitrile; gradient: 0%-100% B over 25.0 min). To the product-containing fraction was added sat. aq. $NaHCO_3$ to reach pH > 7, following which the mixture was extracted with EtOAc (30 mL × 3). The combined organic layers were dried over sodium sulfate, filtered and concentrated under reduced pressure to give S8 (260 mg, 43% yield) as a yellow solid.
$^1$H NMR (400 MHz, $CD_3OD$) δ 7.35 (d, *J* = 8.7 Hz, 1H), 7.14 (d, *J* = 2.3 Hz, 2H), 7.07 – 6.98 (m, 2H), 6.98 – 6.87 (m, 2H), 6.83 (dd, *J* = 8.7, 2.2 Hz, 1H), 4.60 (br. s, 1H), 3.72 (t, *J* = 5.9 Hz, 1H), 3.63 – 3.56 (m, 3H), 3.37 – 3.34 (m, 2H), 3.13 – 3.07 (m, 2H).
LC-MS m/z calc. for $C_{18}H_{18}FN_2O_3$ $[M+H]^+$ 329.1 found 329.2.

methyl (1*R*,3*R*)-1-(benzo[*d*][1,3]dioxol-5-yl)-6-(4-fluorophenoxy)-2,3,4,9-tetrahydro-1*H*-pyrido[3,4-*b*]indole-3-carboxylate (S9)
To S8 (260 mg, 792 µmol, 1.0 equiv) was added HCl (2 M in dioxane, 3 mL, 7.6 equiv) and the mixture was concentrated under reduced pressure. The resulting residue was taken up in MeOH (20 mL), and piperonal (131 mg, 871 µmol, 1.1 equiv) was added. The mixture was warmed to 80 °C and stirred for 16 h. Upon completion, the reaction mixture was diluted with sat. aq. $NaHCO_3$ (20 mL) to maintain pH > 7, and extracted with EtOAc (100 mL × 3). The combined organic layers were dried over sodium sulfate, filtered and concentrated under reduced pressure. The resulting residue was purified by column chromatography ($SiO_2$, petroleum ether/EtOAc = 10:1 to 2:1) to give S9 (120 mg, 32% yield) and (1*S*,3*R*)-*epi*-S9 (120 mg, 32% yield) as yellow solids.

S9:
$^1$H NMR (400 MHz, $CD_3OD$) δ 7.22 (d, *J* = 8.7 Hz, 1H), 7.07 (d, *J* = 2.3 Hz, 1H), 7.03 – 6.97 (m, 2H), 6.93 – 6.86 (m, 3H), 6.86 – 6.80 (m, 2H), 6.77 (dd, *J* = 8.7, 2.3 Hz, 1H), 5.95 (q, *J* = 1.2 Hz, 2H), 5.17 (s, 1H), 4.60 (br.s, 0.5H), 3.91 (dd, *J* = 11.2, 4.3 Hz, 1H), 3.78 (s, 3H), 3.35 (s, 1.5H), 3.09 (ddd, *J* = 15.1, 4.4, 1.8 Hz, 1H), 2.88 (ddd, *J* = 15.2, 11.2, 2.4 Hz, 1H).
LC-MS m/z calc. for $C_{26}H_{22}FN_2O_5$ $[M+H]^+$ 461.2 found 461.1.

*epi*-S9:
$^1$H NMR (400 MHz, $CD_3OD$) δ 7.24 (d, *J* = 8.7 Hz, 1H), 7.09 (d, *J* = 2.3 Hz, 1H), 7.07 – 6.97 (m, 2H), 6.94 – 6.88 (m, 2H), 6.84 – 6.76 (m, 2H), 6.75 – 6.67 (m, 2H), 5.92 (q, *J* = 1.2 Hz, 2H), 5.34

(s, 1H), 3.90 (dd, *J* = 7.8, 5.2 Hz, 1H), 3.71 (s, 3H), 3.13 (ddd, *J* = 15.4, 5.2, 1.1 Hz, 1H), 2.97 (ddd, *J* = 15.4, 7.8, 1.4 Hz, 1H); 2 exchangeable protons not observed.
LC-MS m/z calc. for $C_{26}H_{22}FN_2O_5$ $[M+H]^+$ 461.2 found 461.2.

methyl (1*R*,3*R*)-2-acryloyl-1-(benzo[*d*][1,3]dioxol-5-yl)-6-(4-fluorophenoxy)-2,3,4,9-tetrahydro-1*H*-pyrido[3,4-*b*]indole-3-carboxylate (WX-63c (WX-02-427))
To a precooled (0 ºC) solution of S9 (110 mg, 239 μmol, 1.0 equiv) in THF (3 mL) and $H_2O$ (1 mL) were added $NaHCO_3$ (100 mg, 1.19 mmol, 5.0 equiv) and acryloyl chloride (32.4 mg, 358 μmol, 1.5 equiv). The mixture was stirred at 0 °C for 0.5 hour. Upon reaction completion, the mixture was concentrated under reduced pressure. The residue was purified by prep-HPLC (mobile phase: A: $H_2O$ (10-mM $NH_4HCO_3$), B: acetonitrile; gradient: 42%-72% B over 15.0 min) to give WX-63c (WX-02-427) (53 mg, 43% yield) as a white solid.
$^1H$ NMR (400 MHz, $CDCl_3$) δ 8.26 (br. s, 1H), 7.24 (d, *J* = 8.7 Hz, 1H), 7.18 (d, *J* = 2.3 Hz, 1H), 7.05 – 6.87 (m, 7H), 6.78 – 6.55 (m, 3H), 6.30 (br. d, *J* = 17.1 Hz, 1H), 5.94 – 5.82 (m, 2H), 5.78 (dd, *J* = 10.8, 1.7 Hz, 1H), 5.02 (br. s, 1H), 3.56 (br. d, *J* = 15.8 Hz, 1H), 3.19 (br. s, 3H), 2.98 (ddd, *J* = 15.8, 7.0, 1.7 Hz, 1H).
HRMS m/z calc. for $C_{29}H_{24}FN_2O_6$ $[M+H]^+$ 515.1618 found 515.1638.

*Additional analytical data*

methyl (1*S*,3*S*)-2-acryloyl-1-(benzo[*d*][1,3]dioxol-5-yl)-6-(phenylethynyl)-2,3,4,9-tetrahydro-1*H*-pyrido[3,4-*b*]indole-3-carboxylate (WX-61a (WX-02-418))
$^1H$ NMR (400 MHz, DMSO-$d_6$) δ 11.17 (s, 1H), 7.81 (s, 1H), 7.63 – 7.48 (m, 2H), 7.46 – 7.23 (m, 5H), 7.07 – 6.66 (m, 4H), 6.51 – 6.40 (m, 1H), 6.27 (dd, *J* = 16.6, 2.2 Hz, 1H), 5.98 (d, *J* = 6.8 Hz, 2H), 5.82 (d, *J* = 10.5 Hz, 1H), 5.48 (d, *J* = 6.6 Hz, 1H), 3.50 (d, *J* = 16.2 Hz, 1H), 3.30 (s, 1H), 3.03 (s, 3H).
HRMS m/z calc. for $C_{31}H_{25}N_2O_5$ $[M+H]^+$ 505.1763 found 505.1777.

methyl (1*R*,3*S*)-2-acryloyl-1-(benzo[*d*][1,3]dioxol-5-yl)-6-(phenylethynyl)-2,3,4,9-tetrahydro-1*H*-pyrido[3,4-*b*]indole-3-carboxylate (WX-61b (WX-02-419))
$^1H$ NMR (400 MHz, DMSO-$d_6$) δ 11.32 – 11.10 (m, 1H), 7.73 (s, 1H), 7.62 – 7.48 (m, 2H), 7.46 – 7.17 (m, 5H), 7.17 – 6.67 (m, 4H), 6.28 (br. s, 0.6H), 6.20 – 5.87 (m, 3.4H), 5.68 (d, *J* = 10.5 Hz, 1H), 5.62 (br. s, 0.4H), 5.06 (br. s, 0.6H), 3.62 – 3.45 (m, 3.4H), 3.43 – 3.34 (m, 0.6H), 3.30 – 3.08 (m, 1H); ~6:4 mixture of rotamers.
HRMS m/z calc. for $C_{31}H_{25}N_2O_5$ $[M+H]^+$ 505.1763 found 505.1778.

methyl (1*S*,3*R*)-2-acryloyl-1-(benzo[*d*][1,3]dioxol-5-yl)-6-(phenylethynyl)-2,3,4,9-tetrahydro-1*H*-pyrido[3,4-*b*]indole-3-carboxylate (WX-61d (WX-02-421))
$^1H$ NMR (400 MHz, DMSO-$d_6$) δ 11.35 – 11.09 (m, 1H), 7.74 (s, 1H), 7.59 – 7.49 (m, 2H), 7.48 – 7.16 (m, 5H), 7.15 – 6.68 (m, 4H), 6.28 (br. s, 0.6H), 6.21 – 5.85 (m, 3.4H), 5.68 (dd, *J* = 10.4, 2.2 Hz, 1H), 5.62 (br. s, 0.4H), 5.06 (br. s, 0.6H), 3.62 – 3.46 (m, 3.4H), 3.42 – 3.33 (m, 0.6H), 3.30 – 3.15 (m, 1H); ~6:4 mixture of rotamers.
HRMS m/z calc. for $C_{31}H_{25}N_2O_5$ $[M+H]^+$ 505.1763 found 505.1756.

methyl (1*S*,3*S*)-2-acryloyl-1,6-bis(benzo[*d*][1,3]dioxol-5-yl)-2,3,4,9-tetrahydro-1*H*-pyrido[3,4-*b*]indole-3-carboxylate (WX-62a (WX-02-422))
$^1$H NMR (400 MHz, $CD_3OD$) δ 7.67 (t, *J* = 1.3 Hz, 1H), 7.36 – 7.30 (m, 2H), 7.18 – 7.10 (m, 2H), 7.04 – 6.97 (m, 1H), 6.96 – 6.85 (m, 2H), 6.82 (br. s, 1H), 6.75 – 6.66 (m, 1H), 6.64 – 6.56 (m, 1H), 6.36 – 6.22 (m, 1H), 5.97 (s, 2H), 5.91 (s, 2H), 5.88 – 5.78 (m, 1H), 5.40 – 5.28 (m, 1H), 3.71 – 3.59 (m, 1H), 3.24 – 3.02 (m, 4H); 1 exchangeable proton not observed.
HRMS m/z calc. for $C_{30}H_{25}N_2O_7$ $[M+H]^+$ 525.1662 found 525.1676.

methyl (1*R*,3*S*)-2-acryloyl-1,6-bis(benzo[*d*][1,3]dioxol-5-yl)-2,3,4,9-tetrahydro-1*H*-pyrido[3,4-*b*]indole-3-carboxylate (WX-62b (WX-02-423))
$^1$H NMR (400 MHz, $CD_3OD$) δ 7.59 (t, *J* = 1.3 Hz, 1H), 7.29 (br. s, 2H), 7.13 – 7.05 (m, 2H), 7.01 – 6.64 (m, 5H), 6.32 – 6.05 (m, 2H), 6.03 – 5.80 (m, 4H), 5.71 (dd, *J* = 10.6, 1.9 Hz, 1H), 5.55 – 5.41 (m, 0.3H), 5.10 – 4.98 (m, 0.7H), 3.73 – 3.41 (m, 4.3H), 3.29 – 3.18 (m, 0.7H); 1 exchangeable proton not observed; ~7:3 mixture of rotamers.
HRMS m/z calc. for $C_{30}H_{25}N_2O_7$ $[M+H]^+$ 525.1662 found 525.1672.

methyl (1*S*,3*R*)-2-acryloyl-1,6-bis(benzo[*d*][1,3]dioxol-5-yl)-2,3,4,9-tetrahydro-1*H*-pyrido[3,4-*b*]indole-3-carboxylate (WX-62d (WX-02-425))
$^1$H NMR (400 MHz, $CD_3OD$) δ 7.58 (t, *J* = 1.3 Hz, 1H), 7.36 – 7.21 (m, 2H), 7.12 – 7.04 (m, 2H), 6.98 – 6.66 (m, 5H), 6.30 – 6.04 (m, 2H), 5.99 – 5.78 (m, 4H), 5.70 (dd, *J* = 10.6, 1.9 Hz, 1H), 5.55 – 5.39 (m, 0.3H), 5.09 – 4.97 (m, 0.7H), 3.76 – 3.38 (m, 4.3H), 3.28 – 3.18 (m, 0.7H) ; 1 exchangeable proton not observed; ~7:3 mixture of rotamers.
HRMS m/z calc. for $C_{30}H_{25}N_2O_7$ $[M+H]^+$ 525.1662 found 525.1654.

methyl (1*S*,3*S*)-2-acryloyl-1-(benzo[*d*][1,3]dioxol-5-yl)-6-(4-fluorophenoxy)-2,3,4,9-tetrahydro-1*H*-pyrido[3,4-*b*]indole-3-carboxylate (WX-63a (WX-02-426))
$^1$H NMR (400 MHz, $CDCl_3$) δ 8.36 (br. s, 1H), 7.24 (d, *J* = 8.7 Hz, 1H), 7.18 (d, *J* = 2.4 Hz, 1H), 7.06 – 6.83 (m, 7H), 6.75 – 6.55 (m, 3H), 6.30 (br. d, *J* = 17.0 Hz, 1H), 5.86 (d, *J* = 8.9 Hz, 2H), 5.77 (dd, *J* = 10.8, 1.7 Hz, 1H), 5.02 (br. s, 1H), 3.56 (d, *J* = 15.8 Hz, 1H), 3.19 (s, 3H), 2.99 (ddd, *J* = 15.9, 7.0, 1.7 Hz, 1H).
HRMS m/z calc. for $C_{29}H_{24}FN_2O_6$ $[M+H]^+$ 515.1618 found 515.1634.

methyl (1*R*,3*S*)-2-acryloyl-1-(benzo[*d*][1,3]dioxol-5-yl)-6-(4-fluorophenoxy)-2,3,4,9-tetrahydro-1*H*-pyrido[3,4-*b*]indole-3-carboxylate (WX-63b (WX-02-428))
$^1$H NMR (400 MHz, $CD_3OD$) δ 7.33 – 7.15 (m, 1H), 7.10 – 7.04 (m, 1H), 7.04 – 6.95 (m, 2H), 6.95 – 6.84 (m, 4H), 6.83 – 6.66 (m, 3H), 6.31 – 6.02 (m, 2H), 5.92 (br. s, 2H), 5.74 – 5.66 (m, 1H), 5.53 – 5.38 (m, 0.3H), 5.08 – 4.95 (m, 0.7H), 3.61 (br. s, 3.3H), 3.32 (s, 1H), 3.21 – 3.08 (m, 0.7H); 1 exchangeable proton not observed; ~7:3 mixture of rotamers.
HRMS m/z calc. for $C_{29}H_{24}FN_2O_6$ $[M+H]^+$ 515.1618 found 515.1614.

methyl (1*S*,3*R*)-2-acryloyl-1-(benzo[*d*][1,3]dioxol-5-yl)-6-(4-fluorophenoxy)-2,3,4,9-tetrahydro-1*H*-pyrido[3,4-*b*]indole-3-carboxylate (WX-63d (WX-02-429))
$^1$H NMR (400 MHz, $CD_3OD$) δ 7.33 – 7.16 (m, 1H), 7.09 – 7.04 (m, 1H), 7.04 – 6.96 (m, 2H), 6.95 – 6.85 (m, 4H), 6.85 – 6.67 (m, 3H), 6.31 – 6.03 (m, 2H), 5.97 – 5.78 (m, 2H), 5.70 (dd, *J* = 10.6, 2.0 Hz, 1H), 5.51 – 5.37 (m, 0.3H), 5.07 – 4.97 (m, 0.7H), 3.61 (br. s, 3.3H), 3.32 (s, 1H), 3.21 – 3.07 (m, 0.7H); 1 exchangeable proton not observed; ~7:3 mixture of rotamers.
HRMS m/z calc. for $C_{29}H_{24}FN_2O_6$ $[M+H]^+$ 515.1618 found 515.1632.

methyl (1*S*,3*S*)-2-acryloyl-1-(benzo[*d*][1,3]dioxol-5-yl)-7-(phenylethynyl)-2,3,4,9-tetrahydro-1*H*-pyrido[3,4-*b*]indole-3-carboxylate (WX-71a (WX-02-430))
$^1$H NMR (400 MHz, $CD_3OD$) δ 7.56 – 7.45 (m, 4H), 7.40 – 7.28 (m, 3H), 7.23 (dd, *J* = 8.1, 1.4 Hz, 1H), 7.01 (br. s, 1H), 6.97 – 6.77 (m, 2H), 6.70 (d, *J* = 8.0 Hz, 1H), 6.65 – 6.44 (m, 1H), 6.28 (d, *J* = 16.9 Hz, 1H), 5.91 (s, 2H), 5.83 (d, *J* = 11.1 Hz, 1H), 5.41 – 5.29 (m, 1H), 3.69 – 3.53 (m, 1H), 3.15 (br. s, 3H), 3.09 – 2.98 (m, 1H); 1 exchangeable proton not observed.
HRMS m/z calc. for $C_{31}H_{25}N_2O_5$ $[M+H]^+$ 505.1763 found 505.1765.

methyl (1*R*,3*R*)-2-acryloyl-1-(benzo[*d*][1,3]dioxol-5-yl)-7-(phenylethynyl)-2,3,4,9-tetrahydro-1*H*-pyrido[3,4-*b*]indole-3-carboxylate (WX-71c (WX-02-431))
$^1$H NMR (400 MHz, $CD_3OD$) δ 7.56 – 7.45 (m, 4H), 7.41 – 7.29 (m, 3H), 7.23 (dd, *J* = 8.1, 1.4 Hz, 1H), 7.01 (br. s, 1H), 6.96 – 6.76 (m, 2H), 6.70 (d, *J* = 8.0 Hz, 1H), 6.64 – 6.42 (m, 1H), 6.28 (d, *J* = 16.9 Hz, 1H), 5.91 (s, 2H), 5.83 (d, *J* = 10.8 Hz, 1H), 5.42 – 5.23 (m, 1H), 3.69 – 3.53 (m, 1H), 3.15 (br. s, 3H), 3.09 – 2.95 (m, 1H); 1 exchangeable proton not observed.
HRMS m/z calc. for $C_{31}H_{25}N_2O_5$ $[M+H]^+$ 505.1763 found 505.1775.

methyl (1*R*,3*S*)-2-acryloyl-1-(benzo[*d*][1,3]dioxol-5-yl)-7-(phenylethynyl)-2,3,4,9-tetrahydro-1*H*-pyrido[3,4-*b*]indole-3-carboxylate (WX-71b (WX-02-432))
$^1$H NMR (400 MHz, $CD_3OD$) δ 7.57 – 7.40 (m, 4H), 7.38 – 7.27 (m, 3H), 7.17 (d, *J* = 8.2 Hz, 1H), 6.98 – 6.86 (m, 2H), 6.85 – 6.68 (m, 2H), 6.35 – 6.05 (m, 2H), 5.92 (br. s, 2H), 5.71 (dd, *J* = 10.7, 1.8 Hz, 1H), 5.54 – 5.41 (m, 0.3H), 5.13 – 5.00 (m, 0.7H), 3.61 (br. s, 3.3H), 3.53 – 3.38 (m, 1H), 3.31 – 3.12 (m, 0.7H); 1 exchangeable proton not observed; ~7:3 mixture of rotamers.
HRMS m/z calc. for $C_{31}H_{25}N_2O_5$ $[M+H]^+$ 505.1763 found 505.1773.

methyl (1*S*,3*R*)-2-acryloyl-1-(benzo[*d*][1,3]dioxol-5-yl)-7-(phenylethynyl)-2,3,4,9-tetrahydro-1*H*-pyrido[3,4-*b*]indole-3-carboxylate (WX-71d (WX-02-433))
$^1$H NMR (400 MHz, $CD_3OD$) δ 7.52 – 7.40 (m, 4H), 7.39 – 7.27 (m, 3H), 7.17 (dd, *J* = 8.1, 1.3 Hz, 1H), 7.00 – 6.87 (m, 2H), 6.84 – 6.66 (m, 2H), 6.33 – 6.03 (m, 2H), 5.92 (br. s, 2H), 5.70 (dd, *J* = 10.6, 1.8 Hz, 1H), 5.55 – 5.38 (m, 0.3H), 5.12 – 4.98 (m, 0.7H), 3.61 (br. s, 3.3H), 3.51 – 3.39 (m, 1H), 3.28 – 3.14 (m, 0.7H); 1 exchangeable proton not observed; ~7:3 mixture of rotamers.
HRMS m/z calc. for $C_{31}H_{25}N_2O_5$ $[M+H]^+$ 505.1763 found 505.1766.

methyl (1*S*,3*S*)-2-acryloyl-1,7-bis(benzo[*d*][1,3]dioxol-5-yl)-2,3,4,9-tetrahydro-1*H*-pyrido[3,4-*b*]indole-3-carboxylate (WX-72a (WX-02-434))
$^{1}$H NMR (400 MHz, $CD_3OD$) δ 7.55 (d, *J* = 8.2 Hz, 1H), 7.44 (s, 1H), 7.27 (dd, *J* = 8.2, 1.6 Hz, 1H), 7.16 – 7.06 (m, 2H), 7.00 (br. s, 1H), 6.96 – 6.78 (m, 3H), 6.75 – 6.65 (m, 1H), 6.64 – 6.55 (m, 1H), 6.27 (d, *J* = 16.9 Hz, 1H), 5.96 (s, 2H), 5.90 (s, 2H), 5.82 (d, *J* = 10.8 Hz, 1H), 5.39 – 5.25 (m, 1H), 3.62 (d, *J* = 16.1 Hz, 1H), 3.14 (br. s, 3H), 3.09 – 2.98 (m, 1H); 1 exchangeable proton not observed.
HRMS m/z calc. for $C_{30}H_{25}N_2O_7$ $[M+H]^+$ 525.1662 found 525.1677.

methyl (1*R*,3*R*)-2-acryloyl-1,7-bis(benzo[*d*][1,3]dioxol-5-yl)-2,3,4,9-tetrahydro-1*H*-pyrido[3,4-*b*]indole-3-carboxylate (WX-72c (WX-02-435))
$^{1}$H NMR (400 MHz, $CD_3OD$) δ 7.55 (d, *J* = 8.2 Hz, 1H), 7.44 (s, 1H), 7.27 (dd, *J* = 8.3, 1.6 Hz, 1H), 7.15 – 7.07 (m, 2H), 7.00 (br. s, 1H), 6.95 – 6.79 (m, 3H), 6.77 – 6.65 (m, 1H), 6.64 – 6.56 (m, 1H), 6.27 (d, *J* = 16.8 Hz, 1H), 5.96 (s, 2H), 5.90 (s, 2H), 5.82 (d, *J* = 10.9 Hz, 1H), 5.40 – 5.23 (m, 1H), 3.62 (d, *J* = 16.1 Hz, 1H), 3.14 (br. s, 3H), 3.08 – 3.00 (m, 1H); 1 exchangeable proton not observed.
HRMS m/z calc. for $C_{30}H_{25}N_2O_7$ $[M+H]^+$ 525.1662 found 525.1654.

methyl (1*R*,3*S*)-2-acryloyl-1,7-bis(benzo[*d*][1,3]dioxol-5-yl)-2,3,4,9-tetrahydro-1*H*-pyrido[3,4-*b*]indole-3-carboxylate (WX-72b (WX-02-436))
$^{1}$H NMR (400 MHz, $CD_3OD$) δ 7.47 (d, *J* = 8.3 Hz, 1H), 7.42 (br. s, 1H), 7.22 (d, *J* = 8.3 Hz, 1H), 7.11 – 7.02 (m, 2H), 6.97 – 6.66 (m, 5H), 6.31 – 6.00 (m, 2H), 5.95 (s, 2H), 5.93 – 5.81 (m, 2H), 5.70 (dd, *J* = 10.6, 1.8 Hz, 1H), 5.46 (s, 0.3H), 5.07 – 4.95 (m, 0.7H), 3.70 – 3.52 (m, 3.3H), 3.51 – 3.37 (m, 1H), 3.25 – 3.14 (m, 0.7H); 1 exchangeable proton not observed; ~7:3 mixture of rotamers.
HRMS m/z calc. for $C_{30}H_{25}N_2O_7$ $[M+H]^+$ 525.1662 found 525.1674.

methyl (1*S*,3*R*)-2-acryloyl-1,7-bis(benzo[*d*][1,3]dioxol-5-yl)-2,3,4,9-tetrahydro-1*H*-pyrido[3,4-*b*]indole-3-carboxylate (WX-72d (WX-02-437))
$^{1}$H NMR (400 MHz, $CD_3OD$) δ 7.47 (d, *J* = 8.2 Hz, 1H), 7.42 (br. s, 1H), 7.22 (d, *J* = 8.0 Hz, 1H), 7.11 – 7.02 (m, 2H), 6.99 – 6.66 (m, 5H), 6.32 – 6.04 (m, 2H), 5.95 (s, 2H), 5.94 – 5.78 (m, 2H), 5.71 (dd, *J* = 10.6, 1.8 Hz, 1H), 5.53 – 5.39 (m, 0.3H), 5.09 – 4.94 (m, 0.7H), 3.75 – 3.51 (m, 3.3H), 3.51 – 3.38 (m, 1H), 3.27 – 3.12 (m, 0.7H); 1 exchangeable proton not observed; ~7:3 mixture of rotamers.
HRMS m/z calc. for $C_{30}H_{25}N_2O_7$ $[M+H]^+$ 525.1662 found 525.1663.

methyl (1*S*,3*S*)-2-acryloyl-1-(benzo[*d*][1,3]dioxol-5-yl)-7-(4-fluorophenoxy)-2,3,4,9-tetrahydro-1*H*-pyrido[3,4-*b*]indole-3-carboxylate (WX-73a (WX-02-438))
$^{1}$H NMR (400 MHz, $CD_3OD$) δ 7.51 (d, *J* = 8.5 Hz, 1H), 7.11 – 6.86 (m, 7H), 6.84 – 6.76 (m, 2H), 6.74 – 6.66 (m, 1H), 6.65 – 6.54 (m, 1H), 6.27 (d, *J* = 16.9 Hz, 1H), 5.91 (s, 2H), 5.83 (d, *J* = 11.0 Hz, 1H), 5.39 – 5.25 (m, 1H), 3.66 – 3.55 (m, 1H), 3.15 (br. s, 3H), 3.08 – 2.97 (m, 1H); 1 exchangeable proton not observed.

HRMS m/z calc. for $C_{29}H_{24}FN_2O_6$ $[M+H]^+$ 515.1618 found 515.1613.

methyl (1*R*,3*R*)-2-acryloyl-1-(benzo[*d*][1,3]dioxol-5-yl)-7-(4-fluorophenoxy)-2,3,4,9-tetrahydro-1*H*-pyrido[3,4-*b*]indole-3-carboxylate (WX-73c (WX-02-439))
$^1$H NMR (400 MHz, $CD_3OD$) δ 7.51 (d, *J* = 8.5 Hz, 1H), 7.11 – 6.86 (m, 7H), 6.83 – 6.76 (m, 2H), 6.74 – 6.67 (m, 1H), 6.64 – 6.55 (m, 1H), 6.27 (d, *J* = 16.8 Hz, 1H), 5.91 (s, 2H), 5.83 (d, *J* = 10.8 Hz, 1H), 5.38 – 5.26 (m, 1H), 3.60 (d, *J* = 15.8 Hz, 1H), 3.14 (br. s, 3H), 3.02 (m, 1H); 1 exchangeable proton not observed.
HRMS m/z calc. for $C_{29}H_{24}FN_2O_6$ $[M+H]^+$ 515.1618 found 515.1624.

methyl (1*R*,3*S*)-2-acryloyl-1-(benzo[*d*][1,3]dioxol-5-yl)-7-(4-fluorophenoxy)-2,3,4,9-tetrahydro-1*H*-pyrido[3,4-*b*]indole-3-carboxylate (WX-73b (WX-02-440))
$^1$H NMR (400 MHz, $CD_3OD$) δ 7.42 (d, *J* = 8.5 Hz, 1H), 7.00 (t, *J* = 8.6 Hz, 2H), 6.95 – 6.85 (m, 5H), 6.79 (dd, *J* = 16.7, 10.6 Hz, 2H), 6.73 (dd, *J* = 8.5, 2.1 Hz, 1H), 6.31 – 5.99 (m, 2H), 5.98 – 5.77 (m, 2H), 5.70 (dd, *J* = 10.6, 1.9 Hz, 1H), 5.57 – 5.38 (m, 0.3H), 5.12 – 4.95 (m, 0.7H), 3.75 – 3.53 (m, 3.3H), 3.53 – 3.37 (m, 1H), 3.27 – 3.09 (m, 0.7H); 1 exchangeable proton not observed; ~7:3 mixture of rotamers.
HRMS m/z calc. for $C_{29}H_{24}FN_2O_6$ $[M+H]^+$ 515.1618 found 515.1632.

methyl (1*S*,3*R*)-2-acryloyl-1-(benzo[*d*][1,3]dioxol-5-yl)-7-(4-fluorophenoxy)-2,3,4,9-tetrahydro-1*H*-pyrido[3,4-*b*]indole-3-carboxylate (WX-73d (WX-02-441))
$^1$H NMR (400 MHz, $CD_3OD$) δ 7.42 (d, *J* = 8.5 Hz, 1H), 7.01 (t, *J* = 8.6 Hz, 2H), 6.96 – 6.84 (m, 5H), 6.79 (dd, *J* = 16.7, 10.6 Hz, 2H), 6.73 (dd, *J* = 8.7, 1.2 Hz, 1H), 6.27 – 6.01 (m, 2H), 6.00 – 5.79 (m, 2H), 5.70 (dd, *J* = 10.6, 1.9 Hz, 1H), 5.52 – 5.41 (m, 0.3H), 5.12 – 4.97 (m, 0.7H), 3.71 – 3.52 (m, 3.3H), 3.50 – 3.37 (m, 1H), 3.27 – 3.14 (m, 0.7H); 1 exchangeable proton not observed; ~7:3 mixture of rotamers.
HRMS m/z calc. for $C_{29}H_{24}FN_2O_6$ $[M+H]^+$ 515.1618 found 515.1633.

methyl (1*S*,3*S*)-2-acryloyl-1-(benzo[*d*][1,3]dioxol-5-yl)-6-(1-methyl-1*H*-1,2,3-triazol-4-yl)-2,3,4,9-tetrahydro-1*H*-pyrido[3,4-*b*]indole-3-carboxylate (WX-64a (WX-02-687))
$^1$H NMR (400 MHz, $CD_3OD$) δ 8.21 (s, 1H), 8.01 (s, 1H), 7.59 (dd, *J* = 8.4, 1.7 Hz, 1H), 7.35 (d, *J* = 8.4 Hz, 1H), 7.01 (br. s, 1H), 6.97 – 6.86 (m, 1H), 6.82 (br. s, 1H), 6.69 (d, *J* = 8.1 Hz, 1H), 6.62 – 6.57 (m, 1H), 6.28 (d, *J* = 17.0 Hz, 1H), 5.91 (s, 2H), 5.83 (d, *J* = 10.9 Hz, 1H), 5.43 – 5.27 (m, 1H), 4.16 (s, 3H), 3.67 (d, *J* = 16.1 Hz, 1H), 3.24 – 2.93 (m, 4H); 1 exchangeable proton not observed.
HRMS m/z calc. for $C_{26}H_{24}N_5O_5$ $[M+H]^+$ 486.1777 found 486.1785.

methyl (1*R*,3*R*)-2-acryloyl-1-(benzo[*d*][1,3]dioxol-5-yl)-6-(1-methyl-1*H*-1,2,3-triazol-4-yl)-2,3,4,9-tetrahydro-1*H*-pyrido[3,4-*b*]indole-3-carboxylate (WX-64c (WX-02-688))
$^1$H NMR (400 MHz, $CD_3OD$) δ 8.21 (s, 1H), 8.01 (s, 1H), 7.60 (dd, *J* = 8.4, 1.6 Hz, 1H), 7.36 (d, *J* = 8.5 Hz, 1H), 7.01 (br. s, 1H), 6.93 – 6.89 (m, 1H), 6.82 (br. s, 1H), 6.70 (d, *J* = 7.8 Hz, 1H), 6.65

– 6.55 (m, 1H), 6.28 (d, *J* = 17.2 Hz, 1H), 5.91 (s, 2H), 5.84 (d, *J* = 10.9 Hz, 1H), 5.41 – 5.28 (m, 1H), 4.16 (s, 3H), 3.67 (d, *J* = 16.1 Hz, 1H), 3.26 – 3.00 (m, 4H); 1 exchangeable proton not observed.
HRMS m/z calc. for $C_{26}H_{24}N_5O_5$ $[M+H]^+$ 486.1777 found 486.1788.

methyl (1*R*,3*S*)-2-acryloyl-1-(benzo[*d*][1,3]dioxol-5-yl)-6-(1-methyl-1*H*-1,2,3-triazol-4-yl)-2,3,4,9-tetrahydro-1*H*-pyrido[3,4-*b*]indole-3-carboxylate (WX-64b (WX-02-689))
$^1$H NMR (400 MHz, $CD_3OD$) δ 8.18 (s, 1H), 7.92 (s, 1H), 7.63 – 7.47 (m, 1H), 7.40 – 7.24 (m, 1H), 6.92 (d, *J* = 8.4 Hz, 2H), 6.81 (dd, *J* = 16.7, 10.6 Hz, 2H), 6.35 – 6.05 (m, 2H), 6.05 – 5.78 (m, 2H), 5.71 (dd, *J* = 10.6, 1.8 Hz, 1H), 5.57 – 5.46 (m, 0.3H), 5.13 – 5.03 (m, 0.7H), 4.15 (s, 3H), 3.74 – 3.40 (m, 4.3H), 3.31 – 3.16 (m, 0.7H); 1 exchangeable proton not observed; ~7:3 mixture of rotamers.
HRMS m/z calc. for $C_{26}H_{24}N_5O_5$ $[M+H]^+$ 486.1777 found 486.1785.

methyl (1*S*,3*R*)-2-acryloyl-1-(benzo[*d*][1,3]dioxol-5-yl)-6-(1-methyl-1*H*-1,2,3-triazol-4-yl)-2,3,4,9-tetrahydro-1*H*-pyrido[3,4-*b*]indole-3-carboxylate (WX-64d (WX-02-690))
$^1$H NMR (400 MHz, $CD_3OD$) δ 8.18 (s, 1H), 7.92 (s, 1H), 7.65 – 7.44 (m, 1H), 7.40 – 7.23 (m, 1H), 6.92 (d, *J* = 8.3 Hz, 2H), 6.80 (dd, *J* = 16.7, 10.6 Hz, 2H), 6.35 – 6.03 (m, 2H), 6.01 – 5.80 (m, 2H), 5.71 (dd, *J* = 10.6, 1.8 Hz, 1H), 5.57 – 5.44 (m, 0.3H), 5.16 – 5.00 (m, 0.7H), 4.15 (s, 3H), 3.74 – 3.45 (m, 4.3H), 3.30 (d, *J* = 1.8 Hz, 0.7H); 1 exchangeable proton not observed; ~7:3 mixture of rotamers.
HRMS m/z calc. for $C_{26}H_{24}N_5O_5$ $[M+H]^+$ 486.1777 found 486.1799.

methyl (1*S*,3*S*)-2-acryloyl-1-(benzo[*d*][1,3]dioxol-5-yl)-7-(1-methyl-1*H*-1,2,3-triazol-4-yl)-2,3,4,9-tetrahydro-1*H*-pyrido[3,4-*b*]indole-3-carboxylate (WX-74a (WX-02-691))
$^1$H NMR (400 MHz, $CD_3OD$) δ 8.21 (s, 1H), 7.77 (s, 1H), 7.60 (d, *J* = 8.2 Hz, 1H), 7.50 (d, *J* = 8.2 Hz, 1H), 7.15 – 6.99 (m, 1H), 6.99 – 6.79 (m, 2H), 6.77 – 6.66 (m, 1H), 6.65 – 6.46 (m, 1H), 6.42 – 6.22 (m, 1H), 5.91 (s, 2H), 5.84 (d, *J* = 10.7 Hz, 1H), 5.40 – 5.29 (m, 1H), 4.16 (s, 3H), 3.63 (d, *J* = 15.9 Hz, 1H), 3.14 (br. s, 3H), 3.10 – 2.98 (m, 1H); 1 exchangeable proton not observed.
HRMS m/z calc. for $C_{26}H_{24}N_5O_5$ $[M+H]^+$ 486.1777 found 486.1781.

methyl (1*R*,3*R*)-2-acryloyl-1-(benzo[*d*][1,3]dioxol-5-yl)-7-(1-methyl-1*H*-1,2,3-triazol-4-yl)-2,3,4,9-tetrahydro-1*H*-pyrido[3,4-*b*]indole-3-carboxylate (WX-74c (WX-02-692))
$^1$H NMR (400 MHz, $CD_3OD$) δ 8.19 (s, 1H), 7.77 (s, 1H), 7.59 (d, *J* = 8.1 Hz, 1H), 7.50 (dd, *J* = 8.3, 1.4 Hz, 1H), 7.15 – 6.98 (m, 1H), 6.98 – 6.77 (m, 2H), 6.76 – 6.66 (m, 1H), 6.64 – 6.48 (m, 1H), 6.38 – 6.21 (m, 1H), 5.91 (s, 2H), 5.83 (d, *J* = 10.6 Hz, 1H), 5.43 – 5.19 (m, 1H), 4.15 (s, 3H), 3.63 (d, *J* = 15.7 Hz, 1H), 3.15 (br. s, 3H), 3.12 – 2.96 (m, 1H); 1 exchangeable proton not observed.
HRMS m/z calc. for $C_{26}H_{24}N_5O_5$ $[M+H]^+$ 486.1777 found 486.1787.

methyl (1*R*,3*S*)-2-acryloyl-1-(benzo[*d*][1,3]dioxol-5-yl)-7-(1-methyl-1*H*-1,2,3-triazol-4-yl)-2,3,4,9-tetrahydro-1*H*-pyrido[3,4-*b*]indole-3-carboxylate (WX-74b (WX-02-693))
$^1$H NMR (400 MHz, $CD_3OD$) δ 8.16 (s, 1H), 7.85 – 7.65 (m, 1H), 7.51 (d, *J* = 8.2 Hz, 1H), 7.44 (dd, *J* = 8.3, 1.5 Hz, 1H), 7.01 – 6.86 (m, 2H), 6.86 – 6.65 (m, 2H), 6.34 – 6.03 (m, 2H), 6.02 – 5.77 (m, 2H), 5.71 (dd, *J* = 10.6, 1.9 Hz, 1H), 5.57 – 5.39 (m, 0.3H), 5.11 – 4.97 (m, 0.7H), 4.12 (s, 3H), 3.74 – 3.52 (m, 3.3H), 3.52 – 3.37 (m, 1H), 3.27 – 3.14 (m, 0.7H); 1 exchangeable proton not observed; ~7:3 mixture of rotamers.
HRMS m/z calc. for $C_{26}H_{24}N_5O_5$ $[M+H]^+$ 486.1777 found 486.1786.

methyl (1*S*,3*R*)-2-acryloyl-1-(benzo[*d*][1,3]dioxol-5-yl)-7-(1-methyl-1*H*-1,2,3-triazol-4-yl)-2,3,4,9-tetrahydro-1*H*-pyrido[3,4-*b*]indole-3-carboxylate (WX-74d (WX-02-694))
$^1$H NMR (400 MHz, $CD_3OD$) δ 8.17 (s, 1H), 7.87 – 7.63 (m, 1H), 7.51 (d, *J* = 8.3 Hz, 1H), 7.44 (dd, *J* = 8.3, 1.4 Hz, 1H), 7.06 – 6.88 (m, 2H), 6.88 – 6.67 (m, 2H), 6.38 – 6.03 (m, 2H), 6.03 – 5.80 (m, 2H), 5.71 (dd, *J* = 10.5, 1.8 Hz, 1H), 5.58 – 5.40 (m, 0.3H), 5.11 – 4.98 (m, 0.7H), 4.13 (s, 3H), 3.75 – 3.52 (m, 3.3H), 3.51 – 3.40 (m, 1H), 3.28 – 3.13 (m, 0.7H); 1 exchangeable proton not observed; ~7:3 mixture of rotamers.
HRMS m/z calc. for $C_{26}H_{24}N_5O_5$ $[M+H]^+$ 486.1777 found 486.1797.

methyl (1*S*,3*S*)-2-acryloyl-7-(benzo[*d*][1,3]dioxol-5-yl)-1-(3-methoxy-4-(prop-2-yn-1-yloxy)phenyl)-2,3,4,9-tetrahydro-1*H*-pyrido[3,4-*b*]indole-3-carboxylate (WX-72a-yne (WX-02-891))
$^1$H NMR (400 MHz, $CD_3OD$) δ 7.56 (d, *J* = 8.2 Hz, 1H), 7.45 (s, 1H), 7.28 (dd, *J* = 8.1, 1.6 Hz, 1H), 7.15 – 7.08 (m, 2H), 7.06 (br. s, 1H), 7.00 – 6.80 (m, 4H), 6.72 (d, *J* = 8.4 Hz, 0.8H), 6.58 – 6.47 (m, 0.2H), 6.45 – 6.20 (m, 1H), 5.96 (s, 2H), 5.93 – 5.79 (m, 1H), 5.72 – 5.59 (m, 0.2H), 5.33 (d, *J* = 6.9 Hz, 0.8H), 4.72 (d, *J* = 2.4 Hz, 2H), 3.71 (s, 3H), 3.67 – 3.59 (m, 0.8H), 3.49 – 3.39 (m, 0.2H), 3.28 – 3.17 (m, 0.6H), 3.15 – 2.98 (m, 3.4H), 2.90 (d, *J* = 2.4 Hz, 1H); 1 exchangeable proton not observed; ~8:2 mixture of rotamers.
HRMS m/z calc. for $C_{33}H_{29}N_2O_7$ $[M+H]^+$ 565.1975 found 565.1983.

methyl (1*R*,3*R*)-2-acryloyl-7-(benzo[*d*][1,3]dioxol-5-yl)-1-(3-methoxy-4-(prop-2-yn-1-yloxy)phenyl)-2,3,4,9-tetrahydro-1*H*-pyrido[3,4-*b*]indole-3-carboxylate (WX-72c-yne (WX-02-892))
$^1$H NMR (400 MHz, $CD_3OD$) δ 7.55 (d, *J* = 8.2 Hz, 1H), 7.45 (s, 1H), 7.27 (dd, *J* = 8.2, 1.7 Hz, 1H), 7.15 – 7.08 (m, 2H), 7.05 (br. s, 1H), 7.01 – 6.80 (m, 4H), 6.71 (d, *J* = 8.4 Hz, 0.8H), 6.59 – 6.46 (m, 0.2H), 6.42 – 6.22 (m, 1H), 5.96 (s, 2H), 5.93 – 5.76 (m, 1H), 5.71 – 5.58 (m, 0.2H), 5.31 (d, *J* = 6.7 Hz, 0.8H), 4.71 (d, *J* = 2.4 Hz, 2H), 3.71 (s, 3H), 3.66 – 3.57 (m, 0.8H), 3.49 – 3.38 (m, 0.2H), 3.28 – 3.19 (m, 0.6H), 3.16 – 2.97 (m, 3.4H), 2.90 (t, *J* = 2.4 Hz, 1H); 1 exchangeable proton not observed; ~8:2 mixture of rotamers.
HRMS m/z calc. for $C_{33}H_{29}N_2O_7$ $[M+H]^+$ 565.1975 found 565.1962.

methyl (1*R*,3*S*)-2-acryloyl-7-(benzo[*d*][1,3]dioxol-5-yl)-1-(3-methoxy-4-(prop-2-yn-1-yloxy)phenyl)-2,3,4,9-tetrahydro-1*H*-pyrido[3,4-*b*]indole-3-carboxylate (WX-72b-yne (WX-02-893))
$^1$H NMR (400 MHz, $CD_3OD$) δ 7.55 – 7.31 (m, 2H), 7.22 (d, *J* = 8.2 Hz, 1H), 7.17 – 6.90 (m, 5H), 6.89 – 6.74 (m, 2H), 6.42 – 6.05 (m, 2H), 5.95 (s, 2H), 5.71 (dd, *J* = 10.6, 1.8 Hz, 1H), 5.60 – 5.38 (m, 0.3H), 5.05 – 4.92 (m, 0.7H), 4.70 (br. s, 2H), 3.83 (s, 3H), 3.73 – 3.52 (m, 3.3H), 3.51 – 3.40 (m, 1H), 3.26 – 3.12 (m, 0.7H), 2.89 (br. s, 1H); 1 exchangeable proton not observed; ~7:3 mixture of rotamers.
HRMS m/z calc. for $C_{33}H_{29}N_2O_7$ $[M+H]^+$ 565.1975 found 565.1991.

methyl (1*S*,3*R*)-2-acryloyl-7-(benzo[*d*][1,3]dioxol-5-yl)-1-(3-methoxy-4-(prop-2-yn-1-yloxy)phenyl)-2,3,4,9-tetrahydro-1*H*-pyrido[3,4-*b*]indole-3-carboxylate (WX-72d-yne (WX-02-894))
$^1$H NMR (400 MHz, $CD_3OD$) δ 7.56 – 7.33 (m, 2H), 7.21 (d, *J* = 8.2 Hz, 1H), 7.15 – 6.89 (m, 5H), 6.87 – 6.71 (m, 2H), 6.36 – 6.04 (m, 2H), 5.94 (s, 2H), 5.70 (d, *J* = 10.6 Hz, 1H), 5.57 – 5.38 (m, 0.3H), 5.02 – 4.90 (m, 0.7H), 4.68 (br. s, 2H), 3.82 (s, 3H), 3.76 – 3.51 (m, 3.3H), 3.51 – 3.39 (m, 1H), 3.27 – 3.09 (m, 0.7H), 2.89 (br. s, 1H); 1 exchangeable proton not observed; ~7:3 mixture of rotamers.
HRMS m/z calc. for $C_{33}H_{29}N_2O_7$ $[M+H]^+$ 565.1975 found 565.1976.

(1*S*,3*S*)-2-acryloyl-1-(benzo[*d*][1,3]dioxol-5-yl)-7-(4-fluorophenoxy)-*N*-(prop-2-yn-1-yl)-2,3,4,9-tetrahydro-1*H*-pyrido[3,4-*b*]indole-3-carboxamide (WX-73a-yne (WX-02-905))
$^1$H NMR (400 MHz, $CD_3OD$) δ 7.51 (d, *J* = 8.6 Hz, 1H), 7.08 – 6.84 (m, 8H), 6.83 – 6.68 (m, 3H), 6.32 (d, *J* = 16.8 Hz, 1H), 5.92 (s, 2H), 5.85 (dd, *J* = 10.9, 1.5 Hz, 1H), 5.40 – 5.18 (m, 1H), 3.77 – 3.40 (m, 2H), 3.27 – 3.12 (m, 1H), 3.03 – 2.89 (m, 1H), 2.49 (s, 1H); 2 exchangeable protons not observed.
HRMS m/z calc. for $C_{31}H_{25}FN_3O_5$ $[M+H]^+$ 538.1778 found 538.1803.

(1*R*,3*R*)-2-acryloyl-1-(benzo[*d*][1,3]dioxol-5-yl)-7-(4-fluorophenoxy)-*N*-(prop-2-yn-1-yl)-2,3,4,9-tetrahydro-1*H*-pyrido[3,4-*b*]indole-3-carboxamide (WX-73c-yne (WX-02-906))
$^1$H NMR (400 MHz, $CD_3OD$) δ 7.50 (d, *J* = 8.5 Hz, 1H), 7.11 – 6.84 (m, 8H), 6.82 – 6.67 (m, 3H), 6.31 (d, *J* = 16.7 Hz, 1H), 5.91 (s, 2H), 5.85 (dd, *J* = 10.8, 1.9 Hz, 1H), 5.46 – 5.17 (m, 1H), 3.84 – 3.41 (m, 2H), 3.29 – 3.12 (m, 1H), 3.06 – 2.90 (m, 1H), 2.49 (s, 1H); 2 exchangeable protons not observed.
HRMS m/z calc. for $C_{31}H_{25}FN_3O_5$ $[M+H]^+$ 538.1778 found 538.1799.

(1*R*,3*S*)-2-acryloyl-1-(benzo[*d*][1,3]dioxol-5-yl)-7-(4-fluorophenoxy)-*N*-(prop-2-yn-1-yl)-2,3,4,9-tetrahydro-1*H*-pyrido[3,4-*b*]indole-3-carboxamide (WX-73b-yne (WX-02-907))
$^1$H NMR (400 MHz, $CD_3OD$) δ 7.39 (d, *J* = 8.5 Hz, 1H), 7.00 (t, *J* = 8.7 Hz, 2H), 6.94 – 6.85 (m, 5H), 6.82 – 6.65 (m, 3H), 6.27 (s, 1H), 6.16 (d, *J* = 16.7 Hz, 1H), 5.89 (s, 2H), 5.78 – 5.55 (m, 1H), 5.31 (br. s, 1H), 3.84 (s, 2H), 3.55 – 3.38 (m, 2H), 2.45 (t, *J* = 2.5 Hz, 1H); 2 exchangeable protons not observed.

HRMS m/z calc. for $C_{31}H_{25}FN_3O_5$ $[M+H]^+$ 538.1778 found 538.1785.

(1*S*,3*R*)-2-acryloyl-1-(benzo[*d*][1,3]dioxol-5-yl)-7-(4-fluorophenoxy)-*N*-(prop-2-yn-1-yl)-2,3,4,9-tetrahydro-1*H*-pyrido[3,4-*b*]indole-3-carboxamide (WX-73d-yne (WX-02-908))
$^1$H NMR (400 MHz, $CD_3OD$) δ 7.38 (d, *J* = 8.5 Hz, 1H), 6.99 (t, *J* = 8.7 Hz, 2H), 6.93 – 6.84 (m, 5H), 6.83 – 6.65 (m, 3H), 6.27 (s, 1H), 6.16 (d, *J* = 16.5 Hz, 1H), 5.88 (s, 2H), 5.78 – 5.53 (m, 1H), 5.31 (br. s, 1H), 3.84 (s, 2H), 3.55 – 3.37 (m, 2H), 2.44 (t, *J* = 2.5 Hz, 1H); 2 exchangeable protons not observed.
HRMS m/z calc. for $C_{31}H_{25}FN_3O_5$ $[M+H]^+$ 538.1778 found 538.1787.

(1*S*,3*S*)-2-acryloyl-1,6-bis(benzo[*d*][1,3]dioxol-5-yl)-*N*-(prop-2-yn-1-yl)-2,3,4,9-tetrahydro-1*H*-pyrido[3,4-*b*]indole-3-carboxamide (WX-62a-yne (WX-02-909))
$^1$H NMR (400 MHz, $CD_3OD$) δ 7.68 (d, *J* = 1.3 Hz, 1H), 7.32 (d, *J* = 1.2 Hz, 2H), 7.21 – 6.65 (m, 8H), 6.32 (d, *J* = 16.7 Hz, 1H), 5.97 (s, 2H), 5.94 – 5.90 (m, 2H), 5.85 (d, *J* = 10.7 Hz, 1H), 5.42 – 5.14 (m, 1H), 3.80 – 3.48 (m, 2H), 3.32 – 3.11 (m, 1H), 3.09 – 2.97 (m, 1H), 2.49 (s, 1H); 2 exchangeable protons not observed.
HRMS m/z calc. for $C_{32}H_{26}N_3O_6$ $[M+H]^+$ 548.1822 found 548.1837.

(1*R*,3*R*)-2-acryloyl-1,6-bis(benzo[*d*][1,3]dioxol-5-yl)-*N*-(prop-2-yn-1-yl)-2,3,4,9-tetrahydro-1*H*-pyrido[3,4-*b*]indole-3-carboxamide (WX-62c-yne (WX-02-910))
$^1$H NMR (400 MHz, $CD_3OD$) δ 7.67 (t, *J* = 1.3 Hz, 1H), 7.31 (d, *J* = 1.3 Hz, 2H), 7.19 – 6.57 (m, 8H), 6.31 (d, *J* = 16.8 Hz, 1H), 5.96 (s, 2H), 5.93 – 5.90 (m, 2H), 5.84 (dd, *J* = 10.7, 1.9 Hz, 1H), 5.41 – 5.14 (m, 1H), 3.65 (s, 2H), 3.13 (s, 1H), 3.10 – 2.93 (m, 1H), 2.49 (s, 1H); 2 exchangeable protons not observed.
HRMS m/z calc. for $C_{32}H_{26}N_3O_6$ $[M+H]^+$ 548.1822 found 548.1818.

(1*R*,3*S*)-2-acryloyl-1,6-bis(benzo[*d*][1,3]dioxol-5-yl)-*N*-(prop-2-yn-1-yl)-2,3,4,9-tetrahydro-1*H*-pyrido[3,4-*b*]indole-3-carboxamide (WX-62b-yne (WX-02-911))
$^1$H NMR (400 MHz, $CD_3OD$) δ 7.55 (d, *J* = 1.3 Hz, 1H), 7.32 – 7.17 (m, 2H), 7.11 – 7.01 (m, 2H), 6.99 – 6.90 (m, 2H), 6.84 (d, *J* = 8.0 Hz, 1H), 6.79 – 6.66 (m, 2H), 6.31 (s, 1H), 6.21 – 6.13 (m, 1H), 5.94 (s, 2H), 5.88 (br. s, 2H), 5.75 – 5.55 (m, 1H), 5.31 (br. s, 1H), 3.90 – 3.73 (m, 2H), 3.58 – 3.34 (m, 2H), 2.40 (t, *J* = 2.5 Hz, 1H); 2 exchangeable protons not observed.
HRMS m/z calc. for $C_{32}H_{26}N_3O_6$ $[M+H]^+$ 548.1822 found 548.1813.

(1*S*,3*R*)-2-acryloyl-1,6-bis(benzo[*d*][1,3]dioxol-5-yl)-*N*-(prop-2-yn-1-yl)-2,3,4,9-tetrahydro-1*H*-pyrido[3,4-*b*]indole-3-carboxamide (WX-62d-yne (WX-02-912))
$^1$H NMR (400 MHz, $CD_3OD$) δ 7.55 (s, 1H), 7.34 – 7.20 (m, 2H), 7.10 – 7.01 (m, 2H), 7.00 – 6.89 (m, 2H), 6.84 (d, *J* = 7.9 Hz, 1H), 6.80 – 6.65 (m, 2H), 6.31 (s, 1H), 6.17 (d, *J* = 16.6 Hz, 1H), 5.94 (s, 2H), 5.88 (br. s, 2H), 5.75 – 5.57 (m, 1H), 5.31 (br. s, 1H), 3.93 – 3.69 (m, 2H), 3.59 – 3.34 (m, 2H), 2.39 (t, *J* = 2.5 Hz, 1H); 2 exchangeable protons not observed.
HRMS m/z calc. for $C_{32}H_{26}N_3O_6$ $[M+H]^+$ 548.1822 found 548.1814.

**Analytical data: NMR spectra**

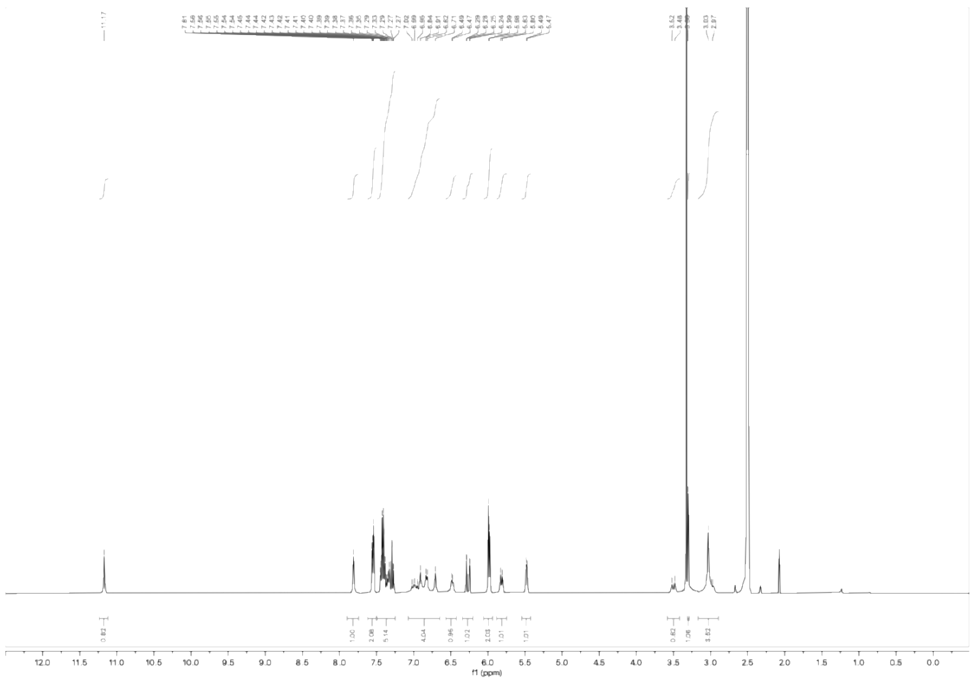

$^1$H NMR spectrum of WX-61a (WX-02-418) (400 MHz, DMSO-$d_6$)

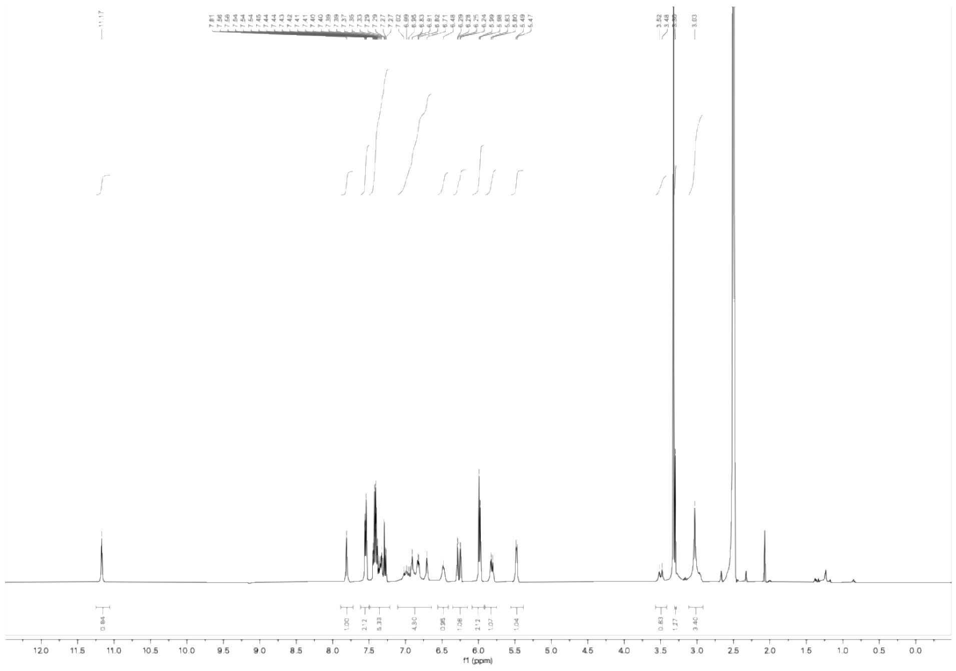

$^1$H NMR spectrum of WX-61c (WX-02-420) (400 MHz, DMSO-$d_6$)

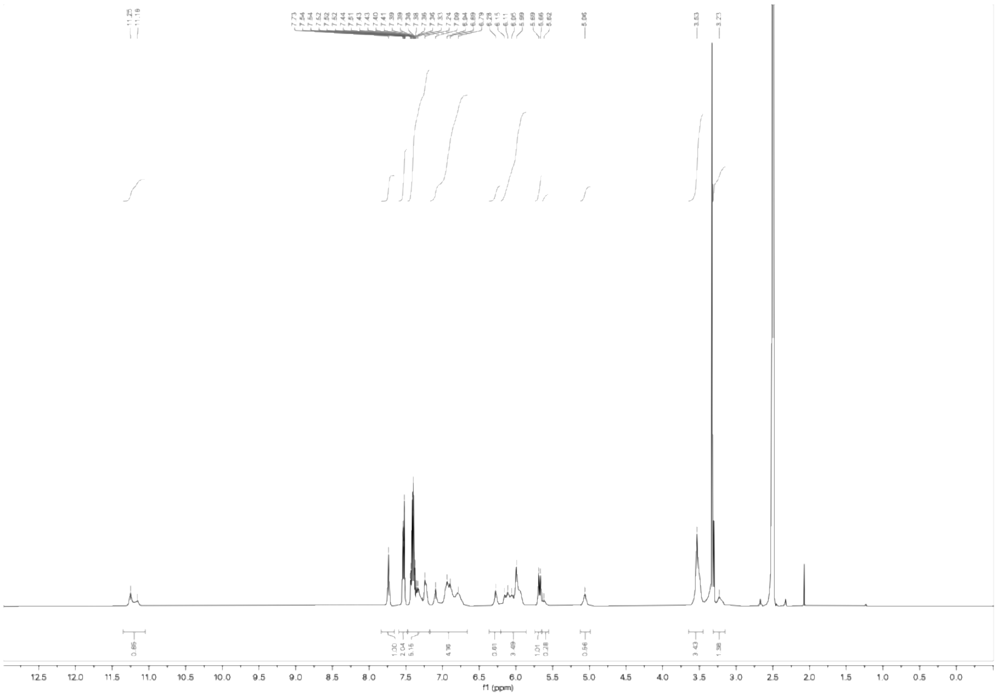


$^1$H NMR spectrum of WX-61b (WX-02-419) (400 MHz, DMSO-$d_6$)

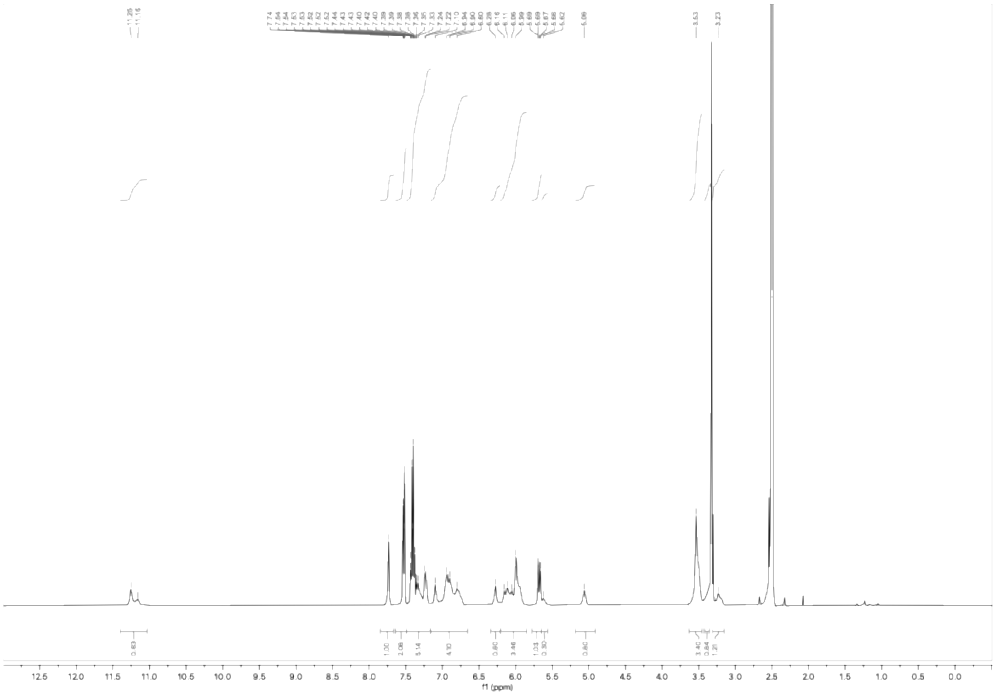


$^1$H NMR spectrum of WX-61d (WX-02-421) (400 MHz, DMSO-$d_6$)

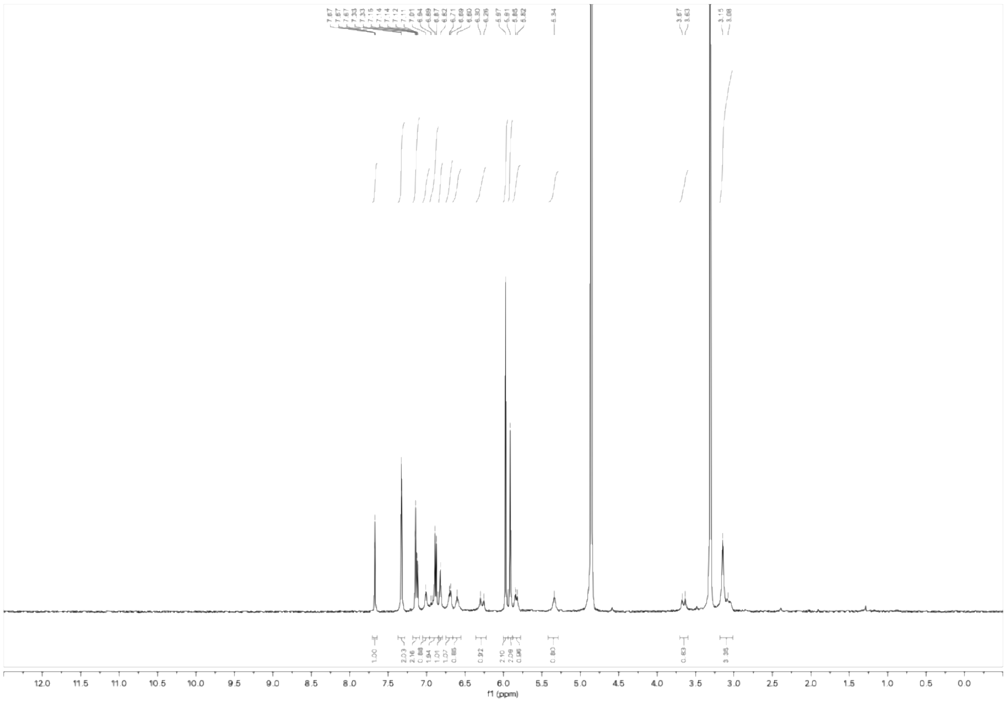

$^1H$ NMR spectrum of WX-62a (WX-02-422) (400 MHz, $CD_3OD$)

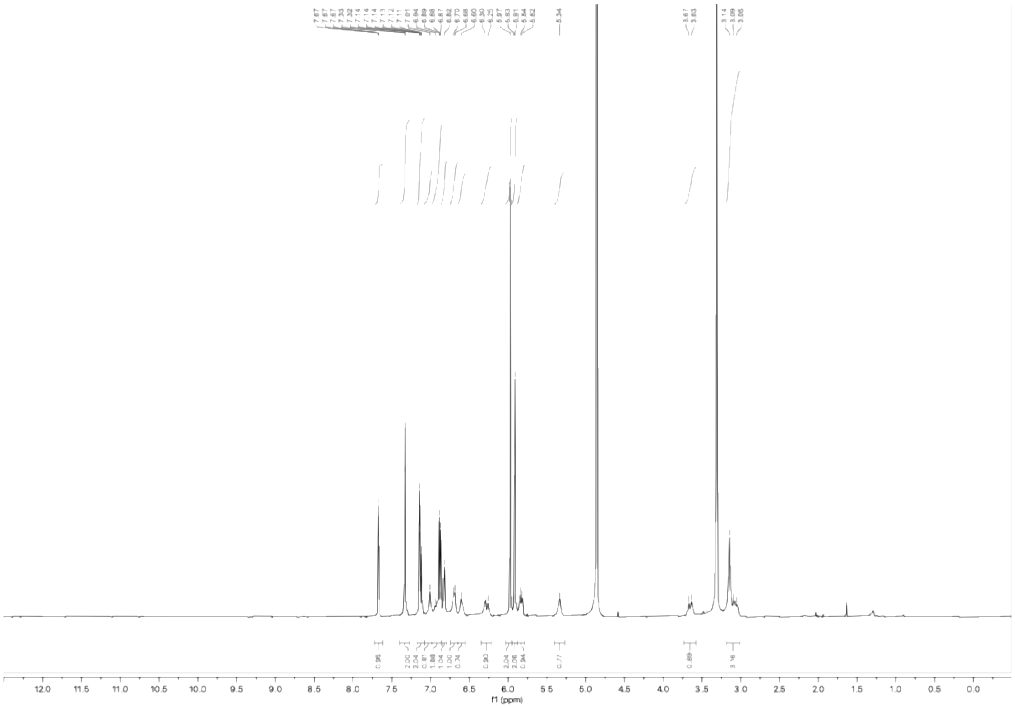

$^1H$ NMR spectrum of WX-62c (WX-02-424) (400 MHz, $CD_3OD$)

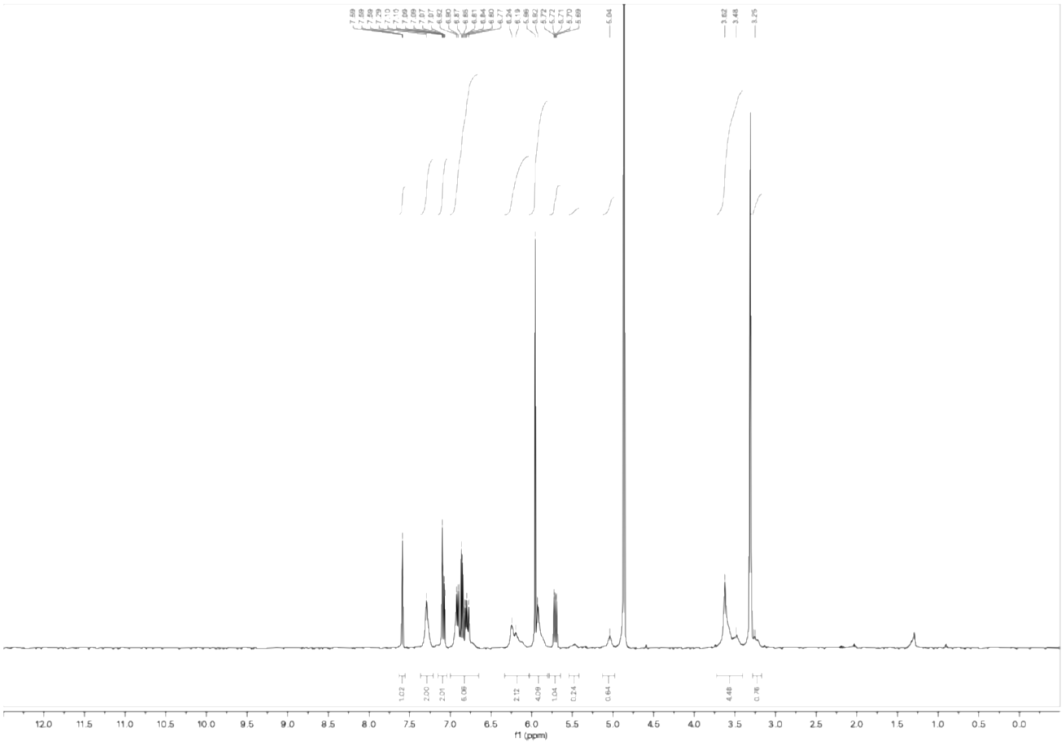

[1]H NMR spectrum of WX-62b (WX-02-423) (400 MHz, $CD_3OD$)

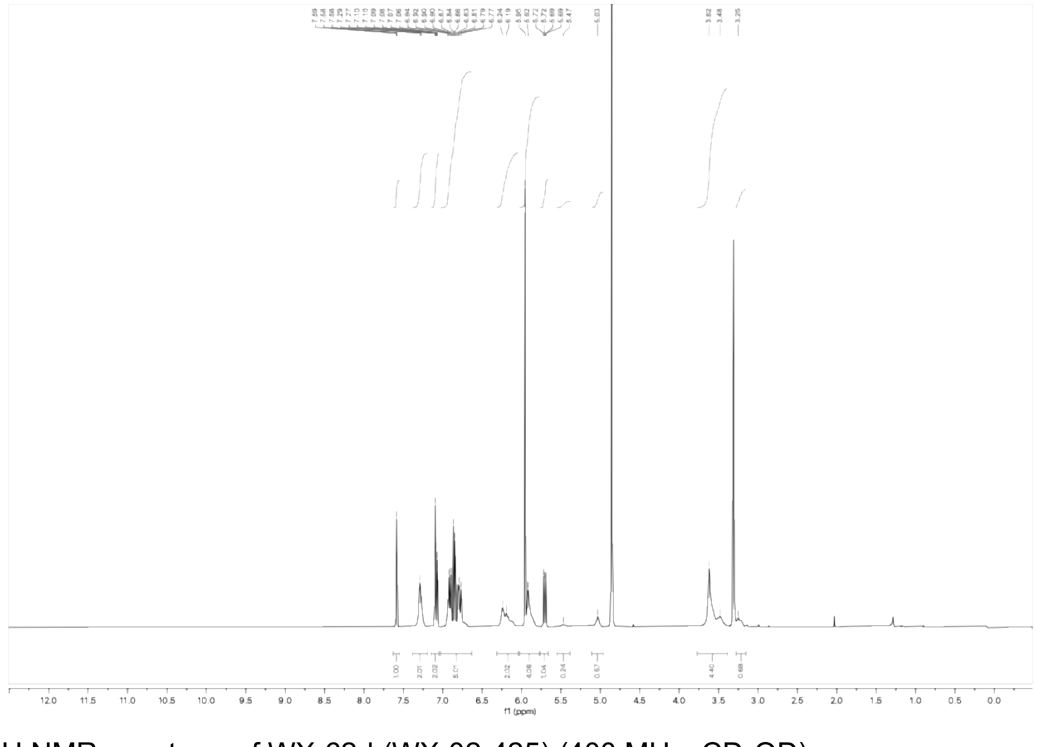

[1]H NMR spectrum of WX-62d (WX-02-425) (400 MHz, $CD_3OD$)

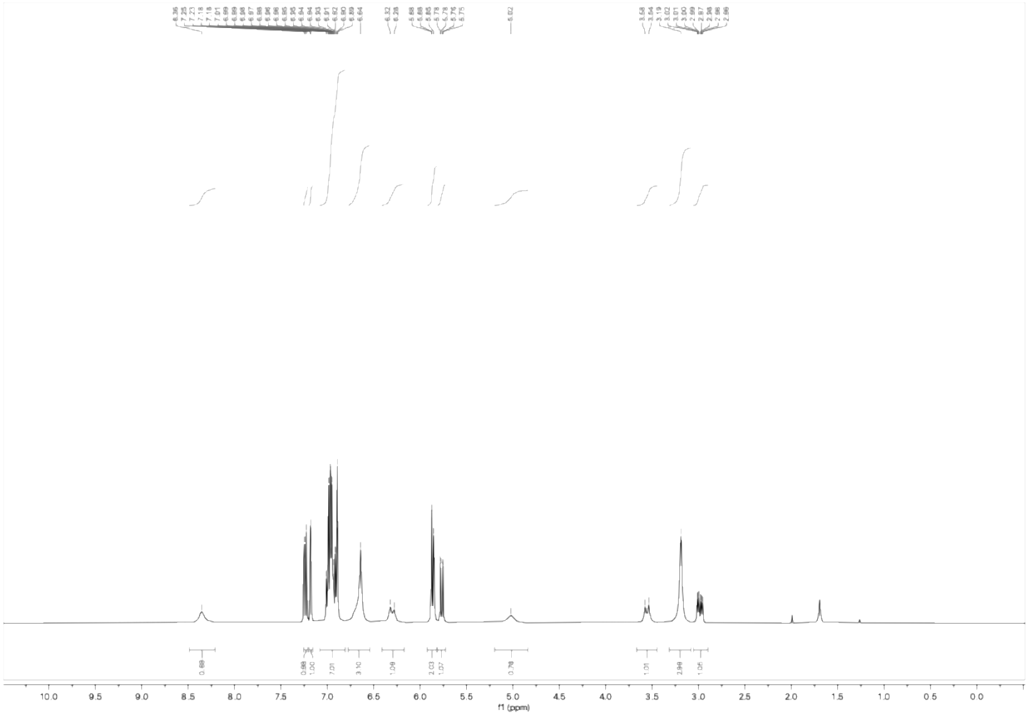

[1]H NMR spectrum of WX-63a (WX-02-426) (400 MHz, $CDCl_3$)

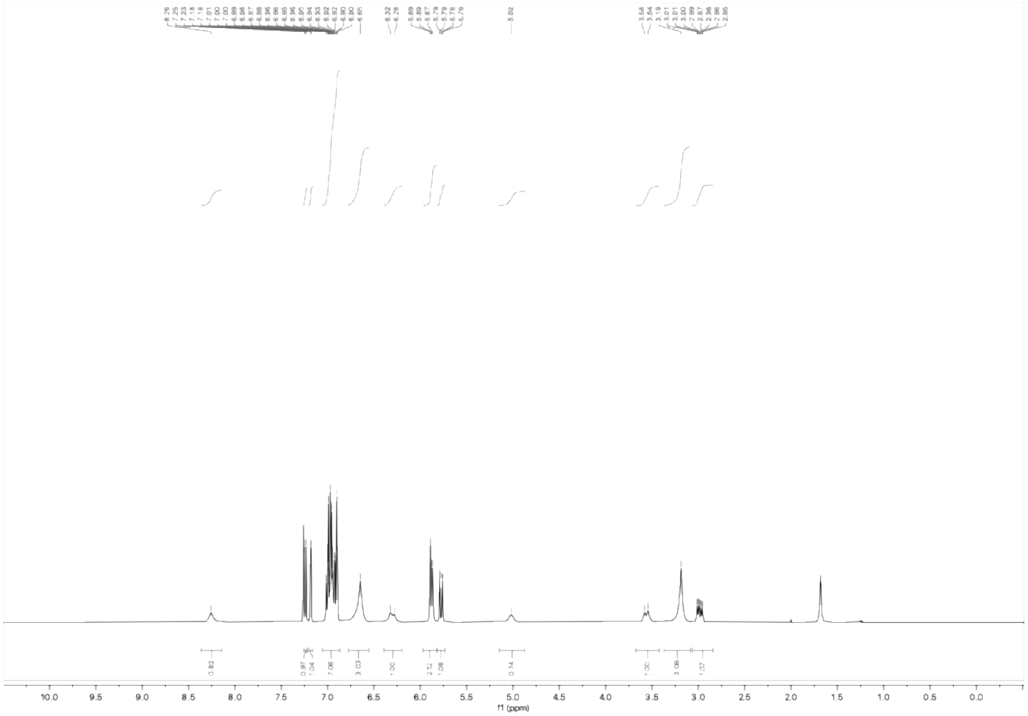

[1]H NMR spectrum of WX-63c (WX-02-427) (400 MHz, $CDCl_3$)

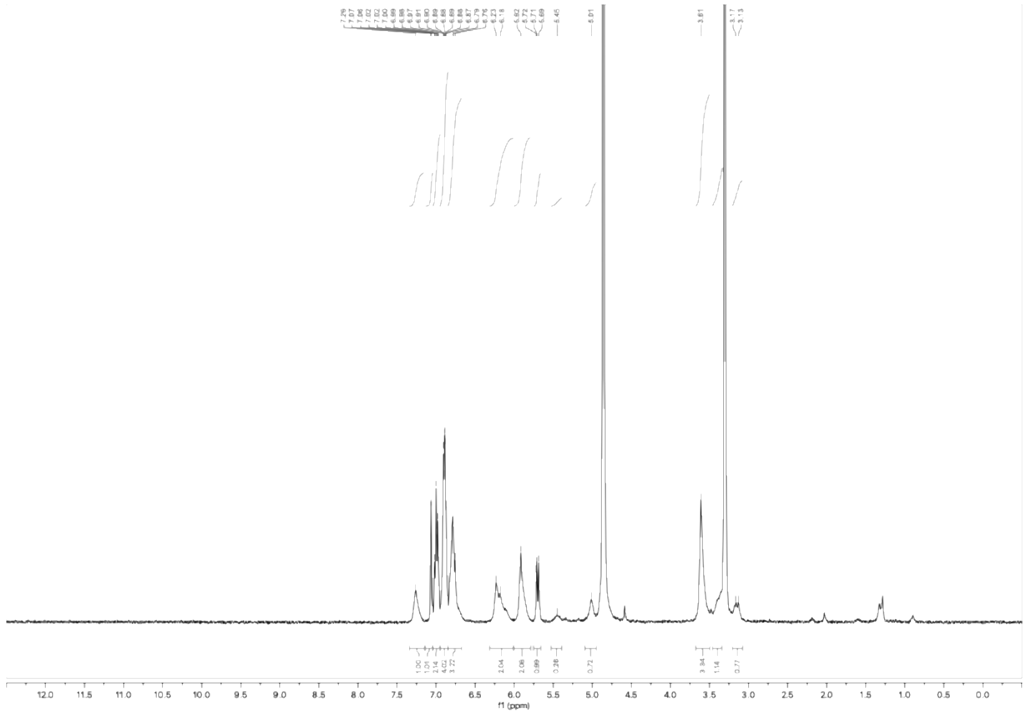

$^{1}H$ NMR spectrum of WX-63b (WX-02-428) (400 MHz, $CD_3OD$)

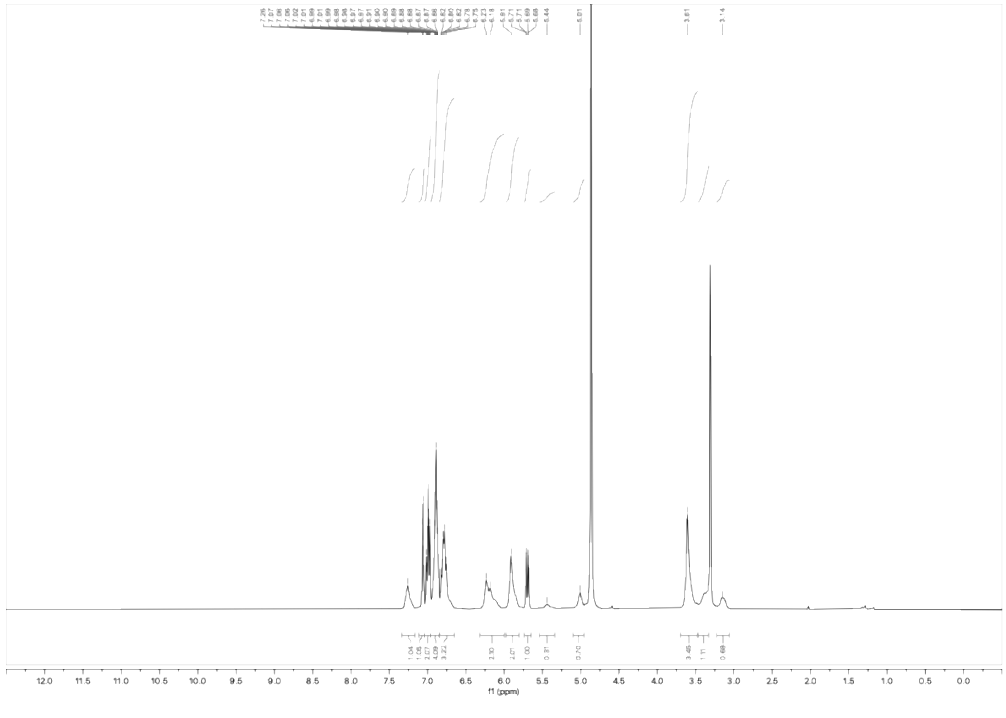

$^{1}H$ NMR spectrum of WX-63d (WX-02-429) (400 MHz, $CD_3OD$)

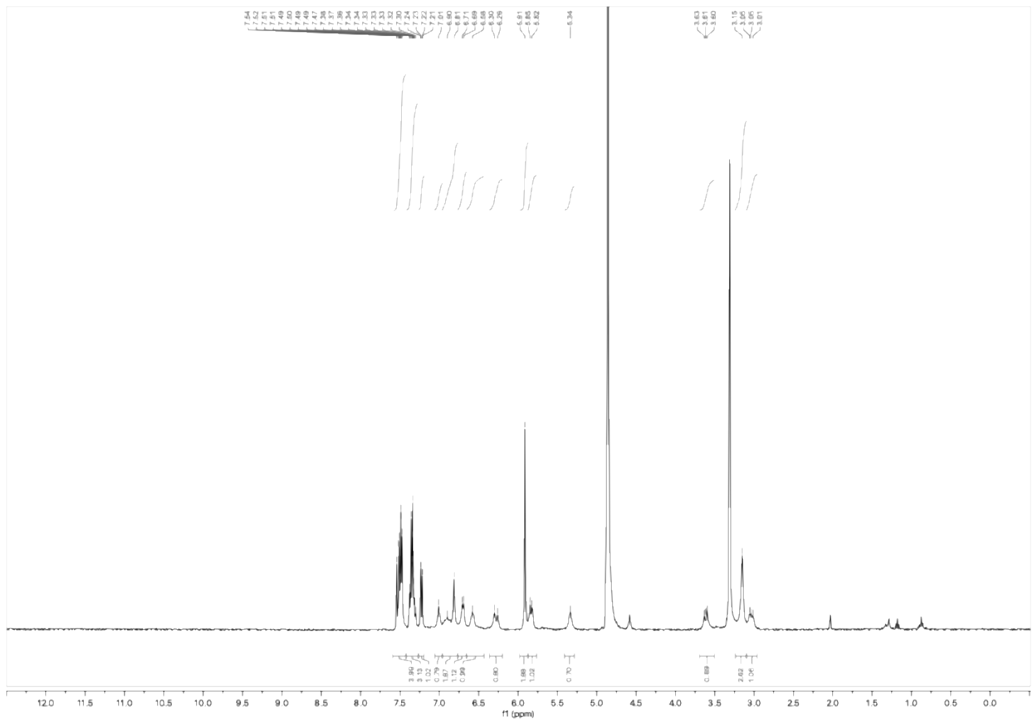

$^{1}H$ NMR spectrum of WX-71a (WX-02-430) (400 MHz, $CD_3OD$)

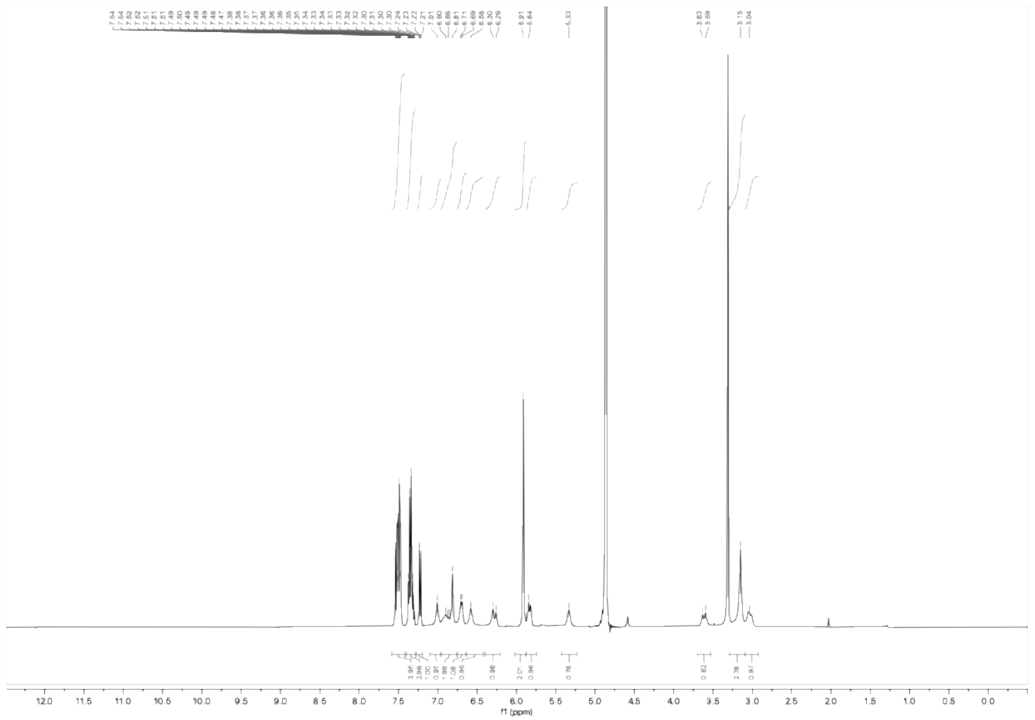

$^{1}H$ NMR spectrum of WX-71c (WX-02-431) (400 MHz, $CD_3OD$)

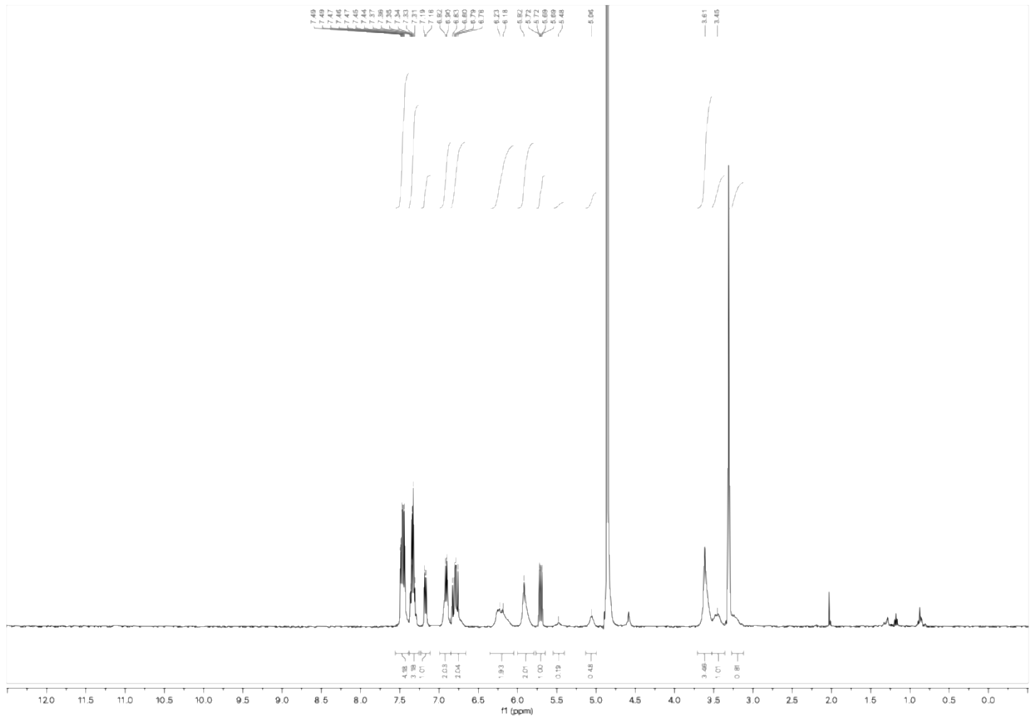

$^{1}H$ NMR spectrum of WX-71b (WX-02-432) (400 MHz, $CD_3OD$)

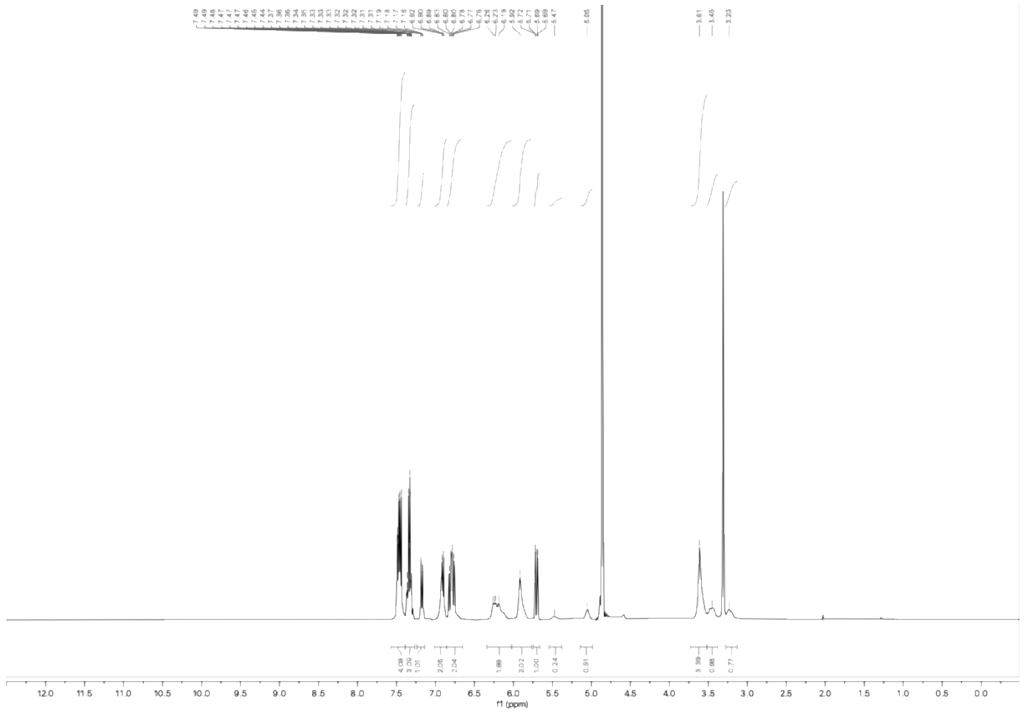

$^{1}H$ NMR spectrum of WX-71d (WX-02-433) (400 MHz, $CD_3OD$)

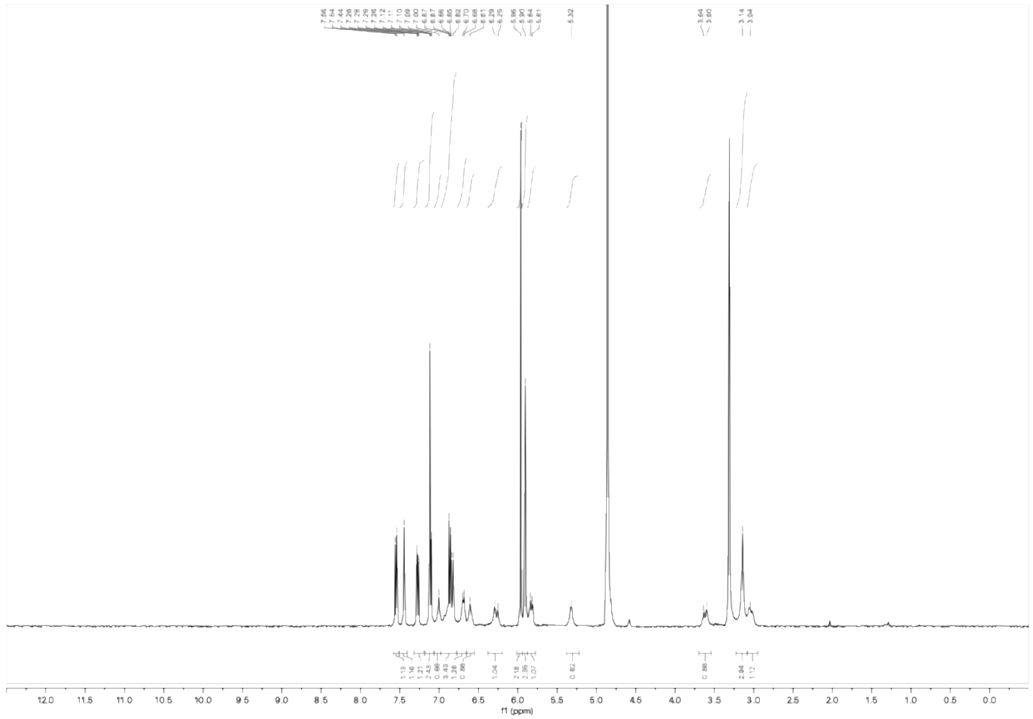

$^{1}H$ NMR spectrum of WX-72a (WX-02-434) (400 MHz, $CD_3OD$)

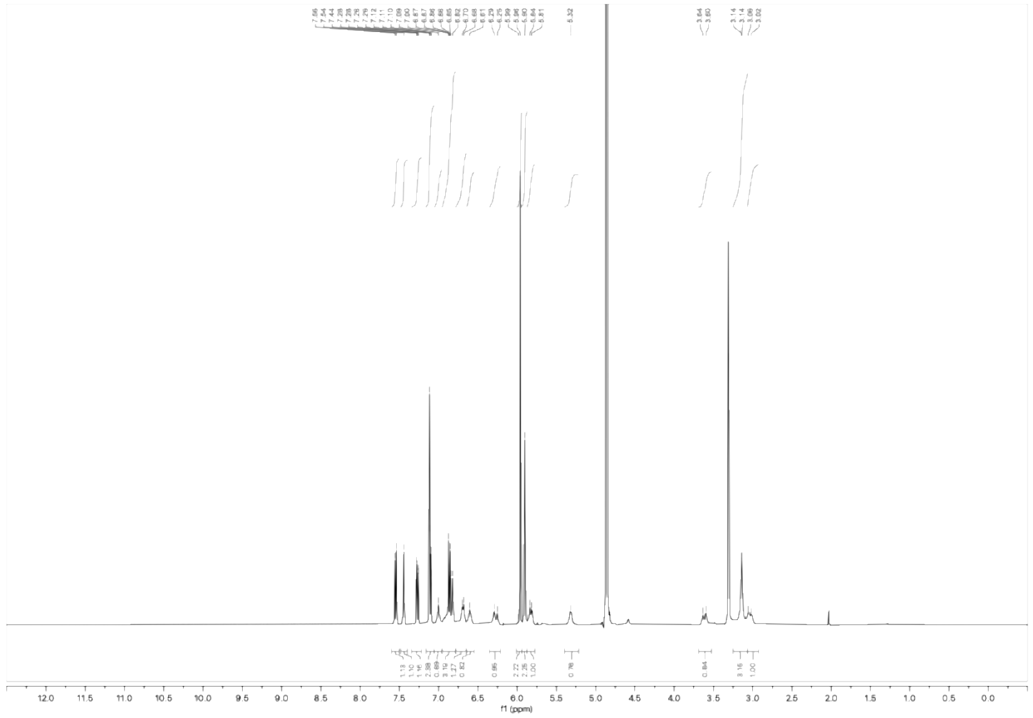

$^{1}H$ NMR spectrum of WX-72c (WX-02-435) (400 MHz, $CD_3OD$)

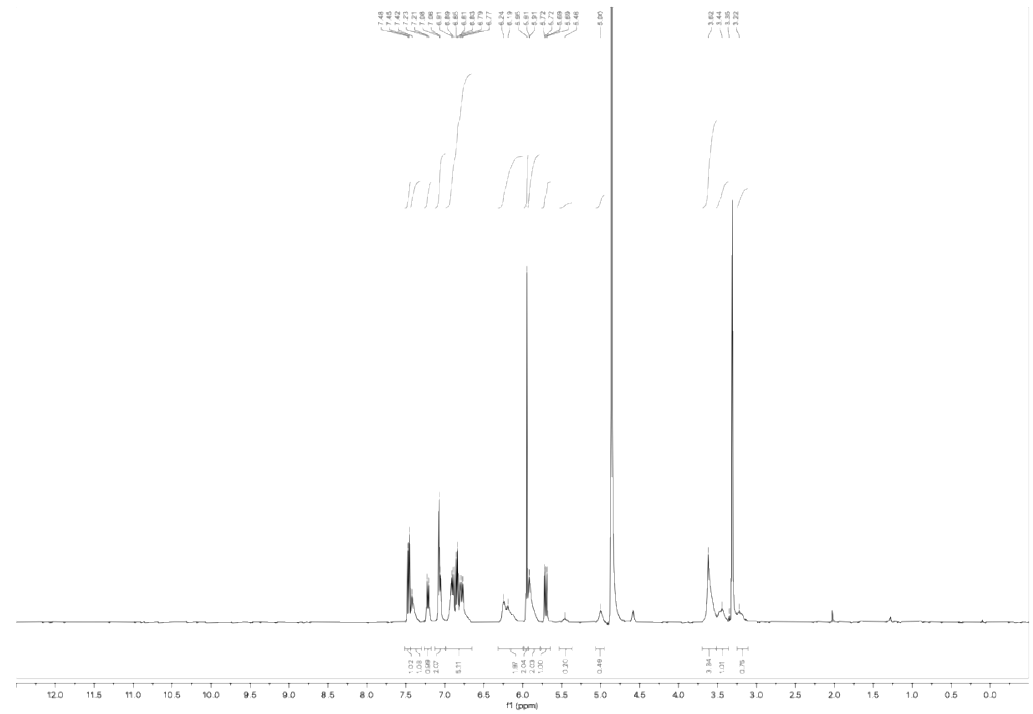


$^1$H NMR spectrum of WX-72b (WX-02-436) (400 MHz, $CD_3OD$)

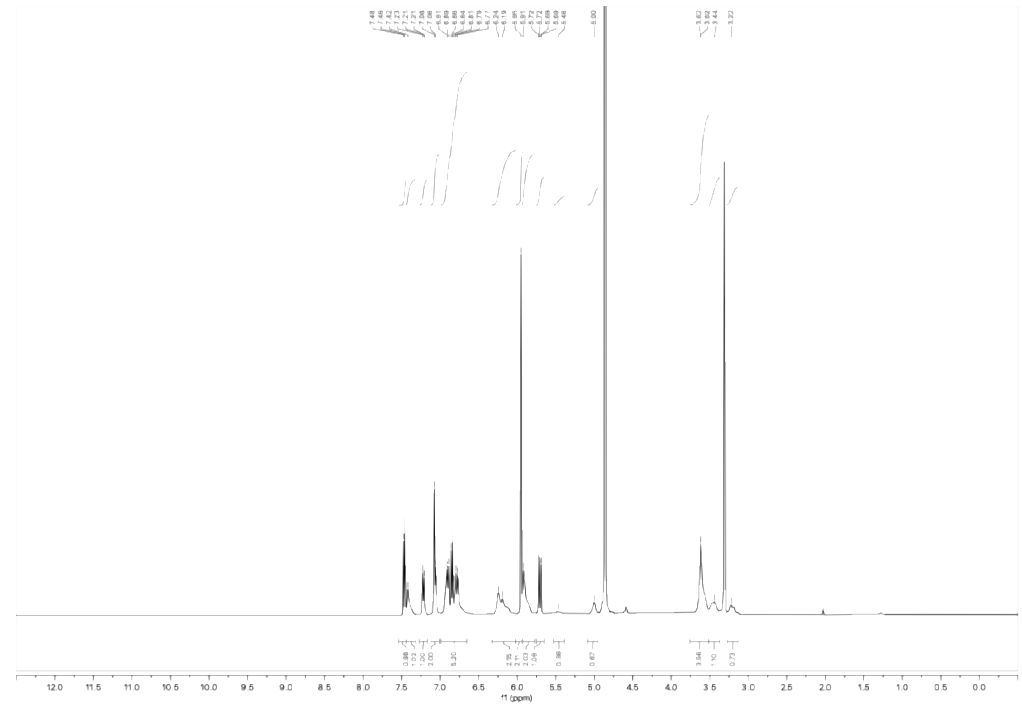


$^1$H NMR spectrum of WX-72d (WX-02-437) (400 MHz, $CD_3OD$)

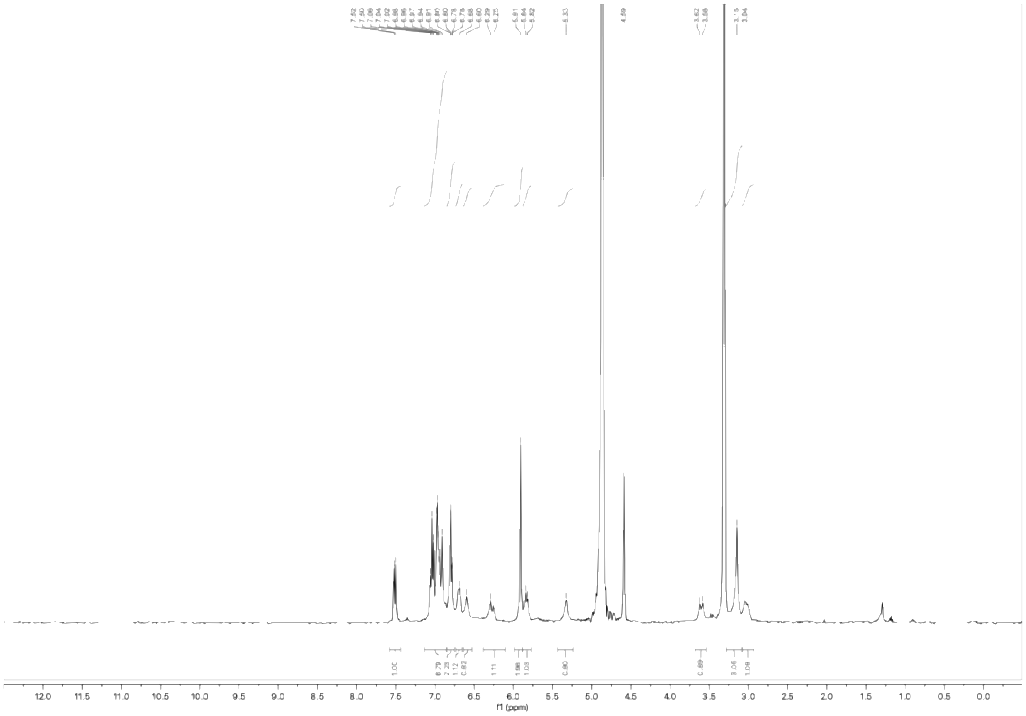


$^1H$ NMR spectrum of WX-73a (WX-02-438) (400 MHz, $CD_3OD$)

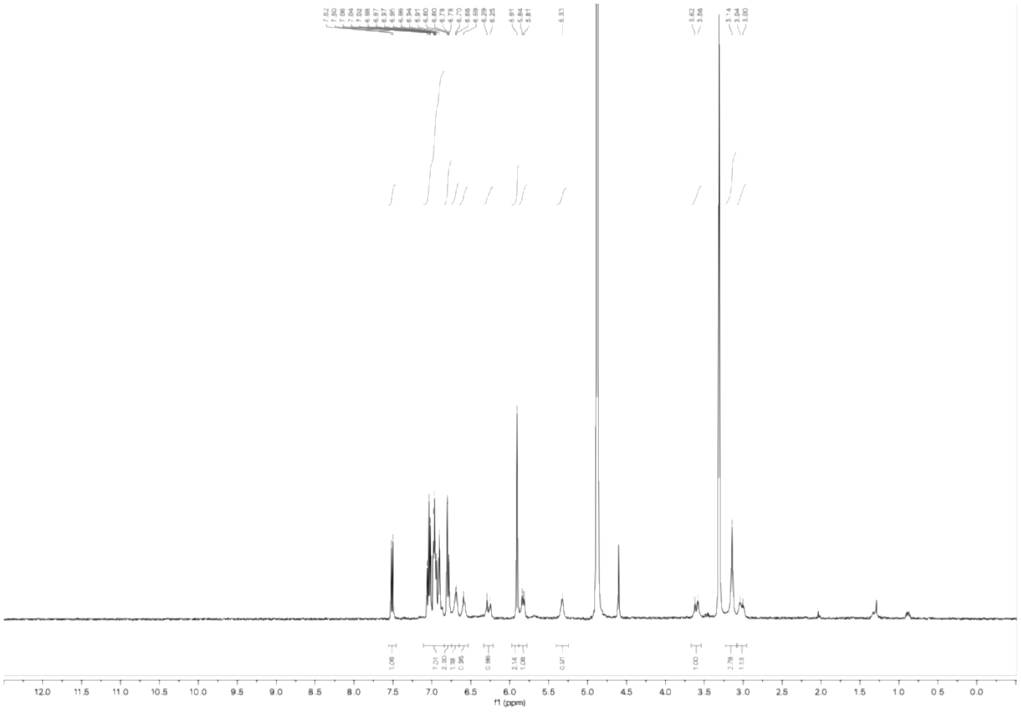


$^1H$ NMR spectrum of WX-73c (WX-02-439) (400 MHz, $CD_3OD$)

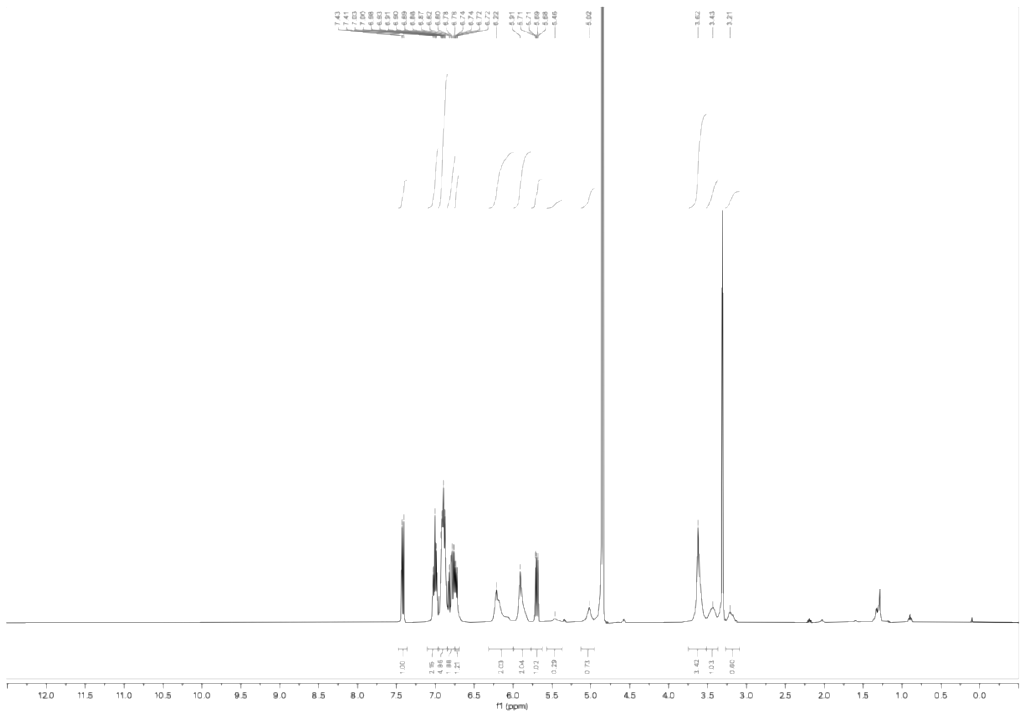

[1]H NMR spectrum of WX-73b (WX-02-440) (400 MHz, $CD_3OD$)

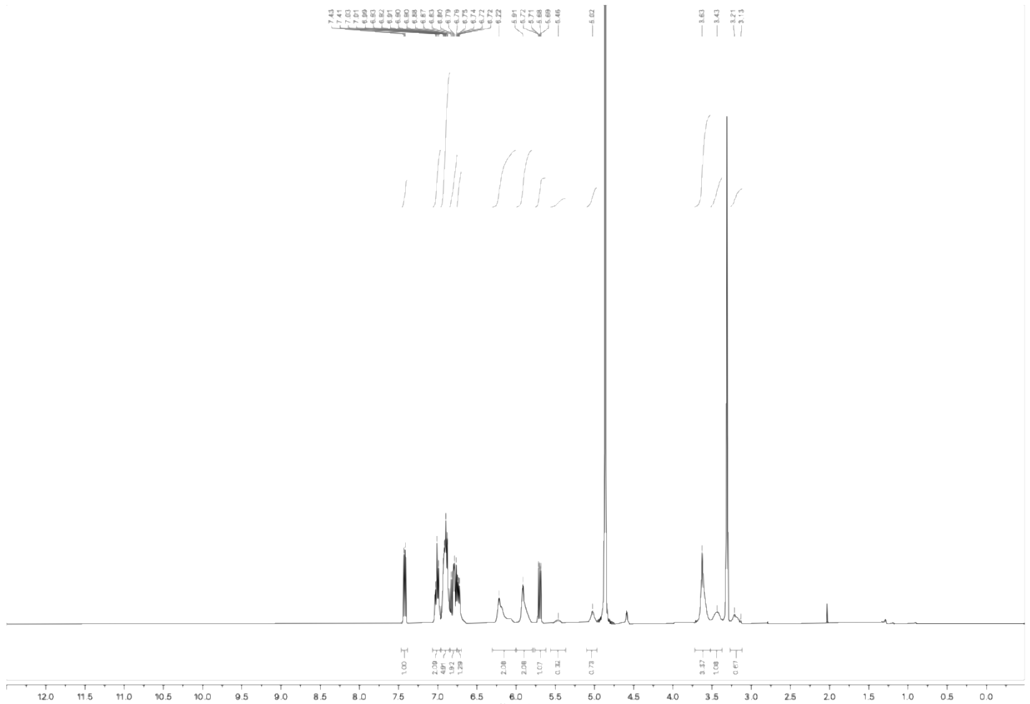

[1]H NMR spectrum of WX-73d (WX-02-441) (400 MHz, $CD_3OD$)

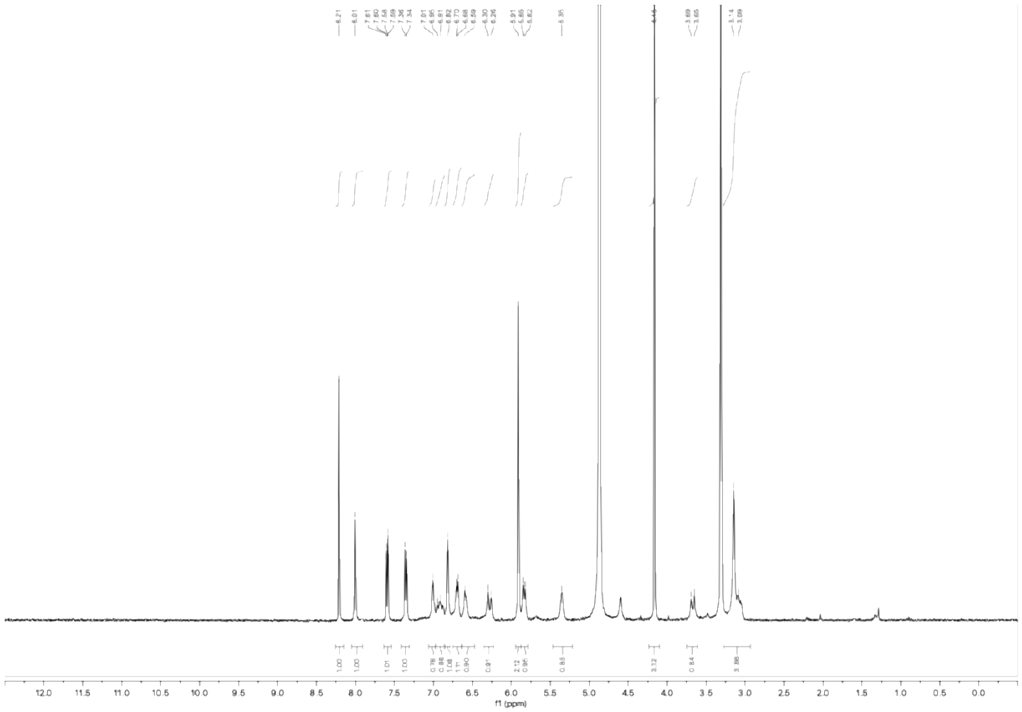

$^{1}H$ NMR spectrum of WX-64a (WX-02-687) (400 MHz, $CD_3OD$)

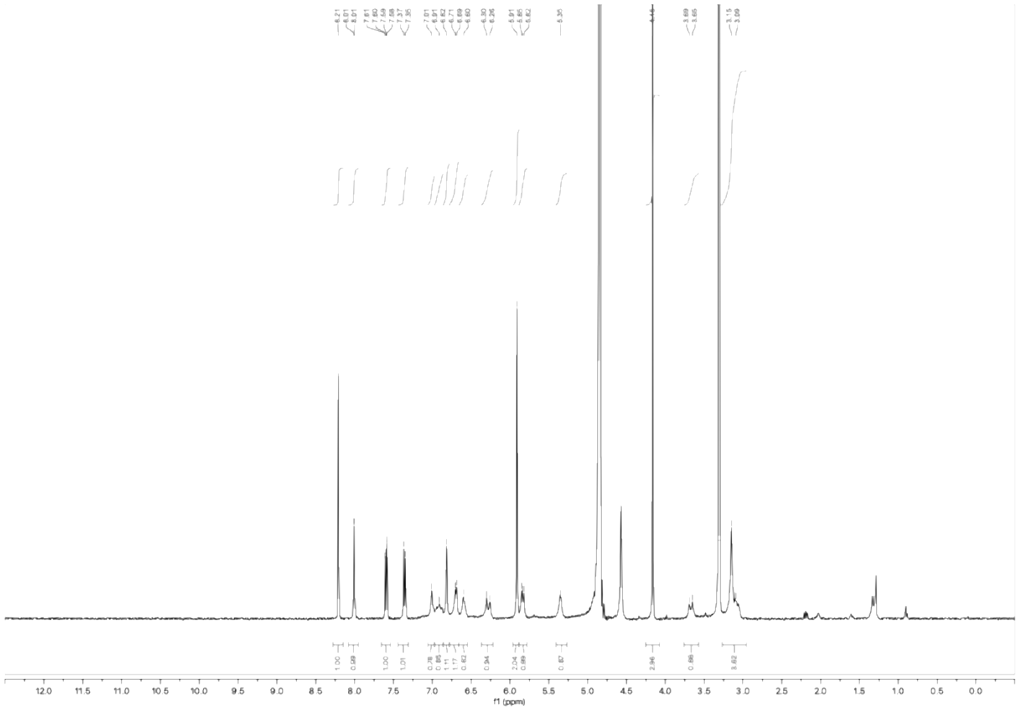

$^{1}H$ NMR spectrum of WX-64c (WX-02-688) (400 MHz, $CD_3OD$)

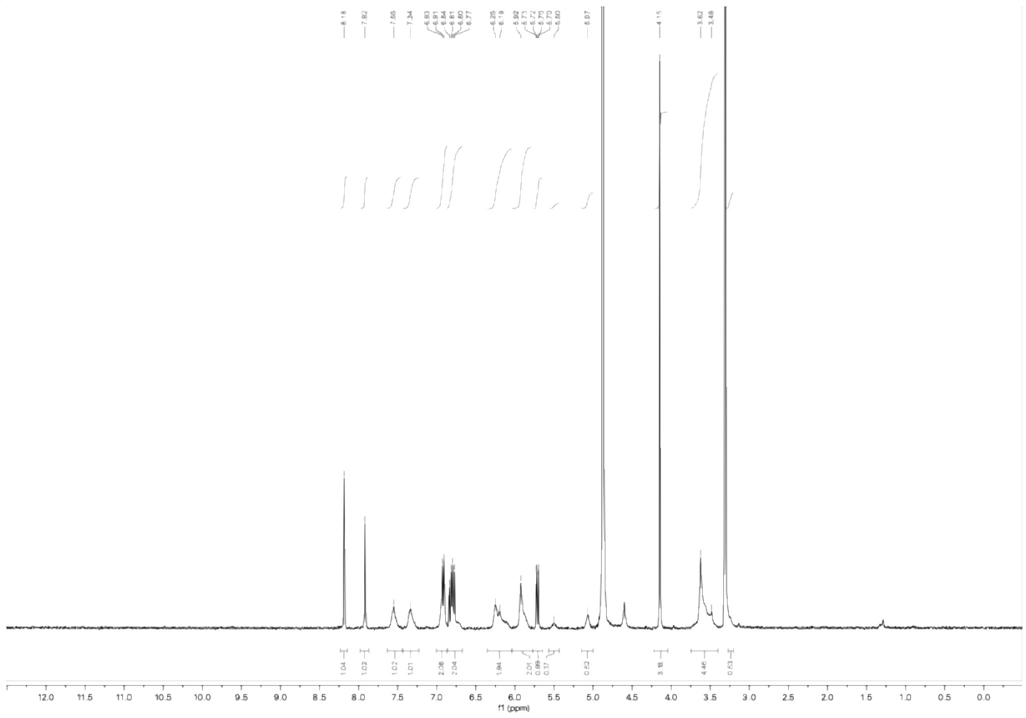

$^1$H NMR spectrum of WX-64b (WX-02-689) (400 MHz, $CD_3OD$)

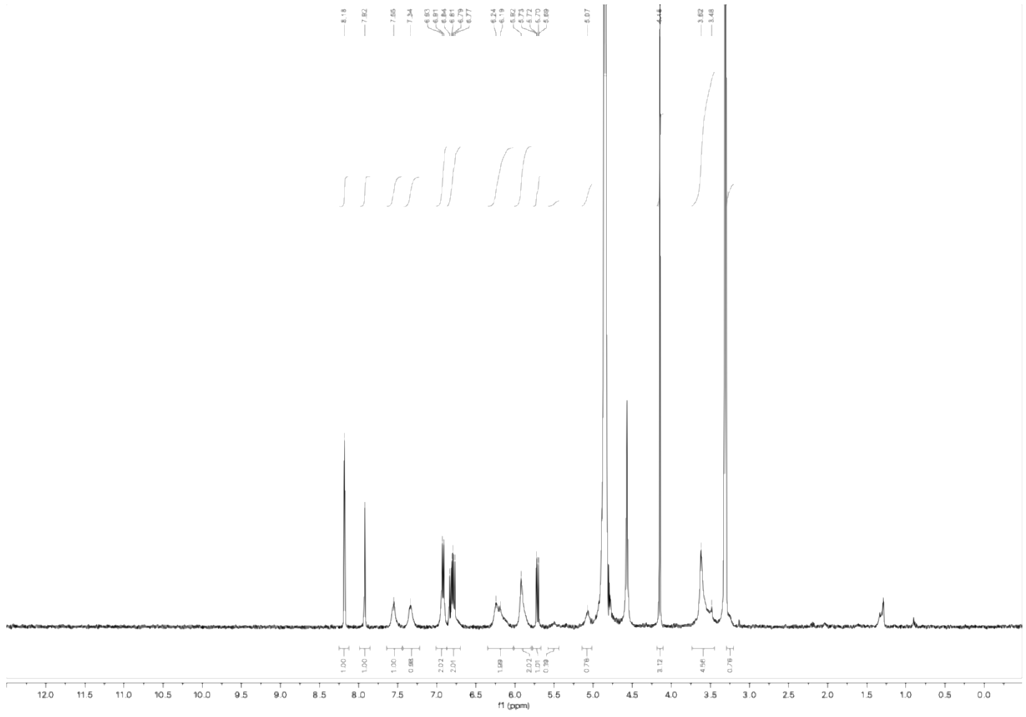

$^1$H NMR spectrum of WX-64d (WX-02-690) (400 MHz, $CD_3OD$)

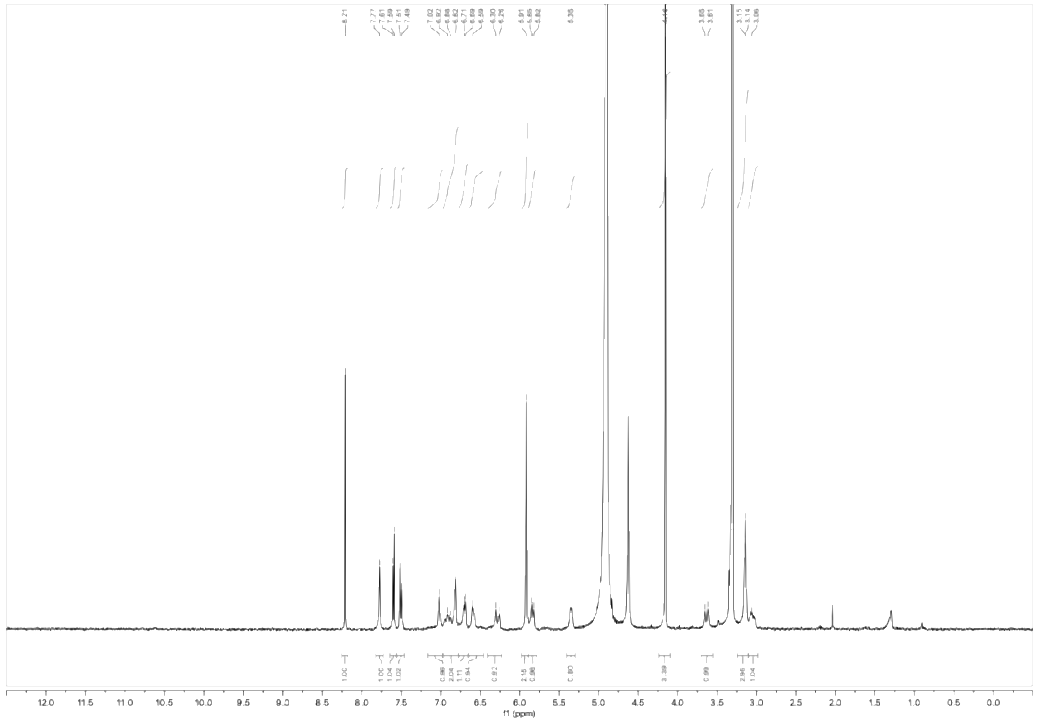

$^1$H NMR spectrum of WX-74a (WX-02-691) (400 MHz, $CD_3OD$)

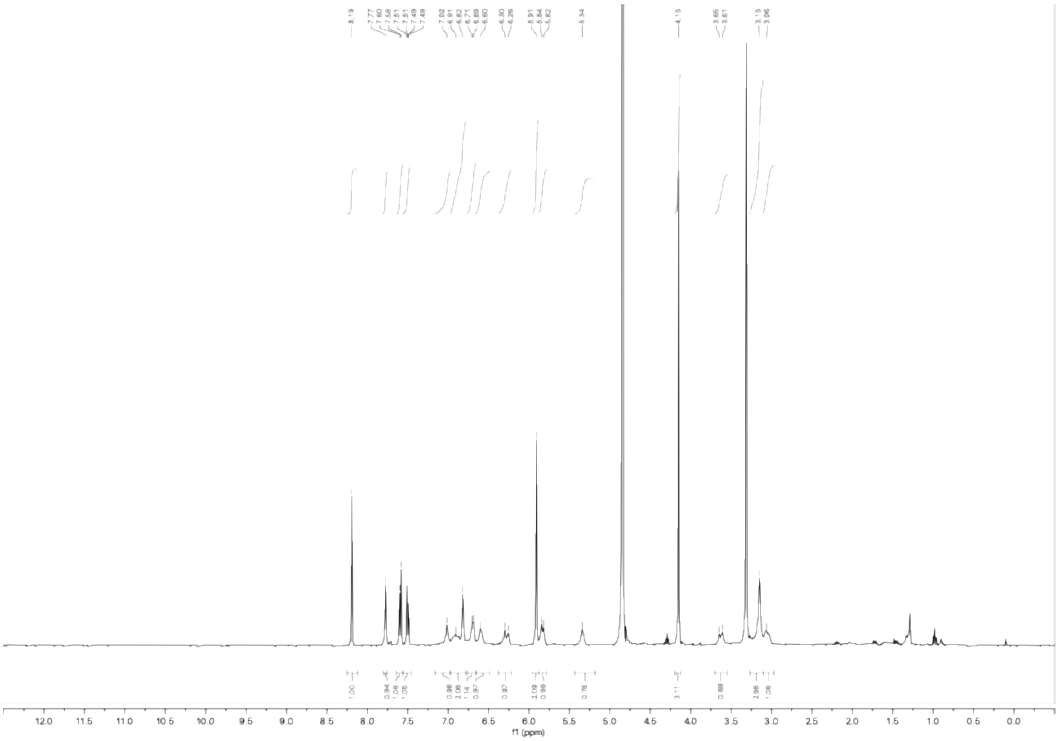

$^1$H NMR spectrum of WX-74c (WX-02-692) (400 MHz, $CD_3OD$)

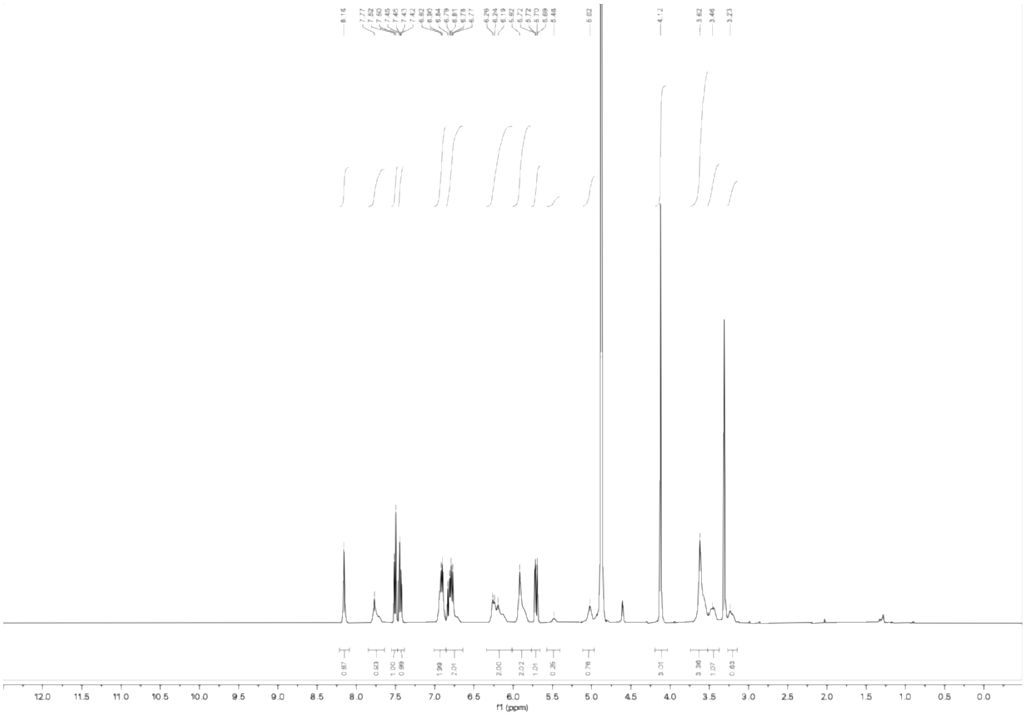

$^{1}H$ NMR spectrum of WX-74b (WX-02-693) (400 MHz, $CD_3OD$)

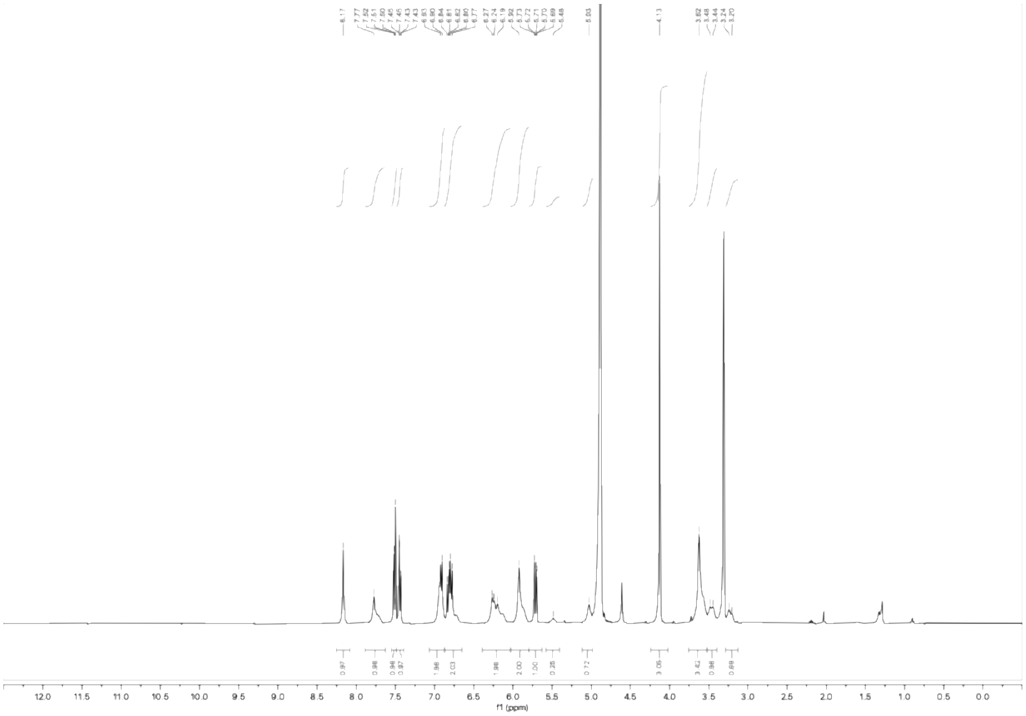

$^{1}H$ NMR spectrum of WX-74d (WX-02-694) (400 MHz, $CD_3OD$)

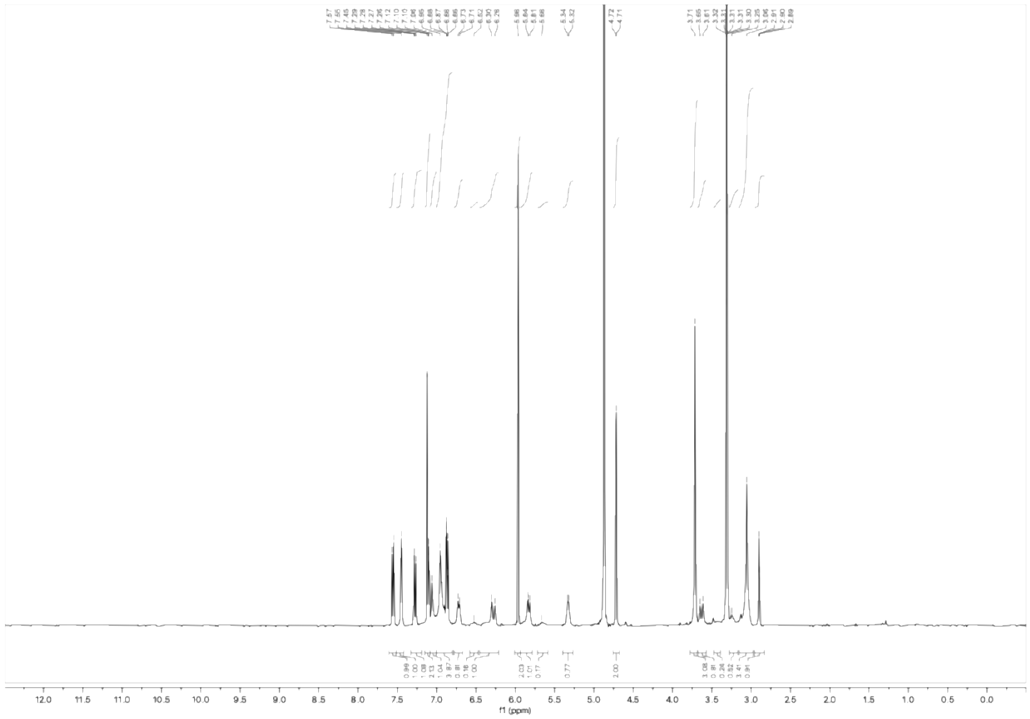

$^1$H NMR spectrum of WX-72a-yne (WX-02-891) (400 MHz, $CD_3OD$)

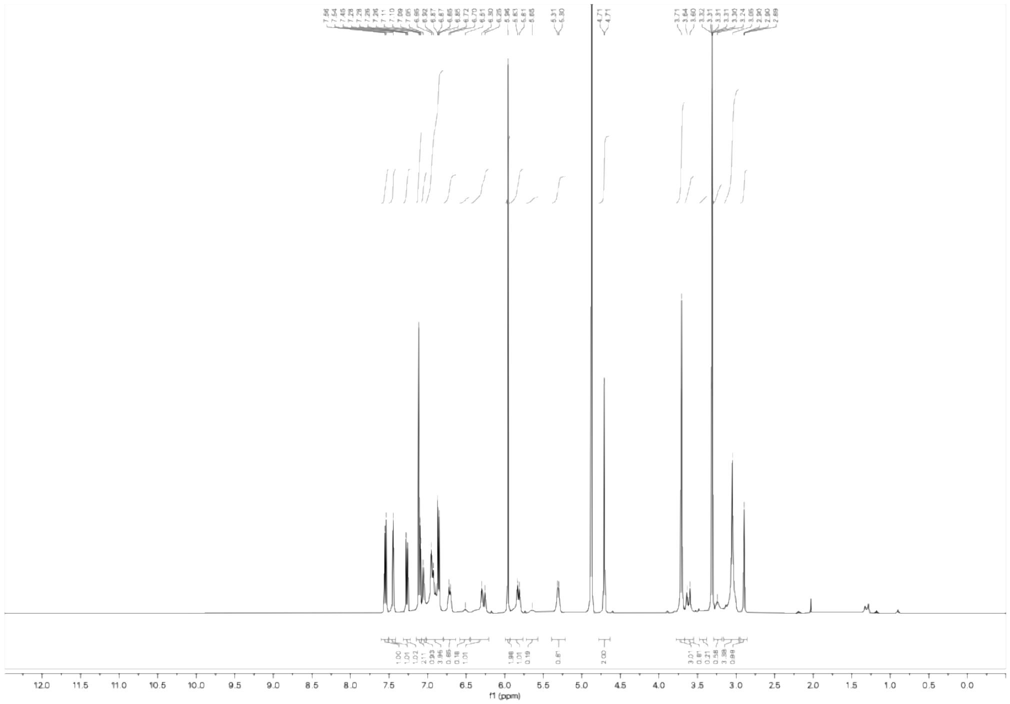

$^1$H NMR spectrum of WX-72c-yne (WX-02-892) (400 MHz, $CD_3OD$)

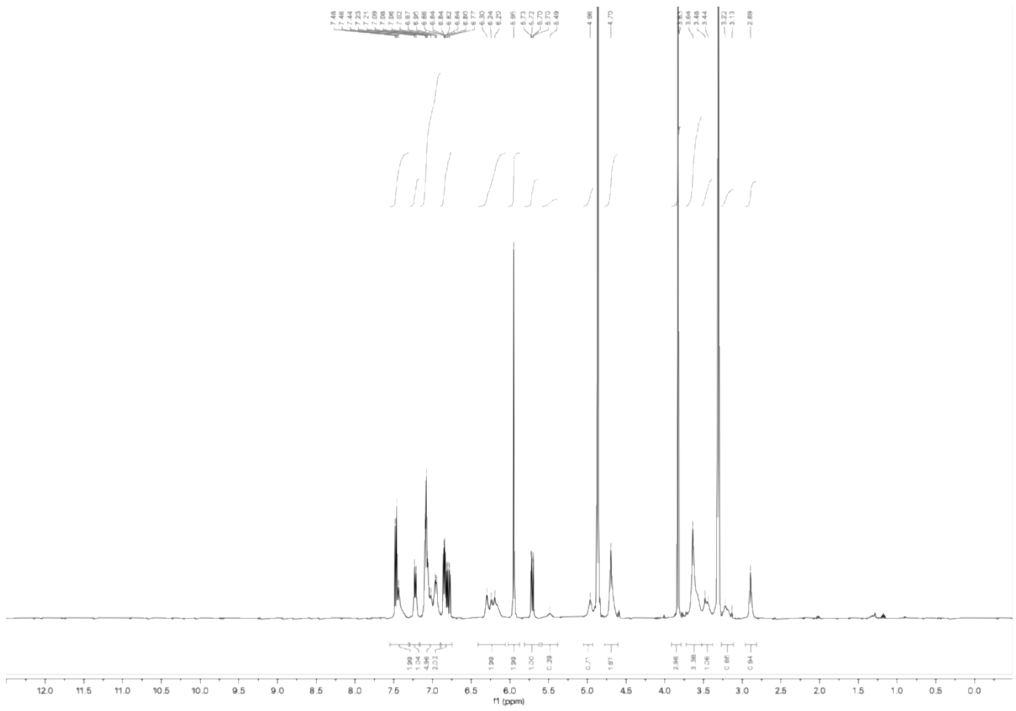

$^{1}H$ NMR spectrum of WX-72b-yne (WX-02-893) (400 MHz, $CD_3OD$)

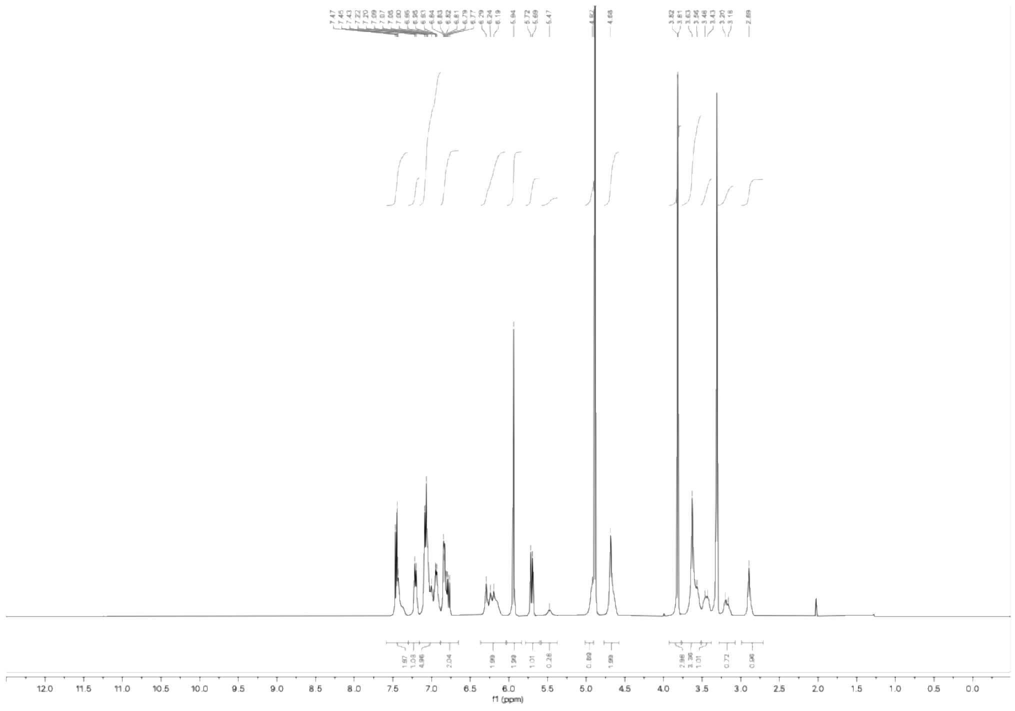

$^{1}H$ NMR spectrum of WX-72d-yne (WX-02-894) (400 MHz, $CD_3OD$)

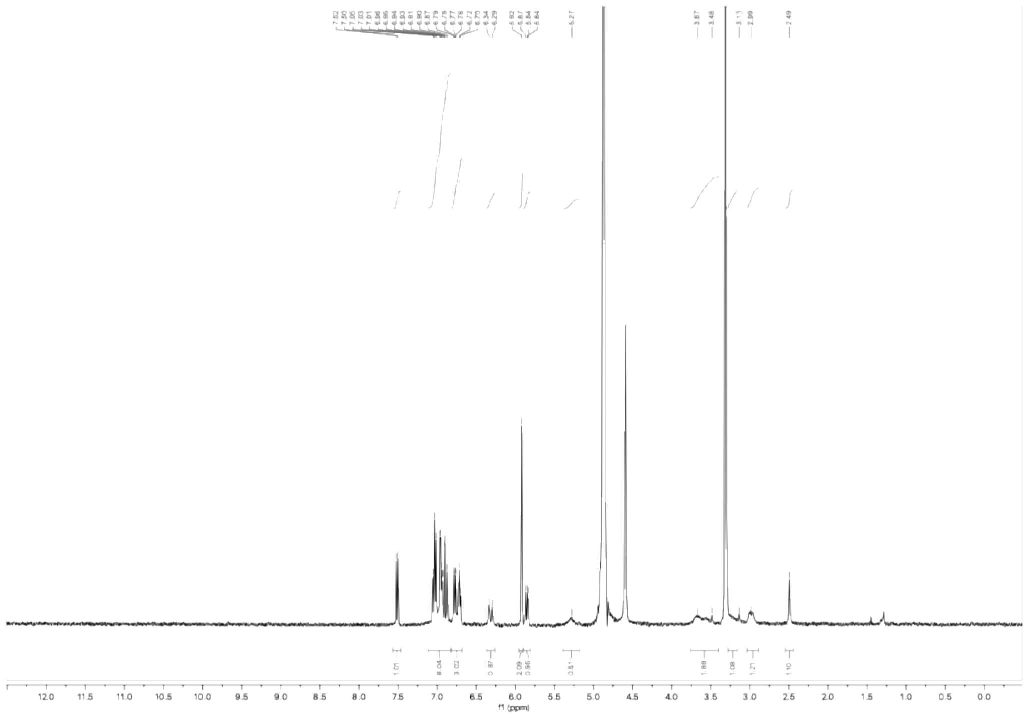

[1]H NMR spectrum of WX-73a-yne (WX-02-905) (400 MHz, $CD_3OD$)

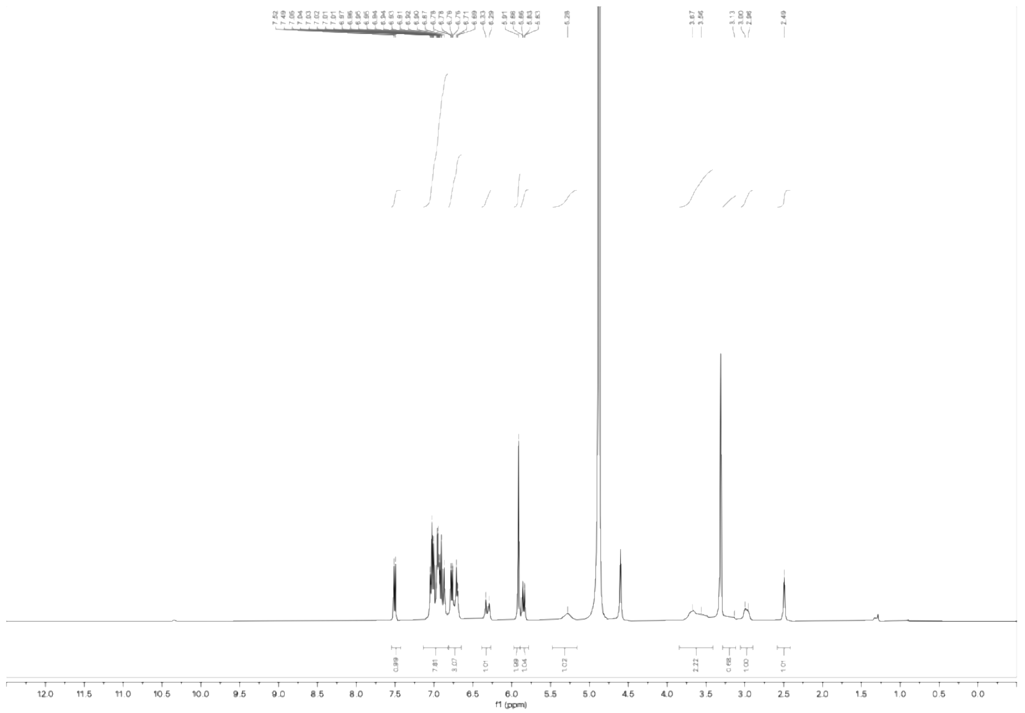

[1]H NMR spectrum of WX-73c-yne (WX-02-906) (400 MHz, $CD_3OD$)

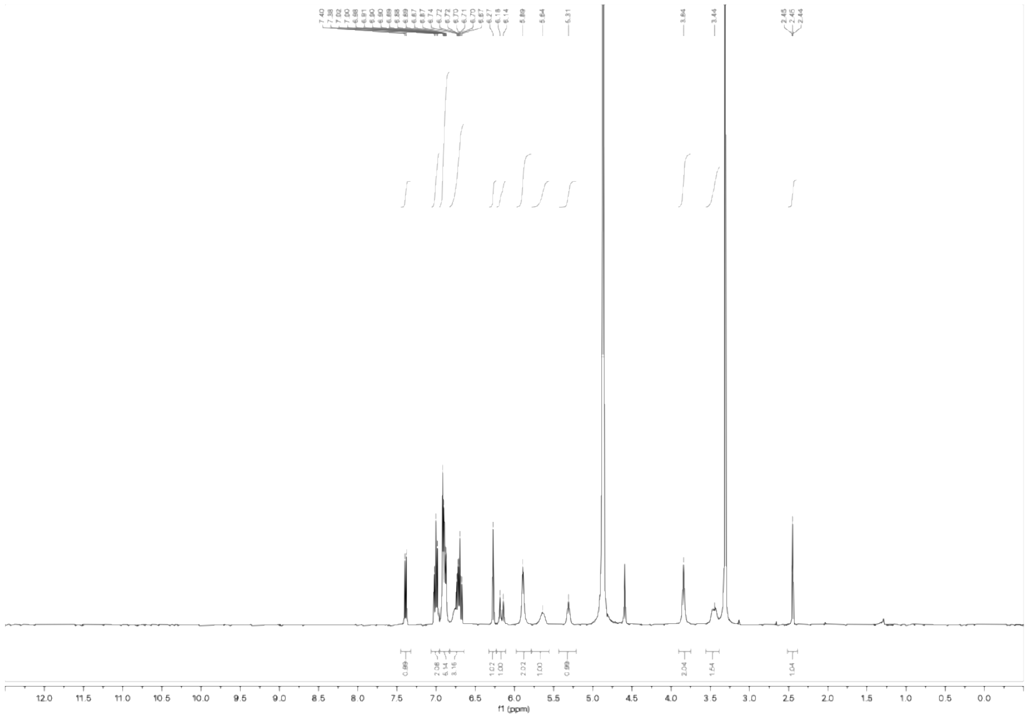

[1]H NMR spectrum of WX-73b-yne (WX-02-907) (400 MHz, $CD_3OD$)

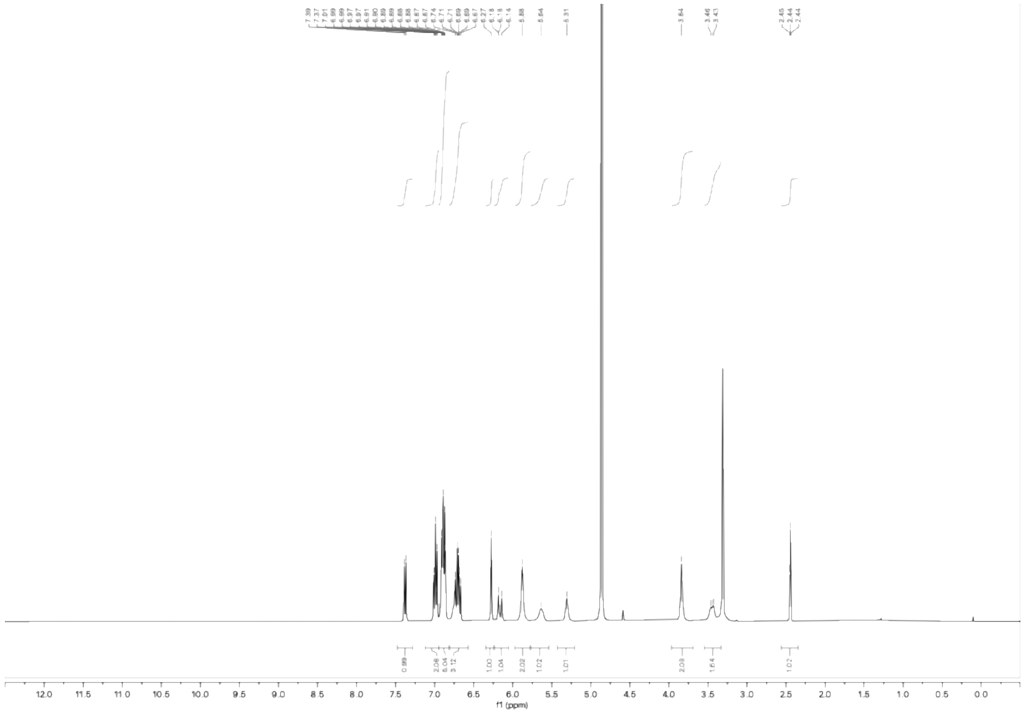

[1]H NMR spectrum of WX-73d-yne (WX-02-908) (400 MHz, $CD_3OD$)

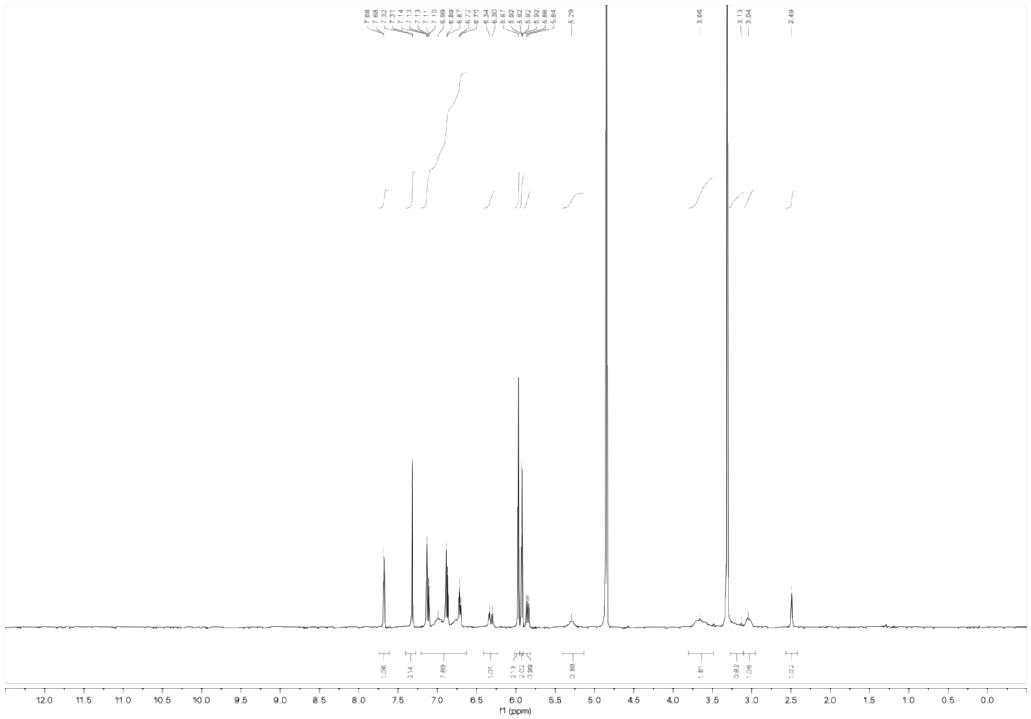

[1]H NMR spectrum of WX-62a-yne (WX-02-909) (400 MHz, $CD_3OD$)

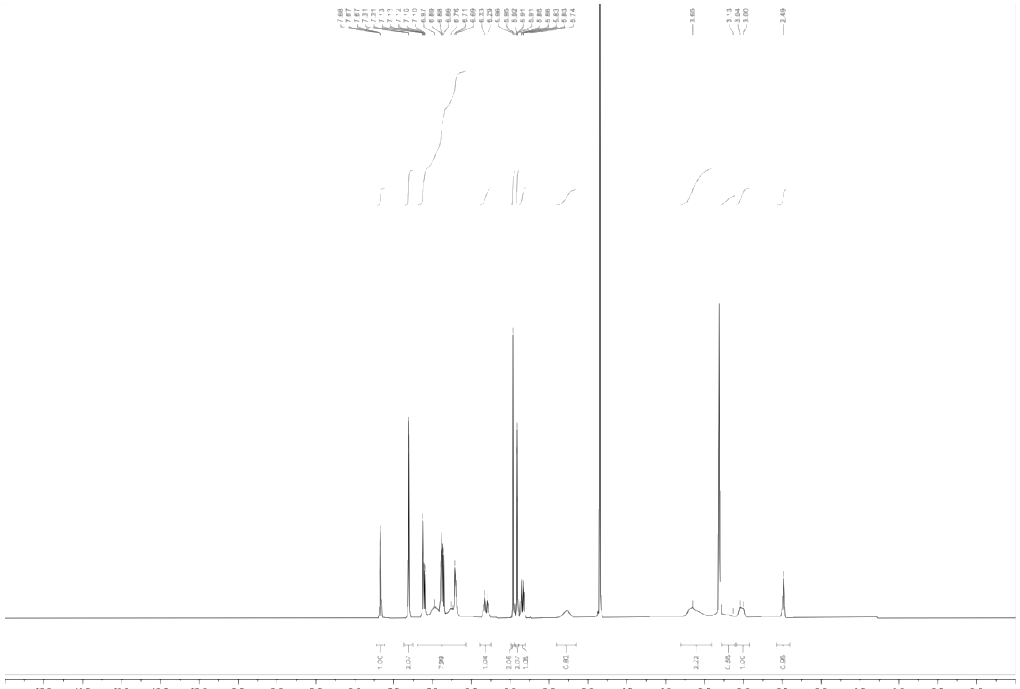

[1]H NMR spectrum of WX-62c-yne (WX-02-910) (400 MHz, $CD_3OD$)

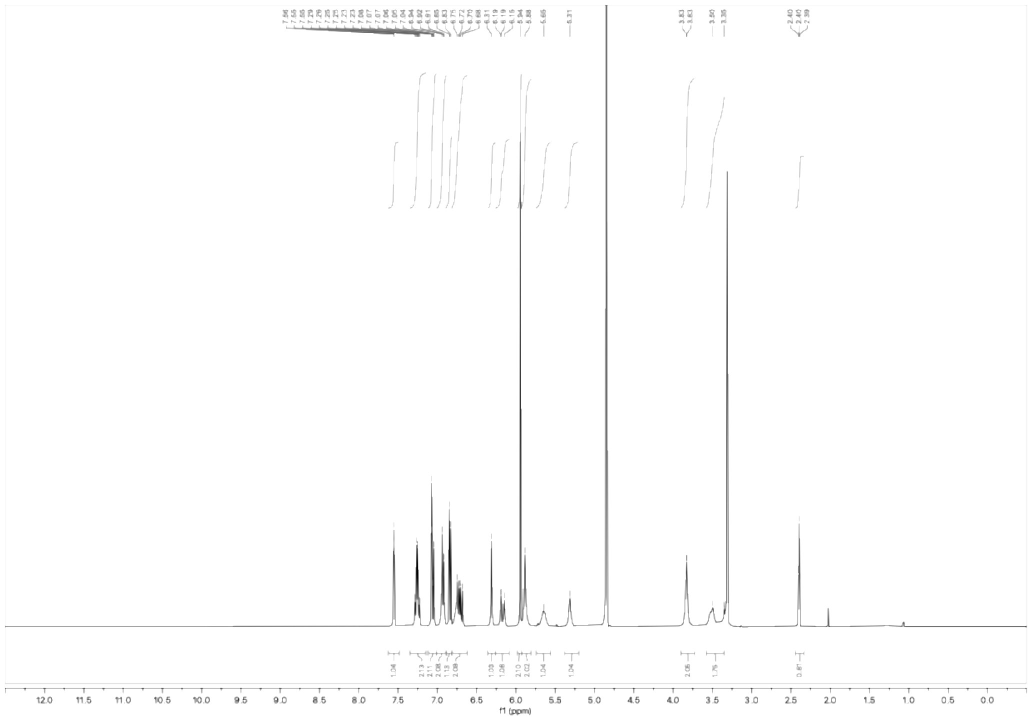

[1]H NMR spectrum of WX-62b-yne (WX-02-911) (400 MHz, $CD_3OD$)

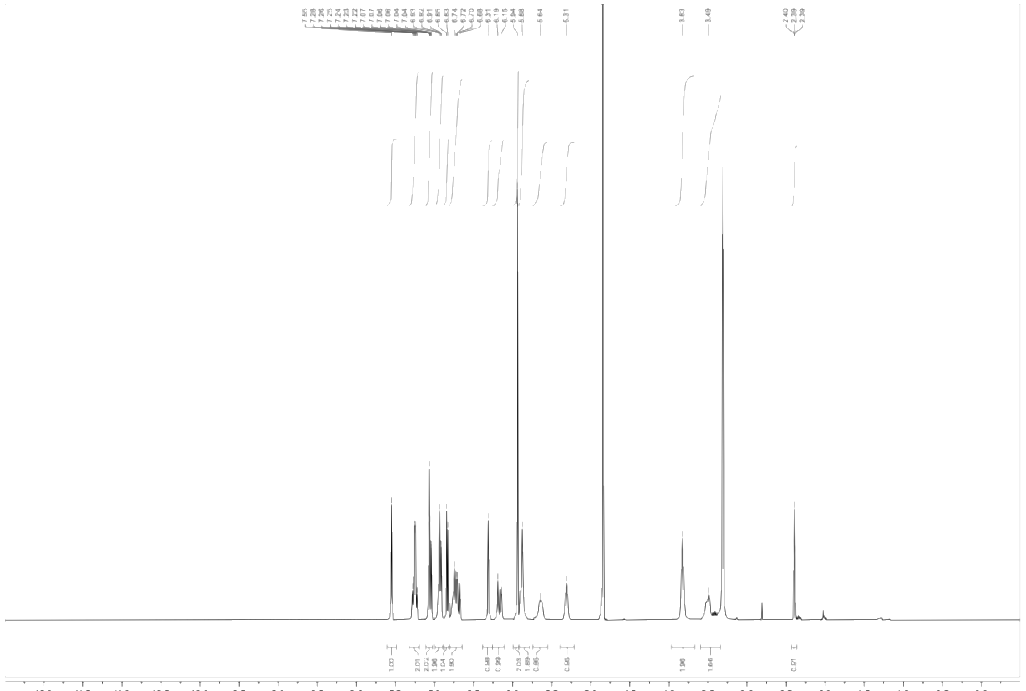

[1]H NMR spectrum of WX-62d-yne (WX-02-912) (400 MHz, $CD_3OD$)

**Analytical data: SFC**

WX-61a (WX-02-418)

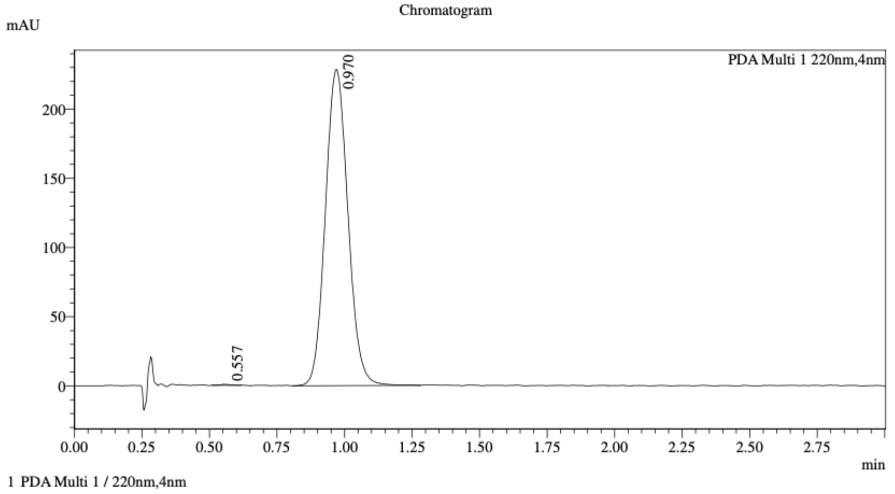


Integration Result

Peak Table

PDA Ch1 220nm

| Peak# | Ret. Time | Height | Height% | Resolution(USP) | Area | Area% |
|---|---|---|---|---|---|---|
| 1 | 0.557 | 625 | 0.275 | -- | 1693 | 0.127 |
| 2 | 0.970 | 226731 | 99.725 | 3.717 | 1329673 | 99.873 |

WX-61c (WX-02-420)

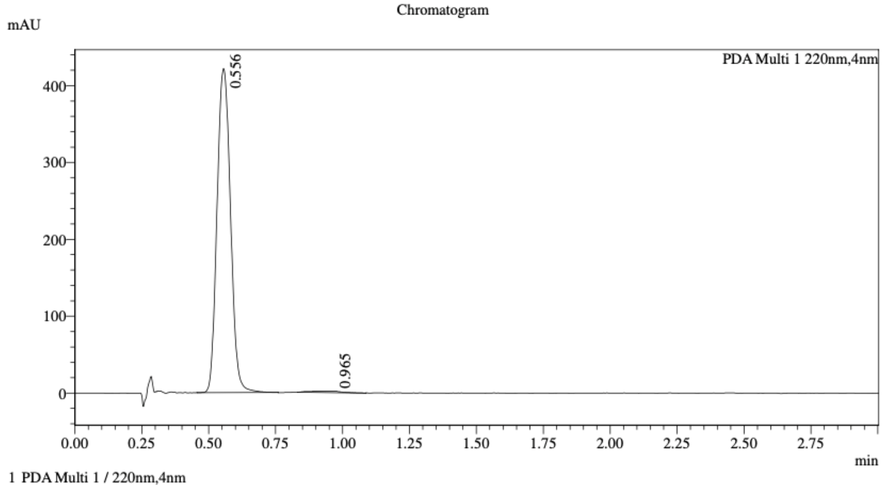


Integration Result

Peak Table

PDA Ch1 220nm

| Peak# | Ret. Time | Height | Height% | Resolution(USP) | Area | Area% |
|---|---|---|---|---|---|---|
| 1 | 0.556 | 419093 | 99.585 | -- | 1478569 | 99.120 |
| 2 | 0.965 | 1746 | 0.415 | 2.903 | 13133 | 0.880 |

Mixture of WX-61a (WX-02-418) & WX-61c (WX-02-420)

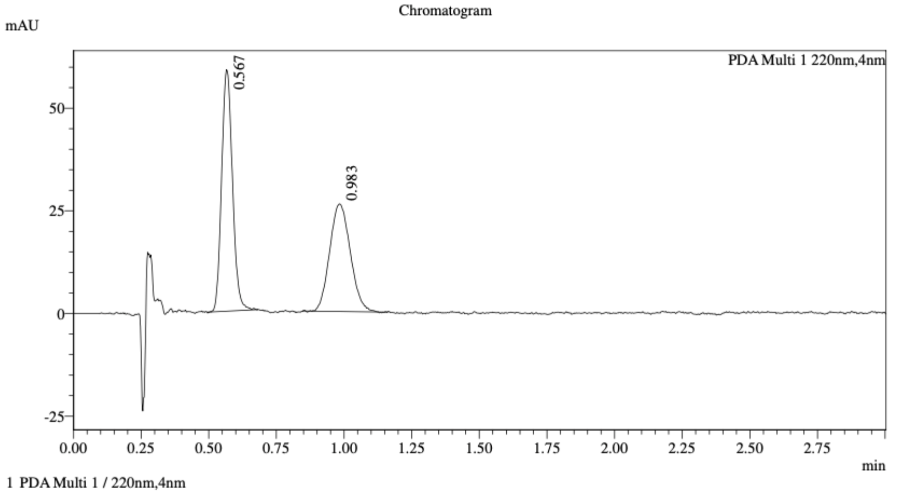


Integration Result

Peak Table

PDA Ch1 220nm

| Peak# | Ret. Time | Height | Height% | Resolution(USP) | Area | Area% |
|---|---|---|---|---|---|---|
| 1 | 0.567 | 58285 | 69.067 | -- | 164221 | 54.221 |
| 2 | 0.983 | 26104 | 30.933 | 3.804 | 138652 | 45.779 |

Method details

Column: Cellulose-4 50×4.6mm I.D., 3 μm
Mobile phase: A: $CO_2$, B: EtOH (0.05% diethylamine)
Elution: 50% B

WX-61b (WX-02-419)

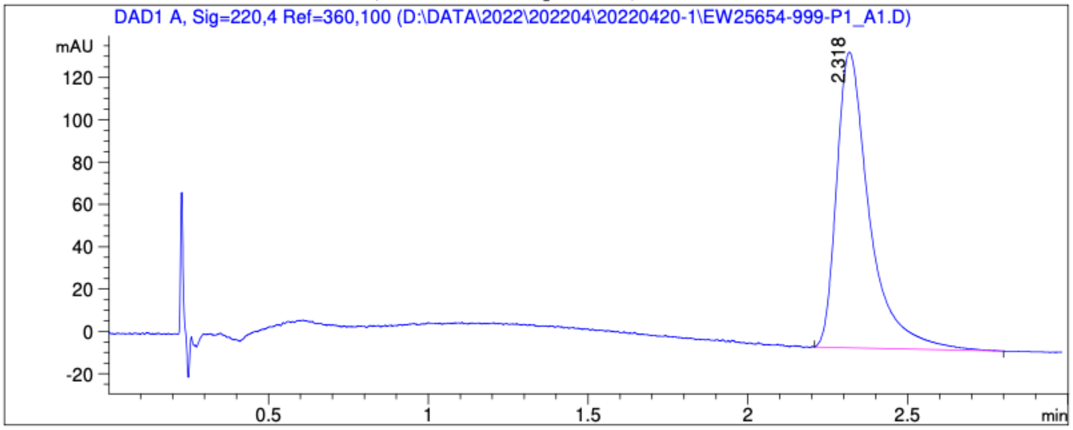


Signal: DAD1 A, Sig=220,4 Ref=360,100

| RetTime [min] | Height | Resolution | Symm. | Area [mAU*s] | Area % |
|---|---|---|---|---|---|
| 2.318 | 139.829 | | 0.671 | 998.515 | 100.000 |

WX-61d (WX-02-421)

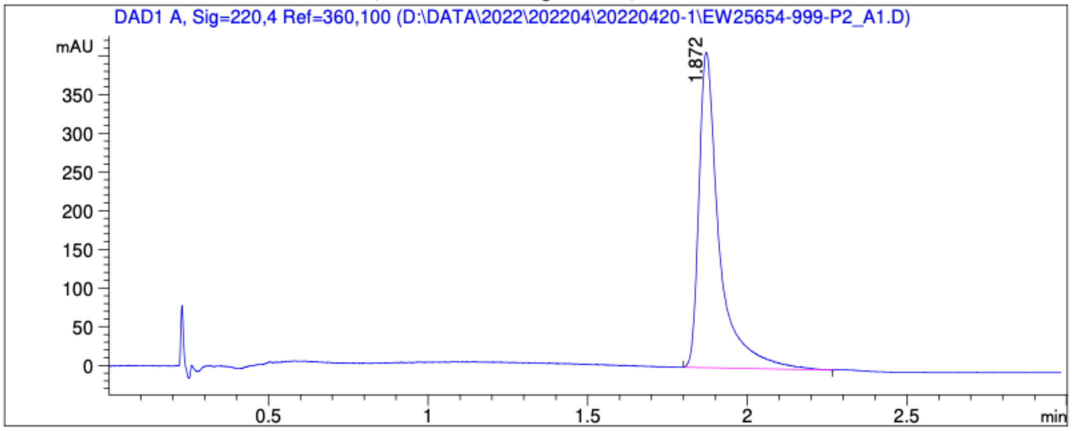


Signal: DAD1 A, Sig=220,4 Ref=360,100

| RetTime [min] | Height | Resolution | Symm. | Area [mAU*s] | Area % |
|---|---|---|---|---|---|
| 1.872 | 408.301 | | 0.634 | 1803.621 | 100.000 |

Mixture of WX-61b (WX-02-419) & WX-61d (WX-02-421)

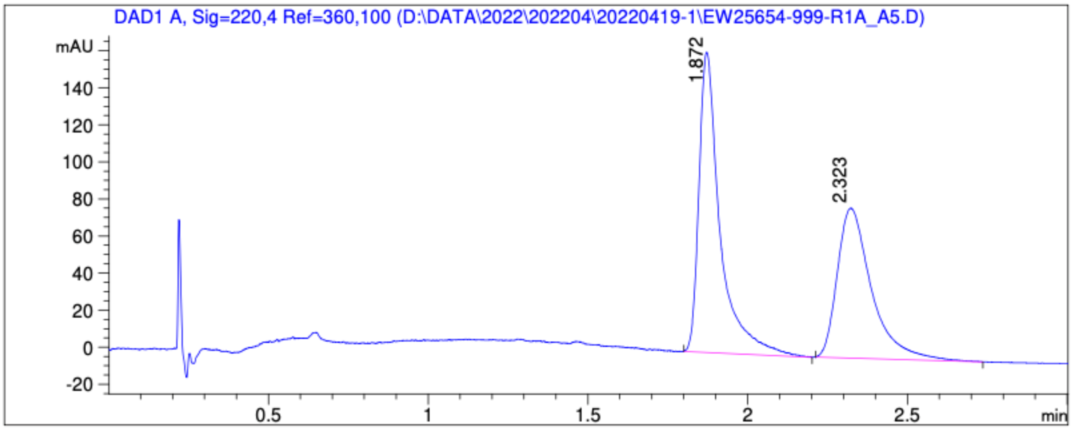


Signal: DAD1 A, Sig=220,4 Ref=360,100

| RetTime [min] | Height | Resolution | Symm. | Area [mAU*s] | Area % |
|---|---|---|---|---|---|
| 1.872 | 162.398 | | 0.612 | 745.010 | 55.273 |
| 2.323 | 80.963 | 3.208 | 0.569 | 602.871 | 44.727 |

Method details

Column: Chiralpak AS-3 50×4.6mm I.D., 3 μm
Mobile phase: A: $CO_2$, B: MeOH (0.05% diethylamine)
Elution: 5% to 40% B (gradient)

WX-62a (WX-02-422)

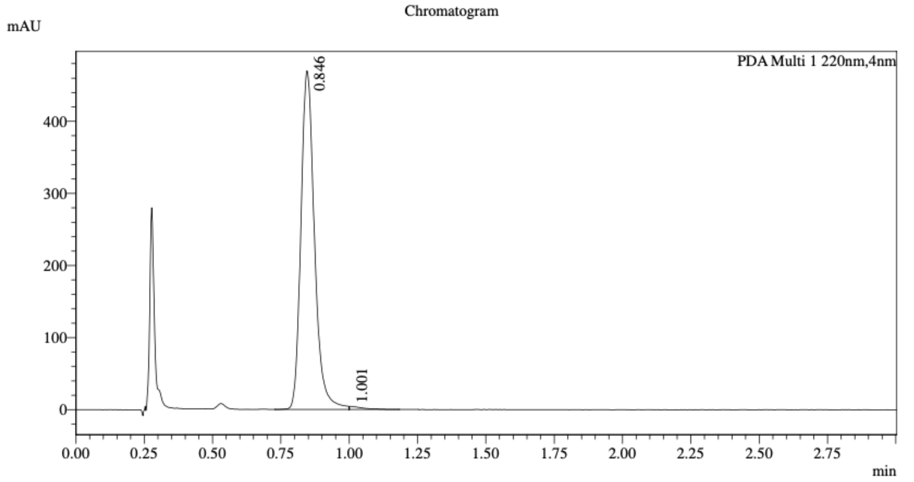


1 PDA Multi 1 / 220nm,4nm

Integration Result

Peak Table

PDA Ch1 220nm

| Peak# | Ret. Time | Height | Height% | Resolution(USP) | Area | Area% |
|---|---|---|---|---|---|---|
| 1 | 0.846 | 467285 | 99.079 | -- | 1624952 | 99.197 |
| 2 | 1.001 | 4346 | 0.921 | -- | 13160 | 0.803 |

WX-62c (WX-02-424)

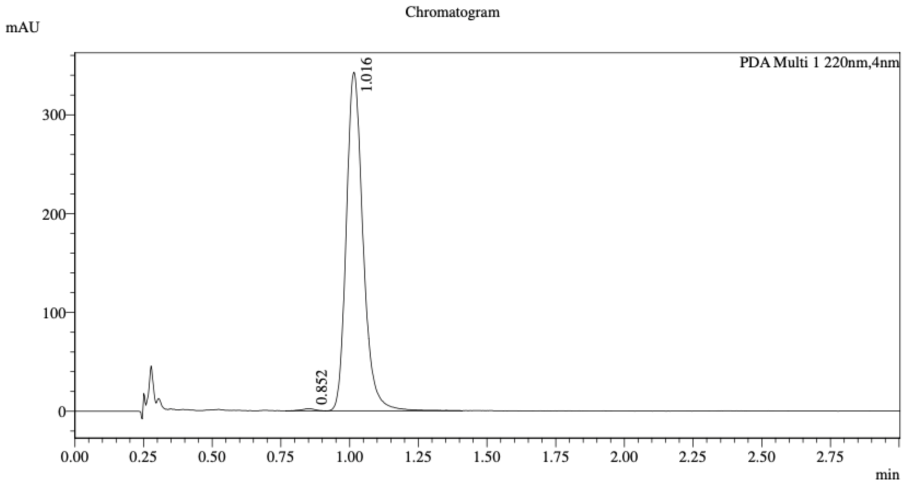


1 PDA Multi 1 / 220nm,4nm

Integration Result

Peak Table

PDA Ch1 220nm

| Peak# | Ret. Time | Height | Height% | Resolution(USP) | Area | Area% |
|---|---|---|---|---|---|---|
| 1 | 0.852 | 2081 | 0.608 | -- | 6547 | 0.449 |
| 2 | 1.016 | 340167 | 99.392 | 1.677 | 1452609 | 99.551 |

Mixture of WX-62a (WX-02-422) & WX-62c (WX-02-424)

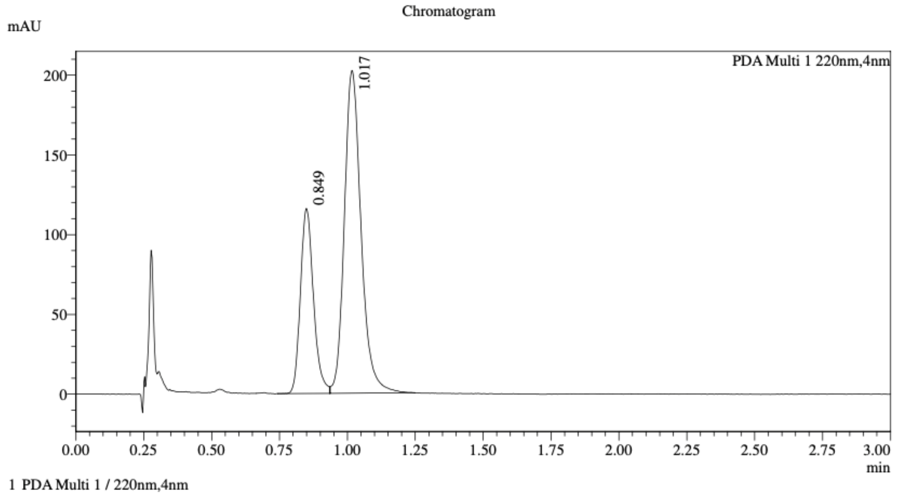


1 PDA Multi 1 / 220nm,4nm

Integration Result

Peak Table

PDA Ch1 220nm

| Peak# | Ret. Time | Height | Height% | Resolution(USP) | Area | Area% |
|---|---|---|---|---|---|---|
| 1 | 0.849 | 113612 | 36.252 | -- | 389002 | 30.978 |
| 2 | 1.017 | 199783 | 63.748 | 1.675 | 866724 | 69.022 |

Method details

Column: Chiralcel OD-3 50×4.6mm I.D., 3 μm
Mobile phase: A: $CO_2$, B: 2:1 MeOH/acetonitrile (0.05% diethylamine)
Elution: 40% B

WX-62b (WX-02-423)

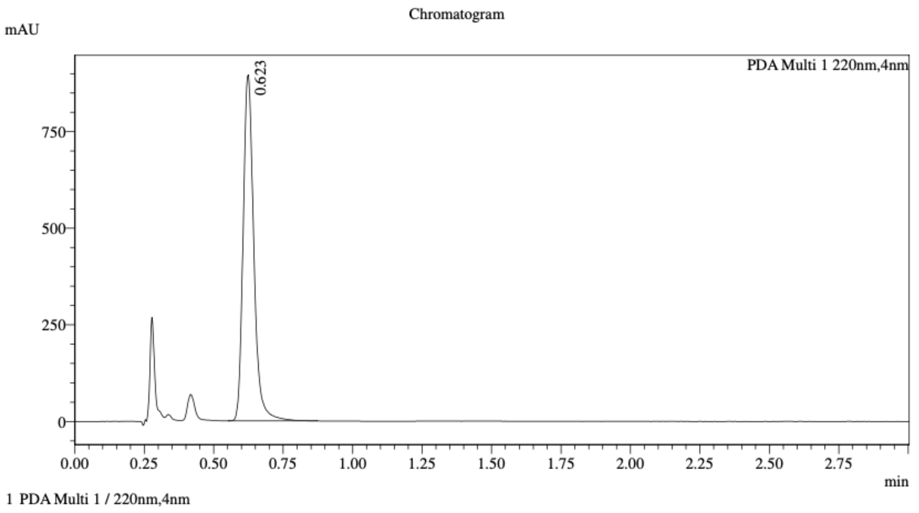


Integration Result

Peak Table

PDA Ch1 220nm

| Peak# | Ret. Time | Height | Height% | Resolution(USP) | Area | Area% |
|---|---|---|---|---|---|---|
| 1 | 0.623 | 882321 | 100.000 | -- | 2395424 | 100.000 |

WX-62d (WX-02-425)

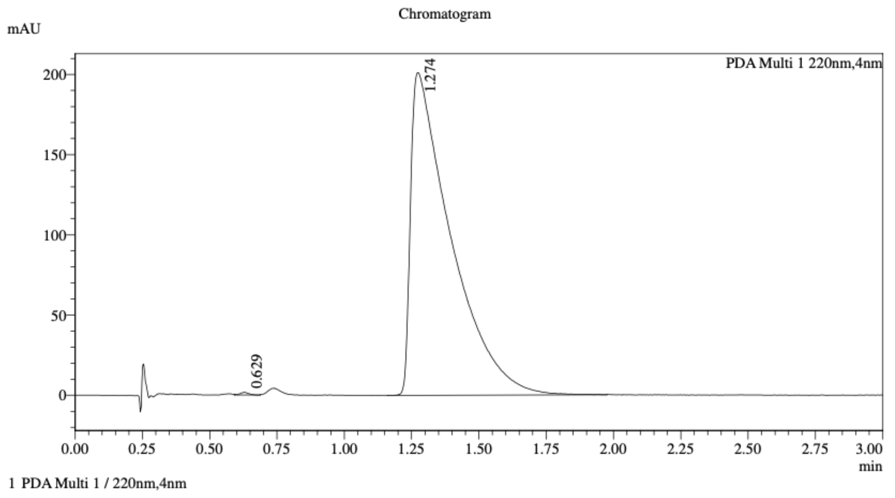


Integration Result

Peak Table

PDA Ch1 220nm

| Peak# | Ret. Time | Height | Height% | Resolution(USP) | Area | Area% |
|---|---|---|---|---|---|---|
| 1 | 0.629 | 1568 | 0.776 | -- | 3460 | 0.165 |
| 2 | 1.274 | 200352 | 99.224 | 3.912 | 2095085 | 99.835 |

Mixture of WX-62b (WX-02-423) & WX-62d (WX-02-425)

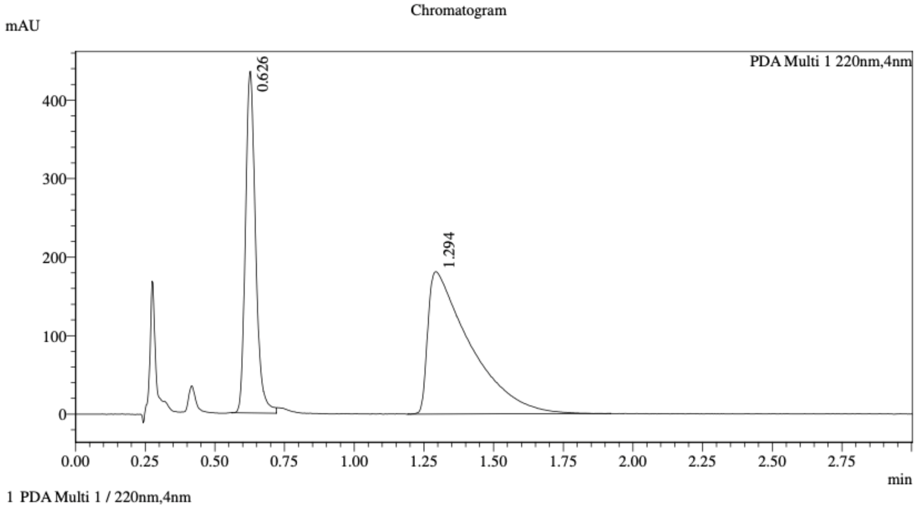


Integration Result

Peak Table

PDA Ch1 220nm

| Peak# | Ret. Time | Height | Height% | Resolution(USP) | Area | Area% |
|---|---|---|---|---|---|---|
| 1 | 0.626 | 423475 | 70.039 | -- | 1098553 | 36.444 |
| 2 | 1.294 | 181152 | 29.961 | 3.796 | 1915835 | 63.556 |

Method details

Column: Chiralcel OD-3 50×4.6mm I.D., 3 μm
Mobile phase: A: $CO_2$, B: 2:1 MeOH/acetonitrile (0.05% diethylamine)
Elution: 40% B

WX-63a (WX-02-426)

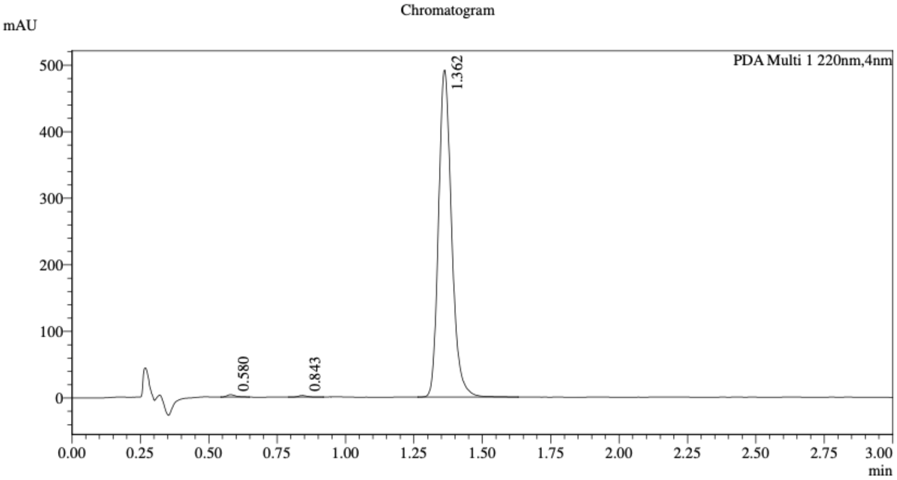


1 PDA Multi 1 / 220nm,4nm

Integration Result

Peak Table

PDA Ch1 220nm

| Peak# | Ret. Time | Height | Height% | Resolution(USP) | Area | Area% |
|---|---|---|---|---|---|---|
| 1 | 0.580 | 3453 | 0.708 | -- | 7501 | 0.456 |
| 2 | 0.843 | 2283 | 0.468 | 4.451 | 5183 | 0.315 |
| 3 | 1.362 | 481685 | 98.823 | 6.970 | 1632319 | 99.229 |

WX-63c (WX-02-427)

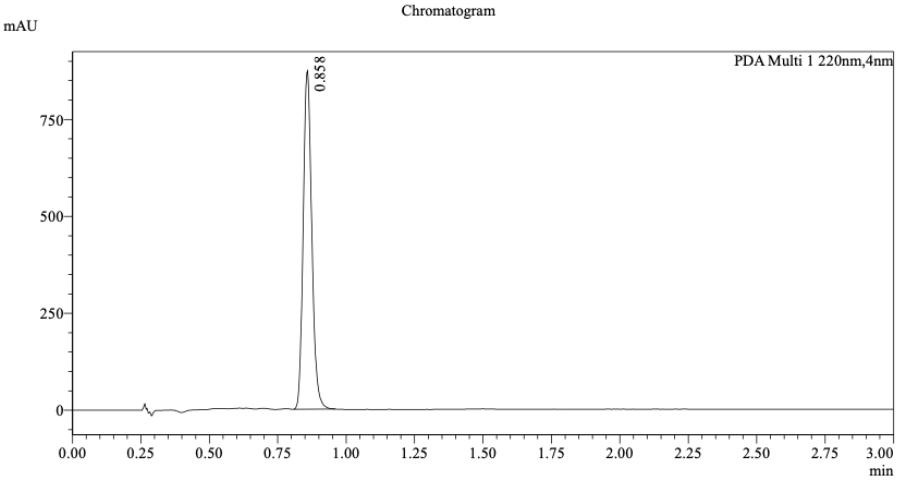


1 PDA Multi 1 / 220nm,4nm

Integration Result

Peak Table

PDA Ch1 220nm

| Peak# | Ret. Time | Height | Height% | Resolution(USP) | Area | Area% |
|---|---|---|---|---|---|---|
| 1 | 0.858 | 837735 | 100.000 | -- | 1909269 | 100.000 |

Mixture of WX-63a (WX-02-426) & WX-63c (WX-02-427)

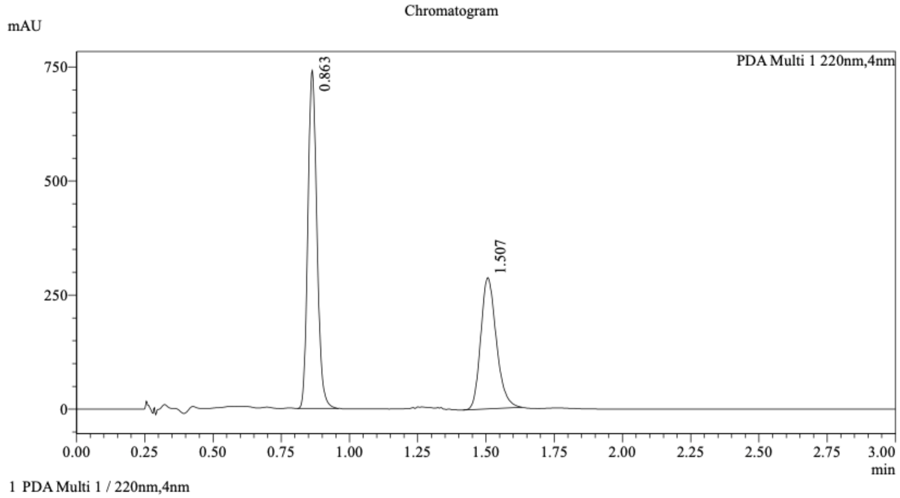


1 PDA Multi 1 / 220nm,4nm

Integration Result

Peak Table

PDA Ch1 220nm

| Peak# | Ret. Time | Height | Height% | Resolution(USP) | Area | Area% |
|---|---|---|---|---|---|---|
| 1 | 0.863 | 727653 | 71.865 | -- | 1714692 | 59.976 |
| 2 | 1.507 | 284868 | 28.135 | 7.590 | 1144278 | 40.024 |

Method details

Column: (*S*,*S*)Whelk-O1 50×4.6mm I.D., 3.5 µm
Mobile phase: A: $CO_2$, B: 2:1 i-PrOH/acetonitrile (0.05% diethylamine)
Elution: 40% B

WX-63b (WX-02-428)

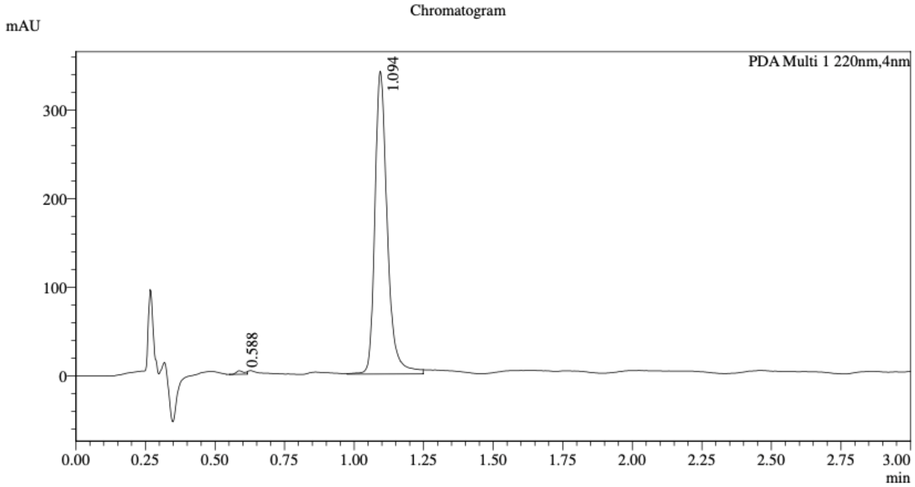


1 PDA Multi 1 / 220nm,4nm

Integration Result

Peak Table

PDA Ch1 220nm

| Peak# | Ret. Time | Height | Height% | Resolution(USP) | Area | Area% |
|---|---|---|---|---|---|---|
| 1 | 0.588 | 3874 | 1.129 | -- | 8019 | 0.748 |
| 2 | 1.094 | 339452 | 98.871 | 7.466 | 1064462 | 99.252 |

WX-63d (WX-02-429)

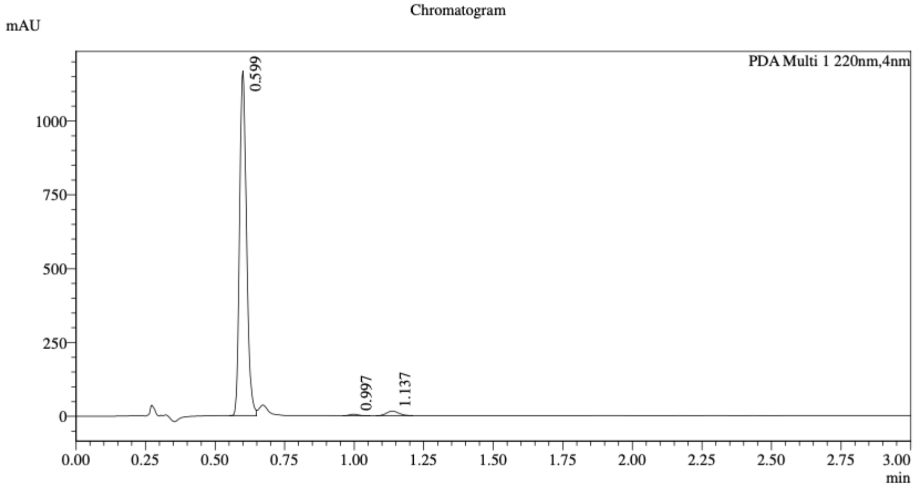


1 PDA Multi 1 / 220nm,4nm

Integration Result

Peak Table

PDA Ch1 220nm

| Peak# | Ret. Time | Height | Height% | Resolution(USP) | Area | Area% |
|---|---|---|---|---|---|---|
| 1 | 0.599 | 1124114 | 98.215 | -- | 2016960 | 97.115 |
| 2 | 0.997 | 5171 | 0.452 | 7.054 | 11738 | 0.565 |
| 3 | 1.137 | 15264 | 1.334 | 1.880 | 48181 | 2.320 |

Mixture of WX-63b (WX-02-428) & WX-63d (WX-02-429)

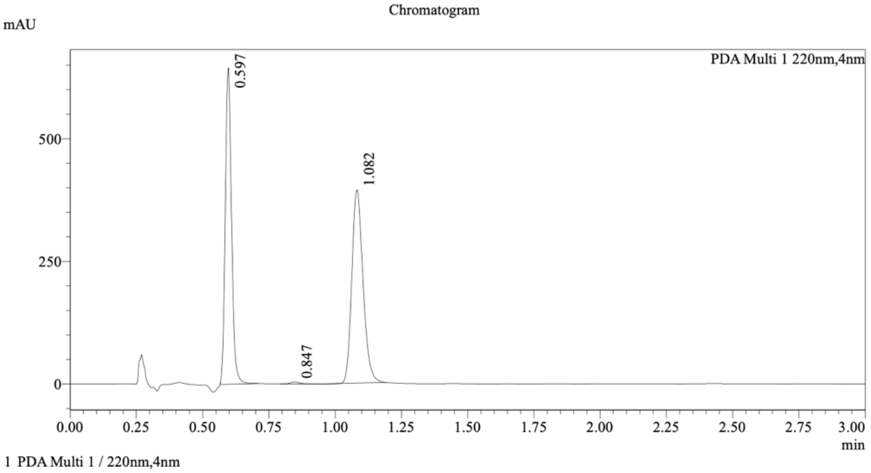


Integration Result

Peak Table

PDA Ch1 220nm

| Peak# | Ret. Time | Width | Height | Resolution | Tailing Factor | Area | Area% |
|---|---|---|---|---|---|---|---|
| 1 | 0.597 | 0.048 | 631181 | -- | 1.156 | 1058332 | 47.466 |
| 2 | 0.847 | 0.062 | 3975 | 4.589 | 1.341 | 9182 | 0.412 |
| 3 | 1.082 | 0.080 | 385858 | 3.329 | 1.207 | 1162156 | 52.122 |

Method details

Column: (*S*,*S*)Whelk-O1 50×4.6mm I.D., 3.5 µm
Mobile phase: A: $CO_2$, B: 2:1 MeOH/acetonitrile (0.05% diethylamine)
Elution: 40% B

WX-71a (WX-02-430)

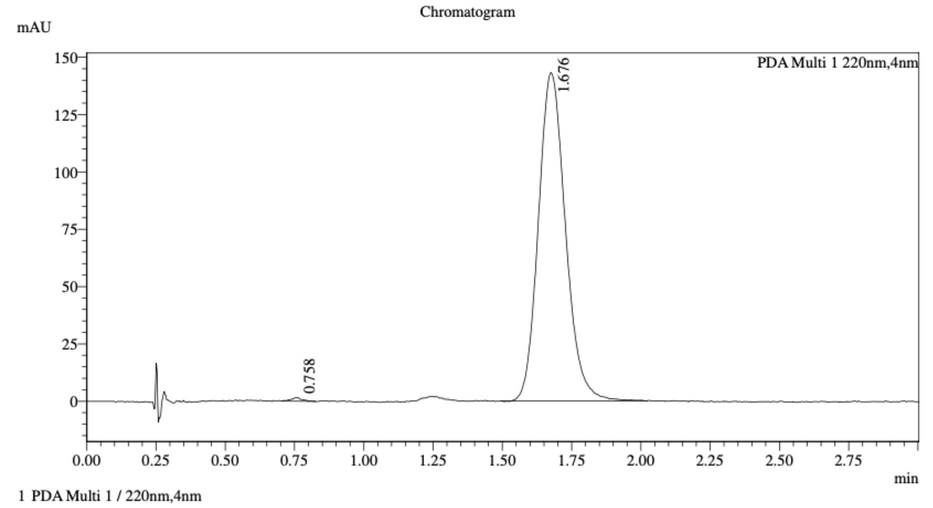


Integration Result

Peak Table

PDA Ch1 220nm

| Peak# | Ret. Time | Height | Height% | Resolution(USP) | Area | Area% |
|---|---|---|---|---|---|---|
| 1 | 0.758 | 1434 | 0.994 | -- | 3977 | 0.407 |
| 2 | 1.676 | 142825 | 99.006 | 7.387 | 973116 | 99.593 |

WX-71c (WX-02-431)

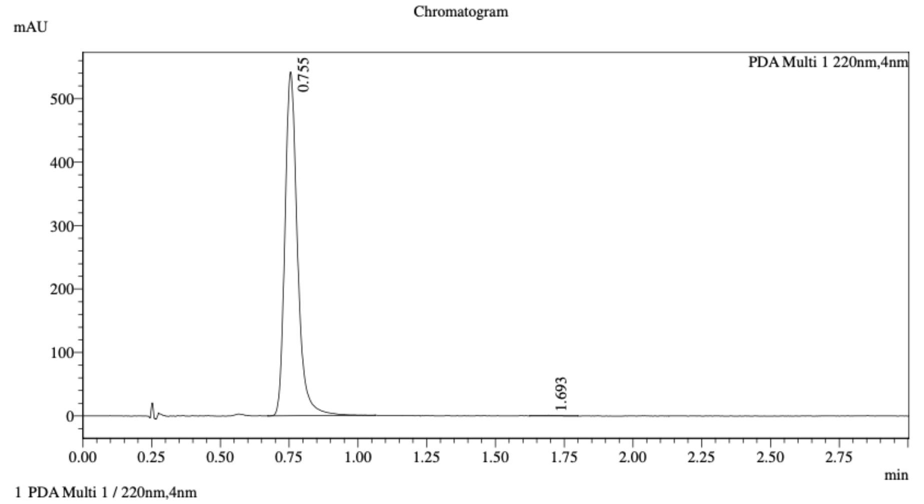


Integration Result

Peak Table

PDA Ch1 220nm

| Peak# | Ret. Time | Height | Height% | Resolution(USP) | Area | Area% |
|---|---|---|---|---|---|---|
| 1 | 0.755 | 535072 | 99.845 | -- | 1715791 | 99.787 |
| 2 | 1.693 | 832 | 0.155 | 8.670 | 3661 | 0.213 |

Mixture of WX-71a (WX-02-430) & WX-71c (WX-02-431)

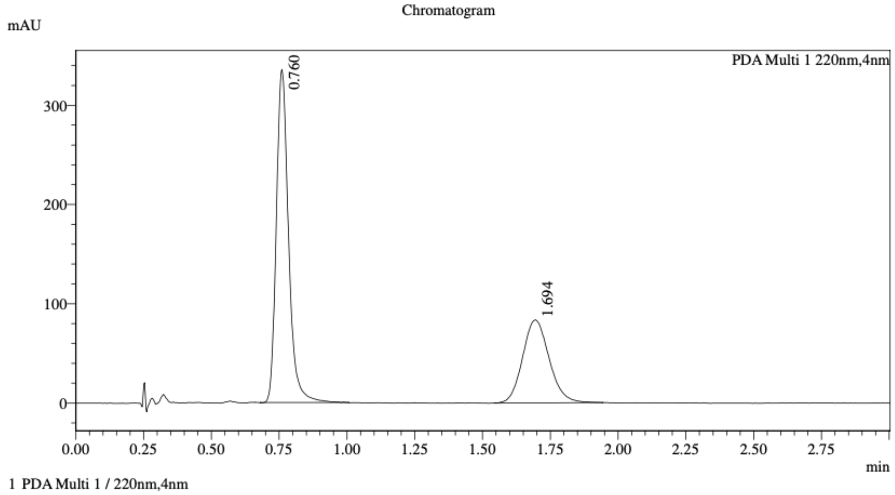

1 PDA Multi 1 / 220nm,4nm

Integration Result

Peak Table

PDA Ch1 220nm

| Peak# | Ret. Time | Height | Height% | Resolution(USP) | Area | Area% |
|---|---|---|---|---|---|---|
| 1 | 0.760 | 330364 | 79.836 | -- | 1052606 | 64.901 |
| 2 | 1.694 | 83442 | 20.164 | 7.142 | 569270 | 35.099 |

Method details

Column: Chiralcel OD-3 50×4.6mm I.D., 3 μm
Mobile phase: A: $CO_2$, B: 2:1 MeOH/acetonitrile (0.05% diethylamine)
Elution: 40% B

WX-71b (WX-02-432)

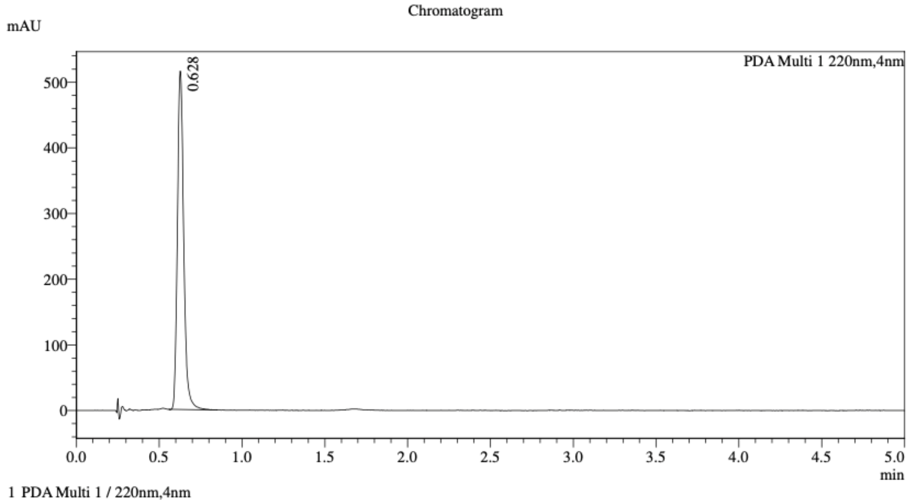


Integration Result

Peak Table

PDA Ch1 220nm

| Peak# | Ret. Time | Height | Height% | Resolution(USP) | Area | Area% |
|---|---|---|---|---|---|---|
| 1 | 0.628 | 507753 | 100.000 | -- | 1362398 | 100.000 |

WX-71d (WX-02-433)

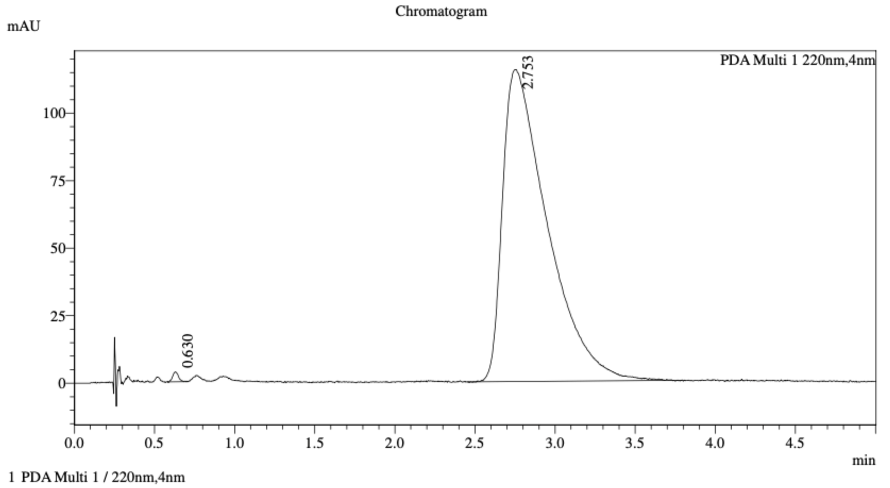


Integration Result

Peak Table

PDA Ch1 220nm

| Peak# | Ret. Time | Height | Height% | Resolution(USP) | Area | Area% |
|---|---|---|---|---|---|---|
| 1 | 0.630 | 3598 | 3.022 | -- | 9294 | 0.411 |
| 2 | 2.753 | 115459 | 96.978 | 7.328 | 2249301 | 99.589 |

Mixture of WX-71b (WX-02-432) & WX-71d (WX-02-433)

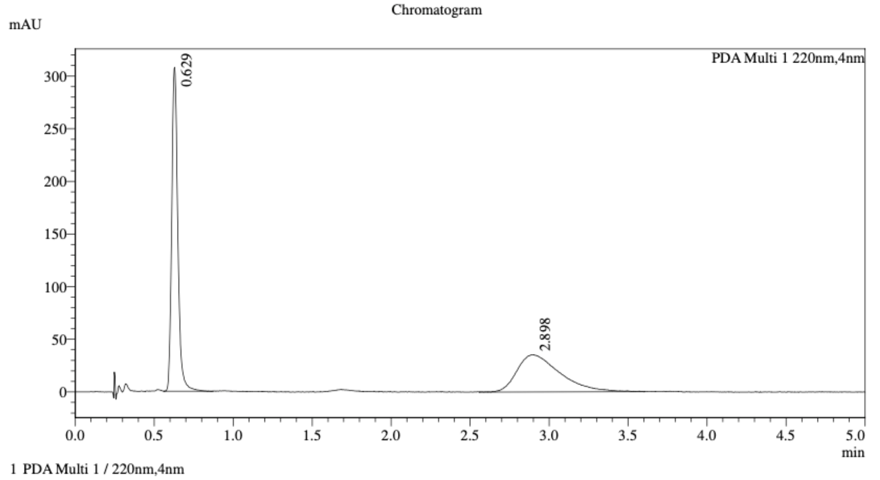


Integration Result

Peak Table

PDA Ch1 220nm

| Peak# | Ret. Time | Height | Height% | Resolution(USP) | Area | Area% |
|---|---|---|---|---|---|---|
| 1 | 0.629 | 304826 | 89.641 | -- | 823057 | 55.232 |
| 2 | 2.898 | 35224 | 10.359 | 8.153 | 667131 | 44.768 |

Method details

Column: Chiralcel OD-3 50×4.6mm I.D., 3 μm
Mobile phase: A: $CO_2$, B: 2:1 MeOH/acetonitrile (0.05% diethylamine)
Elution: 40% B

WX-72a (WX-02-434)

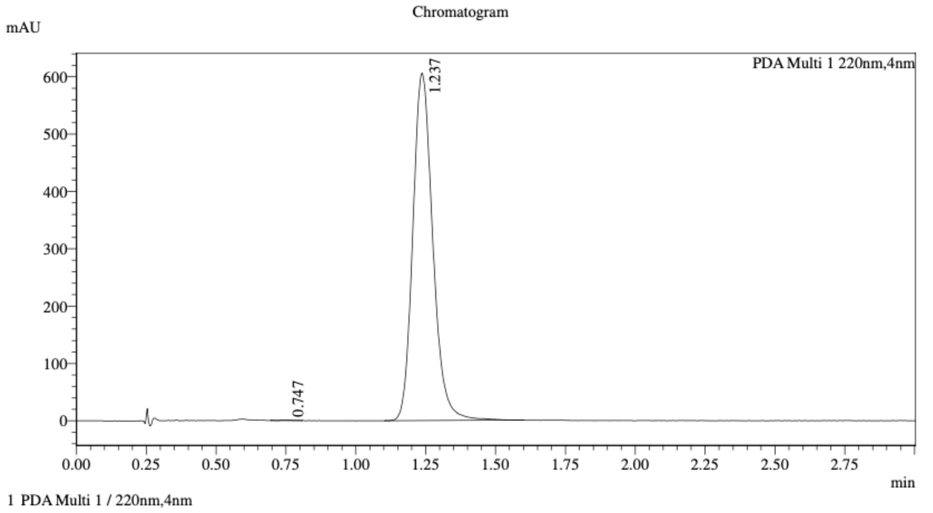


Integration Result

Peak Table

PDA Ch1 220nm

| Peak# | Ret. Time | Height | Height% | Resolution(USP) | Area | Area% |
|---|---|---|---|---|---|---|
| 1 | 0.747 | 496 | 0.082 | -- | 1379 | 0.046 |
| 2 | 1.237 | 604579 | 99.918 | 4.243 | 3016794 | 99.954 |

WX-72c (WX-02-435)

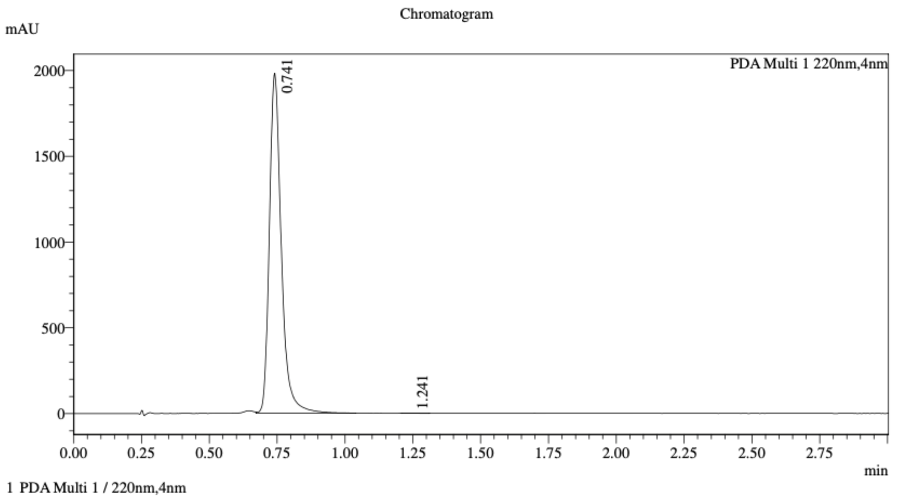


Integration Result

Peak Table

PDA Ch1 220nm

| Peak# | Ret. Time | Height | Height% | Resolution(USP) | Area | Area% |
|---|---|---|---|---|---|---|
| 1 | 0.741 | 1966461 | 99.973 | -- | 6057427 | 99.971 |
| 2 | 1.241 | 535 | 0.027 | 6.325 | 1785 | 0.029 |

Mixture of WX-72a (WX-02-434) & WX-72c (WX-02-435)

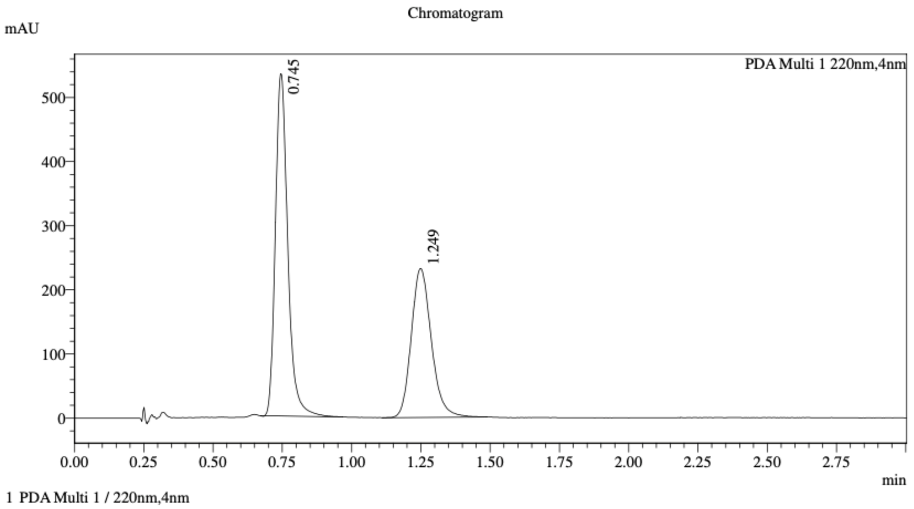


Integration Result

Peak Table

PDA Ch1 220nm

| Peak# | Ret. Time | Height | Height% | Resolution(USP) | Area | Area% |
|---|---|---|---|---|---|---|
| 1 | 0.745 | 521157 | 69.352 | -- | 1659045 | 58.654 |
| 2 | 1.249 | 230305 | 30.648 | 4.722 | 1169467 | 41.346 |

Method details

Column: Chiralcel OD-3 50×4.6mm I.D., 3 µm
Mobile phase: A: $CO_2$, B: 2:1 MeOH/acetonitrile (0.05% diethylamine)
Elution: 40% B

WX-72b (WX-02-436)

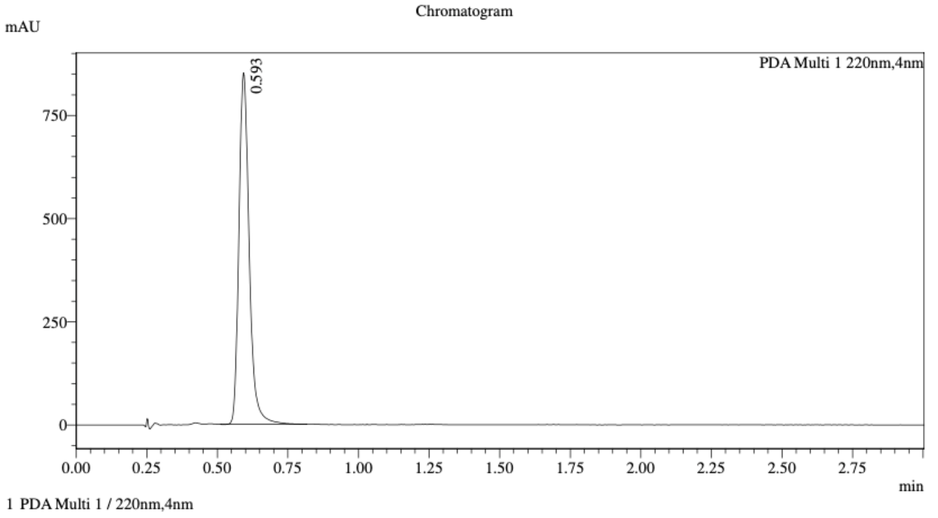


Integration Result

Peak Table

PDA Ch1 220nm

| Peak# | Ret. Time | Height | Height% | Resolution(USP) | Area | Area% |
|---|---|---|---|---|---|---|
| 1 | 0.593 | 823824 | 100.000 | -- | 2160153 | 100.000 |

WX-72d (WX-02-437)

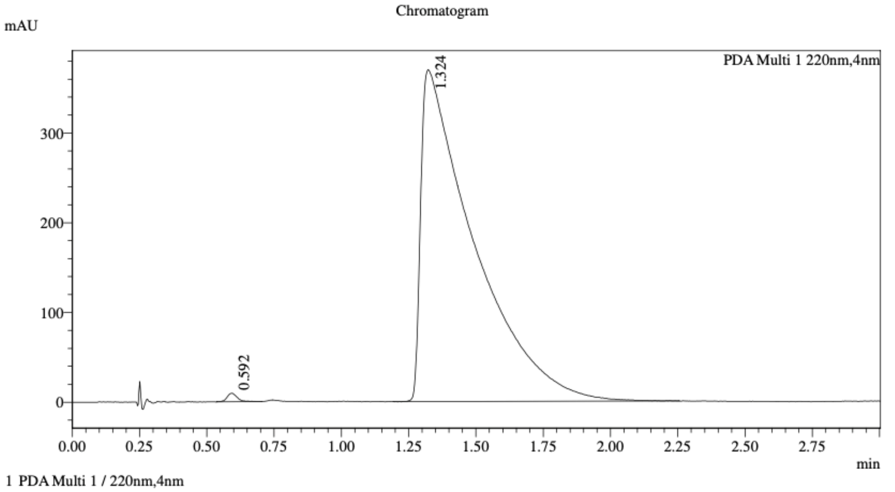


Integration Result

Peak Table

PDA Ch1 220nm

| Peak# | Ret. Time | Height | Height% | Resolution(USP) | Area | Area% |
|---|---|---|---|---|---|---|
| 1 | 0.592 | 9322 | 2.465 | -- | 25048 | 0.492 |
| 2 | 1.324 | 368886 | 97.535 | 3.368 | 5069840 | 99.508 |

Mixture of WX-72b (WX-02-436) & WX-72d (WX-02-437)

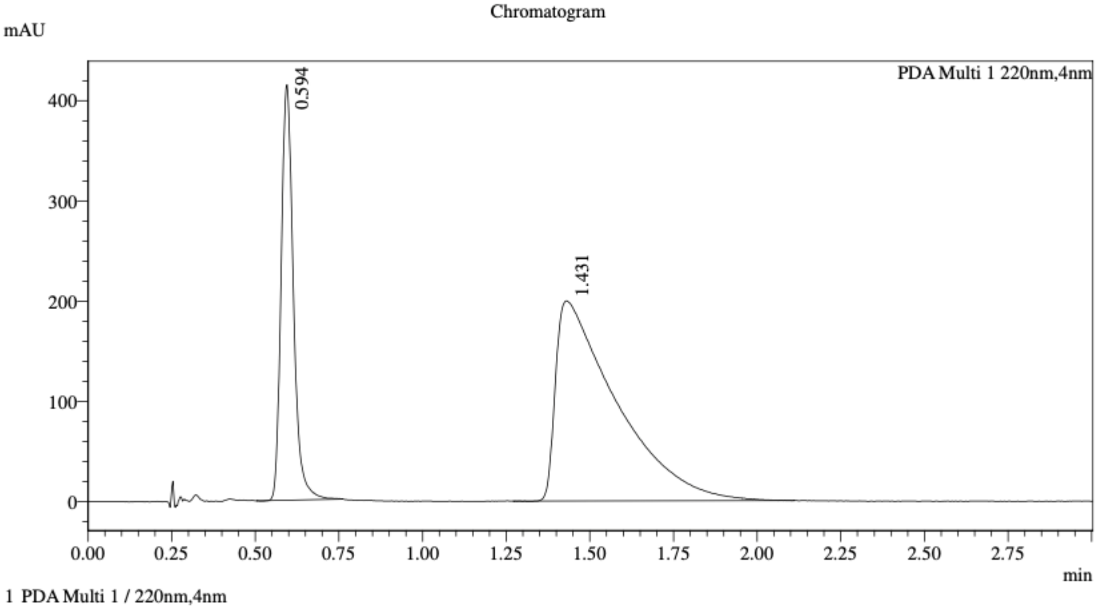


Integration Result

Peak Table

PDA Ch1 220nm

| Peak# | Ret. Time | Height | Height% | Resolution(USP) | Area | Area% |
|---|---|---|---|---|---|---|
| 1 | 0.594 | 402594 | 66.858 | -- | 1054681 | 29.451 |
| 2 | 1.431 | 199569 | 33.142 | 4.116 | 2526408 | 70.549 |

Method details

Column: Chiralcel OD-3 50×4.6mm I.D., 3 μm
Mobile phase: A: $CO_2$, B: 2:1MeOH/acetonitrile (0.05% diethylamine)
Elution: 40% B

WX-73a (WX-02-438)

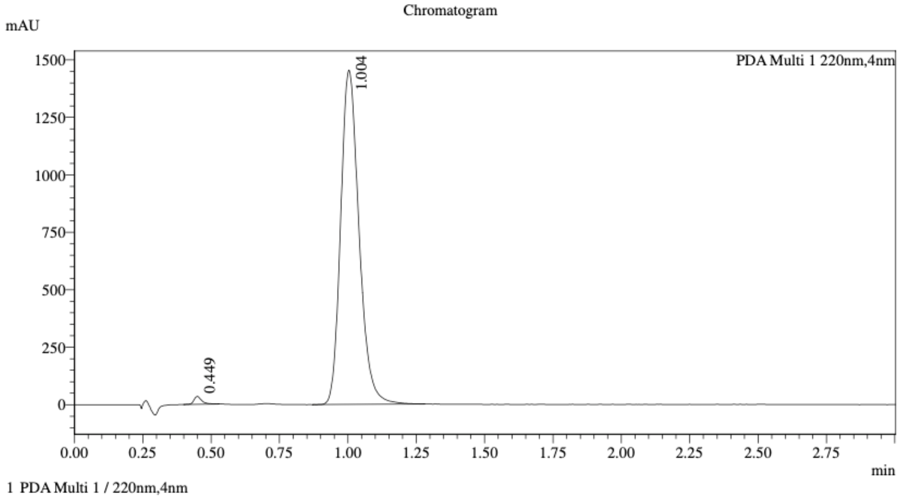


Integration Result

Peak Table

PDA Ch1 220nm

| Peak# | Ret. Time | Height | Height% | Resolution(USP) | Area | Area% |
|---|---|---|---|---|---|---|
| 1 | 0.449 | 31606 | 2.134 | -- | 67359 | 0.981 |
| 2 | 1.004 | 1449217 | 97.866 | 6.271 | 6802183 | 99.019 |

WX-73c (WX-02-439)

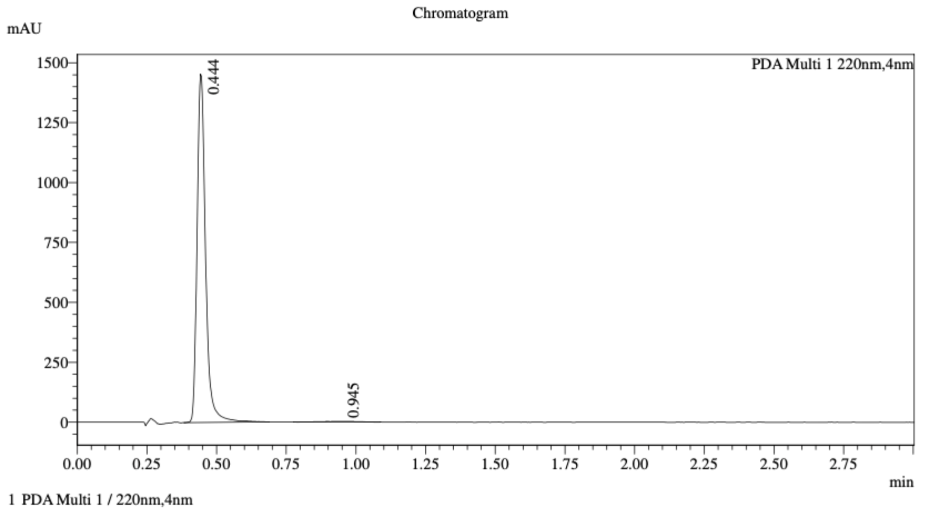


Integration Result

Peak Table

PDA Ch1 220nm

| Peak# | Ret. Time | Height | Height% | Resolution(USP) | Area | Area% |
|---|---|---|---|---|---|---|
| 1 | 0.444 | 1423319 | 99.808 | -- | 3082642 | 99.275 |
| 2 | 0.945 | 2739 | 0.192 | 3.560 | 22524 | 0.725 |

Mixture of WX-73a (WX-02-438) & WX-73c (WX-02-439)

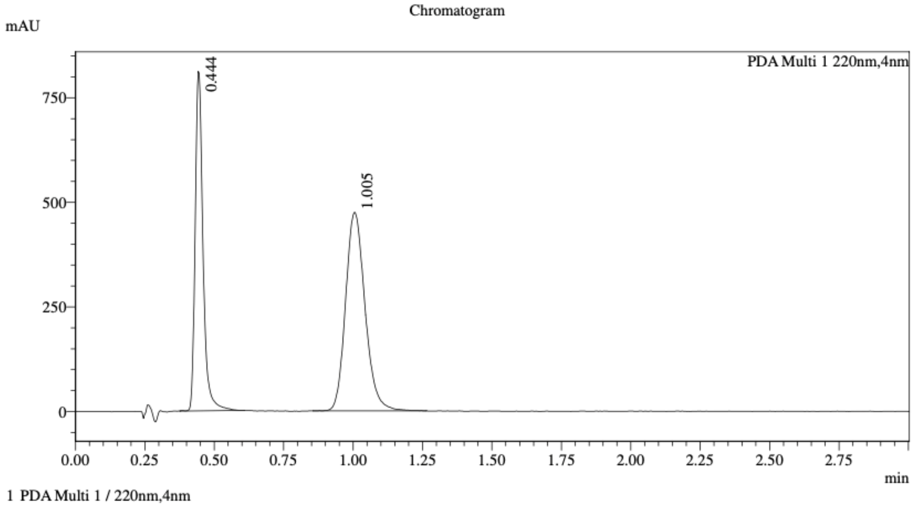


Integration Result

Peak Table

PDA Ch1 220nm

| Peak# | Ret. Time | Height | Height% | Resolution(USP) | Area | Area% |
|---|---|---|---|---|---|---|
| 1 | 0.444 | 793328 | 62.611 | -- | 1592574 | 40.756 |
| 2 | 1.005 | 473752 | 37.389 | 6.198 | 2315007 | 59.244 |

Method details

Column: Chiralcel OD-3 50×4.6mm I.D., 3 μm
Mobile phase: A: $CO_2$, B: X:X i-PrOH/acetonitrile (0.05% diethylamine)
Elution: 40% B

WX-73b (WX-02-440)

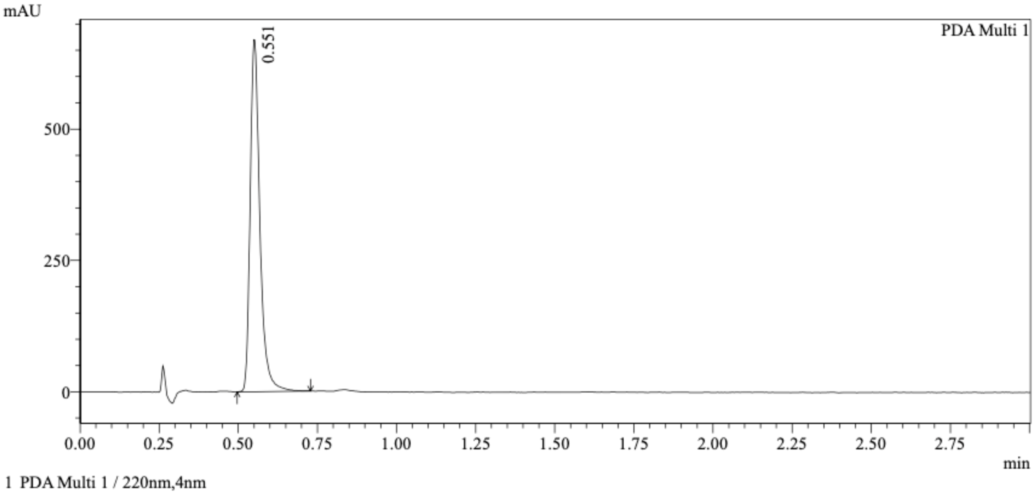


Integration Results

PeakTable

PDA Ch1 220nm

| Peak# | Ret. Time | USP Width | Resolution | Height | Area | Area % |
|---|---|---|---|---|---|---|
| 1 | 0.551 | 0.057 | 0.000 | 659973 | 1428604 | 100.000 |
| Total | | | | 659973 | 1428604 | 100.000 |

WX-73d (WX-02-441)

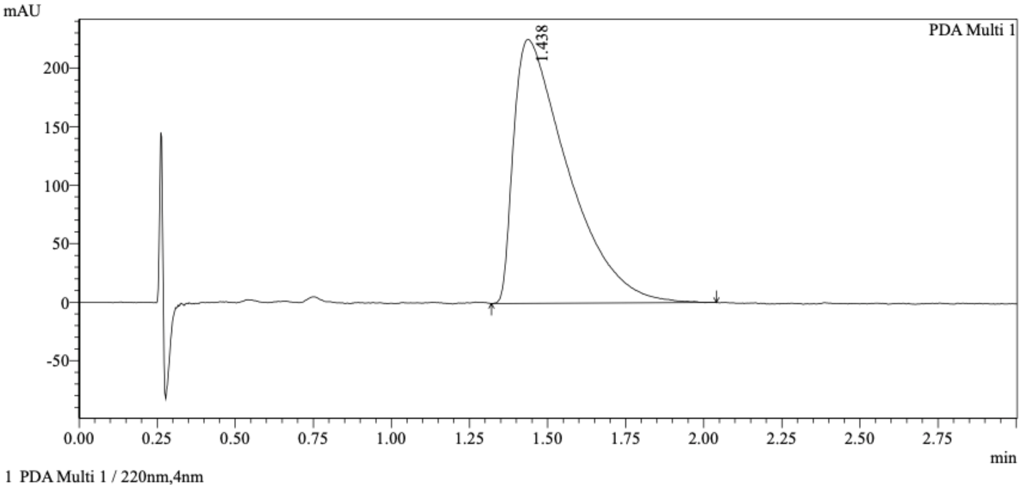


Integration Results

PeakTable

PDA Ch1 220nm

| Peak# | Ret. Time | USP Width | Resolution | Height | Area | Area % |
|---|---|---|---|---|---|---|
| 1 | 1.438 | 0.317 | 0.000 | 225422 | 2733925 | 100.000 |
| Total | | | | 225422 | 2733925 | 100.000 |

Mixture of WX-73b (WX-02-440) & WX-73d (WX-02-441)

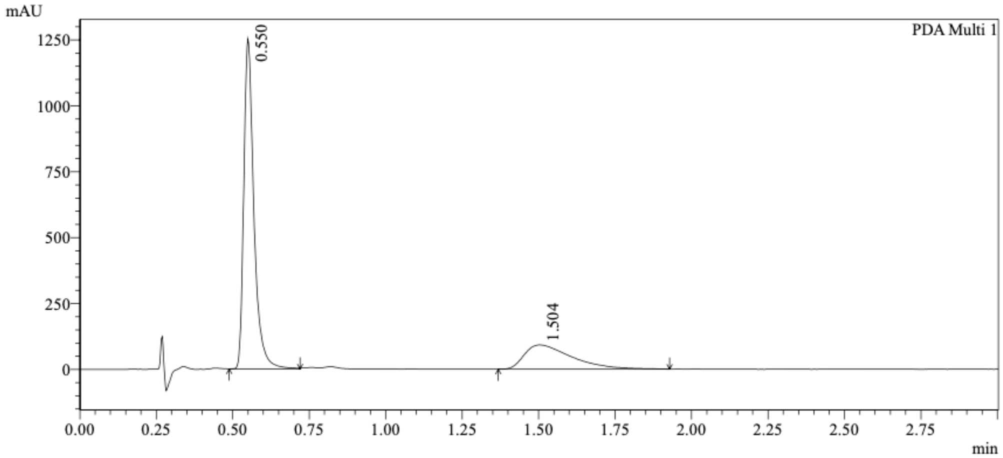


1 PDA Multi 1 / 220nm,4nm

Integration Results

PeakTable

PDA Ch1 220nm

| Peak# | Ret. Time | USP Width | Resolution | Height | Area | Area % |
|---|---|---|---|---|---|---|
| 1 | 0.550 | 0.061 | 0.000 | 1235199 | 2884335 | 74.358 |
| 2 | 1.504 | 0.280 | 5.588 | 92076 | 994635 | 25.642 |
| Total | | | | 1327274 | 3878971 | 100.000 |

Method details

Column: Chiralcel OD-3 50×4.6mm I.D., 3 µm
Mobile phase: A: $CO_2$, B: 2:1 MeOH/acetonitrile (0.05% diethylamine)
Elution: 40% B

WX-64a (WX-02-687)

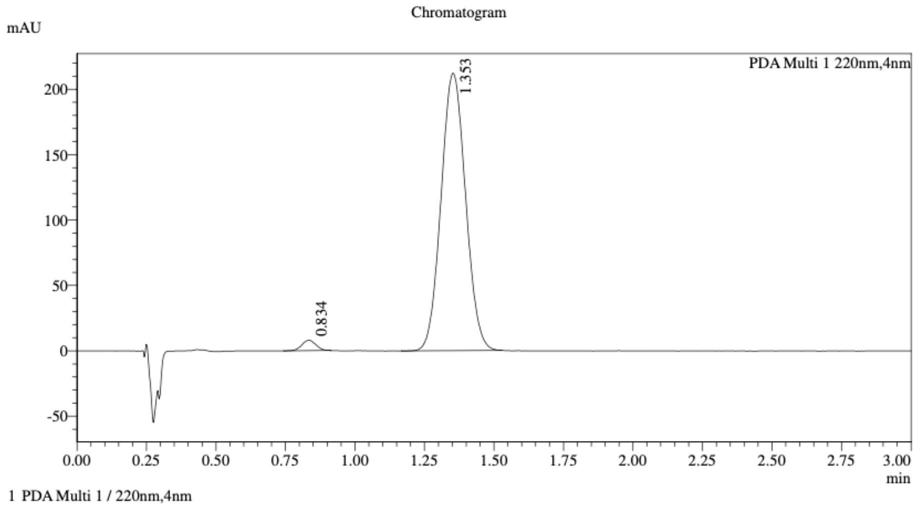


1 PDA Multi 1 / 220nm,4nm

Integration Result

Peak Table

PDA Ch1 220nm

| Peak# | Ret. Time | Height | Height% | Resolution(USP) | Area | Area% |
|---|---|---|---|---|---|---|
| 1 | 0.834 | 7881 | 3.599 | -- | 28180 | 2.119 |
| 2 | 1.353 | 211068 | 96.401 | 3.976 | 1301935 | 97.881 |

WX-64c (WX-02-688)

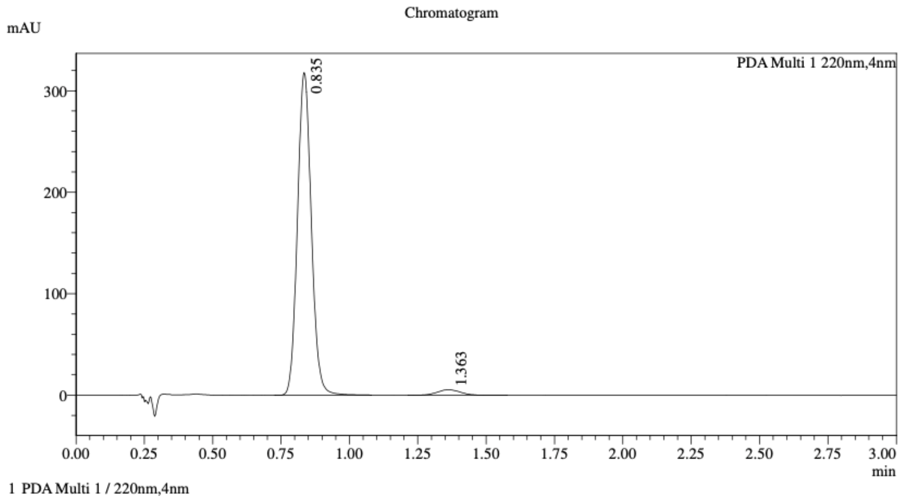


1 PDA Multi 1 / 220nm,4nm

Integration Result

Peak Table

PDA Ch1 220nm

| Peak# | Ret. Time | Height | Height% | Resolution(USP) | Area | Area% |
|---|---|---|---|---|---|---|
| 1 | 0.835 | 314275 | 98.309 | -- | 1121862 | 97.190 |
| 2 | 1.363 | 5407 | 1.691 | 4.165 | 32436 | 2.810 |

Mixture of WX-64a (WX-02-687) & WX-64c (WX-02-688)

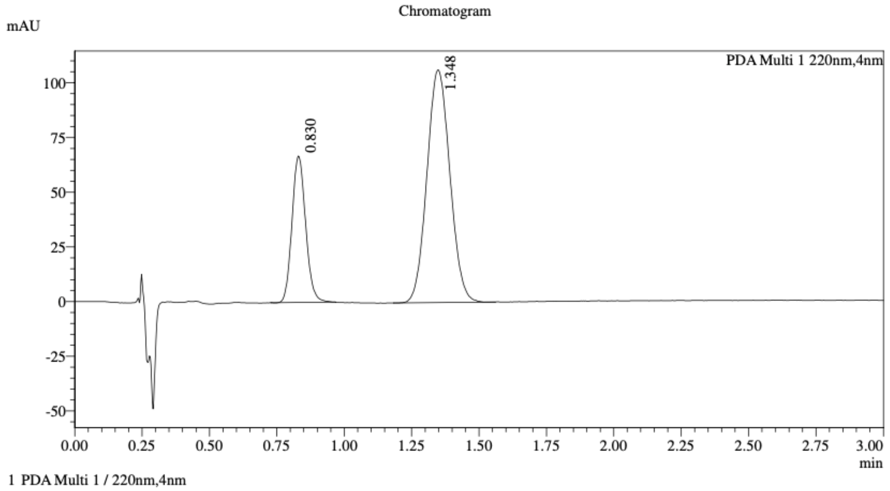


Integration Result

Peak Table

PDA Ch1 220nm

| Peak# | Ret. Time | Height | Height% | Resolution(USP) | Area | Area% |
|---|---|---|---|---|---|---|
| 1 | 0.830 | 66592 | 38.556 | -- | 236080 | 27.258 |
| 2 | 1.348 | 106124 | 61.444 | 4.087 | 630011 | 72.742 |

Method details

Column: Chiralpak AD-3 50×4.6mm I.D., 3 µm
Mobile phase: A: $CO_2$, B: 2:1 i-PrOH/acetonitrile (0.05% diethylamine)
Elution: 50% B

WX-64b (WX-02-689)

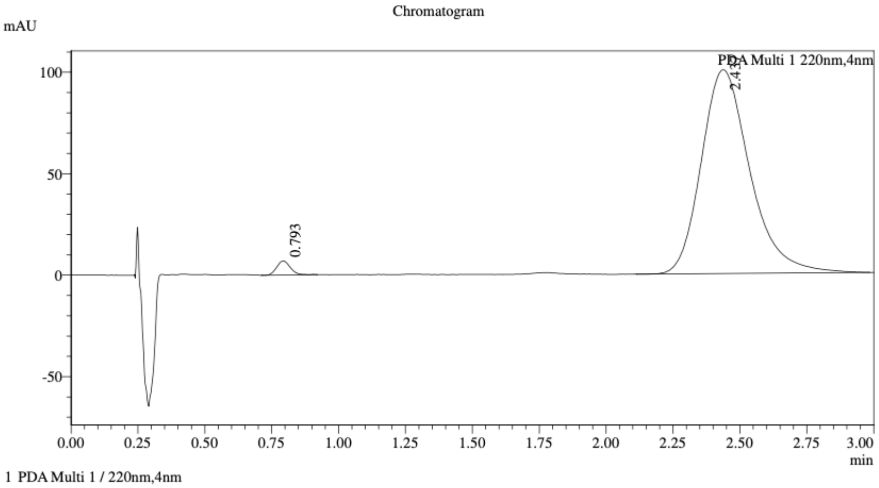


Integration Result

Peak Table

PDA Ch1 220nm

| Peak# | Ret. Time | Height | Height% | Resolution(USP) | Area | Area% |
|---|---|---|---|---|---|---|
| 1 | 0.793 | 6807 | 6.352 | -- | 24394 | 1.889 |
| 2 | 2.437 | 100359 | 93.648 | 7.846 | 1266781 | 98.111 |

WX-64d (WX-02-690)

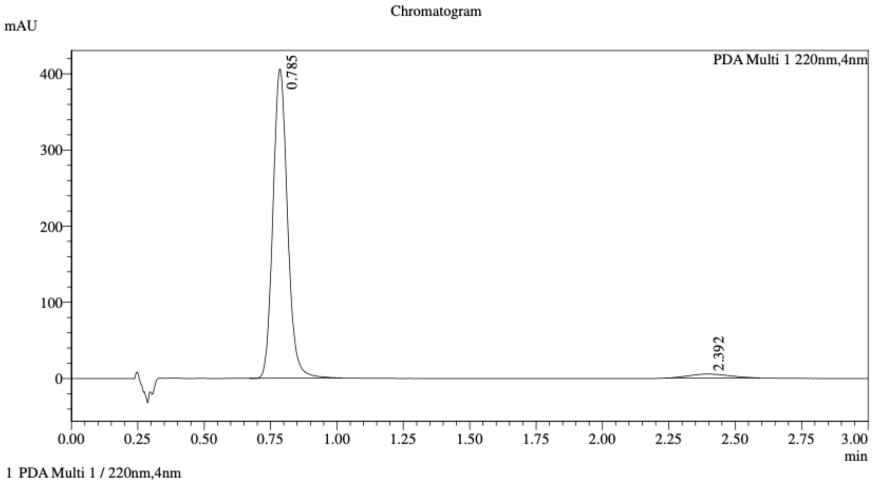


Integration Result

Peak Table

PDA Ch1 220nm

| Peak# | Ret. Time | Height | Height% | Resolution(USP) | Area | Area% |
|---|---|---|---|---|---|---|
| 1 | 0.785 | 398826 | 98.726 | -- | 1545456 | 96.601 |
| 2 | 2.392 | 5145 | 1.274 | 8.316 | 54385 | 3.399 |

Mixture of WX-64b (WX-02-689) & WX-64d (WX-02-690)

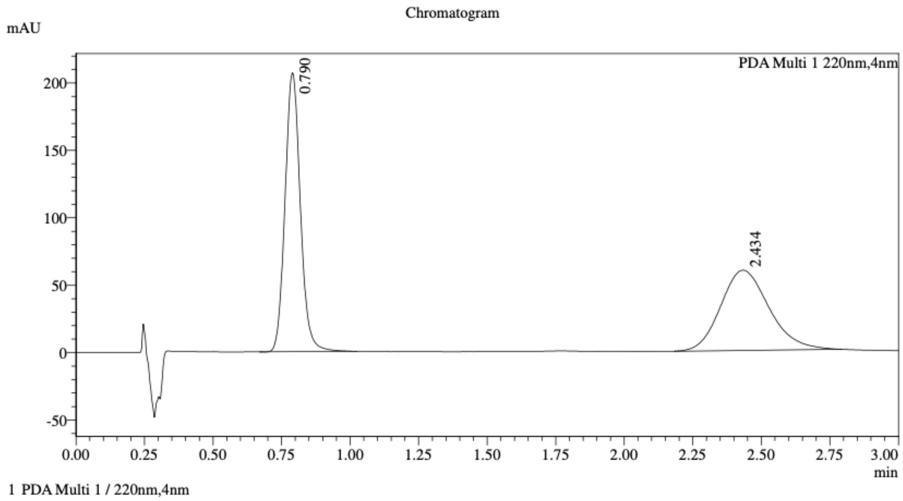


1 PDA Multi 1 / 220nm,4nm

Integration Result

Peak Table

PDA Ch1 220nm

| Peak# | Ret. Time | Height | Height% | Resolution(USP) | Area | Area% |
|---|---|---|---|---|---|---|
| 1 | 0.790 | 206044 | 77.601 | -- | 805181 | 52.373 |
| 2 | 2.434 | 59474 | 22.399 | 7.730 | 732210 | 47.627 |

Method details

Column: Chiralpak AD-3 50×4.6mm I.D., 3 μm

Mobile phase: A: $CO_2$, B: 2:1 i-PrOH/acetonitrile (0.05% diethylamine)

Elution: 40% B

WX-74a (WX-02-691)

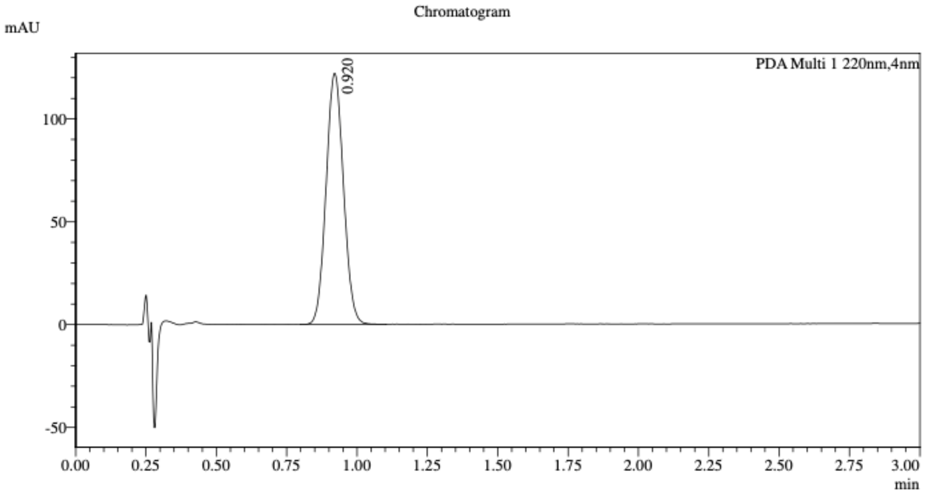


1 PDA Multi 1 / 220nm,4nm

Integration Result

Peak Table

PDA Ch1 220nm

| Peak# | Ret. Time | Height | Height% | Resolution(USP) | Area | Area% |
|---|---|---|---|---|---|---|
| 1 | 0.920 | 121060 | 100.000 | -- | 525449 | 100.000 |

WX-74c (WX-02-692)

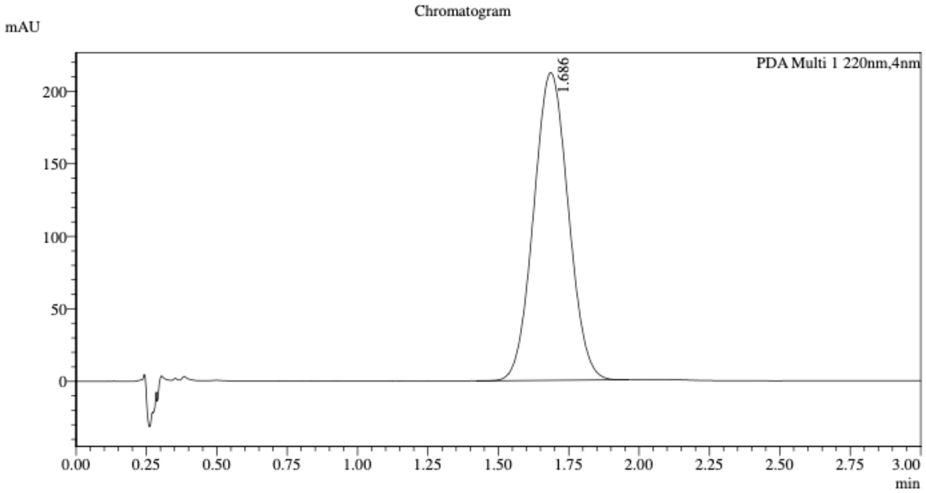


1 PDA Multi 1 / 220nm,4nm

Integration Result

Peak Table

PDA Ch1 220nm

| Peak# | Ret. Time | Height | Height% | Resolution(USP) | Area | Area% |
|---|---|---|---|---|---|---|
| 1 | 1.686 | 212113 | 100.000 | -- | 1820906 | 100.000 |

Mixture of WX-74a (WX-02-691) & WX-74c (WX-02-692)

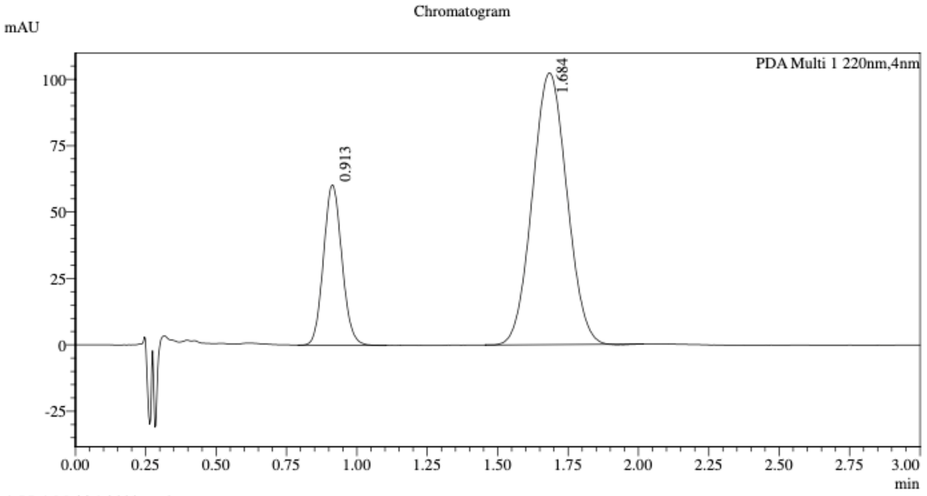


1 PDA Multi 1 / 220nm,4nm

Integration Result

Peak Table

PDA Ch1 220nm

| Peak# | Ret. Time | Height | Height% | Resolution(USP) | Area | Area% |
|---|---|---|---|---|---|---|
| 1 | 0.913 | 59793 | 36.903 | -- | 274652 | 23.938 |
| 2 | 1.684 | 102237 | 63.097 | 4.400 | 872691 | 76.062 |

Method details

Column: Chiralpak AD-3 50×4.6mm I.D., 3 μm
Mobile phase: A: $CO_2$, B: 2:1 i-PrOH/acetonitrile (0.05% diethylamine)
Elution: 60% B

WX-74b (WX-02-693)

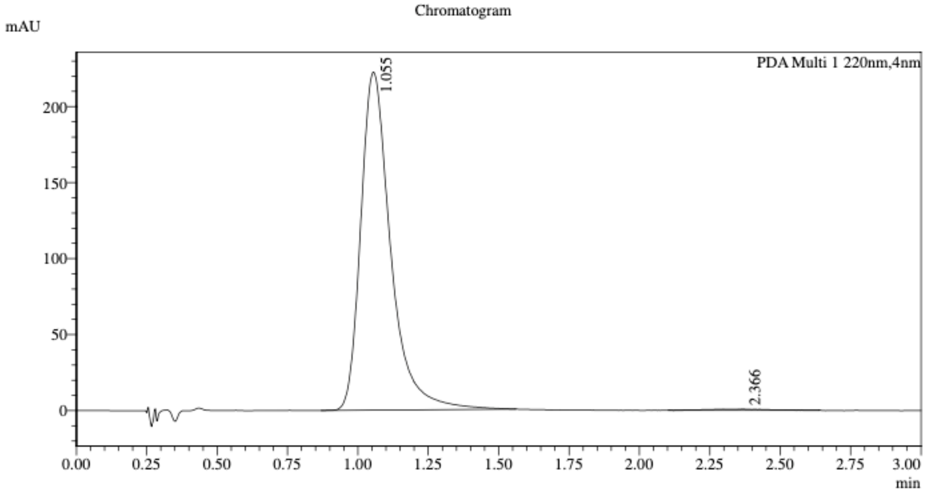


1 PDA Multi 1 / 220nm,4nm

Integration Result

Peak Table

PDA Ch1 220nm

| Peak# | Ret. Time | Height | Height% | Resolution(USP) | Area | Area% |
|---|---|---|---|---|---|---|
| 1 | 1.055 | 221931 | 99.665 | -- | 1644696 | 99.312 |
| 2 | 2.366 | 746 | 0.335 | 4.330 | 11399 | 0.688 |

WX-74d (WX-02-694)

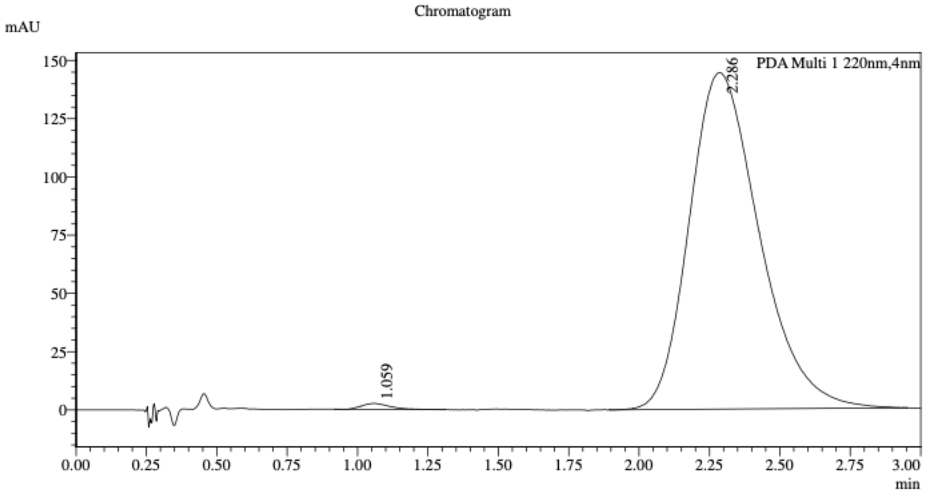


1 PDA Multi 1 / 220nm,4nm

Integration Result

Peak Table

PDA Ch1 220nm

| Peak# | Ret. Time | Height | Height% | Resolution(USP) | Area | Area% |
|---|---|---|---|---|---|---|
| 1 | 1.059 | 2537 | 1.726 | -- | 17374 | 0.690 |
| 2 | 2.286 | 144395 | 98.274 | 3.888 | 2500672 | 99.310 |

Mixture of WX-74b (WX-02-693) & WX-74d (WX-02-694)

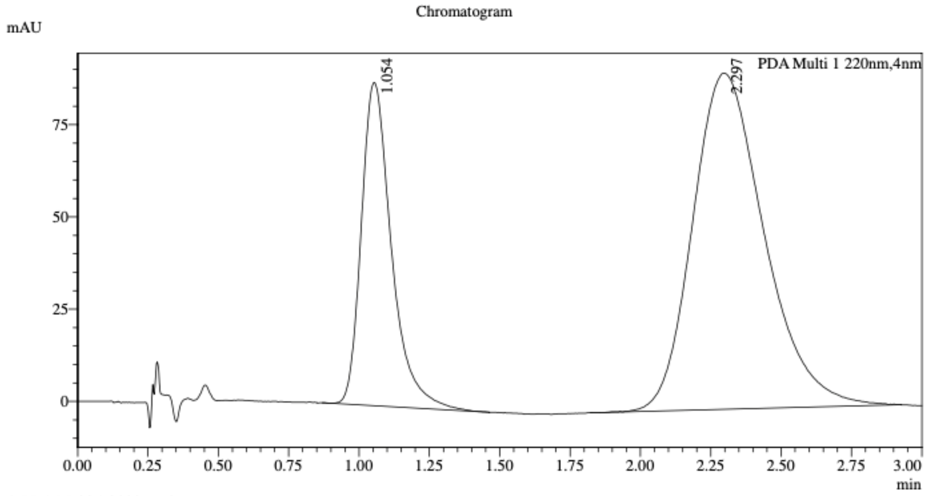


1 PDA Multi 1 / 220nm,4nm

Integration Result

Peak Table

PDA Ch1 220nm

| Peak# | Ret. Time | Height | Height% | Resolution(USP) | Area | Area% |
|---|---|---|---|---|---|---|
| 1 | 1.054 | 87529 | 49.016 | -- | 652454 | 28.817 |
| 2 | 2.297 | 91042 | 50.984 | 3.841 | 1611714 | 71.183 |

Method details

Column: Chiralpak IC-3 50×4.6mm I.D., 3 μm
Mobile phase: A: $CO_2$, B: EtOH (0.05% diethylamine)
Elution: 50% B

WX-72a-yne (WX-02-891)

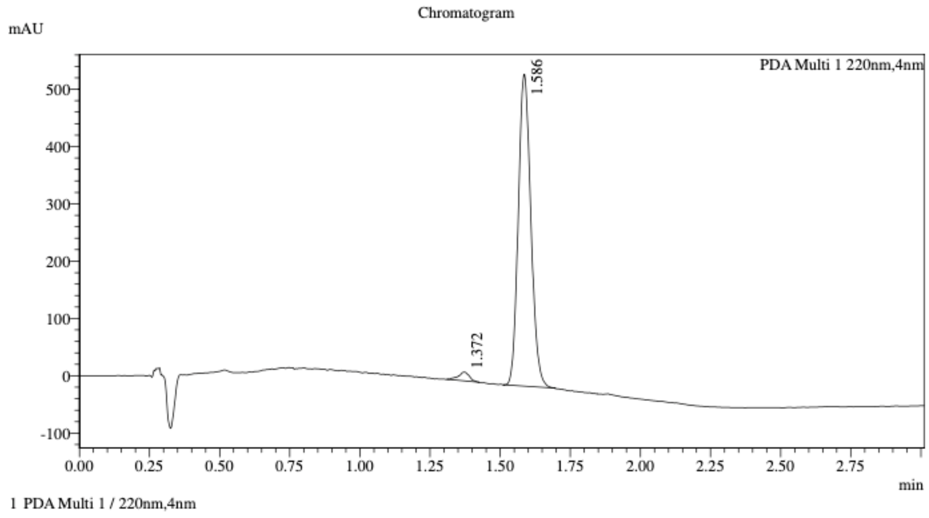


Integration Result

Peak Table

PDA Ch1 220nm

| Peak# | Ret. Time | Height | Height% | Resolution(USP) | Area | Area% |
|---|---|---|---|---|---|---|
| 1 | 1.372 | 15002 | 2.737 | -- | 40525 | 2.334 |
| 2 | 1.586 | 533093 | 97.263 | 2.758 | 1695899 | 97.666 |

WX-72c-yne (WX-02-892)

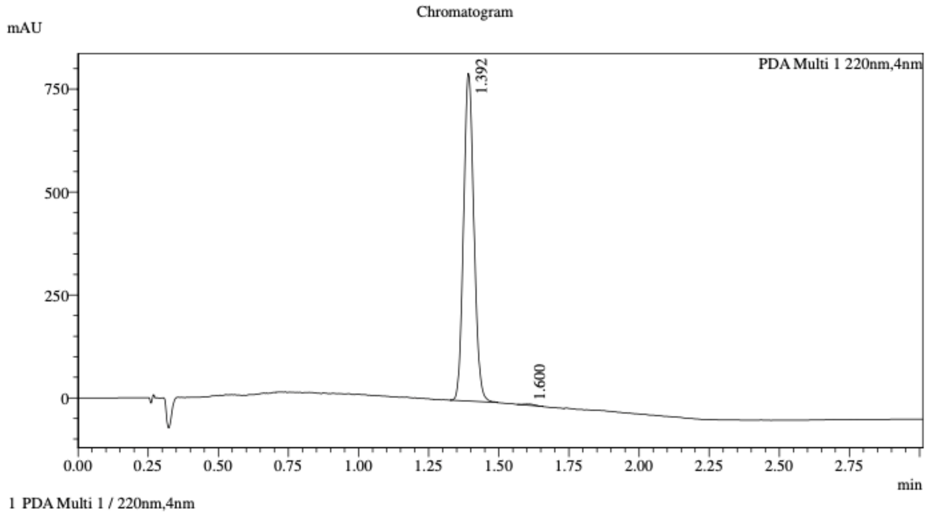


Integration Result

Peak Table

PDA Ch1 220nm

| Peak# | Ret. Time | Height | Height% | Resolution(USP) | Area | Area% |
|---|---|---|---|---|---|---|
| 1 | 1.392 | 779110 | 99.658 | -- | 2039140 | 99.631 |
| 2 | 1.600 | 2676 | 0.342 | 2.859 | 7562 | 0.369 |

Mixture of WX-72a-yne (WX-02-891) & WX-72c-yne (WX-02-892)

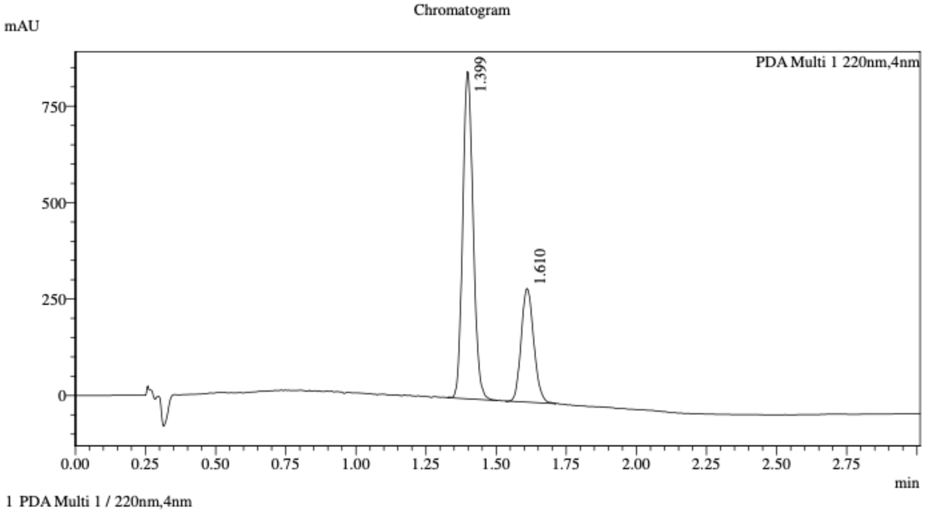


Integration Result

Peak Table

PDA Ch1 220nm

| Peak# | Ret. Time | Height | Height% | Resolution(USP) | Area | Area% |
|---|---|---|---|---|---|---|
| 1 | 1.399 | 840682 | 74.336 | -- | 2192359 | 70.415 |
| 2 | 1.610 | 290234 | 25.664 | 2.723 | 921140 | 29.585 |

Method details

Column: Chiralpak IC-3 50×4.6mm I.D., 3 µm
Mobile phase: A: $CO_2$, B: 2:1 i-PrOH/acetonitrile (0.05% diethylamine)
Elution: 20% to 60% B (gradient)

WX-72b-yne (WX-02-893)

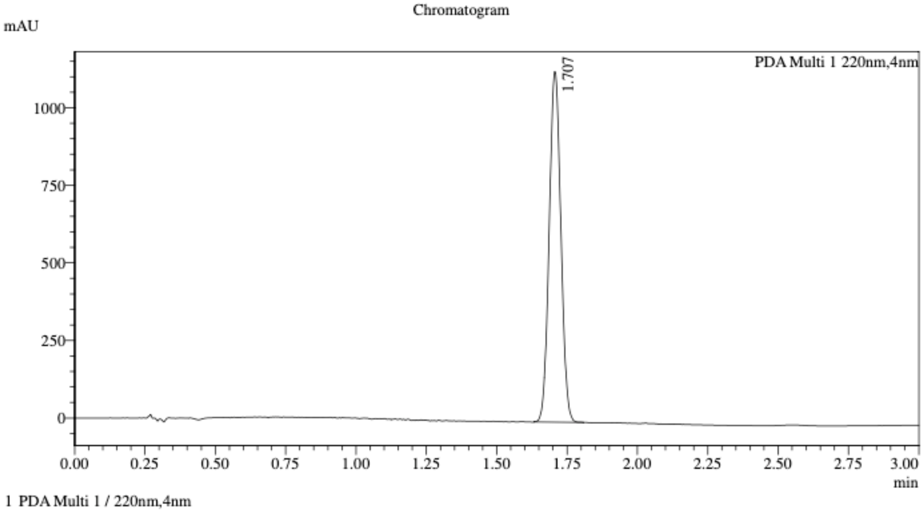


Integration Result

Peak Table

PDA Ch1 220nm

| Peak# | Ret. Time | Height | Height% | Resolution(USP) | Area | Area% |
|---|---|---|---|---|---|---|
| 1 | 1.707 | 1114213 | 100.000 | -- | 3252984 | 100.000 |

WX-72d-yne (WX-02-894)

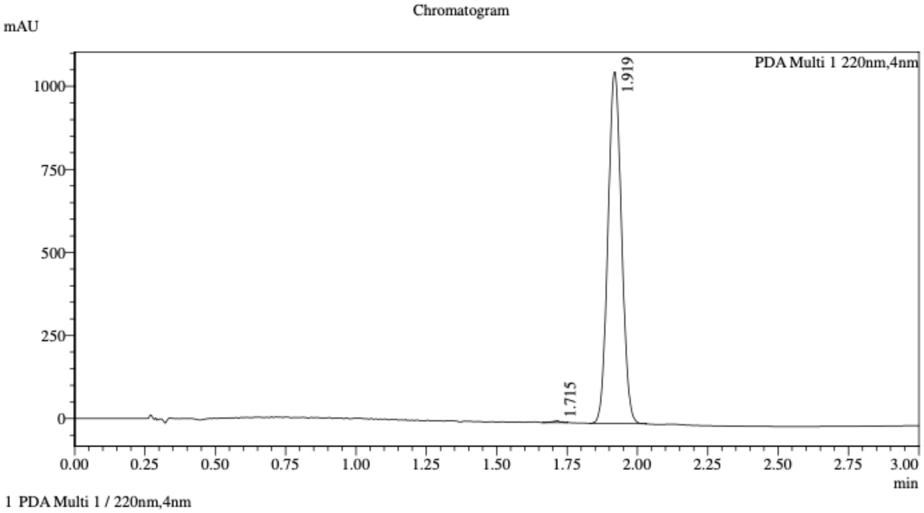


Integration Result

Peak Table

PDA Ch1 220nm

| Peak# | Ret. Time | Height | Height% | Resolution(USP) | Area | Area% |
|---|---|---|---|---|---|---|
| 1 | 1.715 | 4991 | 0.473 | -- | 9158 | 0.258 |
| 2 | 1.919 | 1050820 | 99.527 | 2.783 | 3543260 | 99.742 |

Mixture of WX-72b-yne (WX-02-893) & WX-72d-yne (WX-02-894)

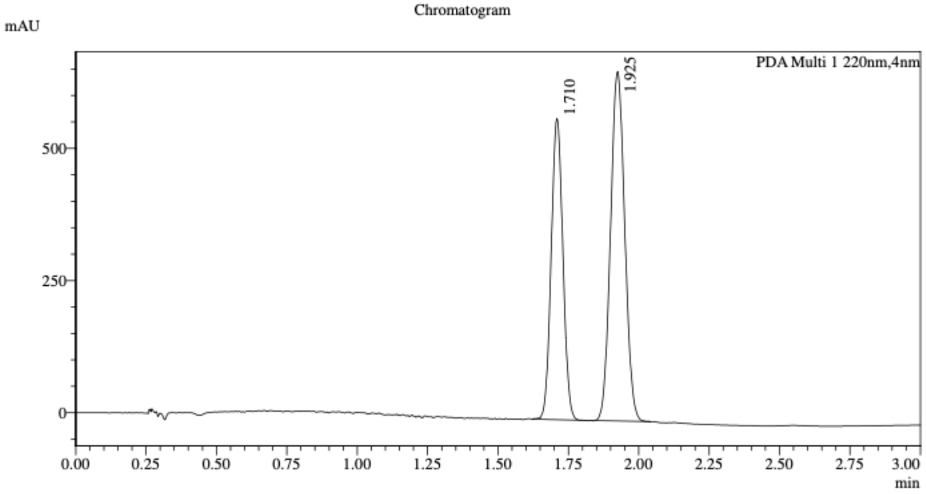


1 PDA Multi 1 / 220nm,4nm

Integration Result

Peak Table

PDA Ch1 220nm

| Peak# | Ret. Time | Height | Height% | Resolution(USP) | Area | Area% |
|---|---|---|---|---|---|---|
| 1 | 1.710 | 565852 | 46.239 | -- | 1644694 | 41.505 |
| 2 | 1.925 | 657900 | 53.761 | 2.455 | 2317973 | 58.495 |

Method details

Column: Chiralpak IK-3 50×4.6mm I.D., 3 μm
Mobile phase: A: $CO_2$, B: EtOH (0.05% diethylamine)
Elution: 5% to 25% B (gradient)

WX-73a-yne (WX-02-905)

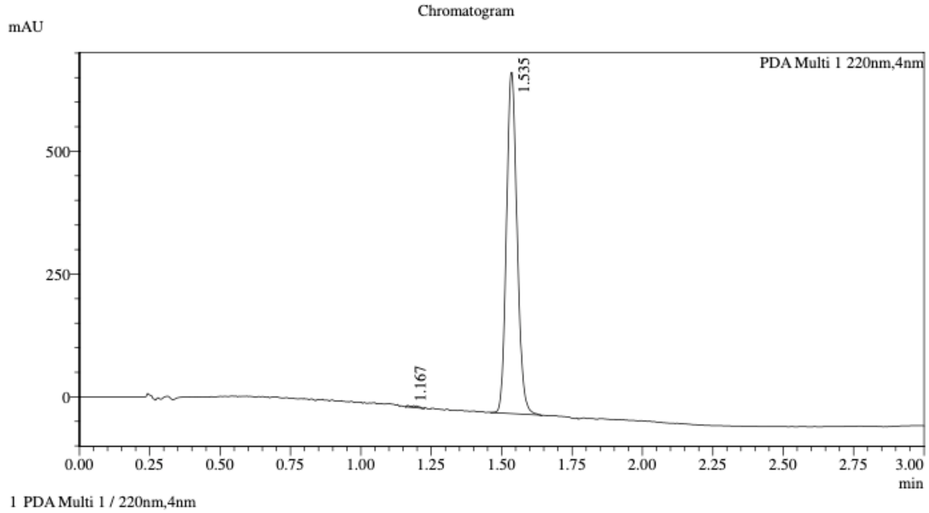


Integration Result

Peak Table

PDA Ch1 220nm

| Peak# | Ret. Time | Height | Height% | Resolution(USP) | Area | Area% |
|---|---|---|---|---|---|---|
| 1 | 1.167 | 2477 | 0.359 | -- | 7702 | 0.421 |
| 2 | 1.535 | 687615 | 99.641 | 5.564 | 1823726 | 99.579 |

WX-73c-yne (WX-02-906)

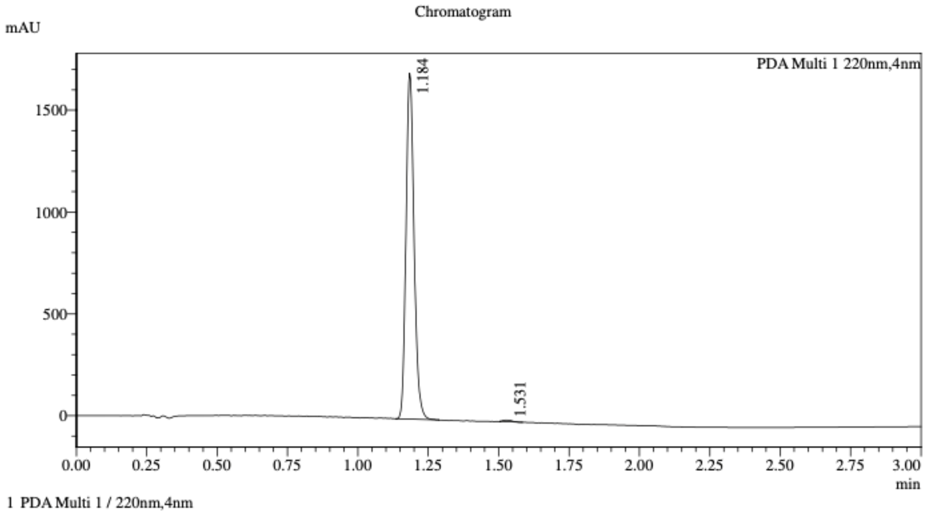


Integration Result

Peak Table

PDA Ch1 220nm

| Peak# | Ret. Time | Height | Height% | Resolution(USP) | Area | Area% |
|---|---|---|---|---|---|---|
| 1 | 1.184 | 1629165 | 99.540 | -- | 3379855 | 99.531 |
| 2 | 1.531 | 7532 | 0.460 | 6.099 | 15931 | 0.469 |

Mixture of WX-73a-yne (WX-02-905) & WX-73c-yne (WX-02-906)

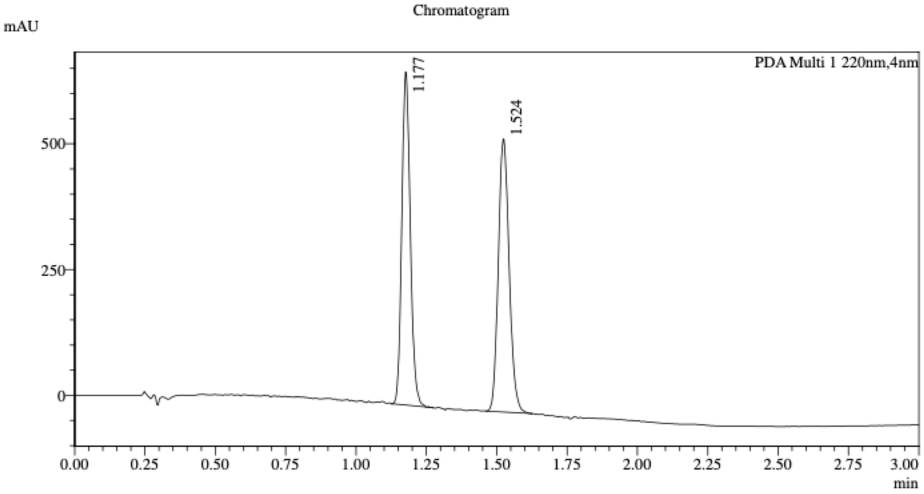


1 PDA Multi 1 / 220nm,4nm

Integration Result

Peak Table

PDA Ch1 220nm

| Peak# | Ret. Time | Height | Height% | Resolution(USP) | Area | Area% |
|---|---|---|---|---|---|---|
| 1 | 1.177 | 629895 | 53.942 | -- | 1387477 | 48.820 |
| 2 | 1.524 | 537823 | 46.058 | 5.242 | 1454530 | 51.180 |

Method details

Column: Chiralcel OD-3 50×4.6mm I.D., 3 µm
Mobile phase: A: $CO_2$, B: EtOH (0.05% diethylamine)
Elution: 20% to 60% B (gradient)

WX-73b-yne (WX-02-907)

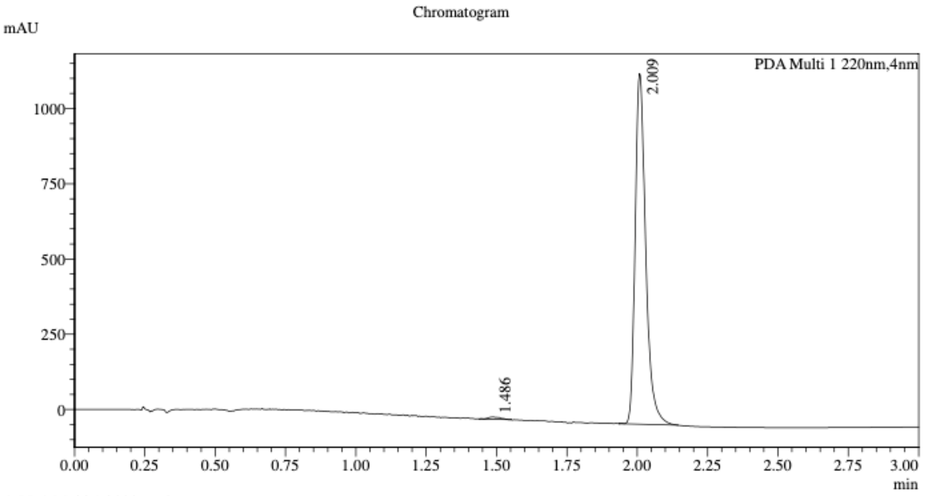


1 PDA Multi 1 / 220nm,4nm

Integration Result

Peak Table

PDA Ch1 220nm

| Peak# | Ret. Time | Height | Height% | Resolution(USP) | Area | Area% |
|---|---|---|---|---|---|---|
| 1 | 1.486 | 6938 | 0.612 | -- | 21362 | 0.700 |
| 2 | 2.009 | 1127094 | 99.388 | 7.051 | 3032079 | 99.300 |

WX-73d-yne (WX-02-908)

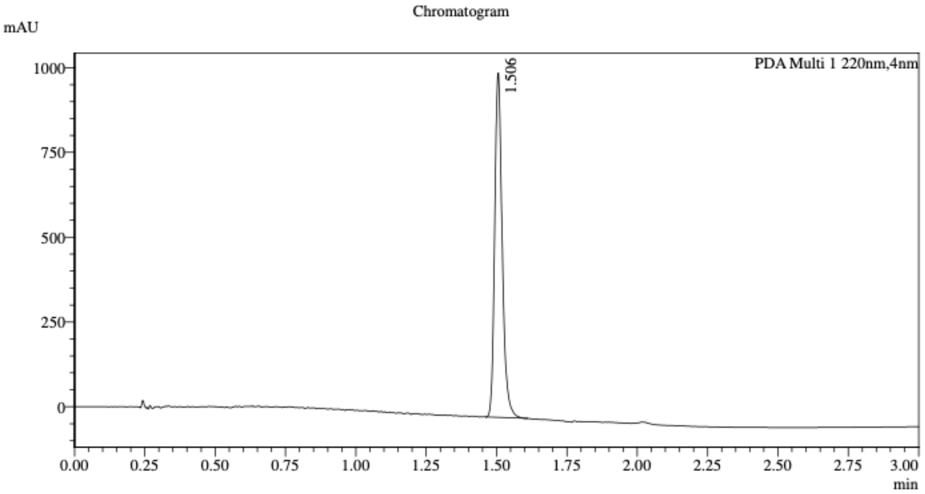


1 PDA Multi 1 / 220nm,4nm

Integration Result

Peak Table

PDA Ch1 220nm

| Peak# | Ret. Time | Height | Height% | Resolution(USP) | Area | Area% |
|---|---|---|---|---|---|---|
| 1 | 1.506 | 954304 | 100.000 | -- | 1887226 | 100.000 |

Mixture of WX-73b-yne (WX-02-907) & WX-73d-yne (WX-02-908)

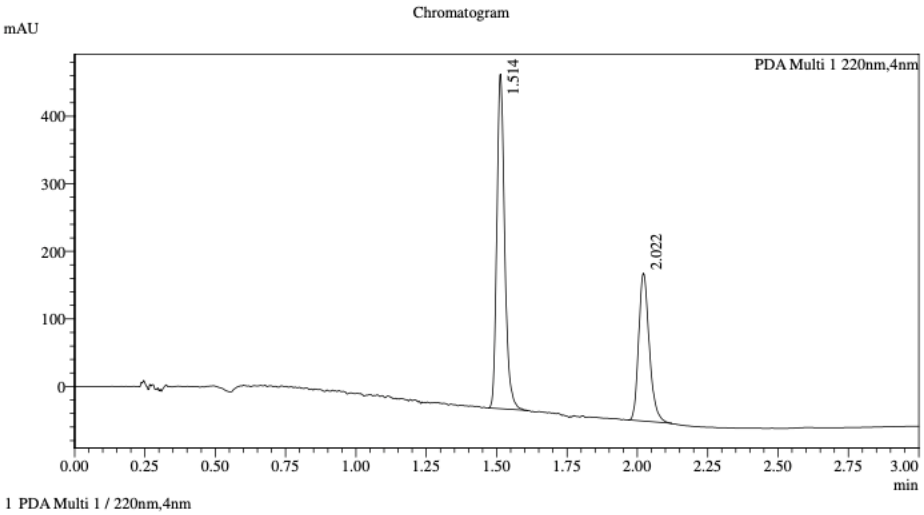


Integration Result

Peak Table

PDA Ch1 220nm

| Peak# | Ret. Time | Height | Height% | Resolution(USP) | Area | Area% |
|---|---|---|---|---|---|---|
| 1 | 1.514 | 466960 | 68.286 | -- | 958785 | 62.552 |
| 2 | 2.022 | 216874 | 31.714 | 8.138 | 574001 | 37.448 |

Method details

Column: (*S*,*S*)Whelk-O1 50×4.6mm I.D., 3.5 μm

Mobile phase: A: $CO_2$, B: EtOH (0.05% diethylamine)

Elution: 20% to 60% B (gradient)

WX-62a-yne (WX-02-909)

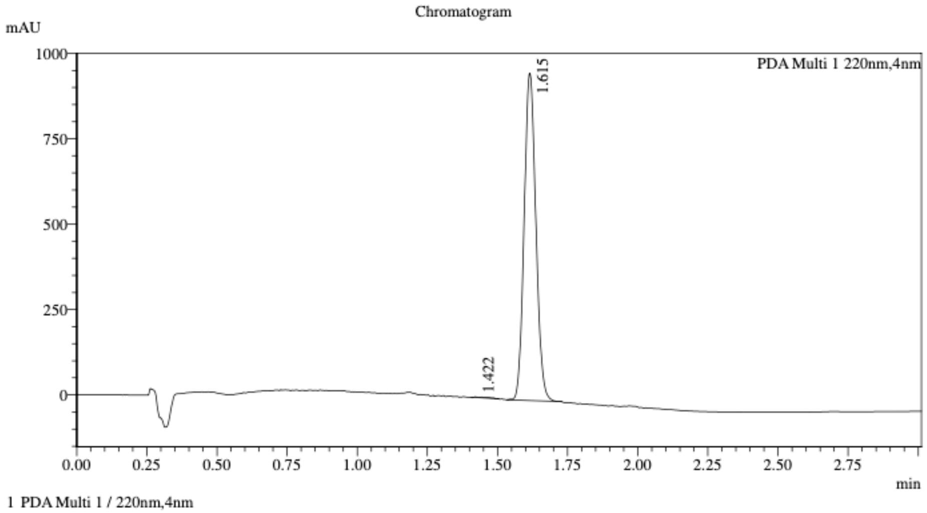


Integration Result

Peak Table

PDA Ch1 220nm

| Peak# | Ret. Time | Height | Height% | Resolution(USP) | Area | Area% |
|---|---|---|---|---|---|---|
| 1 | 1.422 | 1813 | 0.191 | -- | 6683 | 0.240 |
| 2 | 1.615 | 949244 | 99.809 | 0.215 | 2779459 | 99.760 |

WX-62c-yne (WX-02-910)

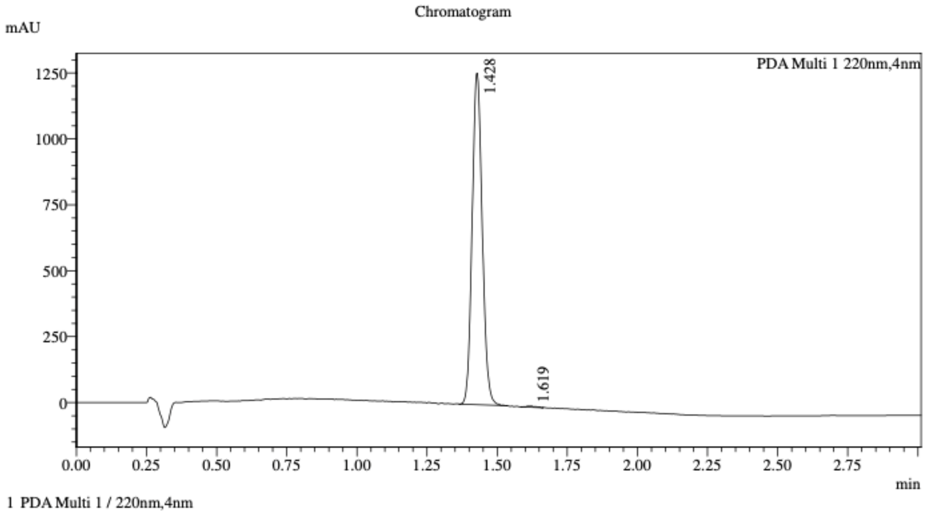


Integration Result

Peak Table

PDA Ch1 220nm

| Peak# | Ret. Time | Height | Height% | Resolution(USP) | Area | Area% |
|---|---|---|---|---|---|---|
| 1 | 1.428 | 1245428 | 99.659 | -- | 3145737 | 99.692 |
| 2 | 1.619 | 4261 | 0.341 | 3.075 | 9731 | 0.308 |

Mixture of WX-62a-yne (WX-02-909) & WX-62c-yne (WX-02-910)

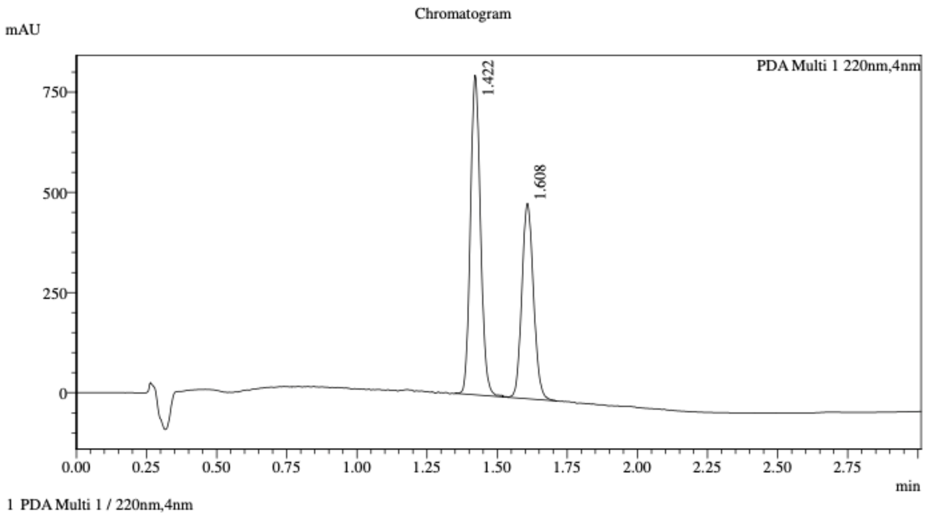


Integration Result

Peak Table

PDA Ch1 220nm

| Peak# | Ret. Time | Height | Height% | Resolution(USP) | Area | Area% |
|---|---|---|---|---|---|---|
| 1 | 1.422 | 790108 | 62.199 | -- | 1995020 | 58.609 |
| 2 | 1.608 | 480186 | 37.801 | 2.516 | 1408908 | 41.391 |

Method details

Column: Chiralpak IC-3 50×4.6mm I.D., 3 µm
Mobile phase: A: $CO_2$, B: 2:1 i-PrOH/acetonitrile (0.05% diethylamine)
Elution: 20% to 60% B (gradient)

WX-62b-yne (WX-02-911)

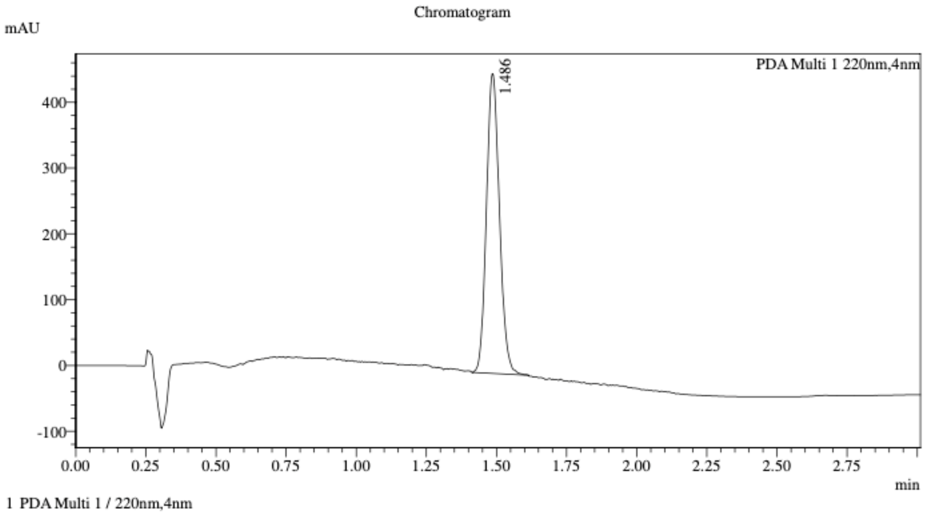


Integration Result

Peak Table

PDA Ch1 220nm

| Peak# | Ret. Time | Height | Height% | Resolution(USP) | Area | Area% |
|---|---|---|---|---|---|---|
| 1 | 1.486 | 453263 | 100.000 | -- | 1470920 | 100.000 |

WX-62d-yne (WX-02-912)

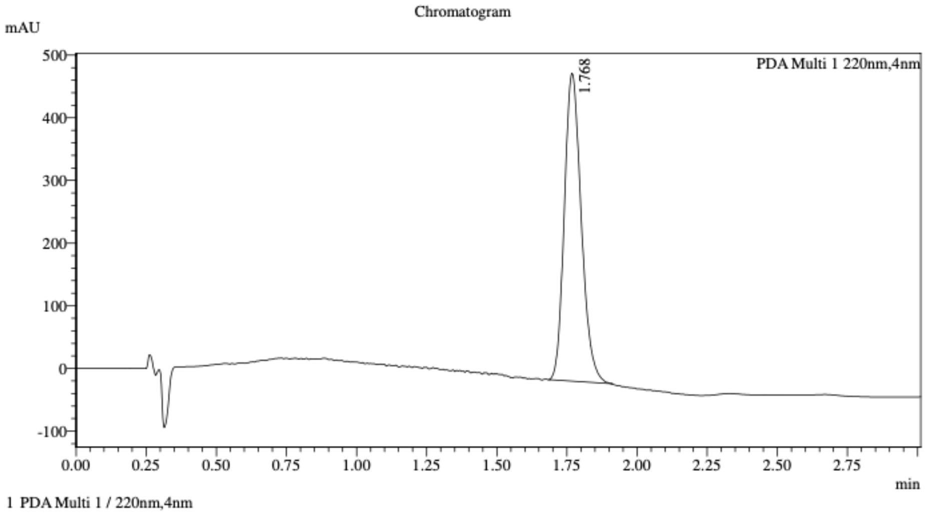


Integration Result

Peak Table

PDA Ch1 220nm

| Peak# | Ret. Time | Height | Height% | Resolution(USP) | Area | Area% |
|---|---|---|---|---|---|---|
| 1 | 1.768 | 486797 | 100.000 | -- | 2063167 | 100.000 |

Mixture of WX-62b-yne (WX-02-911) & WX-62d-yne (WX-02-912)

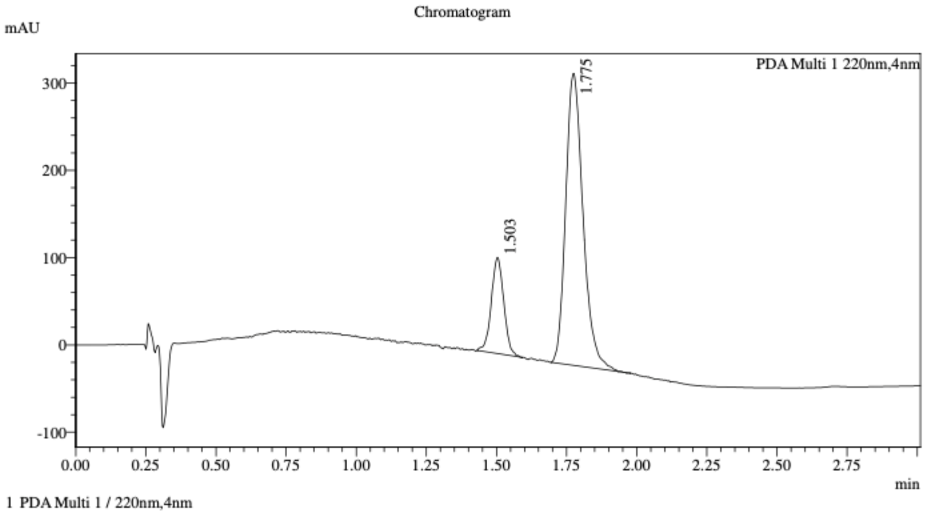


Integration Result

Peak Table

PDA Ch1 220nm

| Peak# | Ret. Time | Height | Height% | Resolution(USP) | Area | Area% |
|---|---|---|---|---|---|---|
| 1 | 1.503 | 108500 | 24.568 | -- | 340187 | 19.152 |
| 2 | 1.775 | 333138 | 75.432 | 2.753 | 1436015 | 80.848 |

Method details

Column: Chiralpak IC-3 50×4.6mm I.D., 3 μm
Mobile phase: A: $CO_2$, B: 2:1 i-PrOH/acetonitrile (0.05% diethylamine)
Elution: 20% to 60% B (gradient)
