## Supplementary Tables for "Tryptoline Stereoprobe Elaboration Identifies Inhibitors of the GRPEL1-HSPA9 Chaperone Complex"

**Supplementary Table 1:** List of reagents not mentioned in the method section, their sources and catalog numbers.

| **Reagent** | **Source** | **Catalog Number** |
|---|---|---|
| Desthiobiotin polyethyleneoxide iodoacetamide (IA-DTB) | Santa Cruz Biotechnology | sc-300424 |
| Biotin-PEG4-azide | ChemPep | 271606 |
| EZ-Link™ NHS-Desthiobiotin | ThermoFisher Scientific | 16129 |
| Urea | Fisher Scientific | M1084871000 |
| Iodoacetamide | Sigma-Aldrich | I1149-25G |
| Dithiothreitol (DTT) | Fisher Bioreagents | BP172-25 |
| Sequencing Grade Modified Trypsin | Promega | V5111 |
| Streptavidin Agarose Resin | Fisher Scientific | 20349 |
| Micro Bio-Spin column | Bio-Rad | 7326204 |
| NonidetTM P40 substitute (IgepalTM CA-630) | USB Corporation | 19628 |
| Triethylammonium bicarbonate buffer | Sigma-Aldrich | T7408-500ML |
| TMT10plex™ Isobaric Label Reagent Set | ThermoFisher Scientific | 90406 |
| TMTpro™ 16plex Label Reagent Set | ThermoFisher Scientific | A44520 |
| Ammonium bicarbonate | Sigma-Aldrich | 09830-500G |
| Hydroxylamine solution | Sigma-Aldrich | 467804-10ML |
| Formic acid, ~98%, for mass spectrometry | Honeywell Fluka | 94318-250ML-F |
| Methanol | Fisher Scientific | A452SK-4 |
| Acetonitrile LC-MS Grade | Fisher Scientific | A955 |
| Water LC-MS Grade | Fisher Scientific | W6-4 |
| Protein LoBind Tubes 1.5 mL | Fisher Scientific | E925000090 |
| EPPS | Millipore | E0276 |
| Chloroform | Fisher Scientific | C297-4 |
| Sep-Pak Vac 1cc C18 cartridges | Waters | WAT054955 |
| Sodium Dodecyl Sulfate (SDS) Biotechnolgy grade, 1kg | BioPioneer | C0065 |
| S-(5'-Adenosyl)-L-methionine iodide | Sigma-Aldrich | A4377-25MG |
| S-(5'-Adenosyl)-L-homocysteine | Sigma-Aldrich | A9384-25MG |
| Glycerol | Fisher Scientific | BP229-4 |
| Benzonase | Sigma-Aldrich | 1016950001 |
| Lys-C | Cell Signaling Technologies | 39003S |
| Tris(benzyltriazolylmethyl)amine (TBTA) | TCI | T2993 |
| Tris(2-carboxyethyl)phosphine HCl (TCEP) | Millipore | 75259 |
| Cupric sulfate, pentahydrate | MP Biomedicals | 195117 |
| Pierce Anti-DYKDDDDK Magnetic Agarose | Fisher Scientific | PIA36798 |
| V5-Trap® Magnetic Agarose (v5tma 200rxns) | Protein Tech | v5tma-200 |
| DMEM for SILAC | Life Technologies | 88364 |

| | | |
|---|---|---|
| Fetal Bovine Serum, Dialyzed | Omega Scientific | FB-03 |
| L-Arginine-$^{13}C_6$,$^{15}N_4$ hydrochloride | Sigma-Aldrich | 608033 |
| L-Lysine-$^{13}C_6$,$^{15}N_2$ hydrochloride | Sigma-Aldrich | 608041 |
| L-Arginine monohydrochloride | Sigma-Aldrich | A6969 |
| L-Lysine dihydrochloride | Sigma-Aldrich | L5751 |
| OPTI-MEM | ThermoFisher Scientific | 31985062 |
| Amicon® Ultra Centrifugal Filter, 30 kDa MWCO | Sigma-Aldrich | UFC503024 |
| Zeba Spin Desalting Columns, 40K MWCO, 0.5 mL, 25 Columns | Life Technologies | A57759 |
| 35 mm Dish \| No. 1.5 Coverslip \| 14 mm Glass Diameter \| Poly-D-Lysine Coated | MatTek | P35GC-1.5-14-C |
| MitoTracker Deep Red FM | Cell Signaling Technologies | 8778 |
| Dulbecco′s Modified Eagle′s Medium | Sigma-Aldrich | D5030 |
| Seahorse XFe96/XF Pro FluxPak Mini | Agilent | 103793-100 |
| Oligomycin | Sigma-Aldrich | 495455 |
| FCCP | Sigma-Aldrich | C2920 |
| Rotenone | Sigma-Aldrich | R8875 |
| Antimycin A | Sigma-Aldrich | A8674 |
| ATF-4 (D4B8) Rabbit mAb | Cell Signaling Technologies | 11815 |
| Doxycycline hyclate cell-culture grade | Sigma-Aldrich | D5207-1G |
| CHOP (L63F7) Mouse mAb | Cell Signaling Technologies | 2895S |
| β-Actin (8H10D10) Mouse mAb | Cell Signaling Technologies | 3700S |
| Monoclonal ANTI-FLAG(R) M2 antibody produced in mouse | Sigma-Aldrich | F1804-1MG |
| Anti-rabbit IgG, HRP-linked Antibody | Cell Signaling Technologies | 7074S |
| anti-mouse IgG, HRP-linked antibody | Cell Signaling Technologies | 7076S |
| GRPEL1 Polyclonal antibody | Protein Tech | 12720-1-AP |
| Gateway BP Clonase II Enzyme mix | Life Technologies | 11791020 |
| Gateway LR Clonase II Enzyme mix | Life Technologies | 11791020 |
| V5 Tag Monoclonal Antibody (SV5-Pk1) | Life Technologies | R96025 |
| Puromycin Dihydrochloride | Gibco | A1113803 |
| Blasticidin, 100mg (10 x 1mL tubes) | invivogen | ant-bl-1 |
| G418, 2 g (20 x 1 ml tubes) | invivogen | ant-gn-2 |
| Mitochondria Isolation Kit for Cultured Cells | Fisher Scientific | 89874 |
| NEB 5-alpha Competent E. coli (High Efficiency) | New England Biolabs | C2987H |
| One Shot Stbl3 Chemically Competent E. coli | Life Technologies | C737303 |
| DMSO | Corning | 25-950-CQC |

**Supplementary Table 2:** List of target genes and sources.

| Gene | Source | Catalog # |
|---|---|---|
| TEX264 | GenScript | OHu00477 |
| NCKAP1L | GenScript | OHu31086 |
| GRPEL1 | OriGene | RC206086 |

**Supplementary Table 3:** List of primers used for cloning.

| Name | Forward Primer | Reverse Primer |
|---|---|---|
| GRPEL1_C124A | GGCAACACAGgctGTTCCAAAAGAAGAAATTAAAG | TTCTCCAGAACGTCTGCC |
| sgGRPEL1_05 | caccgAGCTGAAGGAGACTGTGGTG | aaacCACCACAGTCTCCTTCAGCTc |
| NCKAP1L_C338A | CCAGTTTCATgctCAACGGCGGC | CCACTGTTTGCAATTAC |
| NCKAP1L_gateway | GGGGACAAGTTTGTACAAAAAAGCAGGCTTAatgtctttgacatctgcttaccagc | GGGGACCACTTTGTACAAGAAAGCTGGGTTtcacttatcgtcgtcatccttgtaatcgtt |
| TEX264_C165A | GCGGAAGCTGgctGCCTATCCTC | TCCTTGATGTAGGTGTC |
| TEX264_C68A | CACTGAGAGCgccAGCATCTCTC | AAAAGCCGCCCAGTC |
